# Complexity of Guanine Quadruplex Unfolding Pathways Revealed by Atomistic Pulling Simulations

**DOI:** 10.1101/2023.01.27.525972

**Authors:** Petr Stadlbauer, Vojtěch Mlýnský, Miroslav Krepl, Jiří Šponer

## Abstract

Guanine quadruplexes (GQs) are non-canonical nucleic acid structures involved in many biological processes. GQs formed in single-stranded regions often need to be unwound by cellular machinery, so their mechanochemical properties are important. Here, we performed steered molecular dynamics simulations of human telomeric GQs to study their unfolding. We examined four pulling regimes, including very slow setup with pulling velocity and force load accessible to high-speed atomic force microscopy. We identified multiple factors affecting the unfolding mechanism. The more the direction of force was perpendicular to the GQ channel axis (determined by GQ topology), the more the base unzipping mechanism happened. If the GQ had either all-*anti* or all-*syn* pattern in a strand, strand slippage mechanism was more likely to occur. Importantly, slower pulling velocity led to richer unfolding pathways including partial refolding attempts. We show that GQ may eventually unfold after force drop under forces smaller than those the GQ withstood before the drop. This suggests that proteins *in vivo* might resolve GQs even if their stall forces are smaller than GQ rupture force. Finally, we found out that different unfolding intermediates may have very similar chain end-to-end distance, which reveals some limitations of structural interpretations of single-molecule spectroscopic data.

## INTRODUCTION

Guanine quadruplexes (GQ) are a common non-canonical form of nucleic acids. They are formed by sequences rich in guanine (G), which are widespread – hundreds of thousands known and putative sites have been identified in the human genome (1,2). Notably, GQs are abundant in genes regulatory parts and in the telomeric region, where they play roles in maintaining cell function and genome integrity (3–7). Mutations leading to over- or under-formation of G4s often lead to diseases, such as cancer (8–12) or neuropathologies (13–15).

A basic structural unit of GQ is a G-quartet (quartet). It is composed of four Gs bound in a cyclical arrangement, where each G is connected to two neighboring Gs by *cis*-Watson-Crick-Hoogsteen (*c*WH) H-bonding (Figure 1) (16). GQ is formed by stacking of quartets together. A channel, whose walls are covered by carbonyl O6 atoms, runs through GQ center; positively charged cations (e.g., K^+^ or Na^+^) need to be bound in the channel for the GQ to be stable. Intramolecular GQs have loops, which are nucleotide segments connecting the G-strands (G-stretches, columns) (Figure 1). Loops connecting two neighboring G-strands running in the same direction, i.e. parallel, are called propeller; lateral loops connect two neighboring antiparallel strands, while diagonal loops connect two antiparallel strands across the GQ. G-strand directionality is interdependent with the *syn/anti* conformation of the Gs’ glycosidic torsion angle *χ* (17–19). If two Gs in a quartet are in mutually parallel strands, they have the same *χ* conformation (i.e., either both *anti* or both *syn*). If they are in antiparallel strands, they have opposite *χ* conformation (i.e., one is *anti* and the other is *syn*). Same *χ* conformation of Gs’ in a given G-strand means that the two quartets containing those two Gs’ are of the same quartet directionality, i.e. both are clockwise or counterclockwise, while opposite *χ* conformation leads to opposite quartet directionality. These rules have implications in structural polymorphism and folding pathways of GQs (20). Human telomeric sequence d(GGGTTA)_n_ is a prime example of such a highly polymorphic sequence, forming at least six known stable GQ conformations (21–29).

**Figure 1.**
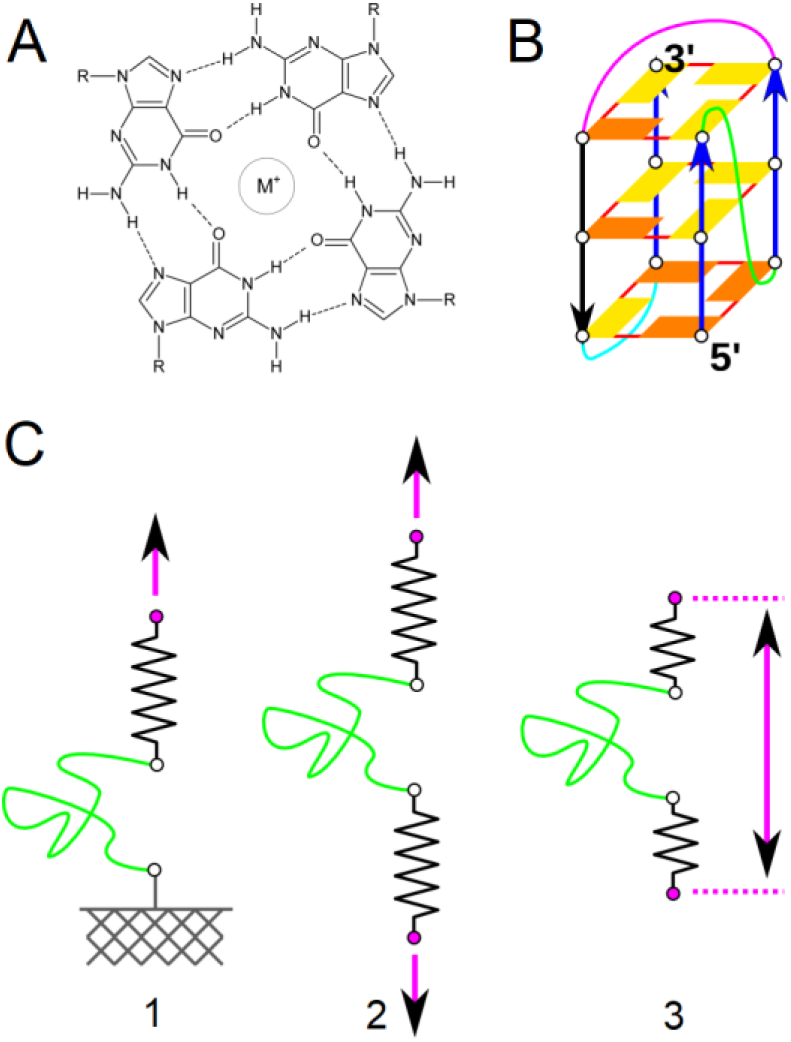
A) G-quartet with a metal cation in the central cavity. “R” represents the sugar-phosphate moiety. B) Illustrative GQ model. *Anti* guanines, i.e., G residues with *anti* orientation of *χ* dihedral angle, are displayed as yellow rectangles, while *syn* guanines (Gs with *syn* conformation of *χ* dihedrals) are shown as orange rectangles. The blue arrows represent strands that are mutually parallel, while the black arrow highlights the remaining strand, which is in antiparallel orientation to the others. Propeller loop is shown as green line, diagonal loop as purple line and lateral loop as cyan line. Solid red lines indicate *cis* Watson-Crick Hoogsteen (*c*WH) base pairing. The bottom quartet is of opposite directionality than the middle and top one. The quartet including G closest to the 5’-end, i.e. the bottom one in this example, is called ‘first’; the middle one is ‘second’ and the quartet on the other side of G-stem is ‘third’ in this paper. Nucleotides in the loops and channel cations are not shown for clarity. C) Illustrative scheme of common force-induced unfolding simulation techniques: (1) One selected point (white circle) of a molecule (green line) is connected by a virtual spring to a virtual particle (purple circle) that is moving in a predefined direction and one point is fixed in space (marked as a connection to the grey wall). (2) Instead of a fixed point, there is another connection by the second spring to a particle moving in the opposite direction. (3) Two points connected to a molecule are moving away from each other, but not in a predefined direction, only their mutual distance is time-dependent. The springs are not independent, but behave as if being just one, i.e., potential energy added by the spring depends on molecule’s end-to-end distance as in (1) but without a fixed point. Pulling protocols used in this work employ the implementation displayed as option (3).

Recent studies have suggested that, globally, GQ folding is a complicated process, generally best described by kinetic partitioning (30–33). We have suggested that the idealized folding process of a GQ ensemble can be divided into two stages (20,34). The first stage is characterized by fast folding of the initial ensemble into various mostly misfolded GQs with non-native *syn/anti* combination and/or reduced number of quartets in comparison to fully folded GQs. Second stage of the folding process is slow refining of the initial GQ population into final (native) GQs population. This stage is slow because it requires partial or complete unfolding of the misfolded GQs, followed by random *syn-anti* flips and then new folding attempts, which may or may not lead to native GQ. Structural transitions between the misfolded and native GQs happen via various ensembles of other structures. For example, depending on target GQ topology, incomplete or perturbed GQs (35–40), G-triplexes (38–51), G-hairpins (36,38,39,41,43,45,52) or cross-like species (34,53–55) have been hypothesized to participate in the process. The exact nature of these ensembles depends on external factors, such as temperature, ionic strength, presence of cosolvents, ligands, specific cations or other nearby structural elements (20).

A way to affect the free-energy landscape area that the GQ folding sequence traverses is to put the molecule under tension. The external force typically pulling the sequence ends away from each other results in exploration of a lower-entropy section of its free-energy surface. External forces acting on DNA are actually common in biological systems and many of its functions are related to its mechanochemical properties (56,57). *In vivo*, GQ forming sequences are not isolated, but are a part of long DNA, which by itself may exert a tug. Cellular machinery causes tension in GQ structures as well, e.g., by superhelical coiling or an effect of specialized enzymes – helicases – whose role is often to resolve GQs; their probably act by moving along a DNA strand, exerting force on the GQ until it eventually unfolds (6,56–60). Force-induced unfolding can be studied in *in vitro* by single-molecule force spectroscopy techniques, such as magnetic, optical and nanopore tweezers or atomic force microscopy (AFM). These have been applied to investigate the conformational and mechanical behavior of biopolymers and their interactions and they can operate using various force-exertion regimes, typically: i) constant velocity, ii) force ramp, iii) force clamp, iv) pulse chase, and v) zig-zag force ramp (61–64). Usually long handles/linkers are connected to the studied biopolymer, one handle is fixed and the other is connected to a moving surface, which is moved towards and away from the fixed end, and the tension is transferred through the linker to the studied biomolecule.

Human telomeric sequence GQs have indeed been widely studied by force spectroscopy techniques and valuable insights have been obtained. It has been shown that the conformations formed in Na^+^ (presumably the basket topology) are mechanically less stable than the topologies folded in K^+^ (3+1 hybrid topology) (e.g., by rupture forces of ~10 pN vs. ~20 pN at loading rates of ~2-5 pN/s) (65–69). The conformational polymorphism of the telomeric sequence in presence of K^+^ has been demonstrated, too, with a detection of a few distinct states (66,70), later extended to at least six distinct states (67). Surprisingly, the authors of the latter study also found out that having more than four repeats of the telomeric sequence actually decreases the conformational diversity (71). A study focused on the effect of pyridostatin has revealed that the ligand promotes folding of the human telomeric GQ (65). Non-telomeric GQs have been investigated, too. It has been shown that GQs with bulges and decreased number of quartets are less stable than complete GQs (72). Conversely, increasing K^+^ concentration has been shown to increase the stability of a three- and four-quartet antiparallel bimolecular GQ (73). High mechanical stability (40-55 pN) of parallel stranded GQs formed by the c-MYC (74), hTERT (75) and BCL-2 (76) promoter sequences has been reported; regarding BCL-2, the same study also suggests that the parallel conformer is more stable than hybrid (20-30 pN) topologies. Similarly high stability of various other parallel-stranded GQs have been reported recently (68). On the other hand, investigation of four-quartet GQs found out that unfolding of antiparallel GQ required higher rupture forces than the parallel one (77,78). In this context, stall (arrest) force of polymerases is ~15 pN (79–81) (for a comparison, stall force of the motor protein kinesin is just ~5 pN (82)), which means that GQs may act as roadblocks and common polymerases have to employ other strategies to overcome the GQ obstacle (58). The coexistence of many GQ structures in the human telomeric sequence and possibly even more folding intermediates is a potential source of ambiguities. For example, several force-spectroscopy studies suggest that the human telomeric GQ unfolds via a G-triplex intermediate (44,46,47), while there is no unambiguous evidence thereof in a recent one (67).

Theoretical modelling approaches are an alternative tool to study the GQ mechanical unfolding with a potential to complement the experimental data. Among the methods, molecular dynamics (MD) simulations have been widely employed. Short potential of mean force simulations of the 15-thrombin binding aptamer revealed unfolding of the GQ by unzipping of its strands one by one, so that a G-triplex and G-hairpin were identified as intermediates under the stretching conditions (83). Steered MD (SMD) simulations of parallel GQ formed by the human telomeric sequence led to identification of two different unfolding pathways, specifically division of the GQ into two G-hairpins and strand detachment with a G-triplex remaining (84). The SMD method was later applied to three human telomeric sequence GQ topologies – parallel, hybrid-1, and antiparallel (85); all the topologies unfolded into a G-triplex (or a perturbed one), but the first two by strand detachment, while the antiparallel GQ experienced a step-wise unzipping of nucleotides from both ends. Comparison of the force curves revealed that the rupture force decreased in the order antiparallel > hybrid > parallel GQ. However, a common drawback of those pioneering computational studies (84,85) is that the authors, due to the limited computational resources, had to resort to short simulations with drastically higher (by approximately twelve orders of magnitude) force loads than those commonly applied in the force spectroscopic studies of GQs, which led to unrealistically high rupture forces. This limitation could have affected the unfolding pathways and prevented detection of important intermediates during the unfolding process. A mesoscopic model has also been developed with the aim to apply lower force loads during SMD simulations of the parallel human telomeric GQ (86). However, the simplistic nature of the model (one bead per nucleotide) led to GQ unfolding via unusual G-quartet rotations (i.e., major changes in GQ helicity) and elongations. Hence, fully atomistic explicit solvent models still remain the gold standard to model structural dynamics of GQs.

In this study, we applied the all-atom SMD method on the parallel, hybrid-1 and antiparallel topology of the human telomeric sequence, as well as three other GQs derived from the native ones, examining four pulling protocols. SMD simulations belong to enhanced sampling methods and are able to drag systems from initial configurations to final ones along predefined degrees of freedom called collective variables (87). SMDs were designed to mimic (and complement) experimental force spectroscopy techniques, with two notable differences in addition to the above mentioned gap in used force loads: (i) molecules may be allowed to freely rotate in space during SMD simulations and thus fully adjust to the direction of an external force, and (ii) presence of additional molecules, called linkers, that are attached to the studied system and could (to unknown degree) affect the studied molecule’s free energy surface during experimental measurements, is not required during SMD simulations (Figure 1C) (88–90). We probed the effect of topology, *syn-anti* patterns, loops and direction of force on the stability of GQs during their unfolding pathways. The investigation was carried out under four different pulling setups: (i) a *fast pulling* scheme used in two previous theoretical studies (84,85), (ii) *very slow pulling* setup with two orders of magnitude decreased pulling velocity and an order of magnitude smaller force constant to get closer to the experimental and *in vivo* pulling conditions, (iii) *slow zig-zag pulling* and (iv) *very slow zig-zag pulling* setups, where pulling force is periodically switched off and on again, allowing the system to briefly relax without any applied tension (61,91). We observed that under comparable conditions (i.e., force stiffness) zig-zag pulling protocols usually provide similar unfolding pathways in comparison with results from common continuous constant velocity pulling simulations. We show that the unfolding pathways are dependent not only on – not unexpectedly – the GQ structure itself, but on pulling velocity itself. Independent simulations of the same system under the *fast pulling* lead to rather uniform unfolding pathways for a given structure without any interesting intermediates. On the other hand, independent *very slow pulling* simulations reveal that a given system may follow multiple unfolding pathways. It is a manifestation of the rugged free energy landscape of GQs into which these simulations provided unique insights. We also show that diverse unfolding intermediates in context of the whole DNA molecule sometimes have the same or very similar end-to-end distance. It indicates that structural interpretation of experimental data based on the molecule’s end-to-end distance may be a quite complex problem.

## MATERIAL AND METHODS

### Starting structures and simulation setup

We investigated six different three-quartet GQ structures (Figure 2); three of them were native human telomeric sequence GQs: (i) parallel-stranded GQ with all-*anti* pattern (PDB ID 1KF1 (26)); (ii) 2+2 antiparallel GQ with *anti-syn-anti* pattern (PDB ID 143D (22)); (iii) 3+1 hybrid GQ with *syn-anti-anti* pattern (PDB ID 2GKU (23)). G-stems with loops were taken from the PDB files and thymine nucleotides were attached at both the 5’- and 3’-ends, which resulted in the same d(T<u>GGG</u>TTA<u>GGG</u>TTA<u>GGG</u>TTAGGGT) sequence for the three native human telomeric quadruplexes. The other three structures we used were modified GQs derived from the previous ones: (iv) antiparallel GQ 143D, where the middle diagonal loop was deleted (named as 143D_noloop_); (v) antiparallel GQ 143D with all four guanines in the middle quartet having flipped their *χ* dihedral angle from *anti* to *syn* orientation in two Gs and vice versa in the other two Gs, to obtain *anti-anti-anti* (and *syn-syn-syn*) patterns in the strands (143D_*syn*_); (vi) parallel GQ 1KF1, with all four guanines from the first quartet (near the 5’-terminal thymine (T_1_)) having flipped *χ* dihedral angle from *anti* to *syn* orientation to obtain *syn-anti-anti* pattern (1KF1*syn*). Coordinates of all starting structures are attached in Supplementary Information (PDB files).

**Figure 2.**
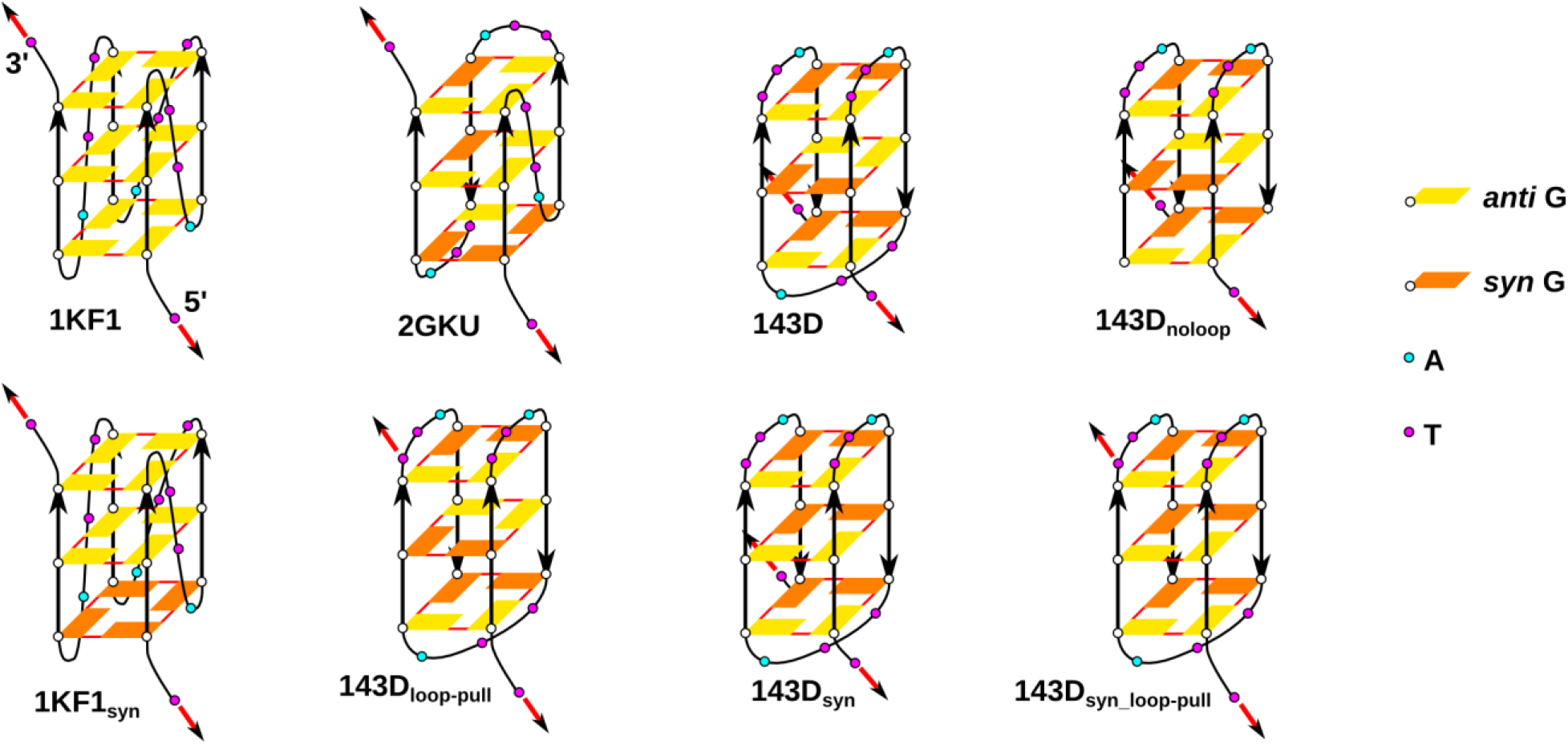
GQ models used in the pulling simulations. The red arrows indicate which two T’s were pulled away and the initial direction of the pulling force.

The starting topologies and coordinates were prepared by using the tLEaP module of AMBER 16 program package (92). Two K^+^ were manually placed inside the GQ channel, if no cations were present in the PDB structure. Models were solvated using a rectangular box with a minimum distance between box walls and solute of 30 Å. Systems were built in OL15 DNA force field (93), SPC/E water model (94) and with ~0.15 M KCl salt using the Joung and Cheatham parameters (95). The systems were subjected to equilibration and thermalization using a common protocol (96). AMBER topologies and coordinates were then converted into GROMACS inputs via PARMED (97) and subsequent pulling simulations were run in GROMACS 2018 (98) in combination with PLUMED 2.5 (99). Pulling simulations were performed at a constant temperature and pressure of 298 K and 1 atm, respectively (100,101). Hydrogen mass repartitioning with a 4-fs integration time step was used (102).

### Pulling simulation strategies

We employed constant velocity pulling using two terminal thymine nucleotides (T1 and T23) of each of the six GQ models and pushed them away from each other by harmonic springs. Specifically, we increased the distance between two centers of mass (COM) formed by C2, C4 and C6 atoms of pyrimidine rings of terminal T_1_ and T_23_ from 5’- and 3’-ends, respectively. In addition, we used modified pulling setup for the two antiparallel 143D and 143D_*syn*_ structures and performed different direction of pulling involving COM of terminal T_1_ and T_17_ from the diagonal loop (instead of the terminal T_23_ nucleobase). Those are labelled as 143D_loop-pull_ and 143D_*syn*_loop-pull_ (derived from 143D and 143D_*syn*_ structures, respectively) and the aim was to employ the same relative direction of force with respect to the G-stem as in 1KF1 and 2GKU. In summary, we investigated forced unfolding in eight different GQ pulling models (Figure 2).

We tried out several testing runs (data not shown) in order to find optimal parameters (mainly force constant and pulling velocity values) with the aim to probe especially initial parts of GQ unfolding (e.g., structural features accompanying breakage of G-quartets). We eventually settled for four pulling setups (Figure 3): (i) *fast pulling* with hard force constant (*k*_0_ = ~1660 pN/nm) and pulling velocity *υ* = ~0.7 nm/ns (setup comparable to previous works by Li *et al.* (84) and Bergues-Pupo *et al.* (85)), (ii) *slow zig-zag pulling* with soft force constant (*k*_0_ = ~150 pN/nm), initial pulling velocity *υ* = ~0.04 nm/ns, followed by a force drop(s) and slower pulling velocity (more information below), (iii) *very slow zig-zag pulling* with the soft force constant (*k*_0_ = ~150 pN/nm), one order of magnitude smaller initial pulling velocity (*υ* = ~0.004 nm/ns), followed by a force drop(s) and slower pulling velocity, and (iv) *very slow pulling* simulations with the soft force constant (*k*_0_ = ~150 pN/nm) and the smaller pulling velocity (*υ* = ~0.004 nm/ns) without any force drop (the latter protocol was used only for 1KF1, 143D and 2GKU systems). Each GQ model was simulated three times under the particular pulling condition, resulting in 81 independent pulling simulations in total (Table 1).

**Figure 3.**
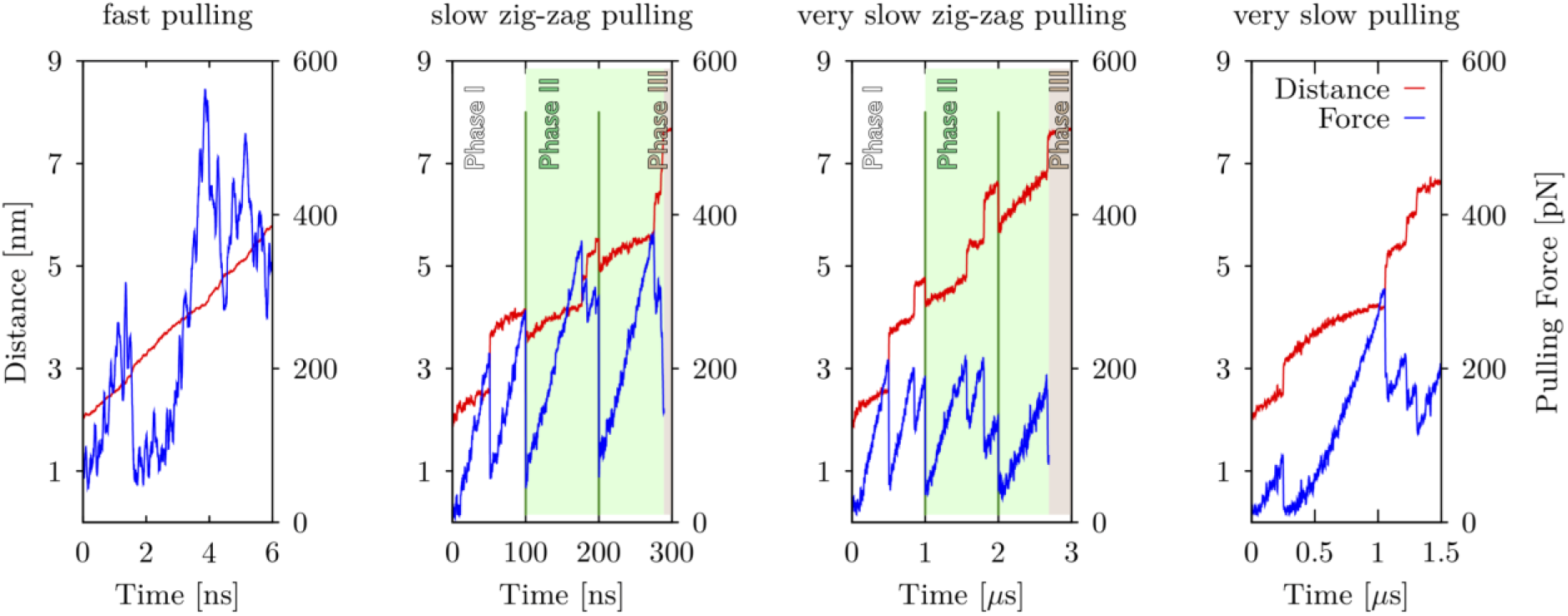
Illustration of four applied pulling protocols as plots showing evolution of the pulling force (in blue) and end-to-end distance (in red) vs. time. Protocols differ mainly by the timescale (horizontal axes) resulting in different pulling velocities (see Methods and Tables S1-S4). Zig-zag pulling protocols (plots in the middle) contain (up to) three different phases. Phase I (white area) resembles common constant velocity pulling (comparable to *fast* and *very slow pulling* protocols). Phase II (green area) is characterized by designed force drops at certain times (green vertical lines), i.e., at 100 ns, 200 ns, 1 μs and 2 μs for *slow zig-zag pulling* and *very slow zig-zag pulling* simulations, respectively. Force drops allowed brief relaxation (often partial refolding) of systems displayed by decrease of distance between pulling centers in the graphs. Occasionally, Phase III is reached, where systems are kept under constant maximum allowed distance between pulling centers; forces are not shown for this stage (brown area; see Methods for details). Notice that data for each plot were chosen from pulling simulations of different GQ topologies and thus, detailed side-by-side comparison of forces and distances in these plots would be misleading.

**Table 1.** Overview of all performed pulling simulations^a^.

| System | Pulling Setup | Simulation Length [μs]<br><sup>b</sup> |
| --- | --- | --- |
| 1KF1 | *fast pulling* | $3 \times 0.006$ |
| 1KF1<sub>syn</sub> | | $3 \times 0.006$ |
| 2GKU | | $3 \times 0.006$ |
| 143D | | $3 \times 0.006$ |
| 143D<sub>syn</sub> | | $3 \times 0.006$ |
| 143D<sub>noloop</sub> | | $3 \times 0.006$ |
| 143D<sub>loop-pull</sub> | | $3 \times 0.006$ |
| 143D<sub>syn_loop-pull</sub> | | $3 \times 0.006$ |
| 1KF1 | *slow*<br>*zig-zag pulling* | $3 \times 0.2^c$ |
| 1KF1<sub>syn</sub> | | $3 \times 0.2^c$ |
| 2GKU | | $3 \times 0.3^c$ |
| 143D | | $3 \times 0.3^c$ |
| 143D<sub>syn</sub> | | $3 \times 0.3^c$ |
| 143D<sub>noloop</sub> | | $3 \times 0.3^c$ |
| 143D<sub>loop-pull</sub> | | $3 \times 0.3^c$ |
| 143D<sub>syn_loop-pull</sub> | | $3 \times 0.3^c$ |
| 1KF1 | *very slow*<br>*zig-zag pulling* | $3 \times 2^d$ |
| 1KF1<sub>syn</sub> | | $3 \times 2^d$ |
| 2GKU | | $3 \times 3^d$ |
| 143D | | $3 \times 4^d$ |
| 143D<sub>syn</sub> | | $3 \times 3^d$ |
| 143D<sub>noloop</sub> | | $3 \times 3^d$ |
| 143D<sub>loop-pull</sub> | | $3 \times 3^d$ |
| 143D<sub>syn_loop-pull</sub> | | $3 \times 2^d$ |
| 1KF1 | *very slow*<br>*pulling* | $3 \times 1.5$ |
| 2GKU | | $3 \times 1.5$ |
| 143D | | $3 \times 1.5$ |

<sup>a</sup> see Methods and Supplementary Tables S1-S4 for details about each system and pulling setups. Pulling directions for each GQ model are visualized in Figure 2 in main text.
<sup>b</sup> " $n \times t$ " means that we performed $n$ independent simulations of a system, each $t$ μs long.
<sup>c</sup> force drops introduced every 0.1 μs
<sup>d</sup> force drops introduced every 1 μs

Trajectories were run for 6 ns and 1.5 μs for *fast pulling* and *very slow pulling*, respectively. *Slow zig-zag pulling* and *very slow zig-zag pulling* simulations were run in a way that introduced force drops at regular intervals of 100 ns and 1 μs, respectively, and the simulations were prolonged as needed (typically twice to reach a simulation length of 300 ns and 3 μs, respectively) until a significant or complete unfolding of the molecule was reached (Figure 3, Table 1). The simulation setup before the first force drop resembles a common constant velocity pulling and is labelled as Phase I, while the whole simulation course with the drops resembles the procedure used in the zig-zag force ramp protocols (61). The pulling velocity was approximately halved after the first drop. The part with decreased pulling velocity and subsequent force drops (if any) until the maximum allowed extension was reached (limitation of the simulation water box) or until the end of the simulation – whichever occurred first – is termed as Phase II. If the maximum allowed extension was reached before the end of simulation, the DNA molecule was kept outstretched at the extension given by the box limit. This final phase (Phase III) is characterized as a *constant end-to-end distance regime*; note that the force magnitude in this phase is meaningless. Notice that it is not possible to predict in advance the appearance of Phase III for particular GQ models, i.e., if or when a given GQ would arrive to the maximum allowed extension range. Side-to-side comparison of applied pulling protocols is shown at Figure (Figure 3). Exact pulling parameters used in particular pulling simulations/phases, namely force constants κ_0_, initial (x_0_) and final (x_τ_) distances between pulling centers, simulation times (τ), and corresponding pulling velocities (ν), are listed in Supplementary Tables S1 – S4.

## RESULTS

### Unfolding intermediates and transitions seen in the simulations

In the course of the pulling simulations, we have seen a wide spectrum of intermediate structures and transitions, which are described in the text below.

Unfolding of GQs usually follows a few characteristic conformational transitions with formation of intermediates (Figure 4). *GQ with reduced number of quartets* is a GQ that has fewer quartets than the native GQ. *G-triplex* is a structure which looks like a GQ, but one G-strand is detached. *5’-triplex* is a G-triplex located near the 5’-end of the sequence, while *3’-triplex* is located near the 3’-end. *Symmetric triplex* is a G-triplex structure, in which the gap left after the missing fourth strand from a GQ is closed by the remaining strands, so that each strand is H-bonded with the other two strands using its Watson-Crick and Hoogsteen edges (42). *G-hairpin* is a structure with only two G-strands connected by the *c*WH or related *t*WW H-bonding. As for the triplex, *5’-hairpin* is located near the 5’-end of the sequence, and *3′-hairpin* is located near the 3’-end; in addition, *middle hairpin* is formed by strands in the middle of the sequence, with both the 5’- and 3’-adjacent strands not involved. In case of 143D and its derivatives, middle hairpin is called *diagonal hairpin,* because the strands were once connected by a diagonal loop and thus were not H-bonded together in the native GQ. *Opened GQ* is a GQ with H-bonding disrupted along one side (groove). The opening occurs between strands that are not connected by a loop (e.g., first and last strand of 2GKU). *Cross-like GQ* (also known as *cross-like structure*) consists of two mutually rotated G-hairpins, which are held together by a network of H-bonds. *Cross-triplex* looks like a cross-like GQ with one G-strand removed. *Cross-hairpin* is just two G-strands held together by H-bonds forming a cross-shape. *Cross-cross structure* encompasses a broad ensemble of structures that are assembling three strands mutually in cross orientation.

**Figure 4.**
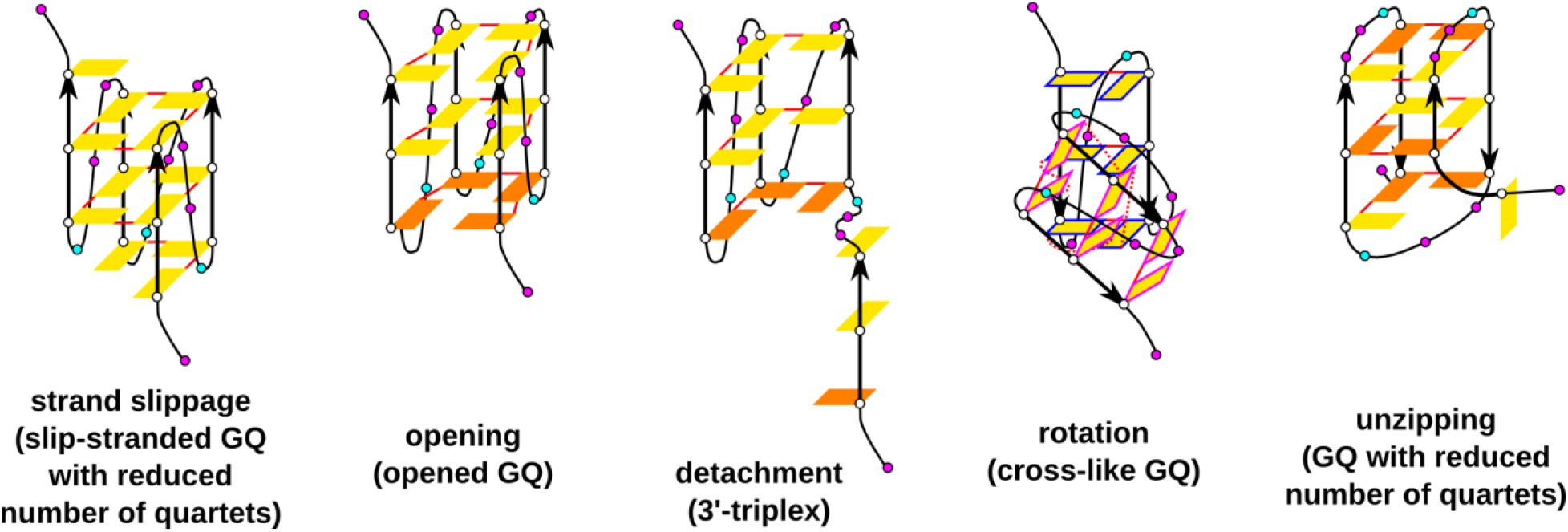
Sketches of common structural transition types and structures observed in the pulling simulations. The initial state of the transitions for all the depicted structures is a native fully folded GQ. In principle, intermediates, such as G-triplexes, may also undergo all the shown transitions (except for opening). In addition to the legend used in Figure 2, the color of perimeter in the cross-like GQ (cross-like structure) indicates which bases are stacked together or are coplanar, solid red line indicated cWH base pairing and the dashed lines indicate non-*c*WH base pairing.

*Strand slippage* is a vertical movement of a G-strand shifting its Gs one or more levels upwards or downwards with respect to the rest of the G-stem (36,53). It is allowed in G-stems if Gs in one strand stacked on each other have the same *χ* orientation. *Spiral structure* is a slip-stranded intermediate, in which the slippage is incomplete in such a way that Gs of the slipping strand are connected to the original quartet at one edge, and the other edge is H-bonded with Gs originally located in the quartet above or below. *Opening* of GQ is disruption of the cyclical bonding of quartets along one side of GQ and it typically occurs between the first and last strand of GQ. An opened GQ may undergo *division*, which is its disintegration into two separated G-hairpins. *Unzipping* is a movement during which a G is pulled out of GQ. Multiple back-to-back unzipping events in a single strand may happen, which is strand unzipping; a special case of strand unzipping is *detachment*, in which all Gs of a single G-strand are unzipped abruptly (almost) at once. *Rotation* of strands is accompanied by partial breaking of native *c*WH H-bonding and potential formation of new non-native H-bonds. The above described motions are not limited to GQs, some of these may happen during unfolding of intermediates themselves, such as G-triplexes. Representative structures of all the mentioned intermediates are attached in Supplementary Information (PDB files).

Not all unfolding/rupture events can be characterized by the transitions and intermediates shown in Figure 4. Some detected events are due to unfolding of structures not directly related to the GQ core, *e.g*., we observed a rupture of a stacked intercalated structure formed by previously unfolded G-strand and the adjacent loop. This further underlines the enormous richness and multidimensionality of the conformational space of the GQs which can be visited upon unfolding and folding processes. We also mention here that, not surprisingly, major unfolding structural transitions mostly coincide with force drops and molecule’s extension (see Supplementary Figures S1-S27). Of note, sometimes the event in the graphs seems lagging behind the force drop; the reason is that the force drop may be associated with, for example, loss of two H-bonds during unzipping, but we take the time of the event when it is fully developed, *i.e*., all four H-bonds holding the G within a quartet are disrupted.

### Antiparallel telomeric GQ is toughest, parallel is weakest

The magnitude of rupture forces of GQs depends on the pulling protocol. The highest rupture forces – and greatest variance of their magnitude between independent simulations of a given system – were found for the *fast pulling* protocol, and the lowest forces in *very slow* and *zig-zag very slow pulling* protocols (Table 2). Among the four tested pulling protocols we consistently observed that the antiparallel basket GQ 143D required the highest rupture forces, followed by the hybrid GQ 2GKU, and the parallel GQ 1KF1 resisted the least under a given pulling scheme.

**Table 2.** GQ rupture force values (in pN) observed in the pulling simulations.^a^

| System<sup>b</sup> | *Fast pulling* | *Slow zig-zag pulling* | *Very slow zig-zag pulling* | *Very slow pulling* |
| --- | --- | --- | --- | --- |
| 1KF1 | 420, 420, 420 | 200, 250, 200 | 150+160, 150+180, 210 | 150+160, 140, 180 |
| 1KF1<sub>syn</sub> | 450, 450, 450 | 200, 270, 270 | 150, 190, 190 | --- |
| 2GKU | 430, 390, 510 | 300, 300, 380 | 250, 220, 220 | 200, 200, 200 |
| 143D | 730, 650, 550 | 380, 360, 340 | 300, 300, 300 | 330, 300, 370 |
| 143D<sub>syn</sub> | 700, 530, 580 | 340+340, 350+370,<br>360+370 | 200, 300, 290 | --- |
| 143D<sub>noloop</sub> | 580, 580, 580 | 150+420, 200+380,<br>290+450 | 200+250,<br>210+320, 200+250 | --- |
| 143D<sub>loop-pull</sub> | 440+800, 580+700,<br>510+680 | 300, 380, 380+410 | 350, 300, 250 | --- |
| 143D<sub>syn_loop-pull</sub> | 510, 510, 510 | 420, 220, 410 | 260, 260, 270 | --- |

<sup>a</sup> The values come from three independent simulations. Two values are shown (X+Y) if the highest force reached in a given simulation does not correspond to the first major GQ structural perturbation; the first value shows the force reached when the first GQ perturbation event occurred while the second value shows the highest observed force in
<sup>b</sup> See Methods and Figure 2 for details about labelling of GQ topologies.

### Quartets of opposite directionality and presence of loops increase mechanical stability

The pulling simulations revealed another two important trends (Table 2): i) GQs containing a mixture of quartet directionalities resist the force better – regardless of the direction of the initial force, as judged from the stability order of the pairs 1KF1 < 1KF1_syn_, 143D_syn_ < 143D, and 143D_syn_loop-pull_ < 143D_loop-pull_,; ii) the diagonal loop of the antiparallel GQ plays a stabilizing role blocking the unzipping as shown by the pair 143D_noloop_ < 143D.

### Force drops in the zig-zag pulling may alter unfolding resistance in both directions

Significant force drops during a molecule’s pulling may happen two ways, either internally after a major unfolding/rupture event, or externally by the zig-zag pulling protocol (the latter being also connected to decreased molecule’s end-to-end extension). The *very slow zig-zag pulling* simulations usually followed similar unfolding pathways as the *very slow pulling* simulations. Nevertheless, a few times in the zig-zag setup, we observed that unfolding occurred after an external force drop at a force smaller than the peak force just before the drop, which indicates a certain degree of structural plasticity. After the force drop the system apparently uses degrees of freedom orthogonal to the pulling collective variable to more efficiently overcome the rupture barrier.

In some simulations of the antiparallel GQ the force needed to disrupt a second or third quartet is similar or even higher than that needed to disrupt the first quartet (Figure 5). This behavior is apparently linked to the zig-zag protocol and may mean that the force drop allows for a relaxation and settling down of the remaining quartets. On the other hand, some simulations of the antiparallel GQ showed rather cooperative unfolding, with the force required to disrupt the first quartet being clearly the highest.

**Figure 5.**
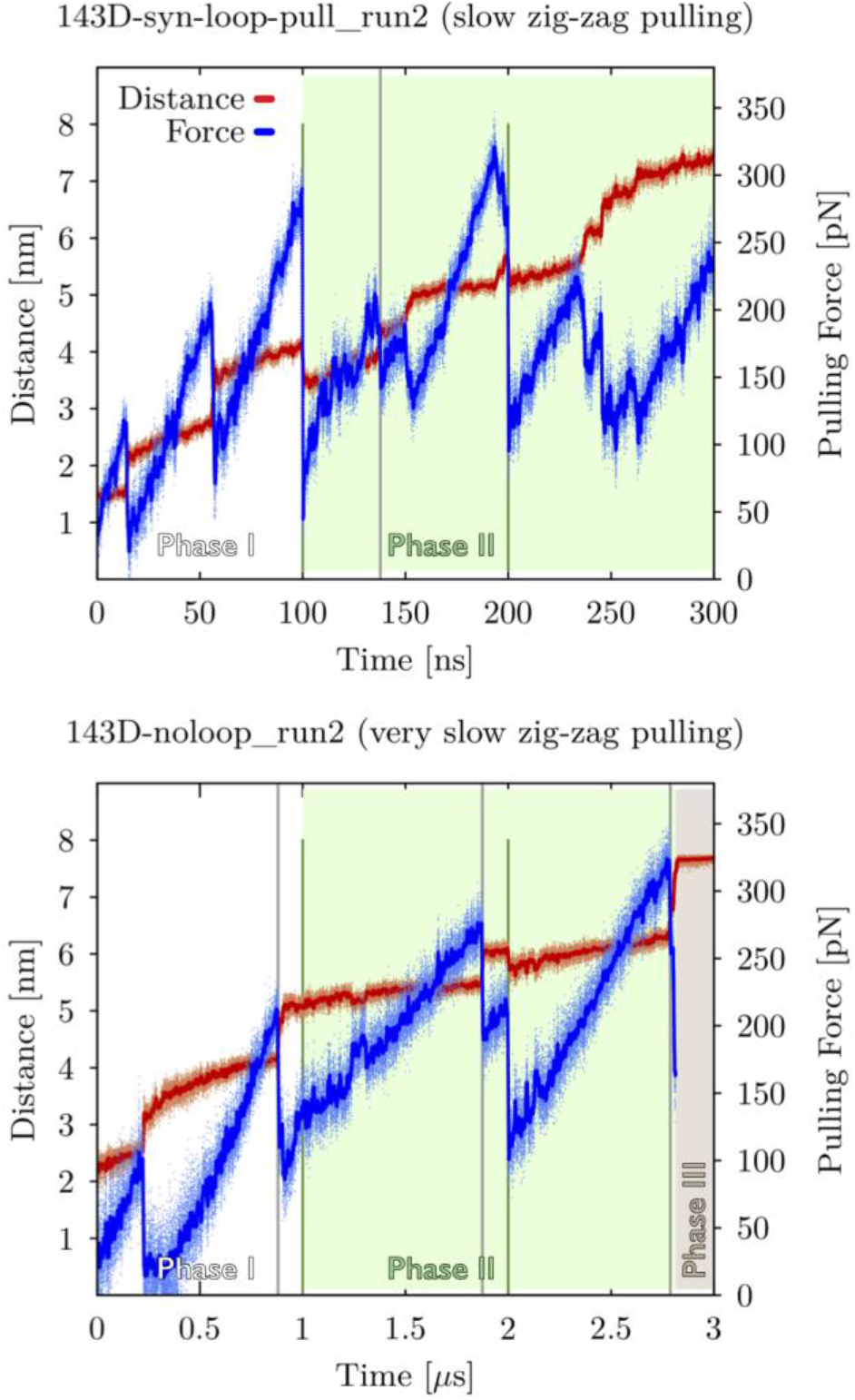
Enforced unfolding of 143D_*syn*_loop-pull_ and 143D_noloop_ GQ models during *slow zig-zag pulling* simulation (top panel) and *very slow zig-zag pulling* simulation (panel at the bottom), respectively. Both plots show evolution of pulling force (in blue) and end-to-end distance (in red) vs. time with pulling phases and intended force drops highlighted by green lines (see Figure 3 and Methods for details). The pulling force reached ~290 pN without having any effect on the 143D_*syn*_loop-pull_ model. After the externally induced force drop at 100 ns, an unfolding event occurred at a force of just ~220 pN (grey vertical line, top panel). The bottom panel shows a case example of second and third quartet resisting a force higher than needed to disrupt the previous quartet; unfolding of the first quartet required ~210 pN force (leftmost grey vertical), the second (middle) quartet resisted to ~280 pN (middle grey vertical line) and the third quartet resisted to a force up to ~320 pN (rightmost grey vertical). Notice that the first two drops of the pulling force in the top panel and the first drop in the bottom panel (and notable prolongation of end-to-end distances) correspond to unstacking of the terminal Ts.

### Two main structural determinants of initial unfolding mechanism – collocation of GQ’s termini and its *syn/anti* pattern

Mutual position of GQ termini translates into the force direction exerted on a given GQ (Figure 2). If both the termini are connected to the same quartet, such as in the antiparallel basket 143D, the force acts across a single quartet, i.e., it is nearly perpendicular to the GQ vertical axis, and base unzipping is the dominant initial unfolding mechanism (Table 3 and Figure 4). If the termini are connected to the quartets at opposite ends of the GQ, the force acts across several quartets in a direction more parallel to the GQ axis, then the *syn/anti* pattern of the GQ is the factor that comes into play: (i) in GQs with no alteration of *syn/anti* Gs within a single G-strand, i.e., when all the quartets are of the same directionality, such as in the parallel-stranded 1KF1, vertical strand slippage takes place commonly, (ii) when a mixture of *syn* and *anti* Gs is present, such as in the 3+1 hybrid 2GKU, the quartets of opposite directionality sterically block the strand slippage and GQ opening is rather followed by rotation into cross-like GQ (Figure 6). Detailed simulation outcome for each GQ system can be found in Supplementary Information (Supplementary Results, Tables S5-S8 and Figures S1-S27).

**Figure 6.**
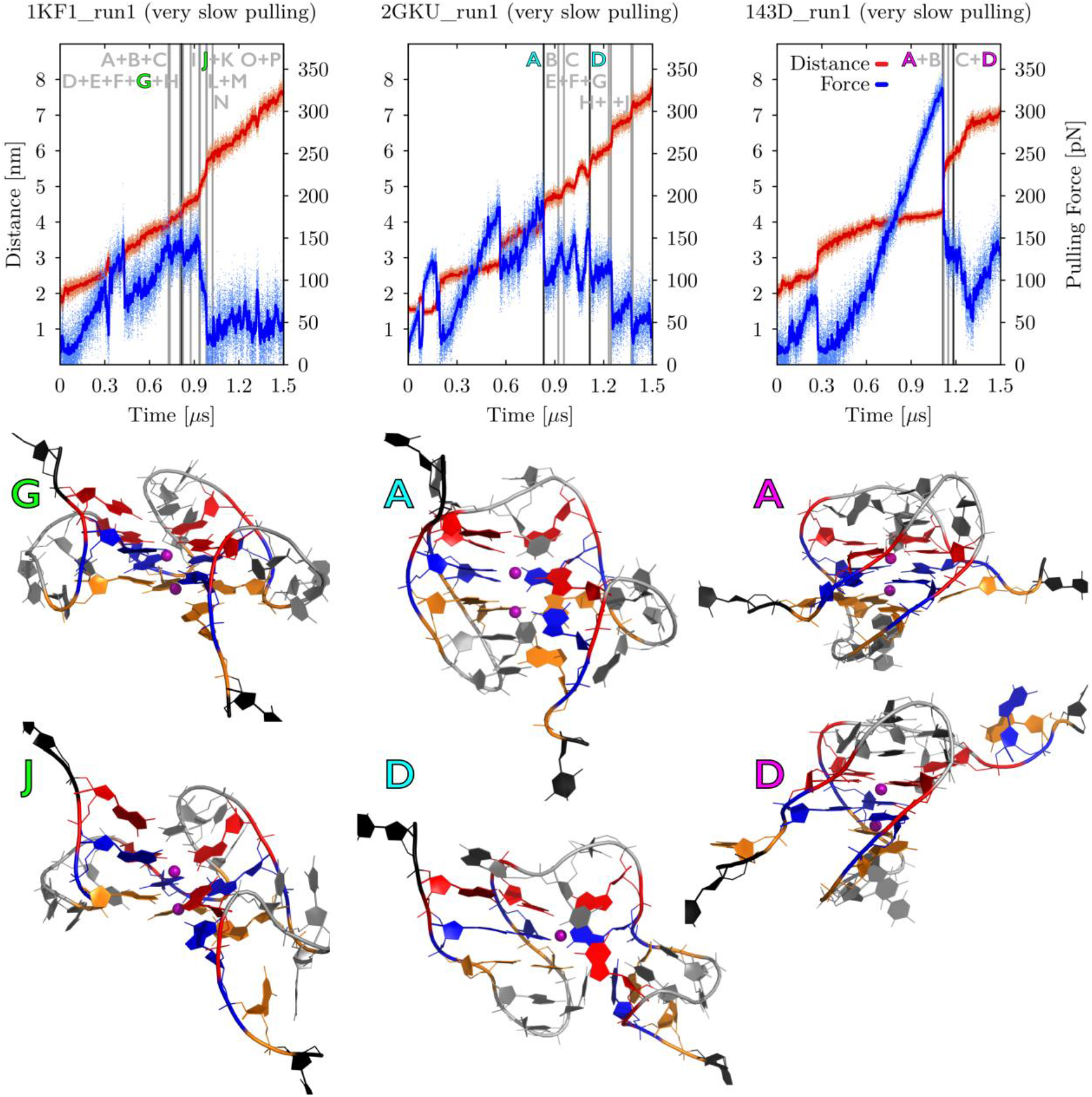
Side-by-side comparison of enforced unfolding of three main GQ systems, i.e., parallel 1KF1, hybrid 2GKU and antiparallel 143D. Plots at the top show evolution of pulling force (in blue) and end-to-end distance (in red) vs. time during *very slow pulling* simulations. It is clearly visible that parallel 1KF1 unfolded the most willingly, whereas antiparallel basket 143D GQ required significantly higher rupture forces (see Table 2 for details). Major structural events are shown (vertical lines) and labelled (capital letters). The most important pulling intermediates are highlighted (black vertical lines with letters in specific colors, i.e., green for 1KF1, cyan for 2GKU and magenta for 143D) and shown as structural snapshots under the plots. Gs from first, second and third G-quartet are highlighted in orange, blue and red, respectively. Both terminal T residues (i.e., pulling centers) are shown in black, remaining DNA residues are in gray and channel K^+^ ions are shown as purple spheres. Hydrogen atoms and water molecules are not shown for clarity. See Supplementary Figures S25-S27 for snapshots of all the unfolding intermediates from these particular simulations and complete data from all three independent pulling simulations.

**Table 3.** Main unfolding pathways and intermediates observed in pulling simulations^a^.

| System<sup>b</sup> | *Fast pulling*<br>(see Table S5) | *Slow zig-zag pulling*<br>(see Table S6) | *Very slow zig-zag pulling*<br>(see Table S7) | *Very slow pulling*<br>(see Table S8) |
| --- | --- | --- | --- | --- |
| 1KF1 | strand slippage leading to G-triplex | strand slippage leading to G-triplex; rotation into cross-like GQ and its division | mixture of strand slippage, strand detachment, base unzipping and rotation into cross-like structures | mixture of base unzipping, strand slippage and strand detachment |
| 1KF1<sub>syn</sub> | GQ opening leading to G-triplex or division into G-hairpins | GQ rotation into cross-like GQ, unfolding to G-triplex or G-hairpin; strand unzipping to G-triplex | GQ opening and rotation into cross-like GQ, followed by complex base unzipping and strand detachments | --- |
| 2GKU | GQ opening and rotation into cross-like GQ | GQ opening and rotation into cross-like GQ, unfolding to G-triplex or G-hairpin | GQ opening and rotation into cross-like GQ, unzipping and detachments, rotation into cross-like structures | GQ opening and rotation into cross-like GQ, then mixture of base unzipping and detachments |
| 143D | GQ opening leading to G-triplex; unzipping of terminal Gs | unzipping of first strand leading to G-triplex, then unzipping to G-hairpins or rotation to cross-like triplex | unzipping of terminal Gs to incomplete triplexes or unstable diagonal hairpins | unzipping of terminal Gs to incomplete G-triplex or diagonal hairpins |
| 143D<sub>syn</sub> | unzipping of terminal Gs; unzipping to G-triplex | unzipping of terminal Gs, strand unzipping to imperfect triplexes | unzipping of terminal strands, followed by detachment of adjacent strand to G-hairpin | --- |
| 143D<sub>noloop</sub> | unzipping of terminal Gs | unzipping of terminal strand, detachment of adjacent strand | unzipping of terminal Gs, detachment of adjacent strand | --- |
| 143D<sub>loop-pull</sub> | GQ opening leading to G-triplex; unzipping of strands leading to G-hairpins | complex unfolding pathways including strand unzipping, GQ opening, rotation into cross-like structures, strand detachment | base unzipping combined with GQ opening and strand detachment, 3'-hairpin remained | --- |
| 143D<sub>syn_loop-pull</sub> | incomplete strand slippage leading to spiral structure, followed by GQ opening and base unzipping | complex unfolding pathways including incomplete strand slippage to spiral structures, base unzipping, GQ opening, rotation into cross-like triplexes | Incomplete strand slippage to spiral structure, then complex unfolding pathways including GQ opening, strand detachment and unzipping | --- |

<sup>a</sup> Detailed outcome of each simulation can be found in Supplementary Tables S5 – S8.

Detachment of both terminal T’s from their stacked position on GQ were the first two unfolding events in the simulations (except for one) and were identified before any GQ unfolding took place. These two events often correspond to the first two major force drops in the force-time graphs (see, e.g., the force drops before the first GQ stem unfolding event in Figure 5; Figures S1-S27). After that, the GQ unfolding was system-specific. In terms of molecular interactions, the GQ stem unfolding in all tested systems was initiated by breaking of H-bonds of a G with neighboring Gs and losing its coordination with the channel cation, regardless of the unfolding mechanism.

### The richest structural dynamics is under the slowest pulling velocity

Comparison of the unfolding pathways between the tested pulling schemes reveals that GQs under high tension and *fast pulling* velocity underwent structural disintegration quickly, but the short time did not allow the GQs to unfold extensively, e.g. into G-hairpins. Structural transitions in individual simulations of a particular system were rather uniform, i.e., independent trajectories of one system followed mostly similar pathways (Supplementary Table S5). As the pulling velocity and spring hardness decreased, the unfolding dynamics became richer (Supplementary Table S6). The *very slow* and *very slow zig-zag pulling* allowed a GQ to follow different unfolding pathways in the independent simulations, with a wider variety of unfolding intermediates. Furthermore, even multiple partial refolding events were observed in some *very slow* and *very slow zig-zag pulling* simulations (Supplementary Tables S7 and S8).

Almost none unfolding events happened in Phase III (i.e., the last “constant distance” phase) of zig-zag pulling simulations, nevertheless, even this regime did not prevent unfolding completely (Supplementary Table S9), which means the molecules still tend to relax and relieve the stress.

### Dynamics in unfolded ensembles – refolding is possible even with increasing end-to-end distance

When molecule unfolds under the force, the unfolded parts do not necessarily behave like extended rigid rods. In fact, the backbone of the unfolded part is where the strain is usually absorbed (after possible GQ rotation in space), by extending to a straighter conformation, because the GQ and its unfolding intermediates are held together by stronger interactions. If the unfolded part is sufficiently long and the whole molecule’s end-to-end extension is short enough, the unfolded part may undergo complex dynamics, even leading to partial refolding events; for example, GQ tearing-up into two G-hairpins may be followed by unfolding of one of the hairpins and the freed G-strand may attach to the other hairpin to form a G-triplex or cross-triplex. The refolding is facilitated by structural memory of the molecule and it requires enough time, so we observed it mostly in *very slow* and *zig-zag very slow pulling* conditions. Refolding attempts usually occurred after major unfolding events, which were accompanied by the relaxation of emerged structures. Only one refolding event was observed in *fast pulling* simulations, where the shape memory is lost quickly and the molecule does not have enough time to relax.

### Molecules rotate in space upon force exertion

Relative direction of the force acting on a given GQ changed during the simulation, as the structure was getting unfolded. The direction of force often differed significantly from the initial direction shown in Figure 2. Because the folded part is stiffer than the unfolded part, the former naturally tended to rotate itself in space while the already unfolded part can undergo conformational changes that relieve the stress. Thus, greatest changes in relative force direction occurred when a loop or multiple G’s at once got ruptured. The molecule’s rotation in space to maximize the length of the folded part in the direction of the force was diminished in the *fast pulling* simulations, because the unfolding was so fast (and thus the simulation time so short) that the GQ did not have enough time to fully respond.

## DISCUSSION

In this work, we present an extended set of atomistic pulling MD simulations of GQs of human telomeric sequence with the aim to better understand their unfolding process under tension and their folding landscape. Results of all 81 individual pulling simulations are fully documented in Supplementary Tables S5-S8 (description of structural transitions) and Supplementary Figures S1-S27 (force-time and extension-time graphs and key structures occurring during the unfolding process). The main outcome of the work is perhaps the richness of conformational space which participate in the unfolding once we apply gentler pulling. Evidently, the slower the process and softer the spring, the more and more structural diversity is seen. In the later stages of pulling, the slower process allows partial relaxations of the previously ruptured structural elements. We suggest a possibility that under common experimental pulling conditions the GQ sequences could experience even much richer dynamics than seen in our simulation.

### Impact of pulling protocol on unfolding

We have used four different pulling schemes (*fast, slow zig-zag, very slow zig-zag* and *very slow*; see Methods). The *fast pulling* protocol used here mimics the setup that was used in previous two SMD studies on human telomeric GQs (84,85), and thus we can compare the previous results with our results and with the other three protocols. In general, our *fast pulling* SMD simulations have provided similar results, i.e., similar magnitude of rupture forces on the order of hundreds of pN (Table 2) and similar unfolding pathways (Table 3), as the previous studies. Differences arise, however, when we compare these results with the other three protocols employing smaller pulling velocities and spring hardnesses (i.e., smaller force loads). Using these updated pulling protocols, rupture forces indeed decreased by a few hundreds of pN (Table 2), and more importantly, richer and more complex structural dynamics was observed (Table 3). In other words, under high force loads, the molecule is being shifted from equilibrium so fast that it does not have time to respond to actual conditions, while smaller force loads enable the molecule to undergo also small conformational changes in a local equilibrium for some time, which relieves the stress and even allows for minor (partial) refolding events.

The length of our slower SMD simulations allowed for extensive unfolding of all the GQ models, so we indeed encountered more intermediate states, such as incomplete G-triplexes, cross-triplexes, G-hairpins and cross-hairpins (Tables S5–S8), which could not be reached in our *fast pulling* SMD protocol, as well as in the comparable protocol used in the previous computational studies (84,85). The claimed dependence of structural richness on pulling velocity is nevertheless primarily based on the greater diversity in the first stages of GQ unfolding in *slow zig-zag, very slow* and *very slow zig-zag pulling* simulations (Tables S5–S8).

GQs follow unfolding pathways with various structural intermediates (Figure 4), which differ for different GQ topologies and *anti*-*syn* patterns. Antiparallel GQs, in which the force acts within a single G-quartet, have been suggested to unfold by sequential unzipping of terminal Gs (78,83,85). Our current SMD simulations corroborate this idea under all unfolding conditions for 143D, 143D_noloop_ and 143D_syn_ models, and furthermore show that the unzipping can be symmetric from both termini or asymmetric, when one terminal strand unfolds more than the other (Tables 3 and S5–S8). The mechanism of unzipping of a G from a complete G-stem indicates that H-bonding is prone to breaking more than stacking when the GQ molecule is being pulled by flanking nucleotides, i.e., via the backbone. The same unzipping initiation was also observed previously (85).

The hybrid-1 GQ 2GKU unfolding was initiated by opening followed by rotation into cross-like GQ in all tested pulling schemes (Tables 3 and S5–S8). On the contrary, previous SMD of the hybrid human telomeric GQ showed strand detachment leading to a 3’-G-triplex (85). Nevertheless, we have also eventually observed this G-triplex intermediate state in the slower pulling schemes.

The impact of pulling protocol is also manifested in SMD of the parallel stranded GQ 1KF1. Under the conditions of *fast pulling,* G-triplex formation (it is unfortunately not clear whether the G-triplex was formed by strand slippage or strand detachment) (84,85) and GQ division into two hairpins (84) have been observed. Our *fast pulling* simulations also lead to G-triplex (by strand slippage in 1KF1 and by strand detachment in 1KF1_syn_; note that strand slippage is sterically blocked in 1KF1_syn_) or to GQ division into G-hairpins (Tables 3 and S5–S8). Nevertheless, the other three pulling protocols have shown additional possible unfolding pathways of 1KF1, namely GQ rotation into cross-like GQ, followed by division into G-hairpins, or a complex mixture of strand slippage (not observed in 1KF1_syn_), base unzipping and strand detachment.

The importance of force direction is further shown by our 143D simulations, when we applied the force in a direction along the groove (143D_loop-pull_ and 143D_syn_loop-pull_ models), i.e., the force acted in a similar direction as in the parallel 1KF1 and hybrid 2GKU GQs. Terminal base unzipping was no longer dominant and other mechanisms, such as GQ opening, strand slippage (in the 143D_syn_loop-pull_ model) and strand detachment, emerged (Tables 3 and S5–S8). While the effect of the force direction on the unfolding mechanism is obvious, unfortunately, we cannot draw conclusions about the magnitude of resistance of GQs to external forces depending on the initial direction of the force; having a look at the antiparallel GQ, the rupture forces for 143D and 143D_loop-pull_ models were mostly similar, but the same did not apply to the pair of 143D_syn_ and 143D_syn_loop-pull_ models, for which the order differed among the pulling schemes (Table 2).

The simulations also provide an example of how applied bias may affect the unfolding pathway. For example, preferential strand slippage in the direction of shortening the distance propeller loop has to span (e.g., strand slippage of the last strand in 3’-direction, Figure 4) was observed in unbiased simulations of the parallel stranded GQ 1KF1 (36), as well as in well-tempered metadynamics simulation with the bias applied to the total number of H-bonds and G-G stacking interactions in the G-stem (37). The same slippage direction was observed also in this work, because it is in the direction of acting force. On the other hand, directionality of rotation into cross-like structure observed here led to prolonged propeller loop (Figure 4), which is opposite to direction of rotation into cross-like intermediates seen in unbiased simulations (34,36,42). The reason is that the direction of exerted force is given by the positions of strand termini and eventually the force overcame the strain in the loop. It is a nice example how the molecule end-to-end distance, acting as a collective variable for pulling, affects exploration of the free energy landscape compared to the free molecule and how unfolding movements orthogonal to the collective variable can be missed.

### Mechanical stability of human telomeric GQs

Comparison of results from available theoretical and experimental studies of GQs is unfortunately not straightforward. SMD simulations provide detailed atomistic description of the unfolding intermediates, but are limited by their timescale (and thus orders of magnitude higher force loads) and unreliable statistics of forces and extensions in comparison to single molecule force spectroscopies experiments, so the two approaches rather complement each other.

The stability of different GQ topologies of human telomeric sequence has been a matter of debate. The mechanical stability trend predicted from our current data (under all four pulling conditions) and also from the previous fast SMD simulations (85) suggest the order antiparallel > hybrid > parallel topology. Given that these trends take place regardless of the pulling velocity and spring hardness, one could expect the same trends to occur even under lower force loads used in the experimental measurements. However, multiple experiments have suggested that the antiparallel human telomeric GQ in Na^+^ solution is mechanically less stable than hybrid and parallel topologies in comparable K^+^ solution (65–68,70).

The contradiction might originate in multiple reasons. First off, the sheer difference in force loads of about ten orders of magnitude might simply mean the simulation results are not transferable to current experiments (and vice versa). Given the structural polymorphism of the human telomeric sequence in conjunction with the limited structural resolution of the single-molecule techniques, the interpretation of the experiments might have been affected by the inability to directly determine which GQ topologies are present in Na^+^ and K^+^ solution; however, we think this option is rather unlikely in the well-studied case of human telomeric sequence. Still, it is possible that the experiments may find some even more complex unfolding-refolding processes than we see in the SMD simulations. Obviously, we cannot rule out that the simulation force field underestimates stability of the parallel stranded quadruplex, possibly because problems in the description of propeller loops (103). On the other hand, although such force field problem could have affected the order of resistance to unfolding in SMD simulations, we do not think it significantly affected the observed unfolding pathways.

Currently, a hope to overcome the barrier between experiments and simulations is aimed at high-speed AFM (104,105), which closes the gap in timescale, pulling velocity and force load between SMD simulations and experiments (89,104). The SMD simulations in the *very slow pulling* regime performed here (force constant *k*_0_ of ~150 pN/nm, pulling velocity *ν* of ~0.004 nm/ns) already belong to the range accessible to high-speed AFM.

Mechanical stability of GQs is important *in vivo*, when GQs are formed in ssDNA chains that are being read by proteins. Stall forces of polymerases are ~15 pN (79–81), which is less than the measured rupture force of GQs (~20-55 pN depending on topology and sequence (67,68,71,74–76)), so these proteins do have difficult times when encountering a GQ and they often need to recruit specialized helicases that resolve GQs (6,56,58). Our zig-zag pulling SMD simulations show a few instances of a GQ being disrupted at lower force after the force drop in Phase II than was the peak force in Phase I (Figure 5). It suggests that a part of GQs could eventually be resolved even by a “weaker” polymerase after repeated unwinding attempts of the protein, i.e., in a sequence of force ramps and drops.

### Mechanical stability of G-triplexes and hairpins

It has been hypothesized that GQ (un)folding may proceed via G-triplex or hairpin intermediates. The existence of parallel G-triplex and hairpin remains questionable (34,42), but there is evidence suggesting stability of antiparallel and hybrid species coming from both experimental and computational studies (35,38,39,41–44,46–52,70,85). Nevertheless, the true importance of these intermediates during forced unfolding has been doubted recently, as their mechanical stability (if any) has been estimated to be below a few pN, while GQs can resist forces an order of magnitude higher (67). The study found no detectable formation of G-triplexes upon truncation of the human telomeric sequence to three hexanucleotide repeats. Our current data rather support possible intermediary role of G-triplexes in unfolding of GQs; the simulations show that G-triplexes and G-hairpins are indeed formed during the mechanical unfolding (and also refolded in a few events) and the mechanical stability of G-triplexes corresponds to non-negligible rupture forces, about two to five times smaller than those of GQ. Still, it could be a consequence of the force loads of the SMD simulations, which could have led to steric clashes increasing the required rupture force of the G-triplex; i.e., the molecule was not capable to fully relax after the first rupture event. Therefore, the role of G-triplexes in non-equilibrium unfolding may depend on the speed of unfolding and does not prove a role of G-triplexes in folding. We reiterate that the pulling protocol biases the free energy landscape by dimensionality reduction (as a collective variable) and the faster is the pulling the larger is the bias. In summary, the simulations suggest role of G-triplexes in fast pulling but it does not imply their role in folding under equilibrium conditions. We also note that due to the kinetic partitioning of the GQ folding landscape, unexpected GQ species might form during the folding phases of some experiments, including weaker two-quartet GQs (20,34,40). We hypothesize that some of them could be occasionally interpreted as G-triplex states.

### Cations binding requires three interacting G-strands

Channel cations are essential for structural stability of GQs. A question remains as of at which stage the cations get bound to the DNA during the folding process, i.e., what is the minimum structure capable of cation binding. Our pulling simulations show that GQs, regardless of the topology and unfolding mechanism, keep binding the channel cations even if a whole G-strand has departed. Upon larger unfolding, if the remaining structure still contains remnants of at least G-triads or a *c*WH G:G pair with another G in the cross-like orientation, it is still able to bind one channel cation. At the G-hairpin stage, cations may remain bound, but when it turns into a cross-hairpin, the binding is lost, unless, rarely, a cation-binding pocket within the cross-hairpin is formed. Our previous standard and enhanced sampling MD simulations starting with G-triplex showed that they do bind cations, even in the cross-triplex state (34,42); G-hairpins, on the other hand, displayed rather weak cation binding both in standard simulations and replica exchange simulations aiming at folding of G-hairpins (34,52). Hence, the presented pulling simulations provide results in agreement with the previous data. It thus seems that the structural cations can be bound essentially anytime during the folding process. However, the probability is low for a hairpin-like intermediate, but increases to almost certainty when three G-strands are involved.

### Determination of structures based on their end-to-end distances is not unambiguous

Structural interpretation of single-molecule experiments is usually based on the molecule’s extension at a given moment. Typically, one measures the end-to-end distance in a GQ (or a putative G-triplex, etc.) in the structure deposited in the PDB database and adds a multiple of standard nucleotide lengths, corresponding to the tether and supposedly unfolded parts of the GQ, and seeks for match between the calculation and actual measurement. However, our simulations suggest that such a structural determination may be ambiguous in some cases; for example, we show that a G-triplex found in one simulation and a G-hairpin in another one may both have very similar end-to-end distance within the whole molecule’s context (G-hairpin actually having slightly shorter distance than G-triplex; Figure 7). While this is certainly not a rule, it demonstrates the variability of nucleic acid chains and therefore the limitations of the end-to-end-calculation based approaches.

**Figure 7.**
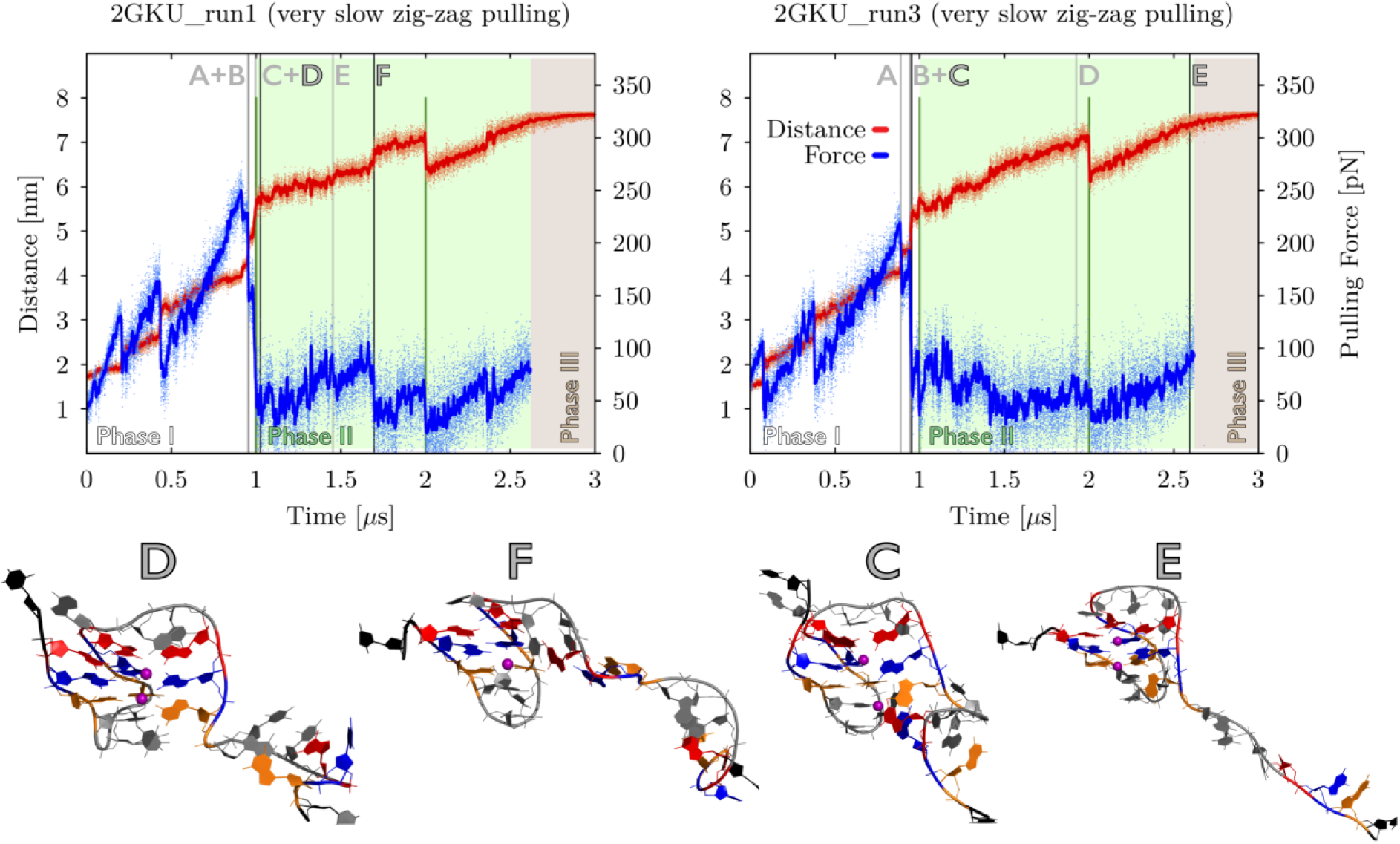
Pulling force vs. time and end-to-end distance vs. time graphs of two very slow zig-zag pulling simulations of the 3+1 hybrid 2GKU. G-hairpin (run1, letter “F”) can have shorter end-to-end distance than G-triplex (run3, letter “E”) when taken in context of the whole molecule. Notice that both the unfolding pathways proceeded via a very similar G-triplex state (run1, letter “D” and run3, letter “C”). Pulling phases and intended force drops in the very slow zig-zag protocol are highlighted (see Figure 3 and Methods for details). Important events are labeled by grey capital letters in the graphs and two key structures from each simulation (corresponding to the highlighted letters) are shown as snapshots below the graphs (see the legend of Figure 5 for details about the coloring scheme). Detailed description and snapshots of all the intermediates are in Supplementary Table S7 and Supplementary Figure S19.

## CONCLUSIONS

We have investigated mechanochemical properties of three human telomeric sequence GQs and three related GQs using SMD simulations. We observed a couple of unfolding transition types and unfolding intermediates (Figure 4). The preferred initial transition type for a given system depended on the relative direction of external force to the GQ. The more perpendicular to the GQ channel axis was the force, the more likely was a step-wise unzipping mechanism, as typically observed in the antiparallel GQ 143D. On the other hand, if the force was more parallel to the GQ channel axis and all the quartets had the same directionality, such as in the parallel GQ 1KF1, strand slippage mechanism could take place. If the quartet directionality was mixed, GQ opening followed by rotation into cross-like GQ was typical, as was the case of hybrid-1 GQ 2GKU. Unfolding pathways following these initial steps were diverse, but we recurrently observed various G-triplexes, G-hairpins and their cross-like analogues, i.e., structures that have been hypothesized to participate in GQ (un)folding before. These structures nevertheless appear to be less stable than GQs.

Importantly, we observed that multiple simulations of a given GQ using *fast pulling* with stronger spring led to monotonous unfolding pathways, but *very slow pulling* with weaker spring, which mimics *in vivo* and *in vitro* conditions more closely, allowed for diversified and more complex pathways, including multiple partial refolding attempts (Table 3). The variety of found pathways, intermediates and transition types is another piece of evidence that the GQ free energy landscape is rugged. In addition, the ruggedness is likely responsible for the fact that we have observed that a GQ might occasionally unfold after a force drop at forces lower than it withstood before the drop (Figure 5), which indicates that stall forces of proteins such as polymerases *in vivo* do not always need to reach declared rupture forces of GQs.

Furthermore, we showed that the GQ unfolding pathways could end up at different intermediates, but the DNA overall end-to-end distance could be almost identical, as manifested by the G-hairpin and G-triplex formed from the hybrid-1 GQ (Figure 7). Such structures could thus be difficult to distinguish in force spectroscopy data interpretation.

Two out of four sets of simulations parameters (pulling velocities and force constants) used in this study were low enough to reach the values accessible to modern high-speed AFM. It promises that our data could be directly complemented by experimental studies of GQs in the near future.

## DATA AVAILABILITY

The data underlying this article will be shared on reasonable request to the corresponding author due to the large size of the raw simulation trajectories.

## SUPPLEMENTARY DATA

Supplementary data are available at NAR Online: Supplementary results, Supplementary Tables S1 – S9 and Figures S1 – S27, and PDB files with coordinates of the starting systems and representative key intermediate structures.

## ACKNOWLEDGEMENT

We acknowledge the use of CESNET data storage facilities [grant no. LM2018140].

## REFERENCES

1. Chambers, V.S., Marsico, G., Boutell, J.M., Di Antonio, M., Smith, G.P. and Balasubramanian, S. (2015) High-Throughput Sequencing of DNA G-Quadruplex Structures in the Human Genome. Nat. Biotech., 33, 877–881.

2. Bedrat, A., Lacroix, L. and Mergny, J.-L. (2016) Re-Evaluation of G-Quadruplex Propensity with G4Hunter. Nucleic Acids Res., 44, 1746–1759.

3. Rhodes, D. and Lipps, H.J. (2015) G-Quadruplexes and Their Regulatory Roles in Biology. Nucleic Acids Res., 43, 8627–8637.

4. Varshney, D., Spiegel, J., Zyner, K., Tannahill, D. and Balasubramanian, S. (2020) The Regulation and Functions of DNA and RNA G-Quadruplexes. Nat. Rev. Mol. Cell Biol., 21, 459–474.

5. Lee, W.T.C., Yin, Y., Morten, M.J., Tonzi, P., Gwo, P.P., Odermatt, D.C., Modesti, M., Cantor, S.B., Gari, K., Huang, T.T. et al. (2021) Single-Molecule Imaging Reveals Replication Fork Coupled Formation of G-Quadruplex Structures Hinders Local Replication Stress Signaling. Nat. Commun., 12, 2525.

6. Lejault, P., Mitteaux, J., Sperti, F.R. and Monchaud, D. (2021) How to Untie G-Quadruplex Knots and Why? Cell Chem. Biol., 28, 436–455.

7. Di Antonio, M., Ponjavic, A., Radzevičius, A., Ranasinghe, R.T., Catalano, M., Zhang, X., Shen, J., Needham, L.-M., Lee, S.F., Klenerman, D. et al. (2020) Single-Molecule Visualization of DNA G-Quadruplex Formation in Live Cells. Nat. Chem., 12, 832–837.

8. Graziella, C.-R., Nadia, Z. and Marco, F. (2016) Emerging Role of G-Quadruplex DNA as Target in Anticancer Therapy. Curr. Pharm. Des., 22, 6612–6624.

9. Tateishi-Karimata, H., Kawauchi, K. and Sugimoto, N. (2018) Destabilization of DNA G-Quadruplexes by Chemical Environment Changes during Tumor Progression Facilitates Transcription. J. Am. Chem. Soc., 140, 642–651.

10. Lee, J., Sung, K., Joo, S.Y., Jeong, J.-H., Kim, S.K. and Lee, H. (2022) Dynamic Interaction of BRCA2 with Telomeric G-Quadruplexes Underlies Telomere Replication Homeostasis. Nat. Commun., 13, 3396.

11. Carvalho, J., Mergny, J.-L., Salgado, G.F., Queiroz, J.A. and Cruz, C. (2020) G-quadruplex, Friend or Foe: The Role of the G-quartet in Anticancer Strategies. Trends Mol. Med., 26, 848–861.

12. Kosiol, N., Juranek, S., Brossart, P., Heine, A. and Paeschke, K. (2021) G-Quadruplexes: A Promising Target for Cancer Therapy. Mol. Cancer, 20, 40.

13. Maizels, N. (2015) G4-Associated Human Diseases. EMBO Rep., 16, 910–922.

14. Balendra, R. and Isaacs, A.M. (2018) C9orf72-Mediated ALS and FTD: Multiple Pathways to Disease. Nat. Rev. Neurol., 14, 544–558.

15. Asamitsu, S., Yabuki, Y., Ikenoshita, S., Wada, T. and Shioda, N. (2020) Pharmacological Prospects of G-Quadruplexes for Neurological Diseases Using Porphyrins. Biochem. Biophys. Res. Commun., 531, 51–55.

16. Stefan, L. and Monchaud, D. (2019) Applications of Guanine Quartets in Nanotechnology and Chemical Biology. Nat. Rev. Chem., 3, 650–668.

17. da Silva, M.W. (2007) Geometric Formalism for DNA Quadruplex Folding. Chem. Eur. J., 13, 9738–9745.

18. Karsisiotis, A.I., O’Kane, C. and da Silva, M.W. (2013) DNA Quadruplex Folding Formalism - A Tutorial on Quadruplex Topologies. Methods, 64, 28–35.

19. Dvorkin, S.A., Karsisiotis, A.I. and Webba da Silva, M. (2018) Encoding Canonical DNA Quadruplex Structure. Sci. Adv., 4, eaat3007.

20. Sponer, J., Islam, B., Stadlbauer, P. and Haider, S. (2020) In Neidle, S. (ed.), Annual Reports in Medicinal Chemistry. Academic Press, Vol. 54, pp. 197–241.

21. Dai, J.X., Carver, M. and Yang, D.Z. (2008) Polymorphism of Human Telomeric Quadruplex Structures. Biochimie, 90, 1172–1183.

22. Wang, Y. and Patel, D.J. (1993) Solution Structure of the Human Telomeric Repeat d[AG(3)(T(2)AG(3))3] G-tetraplex. Structure, 1, 263–282.

23. Luu, K.N., Phan, A.T., Kuryavyi, V., Lacroix, L. and Patel, D.J. (2006) Structure of the Human Telomere in K+ Solution: An Intramolecular (3+1) G-Quadruplex Scaffold. J. Am. Chem. Soc., 128, 9963–9970.

24. Phan, A.T., Kuryavyi, V., Luu, K.N. and Patel, D.J. (2007) Structure of Two Intramolecular G-quadruplexes Formed by Natural Human Telomere Sequences in K+ Solution. Nucleic Acids Res., 35, 6517–6525.

25. Lim, K.W., Amrane, S., Bouaziz, S., Xu, W., Mu, Y., Patel, D.J., Luu, K.N. and Phan, A.T. (2009) Structure of the Human Telomere in K+ Solution: A Stable Basket-Type G-Quadruplex with Only Two G-Tetrad Layers. J. Am. Chem. Soc., 131, 4301–4309.

26. Parkinson, G.N., Lee, M.P.H. and Neidle, S. (2002) Crystal Structure of Parallel Quadruplexes from Human Telomeric DNA. Nature, 417, 876–880.

27. Lim, K.W., Ng, V.C.M., Martin-Pintado, N., Heddi, B. and Phan, A.T. (2013) Structure of the Human Telomere in Na+ Solution: An Antiparallel (2+2) G-quadruplex Scaffold Reveals Additional Diversity. Nucleic Acids Res., 41, 10556–10562.

28. Dai, J., Punchihewa, C., Ambrus, A., Chen, D., Jones, R.A. and Yang, D. (2007) Structure of the Intramolecular Human Telomeric G-quadruplex in Potassium Solution: A Novel Adenine Triple Formation. Nucleic Acids Res., 35, 2440–2450.

29. Palacky, J., Vorlickova, M., Kejnovska, I. and Mojzes, P. (2013) Polymorphism of Human Telomeric Quadruplex Structure Controlled by DNA Concentration: A Raman Study. Nucleic Acids Res., 41, 1005–1016.

30. Sponer, J., Bussi, G., Stadlbauer, P., Kuhrova, P., Banas, P., Islam, B., Haider, S., Neidle, S. and Otyepka, M. (2017) Folding of Guanine Quadruplex Molecules–Funnel-Like Mechanism or Kinetic Partitioning? An Overview From MD Simulation Studies. Biochim. Biophys. Acta Gen. Subj., 1861, 1246–1263.

31. Bessi, I., Jonker, H.R., Richter, C. and Schwalbe, H. (2015) Involvement of Long-Lived Intermediate States in the Complex Folding Pathway of the Human Telomeric G-Quadruplex. Angew. Chem., Int. Ed., 54, 8444–8448.

32. Long, X. and Stone, M.D. (2013) Kinetic Partitioning Modulates Human Telomere DNA G-Quadruplex Structural Polymorphism. PLoS One, 8, e83420.

33. Thirumalai, D., Klimov, D.K. and Woodson, S.A. (1997) Kinetic Partitioning Mechanism as a Unifying Theme in the Folding of Biomolecules. Theor. Chem. Acc., 96, 14–22.

34. Stadlbauer, P., Kuhrova, P., Vicherek, L., Banas, P., Otyepka, M., Trantirek, L. and Sponer, J. (2019) Parallel G-Triplexes and G-Hairpins As Potential Transitory Ensembles in the Folding of Parallel-Stranded DNA G-Quadruplexes. Nucleic Acids Res., 47, 7276–7293.

35. Bian, Y., Ren, W., Song, F., Yu, J. and Wang, J. (2018) Exploration of the Folding Dynamics of Human Telomeric G-Quadruplex with a Hybrid Atomistic Structure-Based Model. J. Chem. Phys., 148, 204107.

36. Stadlbauer, P., Krepl, M., Cheatham, T.E., 3rd, Koca, J. and Sponer, J. (2013) Structural Dynamics of Possible Late-Stage Intermediates in Folding of Quadruplex DNA Studied by Molecular Simulations. Nucleic Acids Res., 41, 7128–7143.

37. Rocca, R., Palazzesi, F., Amato, J., Costa, G., Ortuso, F., Pagano, B., Randazzo, A., Novellino, E., Alcaro, S., Moraca, F. et al. (2020) Folding Intermediate States of the Parallel Human Telomeric G-Quadruplex DNA Explored Using Well-Tempered Metadynamics. Sci. Rep., 10, 3176.

38. Bian, Y., Song, F., Cao, Z., Zhao, L., Yu, J., Guo, X. and Wang, J. (2018) Fast-Folding Pathways of the Thrombin-Binding Aptamer G-Quadruplex Revealed by a Markov State Model. Biophys. J., 114, 1529–1538.

39. Bian, Y.-Q., Song, F., Cao, Z.-X., Yu, J.-F. and Wang, J.-H. (2021) Structure-Based Simulations Complemented by Conventional All-Atom Simulations to Provide New Insights Into the Folding Dynamics of Human Telomeric G-Quadruplex*. Chin. Phys. B, 30, 078702.

40. Marchand, A. and Gabelica, V. (2016) Folding and Misfolding Pathways of G-Quadruplex DNA. Nucleic Acids Res., 44, 10999–11012.

41. Bian, Y., Tan, C., Wang, J., Sheng, Y., Zhang, J. and Wang, W. (2014) Atomistic Picture for the Folding Pathway of a Hybrid-1 Type Human Telomeric DNA G-quadruplex. PLoS Comput. Biol., 10, e1003562.

42. Stadlbauer, P., Trantirek, L., Cheatham, T.E., 3rd, Koca, J. and Sponer, J. (2014) Triplex Intermediates in Folding of Human Telomeric Quadruplexes Probed by Microsecond-scale Molecular Dynamics Simulations. Biochimie, 105, 22–35.

43. Mashimo, T., Yagi, H., Sannohe, Y., Rajendran, A. and Sugiyama, H. (2010) Folding Pathways of Human Telomeric Type-1 and Type-2 G-quadruplex Structures. J. Am. Chem. Soc., 132, 14910–14918.

44. Koirala, D., Mashimo, T., Sannohe, Y., Yu, Z.B., Mao, H.B. and Sugiyama, H. (2012) Intramolecular Folding in Three Tandem Guanine Repeats of Human Telomeric DNA. Chem. Commun., 48, 2006–2008.

45. Hou, X.-M., Fu, Y.-B., Wu, W.-Q., Wang, L., Teng, F.-Y., Xie, P., Wang, P.-Y. and Xi, X.-G. (2017) Involvement of G-Triplex and G-Hairpin in the Multi-Pathway Folding of Human Telomeric G-Quadruplex. Nucleic Acids Res., 45, 11401–11412.

46. Li, W., Hou, X.-M., Wang, P.-Y., Xi, X.-G. and Li, M. (2013) Direct Measurement of Sequential Folding Pathway and Energy Landscape of Human Telomeric G-quadruplex Structures. J. Am. Chem. Soc., 135, 6423–6426.

47. Jiang, H.-X., Cui, Y., Zhao, T., Fu, H.-W., Koirala, D., Punnoose, J.A., Kong, D.-M. and Mao, H. (2015) Divalent Cations and Molecular Crowding Buffers Stabilize G-Triplex at Physiologically Relevant Temperatures. Sci. Rep., 5, 9255.

48. Lu, X.-M., Li, H., You, J., Li, W., Wang, P.-Y., Li, M., Dou, S.-X. and Xi, X.-G. (2018) Folding Dynamics of Parallel and Antiparallel G-Triplexes under the Influence of Proximal DNA. J. Phys. Chem. B, 122, 9499–9506.

49. Limongelli, V., De Tito, S., Cerofolini, L., Fragai, M., Pagano, B., Trotta, R., Cosconati, S., Marinelli, L., Novellino, E., Bertini, I. et al. (2013) The G-Triplex DNA. Angew. Chem., Int. Ed., 52, 2269–2273.

50. Cerofolini, L., Amato, J., Giachetti, A., Limongelli, V., Novellino, E., Parrinello, M., Fragai, M., Randazzo, A. and Luchinat, C. (2014) G-Triplex Structure and Formation Propensity. Nucleic Acids Res., 42, 13393–13404.

51. Yang, C., Kulkarni, M., Lim, M. and Pak, Y. (2017) Insilico Direct Folding of Thrombin-Binding Aptamer G-Quadruplex at All-Atom Level. Nucleic Acids Res., 45, 12648–12656.

52. Stadlbauer, P., Kuhrova, P., Banas, P., Koca, J., Bussi, G., Trantirek, L., Otyepka, M. and Sponer, J. (2015) Hairpins Participating in Folding of Human Telomeric Sequence Quadruplexes Studied by Standard and T-REMD Simulations. Nucleic Acids Res., 43, 9626–9644.

53. Stefl, R., Cheatham, T.E., Spackova, N., Fadrna, E., Berger, I., Koca, J. and Sponer, J. (2003) Formation Pathways of a Guanine-Quadruplex DNA Revealed by Molecular Dynamics and Thermodynamic Analysis of the Substates. Biophys. J., 85, 1787–1804.

54. Havrila, M., Stadlbauer, P., Kuhrova, P., Banas, P., Mergny, J.-L., Otyepka, M. and Sponer, J. (2018) Structural Dynamics of Propeller Loop: Towards Folding of RNA G-Quadruplex. Nucleic Acids Res., 46, 8754–8771.

55. Islam, B., Stadlbauer, P., Krepl, M., Koca, J., Neidle, S., Haider, S. and Sponer, J. (2015) Extended Molecular Dynamics of a c-kit Promoter Quadruplex. Nucleic Acids Res., 43, 8673–8693.

56. Mendoza, O., Bourdoncle, A., Boulé, J.-B., Brosh, R.M., Jr. and Mergny, J.-L. (2016) G-Quadruplexes and Helicases. Nucleic Acids Res., 44, 1989–2006.

57. Hansel-Hertsch, R., Di Antonio, M. and Balasubramanian, S. (2017) DNA G-Quadruplexes in the Human Genome: Detection, Functions and Therapeutic Potential. Nat. Rev. Mol. Cell Biol., 18, 279–284.

58. Estep, N.K., Butler, J.T., Ding, J. and Brosh, M.R. (2019) G4-Interacting DNA Helicases and Polymerases: Potential Therapeutic Targets. Curr. Med. Chem., 26, 2881–2897.

59. Paeschke, K., Bochman, M.L., Garcia, P.D., Cejka, P., Friedman, K.L., Kowalczykowski, S.C. and Zakian, V.A. (2013) Pif1 Family Helicases Suppress Genome Instability at G-Quadruplex Motifs. Nature, 497, 458–462.

60. Postberg, J., Tsytlonok, M., Sparvoli, D., Rhodes, D. and Lipps, H.J. (2012) A Telomerase-associated RecQ Protein-like Helicase Resolves Telomeric G-quadruplex Structures during Replication. Gene, 497, 147–154.

61. Yang, B., Liu, Z., Liu, H. and Nash, M.A. (2020) Next Generation Methods for Single-Molecule Force Spectroscopy on Polyproteins and Receptor-Ligand Complexes. Frontiers in Molecular Biosciences, 7, doi: 10.3389/fmolb.2020.00085.

62. Fang, J., Xie, C., Tao, Y. and Wei, D. (2022) An Overview of Single-Molecule Techniques and Applications in the Study of Nucleic Acid Structure and Function. Biochimie, doi: 10.1016/j.biochi.2022.09.014.

63. Cheng, Y., Zhang, Y. and You, H. (2021) Characterization of G-Quadruplexes Folding/Unfolding Dynamics and Interactions with Proteins from Single-Molecule Force Spectroscopy. Biomolecules, 11, 1579.

64. Laszlo, A.H., Derrington, I.M. and Gundlach, J.H. (2016) MspA Nanopore as a Single-Molecule Tool: From Sequencing to SPRNT. Methods, 105, 75–89.

65. Koirala, D., Dhakal, S., Ashbridge, B., Sannohe, Y., Rodriguez, R., Sugiyama, H., Balasubramanian, S. and Mao, H. (2011) A Single-Molecule Platform for Investigation of Interactions Between G-Quadruplexes and Small-Molecule Ligands. Nat. Chem., 3, 782–787.

66. You, H., Zeng, X., Xu, Y., Lim, C.J., Efremov, A.K., Phan, A.T. and Yan, J. (2014) Dynamics and Stability of Polymorphic Human Telomeric G-Quadruplex under Tension. Nucleic Acids Res., 42, 8789–8795.

67. Mitra, J., Makurath, M.A., Ngo, T.T.M., Troitskaia, A., Chemla, Y.R. and Ha, T. (2019) Extreme Mechanical Diversity of Human Telomeric DNA Revealed by Fluorescence-Force Spectroscopy. Proc. Natl. Acad. Sci. U. S. A., 116, 8350–8359.

68. Cheng, Y., Zhang, Y., Gong, Z., Zhang, X., Li, Y., Shi, X., Pei, Y. and You, H. (2020) High Mechanical Stability and Slow Unfolding Rates Are Prevalent in Parallel-Stranded DNA G-Quadruplexes. J. Phys. Chem. Lett., 11, 7966–7971.

69. Long, X., Parks, J.W., Bagshaw, C.R. and Stone, M.D. (2013) Mechanical Unfolding of Human Telomere G-quadruplex DNA Probed by Integrated Fluorescence and Magnetic Tweezers Spectroscopy. Nucleic Acids Res., 41, 2746–2755.

70. Dhakal, S., Cui, Y., Koirala, D., Ghimire, C., Kushwaha, S., Yu, Z., Yangyuoru, P.M. and Mao, H. (2013) Structural and Mechanical Properties of Individual Human Telomeric G-quadruplexes in Molecularly Crowded Solutions. Nucleic Acids Res., 41, 3915–3923.

71. Mitra, J. and Ha, T. (2019) Streamlining Effects of Extra Telomeric Repeat on Telomeric DNA Folding Revealed by Fluorescence-Force Spectroscopy. Nucleic Acids Res., 47, 11044–11056.

72. Zhang, Y., Cheng, Y., Chen, J., Zheng, K. and You, H. (2021) Mechanical Diversity and Folding Intermediates of Parallel-Stranded G-Quadruplexes with a Bulge. Nucleic Acids Res., 49, 7179–7188.

73. Lynch, S., Baker, H., Byker, S.G., Zhou, D. and Sinniah, K. (2009) Single Molecule Force Spectroscopy on G-Quadruplex DNA. Chem. Eur. J., 15, 8113–8116.

74. You, H., Wu, J., Shao, F. and Yan, J. (2015) Stability and Kinetics of c-MYC Promoter G-Quadruplexes Studied by Single-Molecule Manipulation. J. Am. Chem. Soc., 137, 2424–2427.

75. Yu, Z., Gaerig, V., Cui, Y., Kang, H., Gokhale, V., Zhao, Y., Hurley, L.H. and Mao, H. (2012) Tertiary DNA Structure in the Single-Stranded hTERT Promoter Fragment Unfolds and Refolds by Parallel Pathways via Cooperative or Sequential Events. J. Am. Chem. Soc., 134, 5157–5164.

76. Cheng, Y., Tang, Q., Li, Y., Zhang, Y., Zhao, C., Yan, J. and You, H. (2019) Folding/Unfolding Kinetics of G-Quadruplexes Upstream of the P1 Promoter of the Human BCL-2 Oncogene. J. Biol. Chem., 294, 5890–5895.

77. Yu, Z., Schonhoft, J.D., Dhakal, S., Bajracharya, R., Hegde, R., Basu, S. and Mao, H. (2009) ILPR G-Quadruplexes Formed in Seconds Demonstrate High Mechanical Stabilities. J. Am. Chem. Soc., 131, 1876–1882.

78. de Messieres, M., Chang, J.-C., Brawn-Cinani, B. and La Porta, A. (2012) Single-Molecule Study of G-Quadruplex Disruption Using Dynamic Force Spectroscopy. Phys. Rev. Lett., 109, 058101.

79. Yin, H., Wang, M.D., Svoboda, K., Landick, R., Block, S.M. and Gelles, J. (1995) Transcription Against an Applied Force. Science, 270, 1653–1657.

80. Galburt, E.A., Grill, S.W., Wiedmann, A., Lubkowska, L., Choy, J., Nogales, E., Kashlev, M. and Bustamante, C. (2007) Backtracking Determines the Force Sensitivity of RNAP II in a Factor-Dependent Manner. Nature, 446, 820–823.

81. Mejia, Y.X., Mao, H., Forde, N.R. and Bustamante, C. (2008) Thermal Probing of E. coli RNA Polymerase Off-Pathway Mechanisms. J. Mol. Biol., 382, 628–637.

82. Meyhöfer, E. and Howard, J. (1995) The Force Generated by a Single Kinesin Molecule Against an Elastic Load. Proc. Natl. Acad. Sci. U. S. A., 92, 574–578.

83. Yang, C., Jang, S. and Pak, Y. (2011) Multiple Stepwise Pattern for Potential of Mean Force in Unfolding the Thrombin Binding Aptamer in Complex with Sr2+. J. Chem. Phys., 135, 225104.

84. Li, H., Cao, E.H. and Gisler, T. (2009) Force-Induced Unfolding of Human Telomeric G-quadruplex: A Steered Molecular Dynamics Simulation Study. Biochem. Biophys. Res. Commun., 379, 70–75.

85. Bergues-Pupo, A.E., Arias-Gonzalez, J.R., Moron, M.C., Fiasconaro, A. and Falo, F. (2015) Role of the Central Cations in the Mechanical Unfolding of DNA and RNA G-quadruplexes. Nucleic Acids Res., 43, 7638–7647.

86. Bergues-Pupo, A.E., Gutiérrez, I., Arias-Gonzalez, J.R., Falo, F. and Fiasconaro, A. (2017) Mesoscopic Model for DNA G-Quadruplex Unfolding. Sci. Rep., 7, 11756.

87. Sotomayor, M. and Schulten, K. (2007) Single-Molecule Experiments in Vitro and in Silico. Science, 316, 1144–1148.

88. Franz, F., Daday, C. and Gräter, F. (2020) Advances in Molecular Simulations of Protein Mechanical Properties and Function. Curr. Opin. Struct. Biol., 61, 132–138.

89. Sheridan, S., Gräter, F. and Daday, C. (2019) How Fast Is Too Fast in Force-Probe Molecular Dynamics Simulations? J. Phys. Chem. B, 123, 3658–3664.

90. Stirnemann, G. (2022) Recent Advances and Emerging Challenges in the Molecular Modeling of Mechanobiological Processes. J. Phys. Chem. B, 126, 1365–1374.

91. Jacobson, D.R., Uyetake, L. and Perkins, T.T. (2020) Membrane-Protein Unfolding Intermediates Detected with Enhanced Precision Using a Zigzag Force Ramp. Biophys. J., 118, 667–675.

92. Case, D.A., Betz, R.M., Botello-Smith, W., Cerutti, D.S., T.E. Cheatham, I., Darden, T.A., Duke, R.E., Giese, T.J., Gohlke, H., Goetz, A.W. et al. (2016). AMBER16. University of California, San Francisco, CA.

93. Zgarbova, M., Sponer, J., Otyepka, M., Cheatham, T.E., Galindo-Murillo, R. and Jurecka, P. (2015) Refinement of the Sugar–Phosphate Backbone Torsion Beta for AMBER Force Fields Improves the Description of Z- and B-DNA. J. Chem. Theory Comput., 11, 5723–5736.

94. Berendsen, H.J.C., Grigera, J.R. and Straatsma, T.P. (1987) The Missing Term in Effective Pair Potentials. J. Phys. Chem., 91, 6269–6271.

95. Joung, I.S. and Cheatham, T.E. (2008) Determination of Alkali and Halide Monovalent Ion Parameters for Use In Explicitly Solvated Biomolecular Simulations. J. Phys. Chem. B, 112, 9020–9041.

96. Kuhrova, P., Mlynsky, V., Zgarbova, M., Krepl, M., Bussi, G., Best, R.B., Otyepka, M., Sponer, J. and Banáš, P. (2019) Improving the Performance of the Amber RNA Force Field by Tuning the Hydrogen-Bonding Interactions. J. Chem. Theory Comput., 15, 3288–3305.

97. Shirts, M.R., Klein, C., Swails, J.M., Yin, J., Gilson, M.K., Mobley, D.L., Case, D.A. and Zhong, E.D. (2017) Lessons Learned from Comparing Molecular Dynamics Engines on the SAMPL5 Dataset. J. Comput.-Aided Mol. Des., 31, 147–161.

98. Abraham, M.J., Murtola, T., Schulz, R., Pall, S., Smith, J.C., Hess, B. and Lindahl, E. (2015) GROMACS: High Performance Molecular Simulations through Multi-Level Parallelism from Laptops to Supercomputers. SoftwareX, 1-2, 19–25.

99. Tribello, G.A., Bonomi, M., Branduardi, D., Camilloni, C. and Bussi, G. (2014) PLUMED 2: New Feathers for an Old Bird. Comput. Phys. Commun., 185, 604–613.

100. Bussi, G., Donadio, D. and Parrinello, M. (2007) Canonical Sampling through Velocity Rescaling. J. Chem. Phys., 126, 014101.

101. Parrinello, M. and Rahman, A. (1981) Polymorphic Transitions in Single Crystals: A New Molecular Dynamics Method. J. Appl. Phys., 52, 7182–7190.

102. Hopkins, C.W., Le Grand, S., Walker, R.C. and Roitberg, A.E. (2015) Long-Time-Step Molecular Dynamics through Hydrogen Mass Repartitioning. J. Chem. Theory Comput., 11, 1864–1874.

103. Islam, B., Stadlbauer, P., Gil-Ley, A., Perez-Hernandez, G., Haider, S., Neidle, S., Bussi, G., Banas, P., Otyepka, M. and Sponer, J. (2017) Exploring the Dynamics of Propeller Loops in Human Telomeric DNA Quadruplexes Using Atomistic Simulations. J. Chem. Theory Comput., 13, 2458–2480.

104. Rico, F., Russek, A., González, L., Grubmüller, H. and Scheuring, S. (2019) Heterogeneous and Rate-Dependent Streptavidin–Biotin Unbinding Revealed by High-Speed Force Spectroscopy and Atomistic Simulations. Proc. Natl. Acad. Sci. U. S. A., 116, 6594–6601.

105. Rico, F., Gonzalez, L., Casuso, I., Puig-Vidal, M. and Scheuring, S. (2013) High-Speed Force Spectroscopy Unfolds Titin at the Velocity of Molecular Dynamics Simulations. Science, 342, 741–743.

