## Supplementary Information for "Complexity of Guanine Quadruplex Unfolding Pathways Revealed by Atomistic Pulling Simulations"

### TABLE OF CONTENS

SUPPLEMENTARY RESULTS ........ - 2 -

SUPPLEMENTARY TABLES ........ - 5 -

SUPPLEMENTARY FIGURES ........ - 13 -

REFERENCES ........ - 118 -

### SUPPLEMENTARY RESULTS

**GQ unfolding dynamics**. The results in the following paragraphs describe main developments seen in the simulations. Detailed outcome of all the individual simulations can be found in Tables S5–S8. Detailed plots showing pulling force vs. time and end-to-end distance vs. time from all simulations with corresponding figures summarizing main structural events are shown in Figures S1-S27.

*Parallel-stranded GQ*. The force acting on 1KF1 was almost in the vertical direction along the groove. This, combined with the all-*anti* pattern of the GQ, facilitated vertical strand slippage, making it the dominant unfolding mechanism under all pulling conditions. The slipping strand could be the first or the last one, thus 3'-triplex or 5'-triplex, respectively, was formed. The triplex could unfold further. The only simulation that did not lead to G-triplex went via rotation into the cross-like GQ, which was then divided into two eventually unstable G-hairpins. The richest structural dynamics during the 1KF1 unfolding was observed, as expected, in both *slow zig-zag* and *very slow zig-zag pulling* simulations and also under *very slow pulling* simulations, where we observed even a few refolding attempts. Of note, the direction of rotation into cross-like GQ observed in one pulling simulation was the opposite of unbiased MD simulations; if the molecule is unrestrained, the rotation happens in the direction of the propeller loop spanning shorter distance, while in the pulling simulation the rotation happened in the direction of exerted force.

On the other hand, $1KF1_{syn}$, derived from 1KF1 by flipping its 5'-quartet to all-*syn*, did not undergo any strand slippage, which proves that the *syn-anti* strand mix sterically prohibits such movement. Instead, we observed puling-velocity-dependent scenarios. In *fast pulling* simulations, opening of the GQ was observed. It was followed either by horizontal detachment of the last G-strand, resulting into a G-triplex, or division of the opened GQ in the half into two separated G-hairpins. In *slow zig-zag* and *very slow zig-zag pulling* simulations, the unfolding was more complicated, with multiple unfolding steps seen. Base unzipping, strand detachment and rotations into cross-like ensembles in various order were the most employed mechanisms.

*3+1 hybrid-1 GQ*. 2GKU felt the force in the same direction as 1KF1 and $1KF1_{syn}$ models. All the four tested pulling velocity protocols led most commonly to opening of the GQ followed by formation of a cross-like GQ. Under *slow zig-zag pulling*, the unfolding proceeded further from cross-like GQ to G-hairpin or G-triplex. Both *very slow zig-zag* and *very slow pulling* simulations revealed the most diverse outcomes, i.e., we identified nearly all the movement types described in the main text paragraph *Unfolding intermediates and transitions*, except for strand slippage and division. Also the terminal state in the simulations differed, i.e., we obtained unfolded chain, 3'-hairpin, cross-hairpin and incomplete 5'- or 3'-triplex (two G-triads and one GG pair). Importantly, we found out that the end-to-end distance of the molecule bearing the 3'-hairpin structure was the same as the one with incomplete 3'-triplex (Figure 7 in the main text). This observation is a reminder that structural interpretations based on just end-to-end distances measured in experiments without atomistic resolution may be ambiguous (see the main text for further details).

*2+2 antiparallel GQ*. Unlike in the previous systems, the force acting on 143D is directed diagonally across the 5'-quartet. Under *fast pulling* conditions, formation of 3'-triplex by strand detachment from opened GQ was observed twice, which is similar to 2GKU. In addition, we saw unzipping of the first and last G from the G-stem in the remaining simulation. In *slow zig-zag pulling* simulations, 3'-triplex was formed in all the simulations, but by unzipping of the first strand. Unfolding then proceeded in different ways, so that we obtained cross-triplex, imperfect 3'-triplex and 3'-hairpin. *Very slow zig-zag* and *very slow pulling* simulations revealed another general trend as unzipping of individual bases from the first and last strands happened in all three independent simulations. The unzipping under *very slow* *pulling* conditions ended up by unfolding of four bases from the GQ stem, i.e., either one quartet with two stacked diagonal G:G pairs or an incomplete 3'-triplex remained. *Very slow zig-zag* unfolding continued further and eventually led to formation of 3'-hairpin or unstable diagonal hairpins.

143D$_{noloop}$ simulations, i.e. 143D without the diagonal loop, were designed to see if there was any effect of the diagonal loop, stacked on the terminal quartet, on the unfolding of the 143D model. In all pulling schemes, we consistently observed unzipping of individual bases from the first and last strand, which was followed by formation of triplex and hairpin intermediates under both *slow zig-zag* and *very slow zig-zag pulling* conditions. This behavior was slightly different from *fast pulling* of 143D, where opening was mostly observed, but matches well with results from simulations of 143D model under the same *slow zig-zag*, *very slow zig-zag* and *very slow* pulling setups. In some simulations of the 143D$_{noloop}$ model, we observed flipping of the now terminal 5'-G into the *syn* conformation; the flip happened after the major unfolding events and thus it did not affect the main results. Such *anti/syn* flips of terminal residues are common due to the fact that *syn* states are stabilized by the formation of an intramolecular H-bond between the terminal 5'-OH group and N3 nitrogen of the nucleobase (1).

143D$_{syn}$ model had a strand *syn*/*anti* pattern allowing for strand slippage, however, the direction of the pulling force across the terminal quartet was thought not to promote this unfolding mechanism. Our expectations met with the reality, as unzipping of individual bases was the most common unfolding mechanisms yet again. Nevertheless, the system behave similarly, but not exactly like the 143D model. In *fast pulling* simulations, we observed the formation of 3'-triplex twice but via different mechanism, i.e., by unzipping and not by opening with strand detachment as in the 143D model. Under *slow zig-zag pulling* simulations, base unzipping led to formation of incomplete 5'- and 3'-triplex, while the same conditions in 143D led via G-triplexes to further unfolding. *Very slow zig-zag pulling* led to most extensive unfolding, 3'-hairpin was formed twice and 5'-hairpin, which further unfolded, once. Diagonal hairpin, like in the 143D model, was not found. 143D has both the first and last GpG dinucleotide step, which is disrupted by unzipping, in the same *anti*-*syn* order, while the 143D$_{syn}$ model has the first step *anti*-*anti* and the last one *syn*-*syn*. Therefore, one might expect that *syn*-*syn* step, which is thought to be less stable (1,2), would unzip preferentially; however, this was not the case in our simulations.

In 143D$_{loop\text{-}pull}$ simulations, the force acted on the GQ in the same relative direction as on the parallel and hybrid topologies (Figure 2). The most common outcome under *fast pulling* and *slow zig-zag pulling* conditions was formation of diagonal hairpin by an unzipping and/or detaching of the first and last strand, followed by unfolding of the diagonal hairpin and spontaneous refolding of 3'-hairpin. *Very slow zig-zag pulling* simulations offered the most complex unfolding dynamics including unzipping, strand detachment, cross-like GQ or G-triplex formation. Description of all those unfolding processes can be simplified as unbinding of the first strand, followed by the second strand, so that 3'-hairpin remained. Notice that the final unfolding product was the same in all three pulling schemes. However, unfolding pathways identified during the *very slow zig-zag pulling* protocol were fundamentally different from those observed in the *fast pulling* and *slow zig-zag pulling* schemes.

143D$_{syn_loop\text{-}pull}$ model was under the same pulling force direction as the 143D$_{loop\text{-}pull}$ model, but its modified *syn*/*anti* pattern theoretically allowed strand slippage, as in the 1KF1 model. *Fast pulling* setup provided diverse outcomes; unzipping of bases in the first and third strand, GQ opening, and incomplete strand slippage leading to spiral structures. *Slow zig-zag pulling* showed a similar picture, including spiral intermediates, extended by additional unfolding. The first strand was always ultimately removed and the simulations led to partially unfolded 3'-triplexes or 3'-hairpin structures. *Very slow zig-zag pulling* protocol offered the most complex scenarios, again. Apart from GQ division, all the described unfolding motions were observed. Unfolding reached the greatest extent in these simulations, leading to 3'-hairpins and several partial refolding events were also identified. We found similar spiral intermediates as observed during fast and *slow zig-zag pulling* simulations and also extra events of proper strand slippage by one level. We suspect that complete strand slippage is unlikely on this time scale (few μs), because as the distance between the connected strand ends increases during the slippage, so has to extend the lateral loop backbone, whose flexibility is limited by interaction of the loop bases with the G-stem. In comparison, the distance between connected ends during strand slippage decreases in the 1KF1 model, so the propeller loop does not need to be extended (strand slippage in the other direction, in which the distance would increase, has not been observed). In overall, $143D_{syn\_loop\text{-}pull}$ system behaved like a mixture of 1KF1 and 2GKU models.

### SUPPLEMENTARY TABLES

**Table S1**: Detailed pulling parameters for *fast pulling* simulations of all GQ models.[a]

| System | $\kappa_0$ [pN·nm$^{-1}$] | $x_0$ [nm] | $x_\tau$ [nm] | $\tau$ [ns] | $\upsilon$ [nm/ns] |
|---|---|---|---|---|---|
| 1KF1 | 1660 | 2.0 | 6.0 | 6 | 0.667 |
| 1KF1$_{syn}$ | 1660 | 2.0 | 6.0 | 6 | 0.667 |
| 2GKU | 1660 | 2.0 | 6.0 | 6 | 0.667 |
| 143D | 1660 | 2.0 | 6.0 | 6 | 0.667 |
| 143D$_{syn}$ | 1660 | 2.0 | 6.0 | 6 | 0.667 |
| 143D$_{\mathrm{noloop}}$ | 1660 | 2.4 | 6.0 | 6 | 0.600 |
| 143D$_{\mathrm{loop\text{-}pull}}$ | 1660 | 2.1 | 6.0 | 6 | 0.650 |
| 143D$_{syn_\mathrm{loop\text{-}pull}}$ | 1660 | 1.7 | 6.0 | 6 | 0.717 |

[a] We ran three independent simulations (identical pulling settings) for each GQ model characterized by force constant ($\kappa_0$), initial ($x_0$) and final ($x_\tau$) distances between pulling centers, simulation timescale ($\tau$), and corresponding pulling velocity ($\upsilon$).

**Table S2**. Detailed pulling parameters for *slow zig-zag pulling* simulations of all GQ models.[a]

| System | Phase I | | | | | Phase II | | | | |
|---|---|---|---|---|---|---|---|---|---|---|
| | $\kappa_0$ [pN·nm$^{-1}$] | $x_0$ [nm] | $x_\tau$ [nm] | $\tau$ [ns] | $\upsilon$ [nm/ns] | $\kappa_0$ [pN·nm$^{-1}$] | $x_0$ [nm] | $x_\tau$ [nm] | $\tau$ [ns] | $\upsilon$ [nm/ns] |
| 1KF1 | 150 | 2.0 | 6.0 | 100 | 0.040 | 150 | 6.0 | 7.5 | 100 | 0.015 |
| 1KF1$_{syn}$ | 150 | 2.0 | 6.0 | 100 | 0.040 | 150 | 6.0 | 7.5 | 100 | 0.015 |
| 2GKU | 150 | 2.0 | 6.0 | 100 | 0.040 | 150 | 6.0 | 9.0 | 200 | 0.015 |
| 143D | 150 | 2.0 | 6.0 | 100 | 0.040 | 150 | 6.0 | 9.0 | 200 | 0.015 |
| 143D$_{syn}$ | 150 | 2.0 | 6.0 | 100 | 0.040 | 150 | 4.0 | 9.0 | 200 | 0.025 |
| 143D$_{\mathrm{noloop}}$ | 150 | 2.4 | 6.0 | 100 | 0.036 | 150 | 6.0 | 9.0 | 200 | 0.015 |
| 143D$_{\mathrm{loop\text{-}pull}}$ | 150 | 2.1 | 6.0 | 100 | 0.039 | 150 | 6.0 | 9.0 | 200 | 0.015 |
| 143D$_{syn_\mathrm{loop\text{-}pull}}$ | 150 | 1.7 | 6.0 | 100 | 0.043 | 150 | 4.0 | 9.0 | 200 | 0.025 |

[a] We ran three independent simulations (identical pulling settings) for each GQ model. Pulling Phases I and II (see Methods in main text) are characterized by force constant ($\kappa_0$), initial ($x_0$) and final ($x_\tau$) distances between pulling centers, simulation timescale ($\tau$), and corresponding pulling velocity ($\upsilon$).

**Table S3**. Detailed pulling parameters for *very slow zig-zag pulling* simulations of all GQ models.[a]

| System | Phase I | | | | | Phase II | | | | |
|---|---|---|---|---|---|---|---|---|---|---|
| | $\kappa_0$ [pN·nm$^{-1}$] | $x_0$ [nm] | $x_\tau$ [nm] | $\tau$ [ns] | $\upsilon$ [nm/ns] | $\kappa_0$ [pN·nm$^{-1}$] | $x_0$ [nm] | $x_\tau$ [nm] | $\tau$ [ns] | $\upsilon$ [nm/ns] |
| 1KF1 | 150 | 2.0 | 6.0 | 1000 | 0.004 | 150 | 6.0 | 7.5 | 1000 | 0.002 |
| 1KF1$_{syn}$ | 150 | 2.0 | 6.0 | 1000 | 0.004 | 150 | 6.0 | 7.5 | 1000 | 0.002 |
| 2GKU | 150 | 2.0 | 6.0 | 1000 | 0.004 | 150 | 6.0 | 9.0 | 2000 | 0.002 |
| 143D | 150 | 2.0 | 6.0 | 1000 | 0.004 | 150 | 6.0 | 9.0 | 3000 | 0.001 |
| 143D$_{syn}$ | 150 | 2.0 | 6.0 | 1000 | 0.004 | 150 | 4.5 | 9.0 | 2000 | 0.002 |
| 143D$_{\mathrm{noloop}}$ | 150 | 2.4 | 6.0 | 1000 | 0.004 | 150 | 6.0 | 9.0 | 2000 | 0.002 |
| 143D$_{\mathrm{loop\text{-}pull}}$ | 150 | 2.1 | 6.0 | 1000 | 0.004 | 150 | 6.0 | 9.0 | 2000 | 0.002 |
| 143D$_{syn_\mathrm{loop\text{-}pull}}$ | 150 | 1.7 | 6.0 | 1000 | 0.004 | 150 | 4.5 | 7.5 | 1000 | 0.003 |

[a] We ran three independent simulations (identical pulling settings) for each GQ model. Pulling Phases I and II are characterized by force constant ($\kappa_0$), initial ($x_0$) and final ($x_\tau$) distances between pulling centers, simulation timescale ($\tau$), and corresponding pulling velocity ($\upsilon$).

**Table S4**. Detailed pulling parameters for *very slow pulling* simulations of three GQ models.[a]

| System | $\kappa_0$ [pN·nm$^{-1}$] | $x_0$ [nm] | $x_\tau$ [nm] | $\tau$ [ns] | $\upsilon$ [nm/ns] |
|---|---|---|---|---|---|
| 1KF1 | 150 | 2.0 | 8.0 | 1500 | 0.004 |
| 2GKU | 150 | 2.0 | 8.0 | 1500 | 0.004 |
| 143D | 150 | 2.0 | 8.0 | 1500 | 0.004 |

[a] We ran three independent simulations (identical pulling settings) for 1KF1, 2GKU and 143D GQ models. Simulations are characterized by force constant ($\kappa_0$), initial ($x_0$) and final ($x_\tau$) distances between pulling centers, simulation timescale ($\tau$), and corresponding pulling velocity ($\upsilon$).

**Table S5.** Outcome of simulations with the *fast pulling* setup.[a]

| System | Outcome | Figures |
|---|---|---|
| 1KF1 | run1: first strand slipped three levels in the 5'-direction (A) leading to 3'-triplex | S1B |
| | run2: spiral structure formed and GQ opened (A), last G unzipped (B), last strand slipped in the 3'-direction (C), first strand slipped in the 5'-direction (D), third and fourth strand slipped in the 3'-direction two levels (E), 5'-hairpin unfolded (F) | S1C |
| | run3: last strand slipped three levels in the 3'-direction (A) leading to 5'-triplex | S1B |
| $1KF1_{syn}$ | run1: GQ opening and formation of cross-like GQ (A), last G unzipped (B) | S2B |
| | run2: GQ opening and formation of cross-like GQ (A), last G unzipped (B), last strand detached (C), so that 5'-cross-triplex remained | S2B |
| | run3: GQ opening and formation of cross-like GQ (A), last G unzipped (B), last strand detached (C), and 5'-triplex refolded | S2C |
| 2GKU | run1: GQ opening and formation of cross-like GQ (A) | S3B |
| | run2: GQ opening and formation of cross-like GQ (A) | S3B |
| | run3: GQ opening and formation of cross-like GQ (A), last (B) and first G unzipped (C) | S3B |
| 143D | run1: first (A) and second G unzipped (B) | S4B |
| | run2: first (A) and second G unzipped (B) | S4B |
| | run3: first (A) and last G unzipped (B) | S4B |
| $143D_{syn}$ | run1: first (A) and last (B) G unzipped; | S5B |
| | run2: first (A), and second G unzipped (B); | S5B |
| | run3: first (A), and second G unzipped (B) | S5B |
| $143D_{noloop}$ | run1: first (A) and last G unzipped (B) | S6B |
| | run2: first and last G unzipped (A) | S6B |
| | run3: first (A) and last G unzipped (B) | S6B |
| $143D_{loop\text{-}pull}$ | run1: first G unzipped (A), GQ opening (B); | S7B |
| | run2 first G unzipped (A), GQ opening (B), first strand rotated into cross-orientation (C); | S7B |
| | run3: first G unzipped (A), GQ opening (B), last G of third strand unzipped (C) | S7C |
| $143D_{syn\_loop\text{-}pull}$ | run1: GQ opening (A), first (B) and last G of the third strand unzipped (C); | S8B |
| | run2: first strand slipped in the 5'-driection to form spiral structure (A), last G of third strand unzipped (B), GQ opening (C), first G unzipped (D); | S8C |
| | run3: GQ opening (A), first strand rotated into cross-orientation (B) | S8C |

[a] The capital letters (A, B, C…) denote the events and structural snapshots in the respective Supplementary Figures.

**Table S6.** Outcome of simulations with the *slow zig-zag pulling* setup.[a]

| System | Outcome | Figures |
|---|---|---|
| 1KF1 | run1: first strand rotated into cross-orientation to the last strand (A), then triple strand slippage of the first strand in the 5'-direction (B), so that 3'-hairpin remained; | S9B |
| | run2: spiral structure formed (A), last G unzipped (B), GQ opening and rotation into cross-like GQ (C), triple strand slippage of the last strand in the 3'-direction, (**II**) the other strands reformed 5'-triplex (D), last G of third strand unzipped (E); | S9C |
| | run3: third and last strand slipped in the 3'-direction (A), last strand rotated into cross-orientation to the first strand (B), last strand slipped in the 3'-direction (C), first G unzipped (D), third strand detached from second strand (E), so that 5'-hairpin and 3'-cross-hairpin are formed, (**II**) the latter unfolds by strand detachment (F), the former eventually turns into cross-hairpin | S9D |
| $1KF1_{syn}$ | run1: GQ opening and rotation into cross-like GQ (A), last (B), (**II**) first (C), second to last (D) and third to last G unzipped (E), 5'-triplex with two triplets refolded; | S10B |
| | run2: spiral structure formed (A), (**II**) last G unzipped (B), GQ opening and rotation into cross-like GQ (C), first G unzipped (D), first (E) and last strand detached (F), second G reattached (G), middle hairpin turned into cross-hairpin (H), third G reattached (I), third strand detached (J); | S10C |
| | run3: (**II**) first G unzipped (A), GQ opening and rotation of first strand (with first G of second strand stacked onto second G) into cross-like orientation (B), first strand detached (C), so that 3'-triplex with two triplets remained | S10D |
| 2GKU | run1: (**II**) GQ opening and rotation into cross-like GQ (A), first (B) and last G unzipped (C), division into two G-hairpins (D), 5'-hairpin unfolded (E), perturbed 3'-hairpin remained; | S11B |
| | run2: (**II**) last G unzipped (A), GQ opening and rotation into cross-like GQ (B), second to last G unzipped and 5'-triplex with one quartet refolded (C), third to last G unzipped (D); | S11C |
| | run3: GQ opening and rotation into cross-like GQ (A), (**II**) first G unzipped (B), structure divided into two hairpins (C), 5'-hairpin unfolded (D), so that 3'-hairpin remained | S11D |
| 143D | run1: (**II**) first (A), second (B) and third G unzipped (C), then rotation of second strand, resulting into cross-like triplex (D); | S12B |
| | run2: (**II**) first (A), second (B), last (C) and third G unzipped (D), so that 3'-triplex with two triplets remained; | S12C |
| | run3: (**II**) first (A), second (B), third (C) and (**III**) last G unzipped (D), second strand rotated to form a cross-triplex (E), then detached (F), resulting into imperfect 3'-hairpin | S12D |
| $143D_{syn}$ | run1: (**II**) first (A), last (B), second (C), and second to last G unzipped (D); | S13B |
| | run2: (**II**) last (A), first (B), second (C), and third G unzipped (D), second strand slipped in 5'-direction (E), resulting into imperfect 3'-G-triplex; | S13C |
| | run3: (**II**) first (A), last (B), second to last (C), and third to last G unzipped (D), resulting into imperfect 5'-G-triplex | S13D |
| $143D_{noloop}$ | run1: first (A), last (B), second (C) and (**II**) third G unzipped (D), then second strand detached, migrated to third strand (E) and rotated to form a cross-like 3'-triplex (F); | S14B |

| | | |
|---|---|---|
| | run2: first (A), last (B), second (C) and (**II**) third G unzipped (D), then second strand detached (E), so the structure was separated into parts, 3'-hairpin remained; | S14C |
| | run3: (**II**) first (A), last (B) and second to last G unzipped (C) | S14D |
| 143D$_{loop-pull}$ | run1: last G of third strand unzipped (A),* GQ opening (B), first strand rotated into cross position (C), second G of third strand unzipped (D), first and second G detached (E), (**II**) second strand rotated into cross position (F), third G unzipped (G), second strand detached (H), 3'-hairpin refolded (I); | S15B |
| | run2: (**II**) last G of third strand (A) and second G of third strand unzipped (B), first and second G detached (C), third G unzipped (D), second strand detached (E), 3'-hairpin refolded (F); | S15C |
| | run3: (**II**) first (A), second (B), and third G unzipped (C), then gradual opening between third and fourth strand, third strand rotated into cross orientation (D), then last G of third strand unzipped (E) and first and second G of third strand detached (F), leaving middle hairpin, which turned into cross-hairpin (G), third strand eventually gradually bound to fourth strand in cross orientation, so that cross-cross structure was formed | S15D |
| 143D$_{syn_loop-pull}$ | run1: (**II**) first and second G detached (A), third G rotated into cross orientation (B), last G of third strand (C) and third G unzipped (D), then last G of third strand incorporated back into 3'-triplex (E), and unzipped again (F), (**III**) first G of second strand unzipped (G), so that 3'-triplex with two triads remained; | S16B |
| | run2: (**II**) spiral structure formed (A), first G unzipped (B), second and third G detached (C), then third strand (only) slipped in the 3'-direction and rotated (D), so 3'-cross-triplex was formed; | S16C |
| | run3: (**II**) spiral structure formed (A), first G unzipped (B), GQ opening (C), last G of third strand unzipped and GQ closed the opening (D), second and third G detached (E), last G of third strand incorporated back into 3'-triplex (F), second strand rotates to form 3'-cross-triplex (G) | S16D |

[a] The capital letters (A, B, C…) denote the events and structural snapshots in the respective Supplementary Figures. Roman numerals II and III denote events that happened after the first external force drop, i.e. in phase II, and after reaching maximum allowed extension, i.e. phase III, respectively.
*T1 unzipped after the marked unfolding event

**Table S7.** Outcome of simulations with the *very slow zig-zag pulling* setup.[a]

| System | Outcome | Figures |
|---|---|---|
| 1KF1 | run1: last G unzipped (A), the last strand slipped in the 3’-direction (B) and got detached (C), (**II**) the resultant G-triplex turned into a cross-triplex by rotation of the third strand (D), the first strand gradually unzipped (E), the remaining cross-hairpin unfolded (F) | S17B |
| | run2: last strand slipped in the 3’-direction (A), first strand slipped in the 5’-direction (B), last strand unzipped (C), (**II**) the resultant G-triplex turned into cross-triplex by rotation of first strand (D), then briefly turned into symmetric triplex (E), cross-triplex was formed by rotation of second strand (F), reformation of G-triplex (G) was followed by strand slippage of the third strand in the 3’-direction (H), then cross-triplex reappeared by rotation of first strand (I), which was then detached (J), the remaining middle hairpin unfolded via cross-hairpin (K) | S17C |
| | run3: last G unzipped (A), followed by incomplete strand slippage resulting into the spiral structure (B), first G unzipped (C), the last strand completed the slippage in the 3’-direction (D) and (**II**) then detached (E), 5’-G-triplex unfolded and refolded several time by back-and-forth detaching and reattaching first strand (without first G), eventually whole third strand unzipped (F), the resultant 5’-G-hairpin unfolded by strand slippage (G) | S17D |
| 1KF1$_{syn}$ | run1: first G unzipped (A), GQ opening and rotation into cross-like GQ (B), (**II**) last strand gradually unzipped and cross-triplex formed (C), first strand and first G of second strand detached (first G of second strand was stacked onto second and third G of first strand) (D), remaining middle G-hairpin eventually unfolded (E) | S18B |
| | run2: first (A) and last G unzipped (B), cross-like GQ formed (C), (**II**) second to last G unzipped (D), first strand and first G of second strand detached (E), third to last G unzipped (F), resultant middle G-hairpin unfolded (G) | S18C |
| | run3: first G unzipped (A), GQ opening and rotation into cross-like GQ (B) first strand and first G of second strand detached, second strand rotated back to form G-triplex (C), (**II**) first G of the second strand unzipped, last strand gradually unzipped (D), resultant middle G-hairpin H-bonded with last G of the first strand | S18D |
| 2GKU | run1: GQ opening (A), last G unzipped (B), first strand gradually unzipped (C), (**II**) last G incorporated back into the 3’-G-triplex (D), which later turned into a cross-triplex by rotation of second strand (E), eventually second strand detached (F) and 3’-G-hairpin remained | S19B |
| | run2: first G unzipped (A), GQ opening and rotation into cross-like GQ (B), (**II**) first strand unzipped and cross-triplex formed (C), last two Gs unzipped (D), middle cross-hairpin with first G of the last strand remained | S19C |
| | run3: first G unzipped (A), GQ opening and rotation into cross-like GQ (B), first strand detached and 3’-triplex remained (C), then (**II**) back-and-forth transitions to cross-like triplex (D), first G of the second strand unzipped (E) | S19D |
| 143D | run1: (**II**) first (A) and second G unzipped (B), last (C) and second to last G unzipped (D), third G unzipped (E), resultant diagonal hairpin gradually unfolded | S20B |
| | run2: (**II**) first and last G unzipped (A), second G unzipped (B), third G unzipped (C), then second strand slipped by one level in the 5’-direction to form a misfolded G-triplex (*t*WH pairing; D) and eventually detached (E), so that 3’-G-hairpin remained | S20C |
| | run3: (**II**) first (A) and last G unzipped (B), second G unzipped (C), second to last G unzipped (D), third G unzipped (E), remaining structure gradually unfolded | S20D |

| | | |
|---|---|---|
| $143D_{syn}$ | run1: first (A), (**II**) second (B) and third G unzipped (C), then second strand detached (D), so that 3'-hairpin remained | S21B |
| | run2: (**II**) last (A), first (B), second (C) and third G unzipped (D), second strand detached (E), then bound to third strand to form a cross-triplex (F) with brief visits of G-triplex, (**III**) eventually second strand detached (G), so that incomplete 3'-hairpin remained | S21C |
| | run3: (**II**) first (A), last (B), second to last (C), third to last G unzipped (D), third strand detached (E), resultant 5'-hairpin unfolded | S21D |
| $143D_{noloop}$ | run1: first and last G unzipped (A), (**II**) second G unzipped (B), third G unzipped (C), symmetric triplex formed (D), then second strand unbound from last strand and bound to third strand to form a new G-triplex (E), (**III**) first G of second strand unzipped (F), rotation of second strand to form a cross-like triplex (G), eventually second strand detached and the two molecule parts got separated (H); the 5'-G reversibly visited the *syn* conformation | S22B |
| | run2: last (A) and first G unzipped (B), (**II**) second G unzipped (C), second to last G unzipped (D), third G unzipped (E), the remainder unfolded and the two molecule parts got separated (F) | S22C |
| | run3: first (A) and last G unzipped (B), then (**II**) 5'-G flipped to *syn*, second (C), second to last (D), third (E), and (**III**) third to last G unzipped (F), so middle (diagonal) hairpin remained | S22D |
| $143D_{loop\text{-}pull}$ | run1: (**II**) first G unzipped (A), GQ opening (B), first strand detached (C), second strand detached (D), so that 3'-hairpin remained | S23B |
| | run2: third G of the third strand unzipped (A), first G unzipped (B), (**II**) rotation of strands into cross-like GQ (C), then second (D) and third G unzipped (E), second strand detached (F), so that perturbed 3'-hairpin remained | S23C |
| | run3: third (A) and second G of the third strand unzipped (B), first strand detached (C), (**II**) 3'-triplex with two G-triplets refolded (D), second strand detached (E), so that perturbed 3'-hairpin remained | S23D |
| $143D_{syn\_loop\text{-}pull}$ | run1: spiral structure formed (A), GQ opening (B) followed by strand rotation to cross-like GQ, (**II**) last G of third strand unzipped (C), spiral structure then partially refolded, second G of the third strand unzipped (D) and 5'-triplex with one quartet refolded, spiral structure formed (E), first G of the third strand slipped in the 3'-direction (F), first G of the last strand unzipped (G), first G of the third strand unzipped and fourth strand rotated into cross-triplex (H), first and second G slipped in the 5'-direction (I), last strand detached (J), 5'-G-hairpin gradually unfolded | S24B |
| | run2: (**II**) spiral structure formed (A), last G of the third strand pulled out (B), GQ opening (C), third strand detached (D), first strand detached (E), 3'-triplex refolded (F), second strand detached (G), 3'-hairpin began to unfold | S24C |
| | run3: spiral structure formed (A), then (**II**) first (B), second (C) and third G unzipped (D) and 3'-triplex refolded, second strand detached (E), so that 3'-G-hairpin remained | S24D |

[a] The capital letters (A, B, C…) denote the events and structural snapshots in the respective Supplementary Figures. Roman numerals II and III denote events that happened after the first external force drop, i.e. in phase II, and after reaching maximum allowed extension, i.e. phase III, respectively.

**Table S8.** Outcome of simulations with the *very slow pulling* setup.[a]

| System | Outcome | Figures |
|---|---|---|
| 1KF1 | run 1: GQ opening (A), last G unzipped + GQ refolded (B), last G stacked under third strand (C), GQ opening between third and fourth strand (D), GQ reformed (E), quick opening and reformation (F), last strand slipped down one level (G), last G unstacked (H), first G unzipped (I), spiral 1-4 formed (J), opening (K), last strand slipped one level downwards (L), last strand detached (M), third strand slipped one level downwards (N), third strand slipped one level downwards (O), third strand detached (P), incomplete 5'-G-hairpin remained | S25B |
| | run2: last G unzipped (A), third+last strand slipped down one level (B), first G unzipped (C), last strand slipped one level downwards (D), second to last G detached (E), last strand detached (F), incomplete 5'-G-triplex remained | S25C |
| | run3: spiral structure (A), first strand slipped upwards (B), first G unzipped (C), last G unzipped (D), first strand slipped upwards (E), first strand detached (F), last G reattached to reform a perfect G-triplex (G), first G of second strand unzipped (H), last G slipped downwards one level (I), last strand slipped one level downwards (J), second strand slipped one level downwards (K), last strand detached (L), second strand slipped one level upwards (M), incomplete middle G-hairpin remained | S25D |
| 2GKU | run 1: GQ opening and rotation into cross-like GQ (A), first G unzipped (B), first G of second strand detached + second G unzipped (lever to detach first G of second strand) (C), last G unzipped (D), first strand + second G of second strand detached (E), last G rebound (F), last G of second strand unzipped and 3'-hairpin remained (G), last G unzipped (H), second to last G unzipped (I), last strand unzipped (J), attempts to refold 3'-hairpin until the end of the simulation, up to two G:G pairs formed | S26B |
| | run2: GQ opening and rotation into cross-like GQ (A), last G unzipped (B), second to last G unzipped (C), fist G unzipped (D), last strand unzipped (E) and third strand derotated to form an incomplete 5'-triplex, last G of third strand detached (F), imperfect 5'-triplex remained | S26C |
| | run3: GQ opening and rotation into cross-like GQ (A), first G unzipped (B), last G unzipped (C), first strand + first G of second strand detached (D), imperfect 3'-triplex remained | S26D |
| 143D | run1: first G unzipped (A), second G unzipped (B), last G unzipped (C), third G unzipped (D), so that an incomplete 3'-triplex remained | S27B |
| | run2: first G unzipped (A), last G unzipped (B), second G unzipped (C), second to last G unzipped (D), so that a quartet + diagonal hairpin remained – *had three cations in the "channel"* | S27C |
| | run3: first G unzipped (A), last G unzipped (B), second to last G unzipped (C), second G unzipped (D), so that a quartet + diagonal hairpin remained – *had three cations in the "channel" (at different sites)* | S27D |

[a] The capital letters (A, B, C…) denote the events and structural snapshots in the respective Supplementary Figures.

**Table S9.** Events counts in each phase of the simulations with the *slow and very slow zig-zag pulling* schemes.[a]

| Phase | 1KF1 | $1KF1_{syn}$ | 2GKU | 143D | $143D_{syn}$ | $143D_{noloop}$ | $143D_{loop-pull}$ | $143D_{syn\_loop-pull}$ |
|---|---|---|---|---|---|---|---|---|
| *Slow zig-zag pulling* [b] | | | | | | | | |
| I | 11 | 3 | 1 | | | 4 | 5 | |
| II | 2 | 15 | 12 | 11 | 13 | 11 | 17 | 17 |
| III | | | | 3 | | | | 1 |
| *Very slow zig-zag pulling* [c] | | | | | | | | |
| I | 11 | 8 | 8 | | 1 | 5 | 5 | 3 |
| II | 13 | 8 | 7 | 15 | 14 | 12 | 10 | 20 |
| III | | | | | 1 | 4 | | |

[a] Events from three independent simulations are counted together for each GQ model.

[b] Phase I: 0-100 ns; Phase II: 100 ns – until box limit reached (or simulation end reached, if box limit not reached); Phase III: molecule kept extended at max distance allowed by the box

[c] Phase I: 0-1000 ns; Phase II: 1000 ns – until box limit reached (or simulation end reached, if box limit not reached); Phase III: molecule kept extended at max distance allowed by the box

#### SUPPLEMENTARY FIGURES

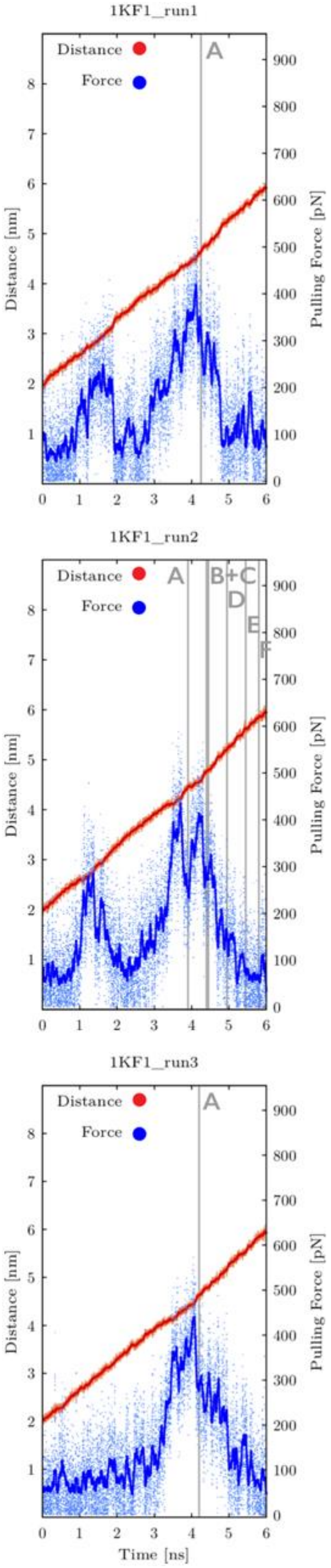

**Figure S1A:** Time evolution of distance between pulling centers and pulling force during three independent *fast pulling* simulations of 1KF1 GQ system. Snapshots were saved every 0.6 ps and plots are showing both instantaneous values (orange and light-blue dots for distance and force, respectively) and smoothing, i.e., averaging over 100 consecutive snapshots (red and blue lines for distance and force, respectively). Main structural events are highlighted as grey vertical lines with labels (capital letters). See Figures S1B and S1C for inspection of structures corresponding to main structural events. Note that first major drops of the pulling force before the GQ unfolding event "A" are connected with repositioning of terminal T residues.

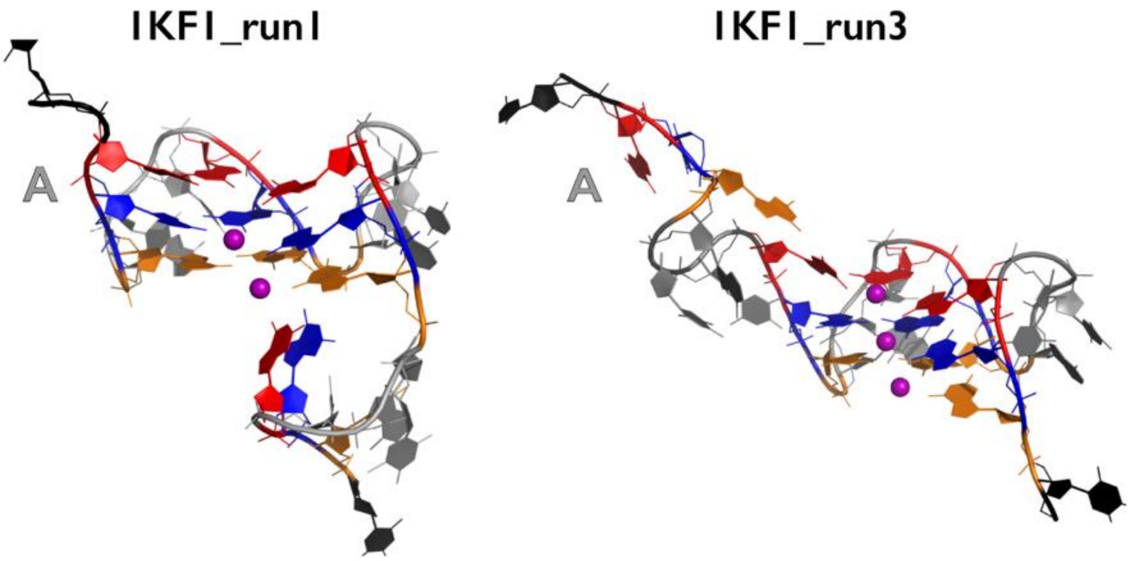

**Figure S1B:** Most important structural events during first and third independent *fast pulling* simulations of 1KF1 GQ system. G residues from first (5'-end), second and third quartet are highlighted in orange, blue and red, respectively. Pulling centers, i.e., either both terminal T residues or one terminal and other T residue from the loop (in some structures, see Methods in the main text for details), are shown in black. Remaining DNA residues are in gray and channel $K^+$ ions are shown as purple spheres. H-atoms and water molecules are not shown for clarity.

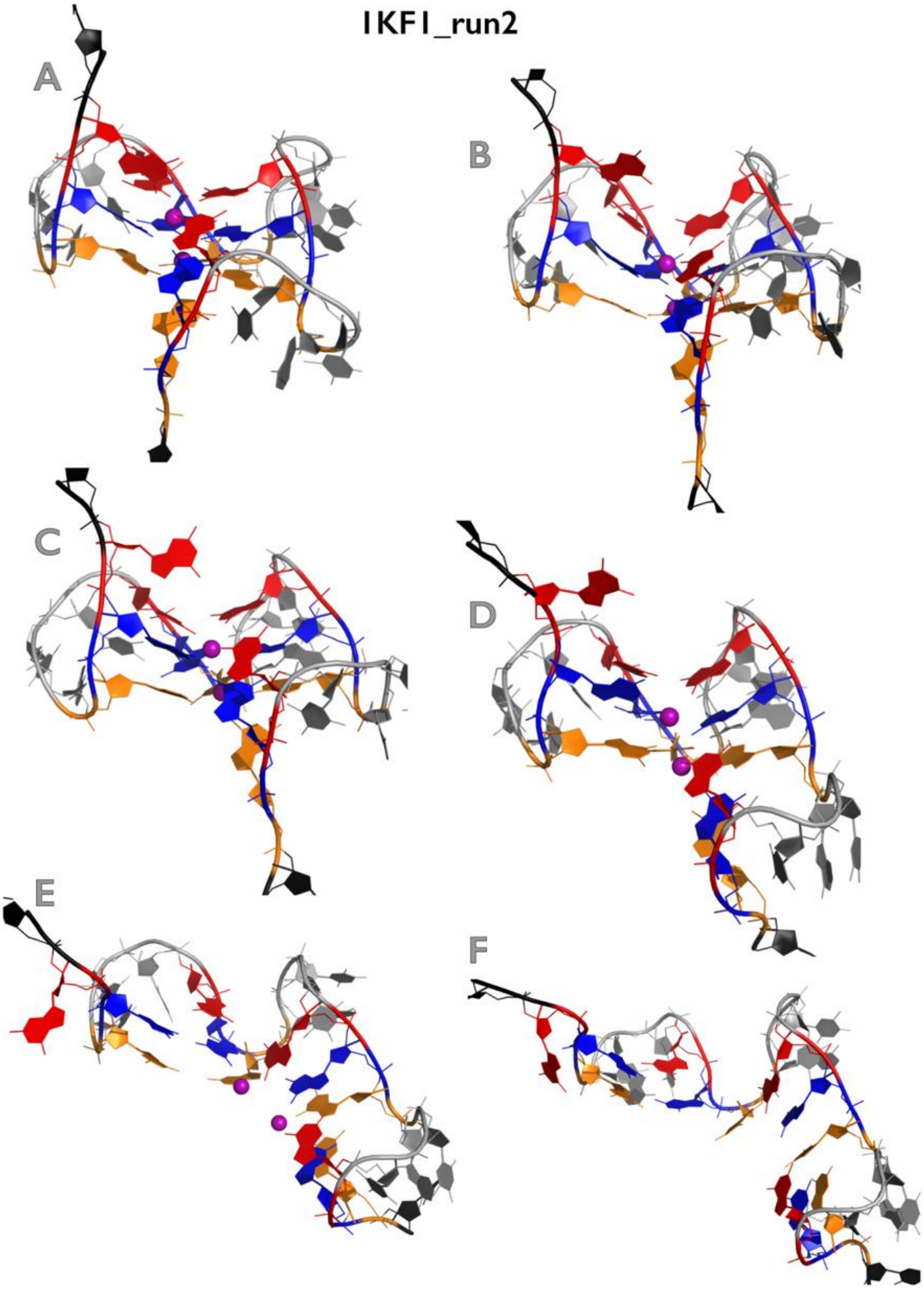

**Figure S1C:** Most important structural events during second independent *fast pulling* simulation of 1KF1 GQ system. See legend of Figure S1B for more details.

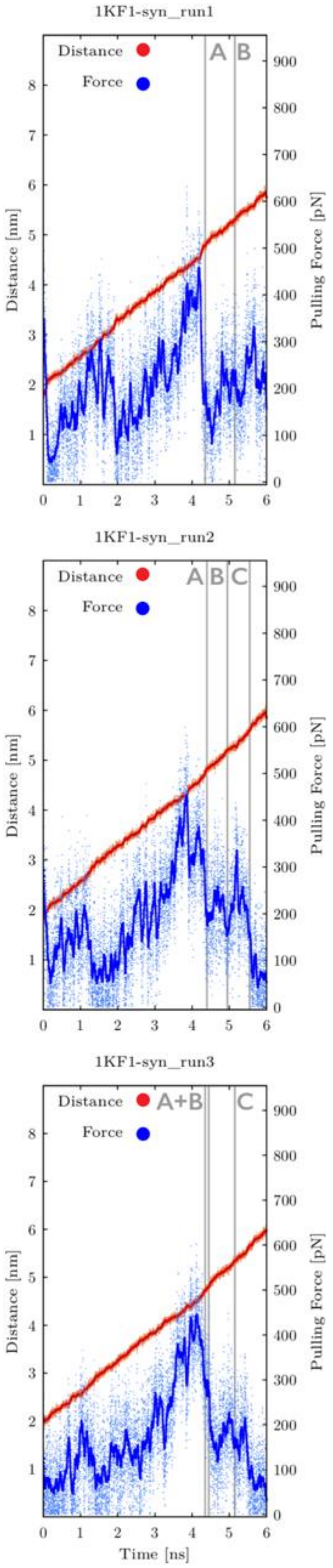

**Figure S2A:** Time evolution of distance between pulling centers and pulling force during three independent *fast pulling* simulations of $1KF1_{syn}$ GQ system (see legend of Figure S1A for more details). See Figures S2B and S2C for inspection of structures corresponding to main structural events.

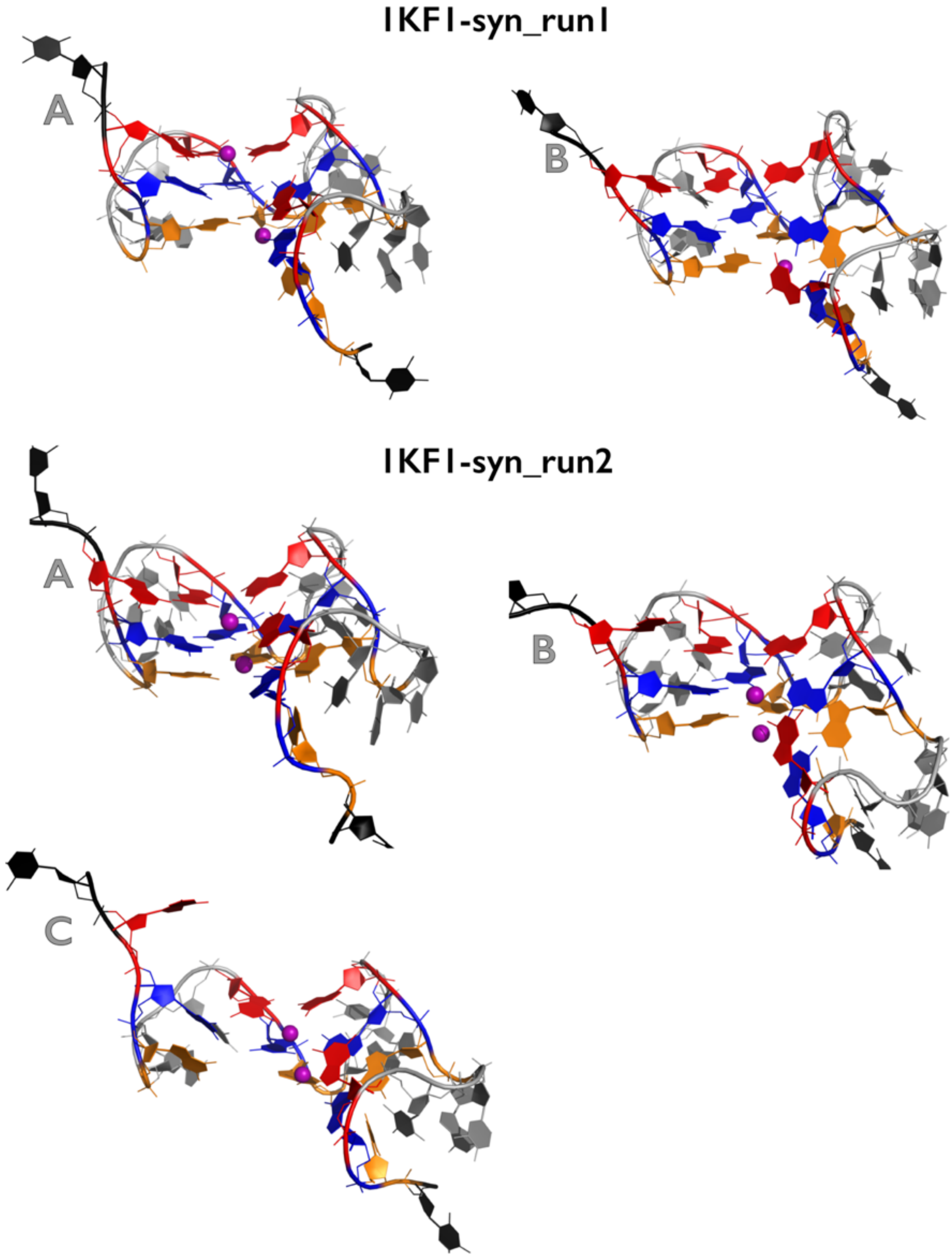

**Figure S2B:** Most important structural events during first and second independent *fast pulling* simulations of 1KF1$_{syn}$ GQ system. See legend of Figure S1B for more details.

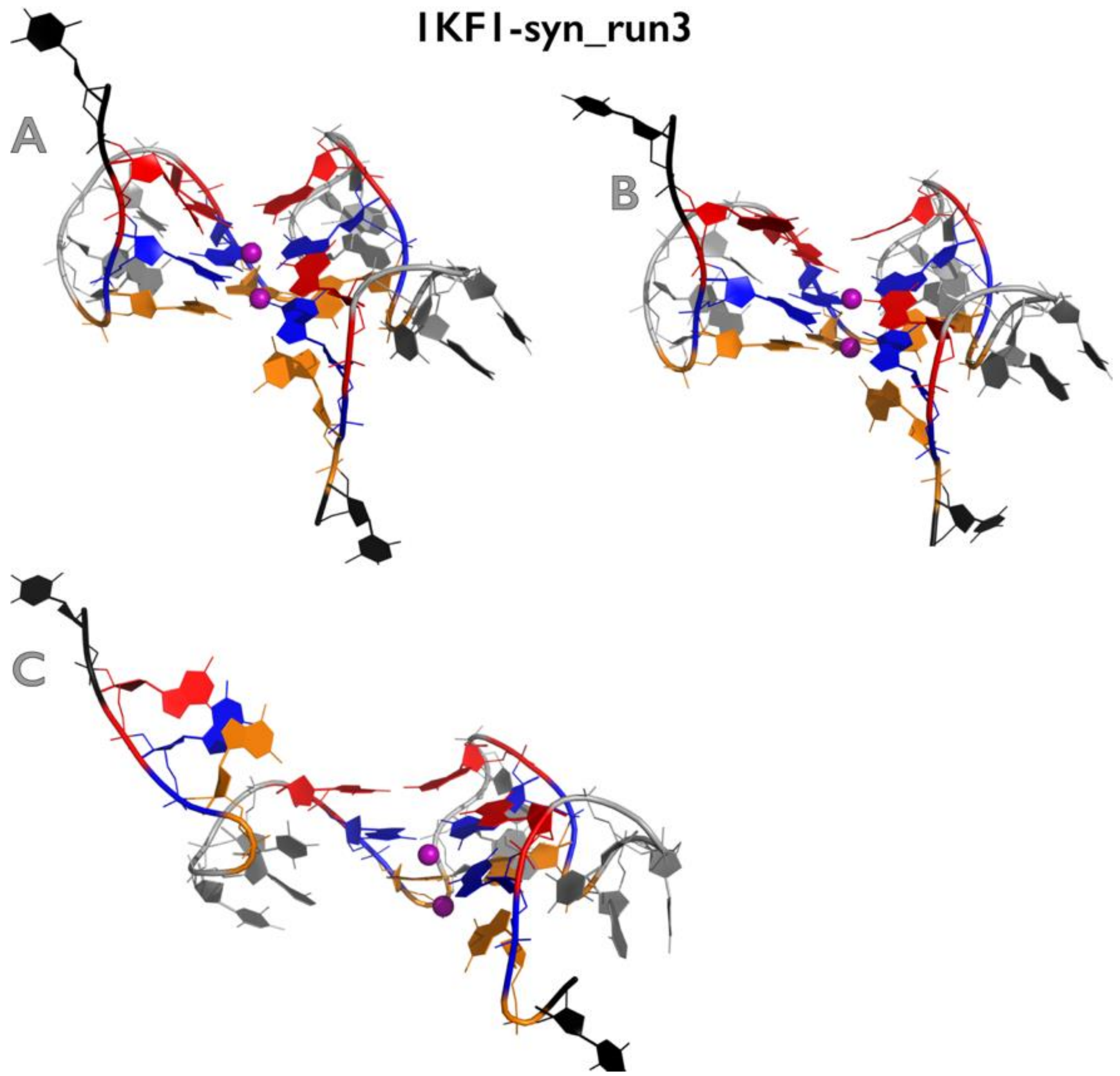

**Figure S2C:** Most important structural events during third independent fast pulling simulation of 1KF1$_{syn}$ GQ system. See legend of Figure S1B for more details.

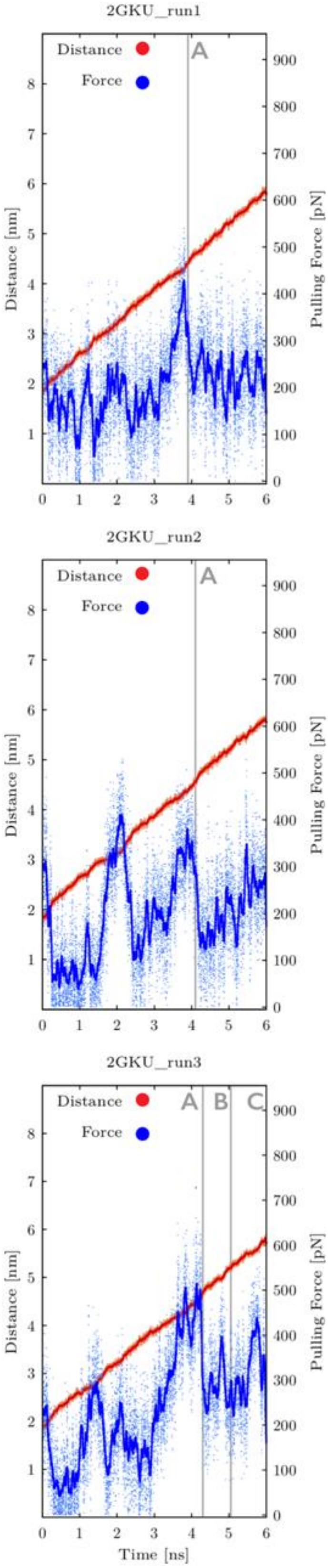

**Figure S3A:** Time evolution of distance between pulling centers and pulling force during three independent *fast pulling* simulations of 2GKU GQ system (see legend of Figure S1A for more details). See Figure S3B for inspection of structures corresponding to main structural events.

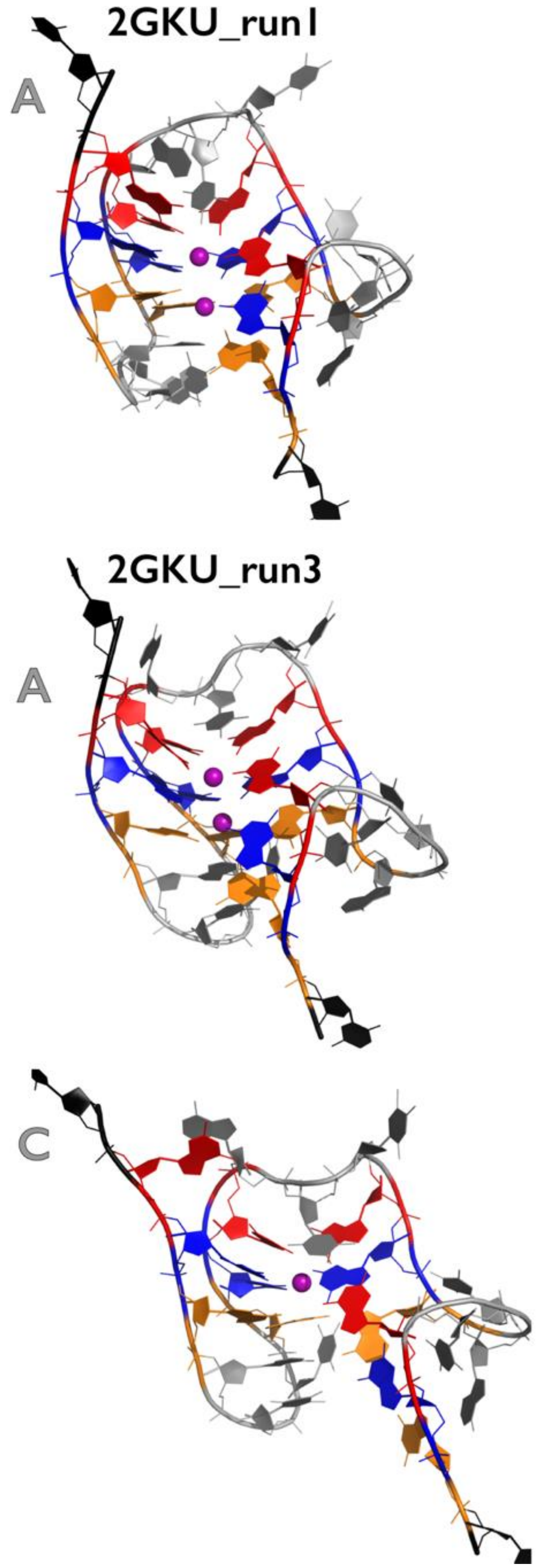

**Figure S3B:** Most important structural events during three independent *fast pulling* simulations of 2GKU GQ system. See legend of Figure S1B for more details.

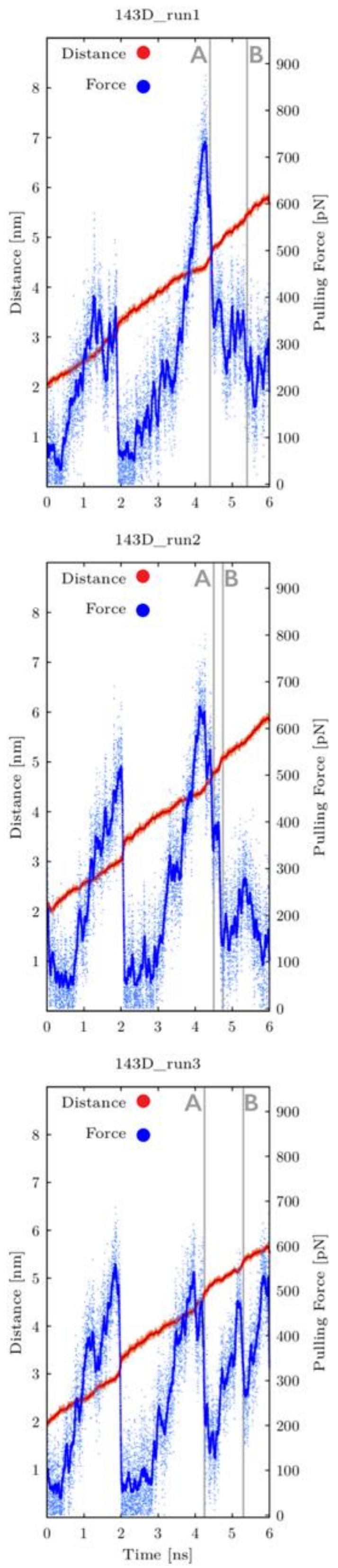

**Figure S4A:** Time evolution of distance between pulling centers and pulling force during three independent *fast pulling* simulations of 143D GQ system (see legend of Figure S1A for more details). See Figure S4B for inspection of structures corresponding to main structural events.

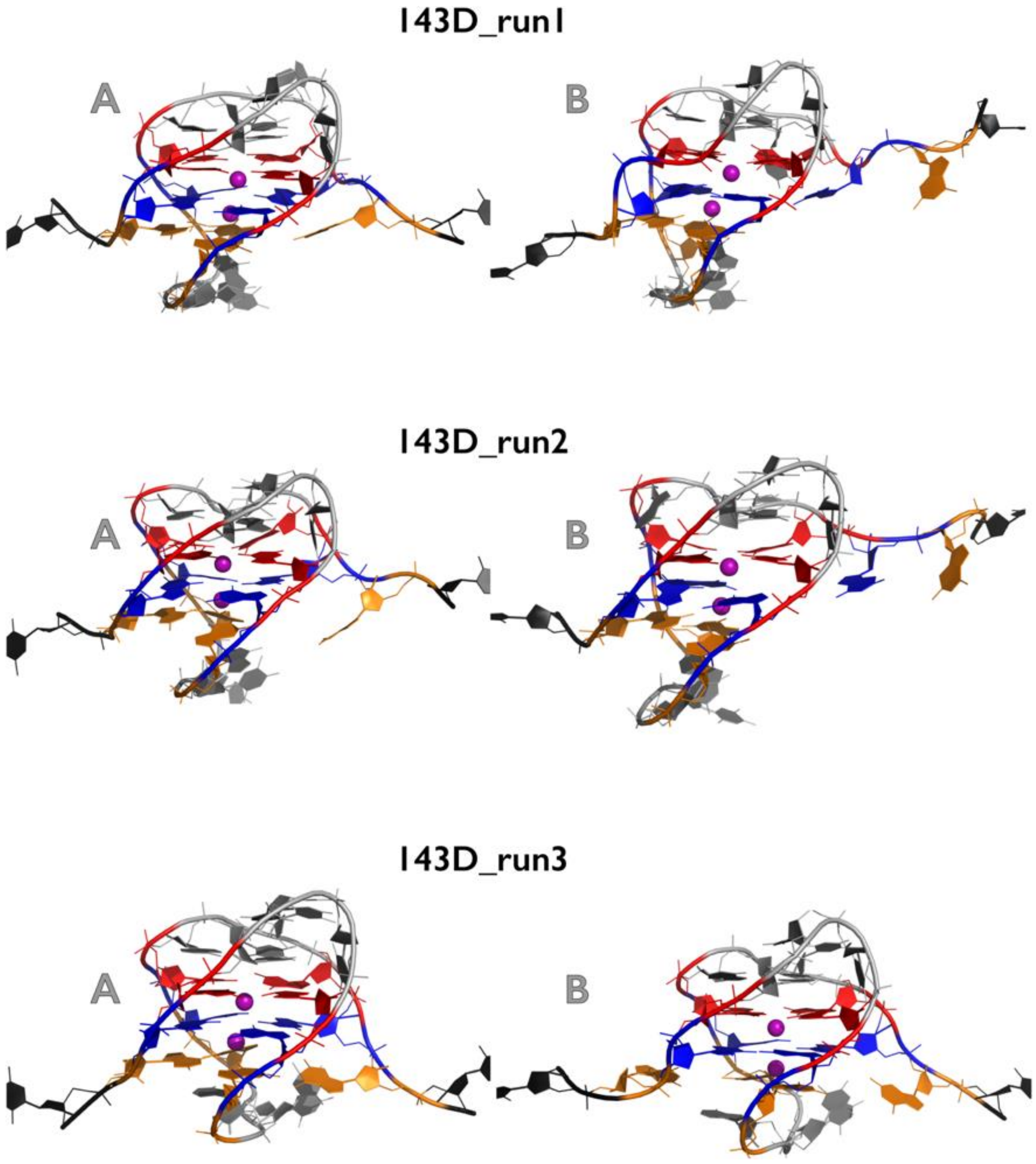

**Figure S4B:** Most important structural events during three independent *fast pulling* simulations of 143D GQ system. See legend of Figure S1B for more details.

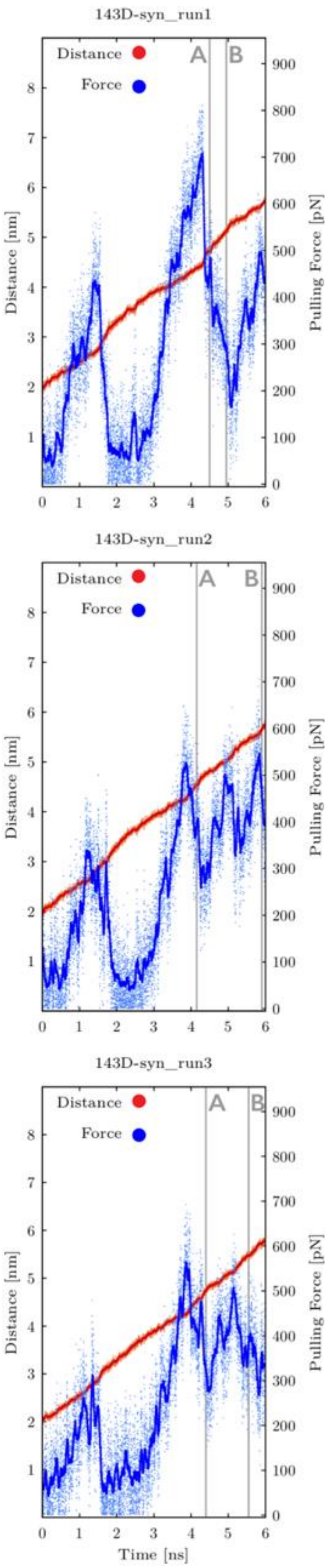

**Figure S5A:** Time evolution of distance between pulling centers and pulling force during three independent *fast pulling* simulations of 143D$_{syn}$ GQ system (see legend of Figure S1A for more details). See Figure S5B for inspection of structures corresponding to main structural events.

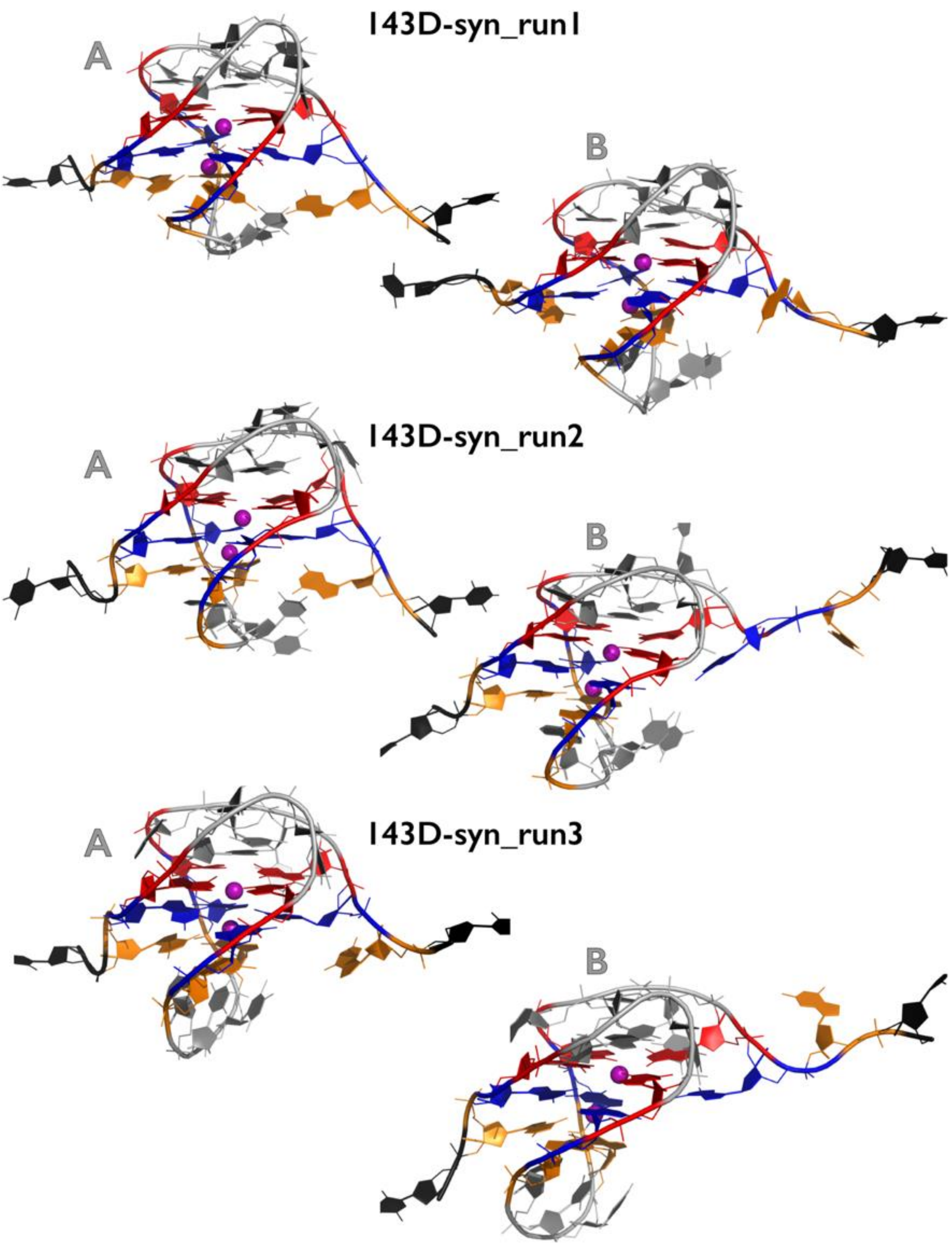

**Figure S5B:** Most important structural events during three independent *fast pulling* simulations of 143D$_{syn}$ GQ system. See legend of Figure S1B for more details.

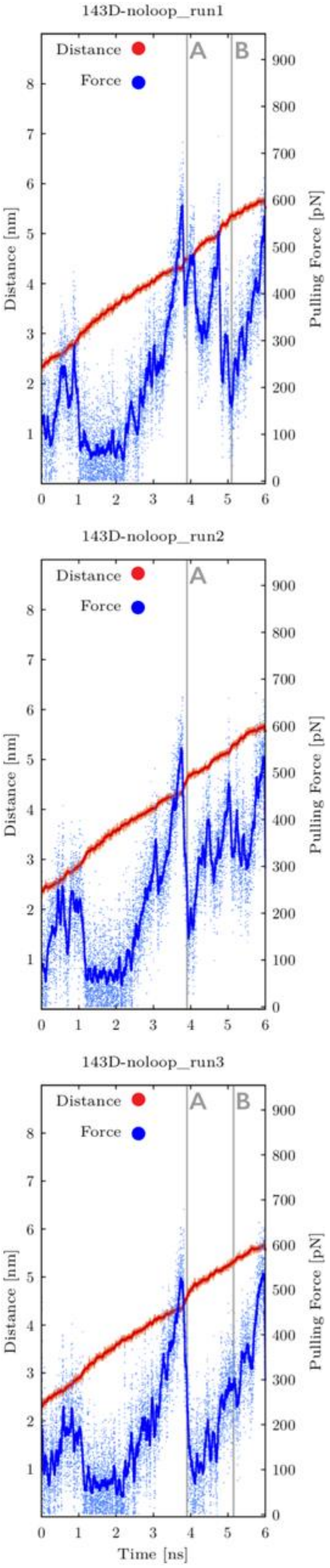

**Figure S6A:** Time evolution of distance between pulling centers and pulling force during three independent *fast pulling* simulations of $143D_{noloop}$ GQ system (see legend of Figure S1A for more details). See Figure S6B for inspection of structures corresponding to main structural events.

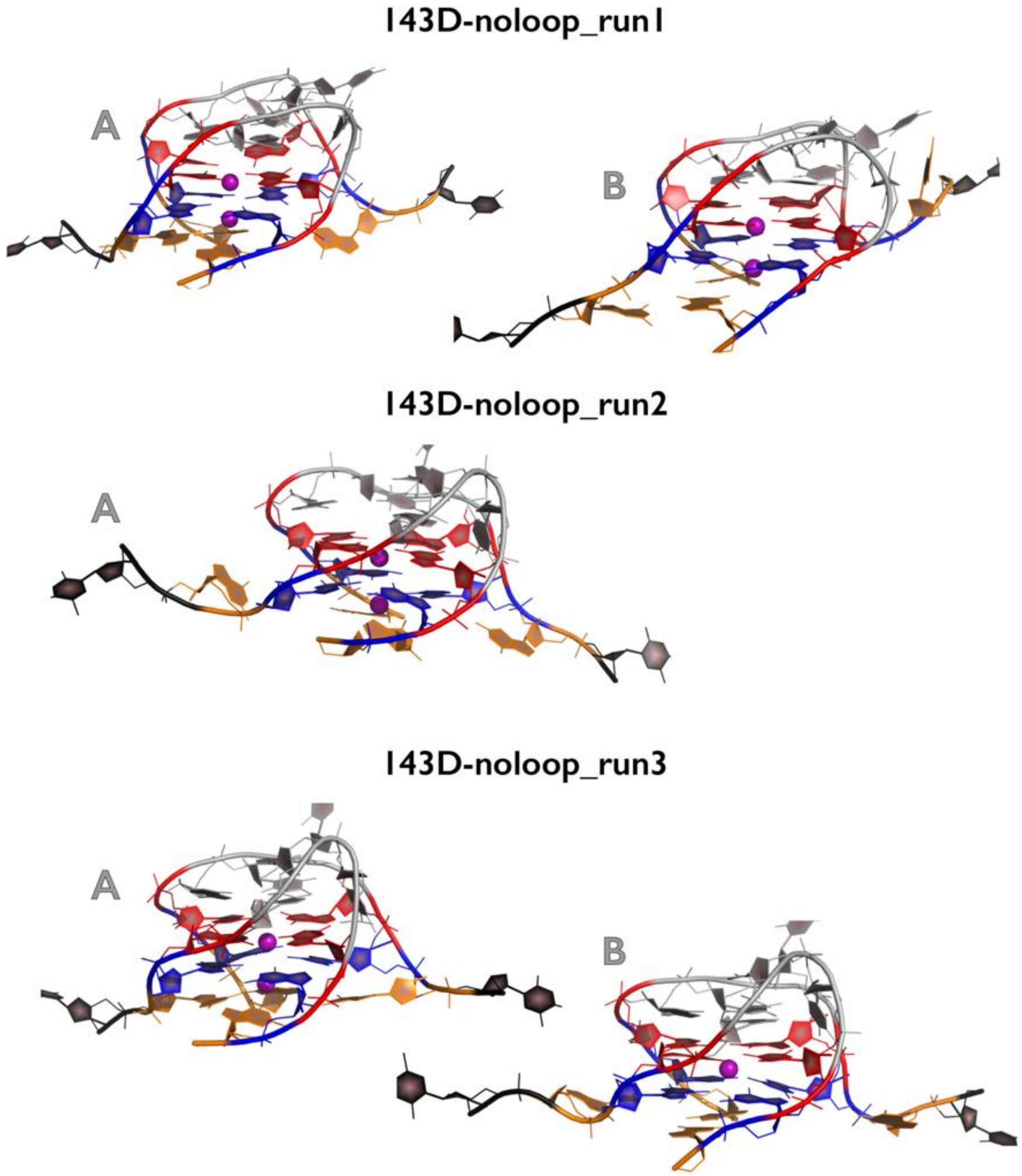

**Figure S6B:** Most important structural events during three independent *fast pulling* simulations of $143D_{noloop}$ GQ system. See legend of Figure S1B for more details.

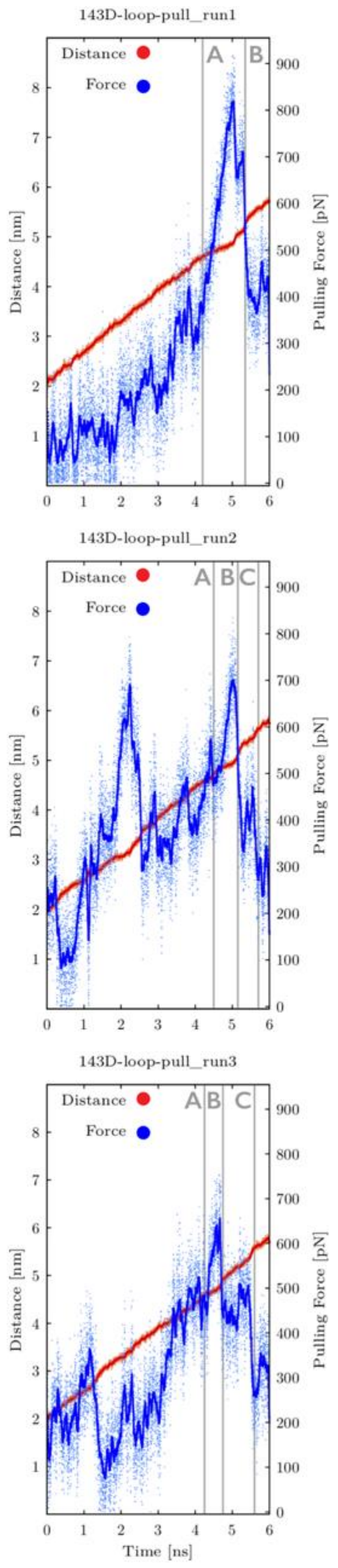

**Figure S7A:** Time evolution of distance between pulling centers and pulling force during three independent *fast pulling* simulations of $143D_{loop\text{-}pull}$ GQ system (see legend of Figure S1A for more details). See Figures S7B and S7C for inspection of structures corresponding to main structural events.

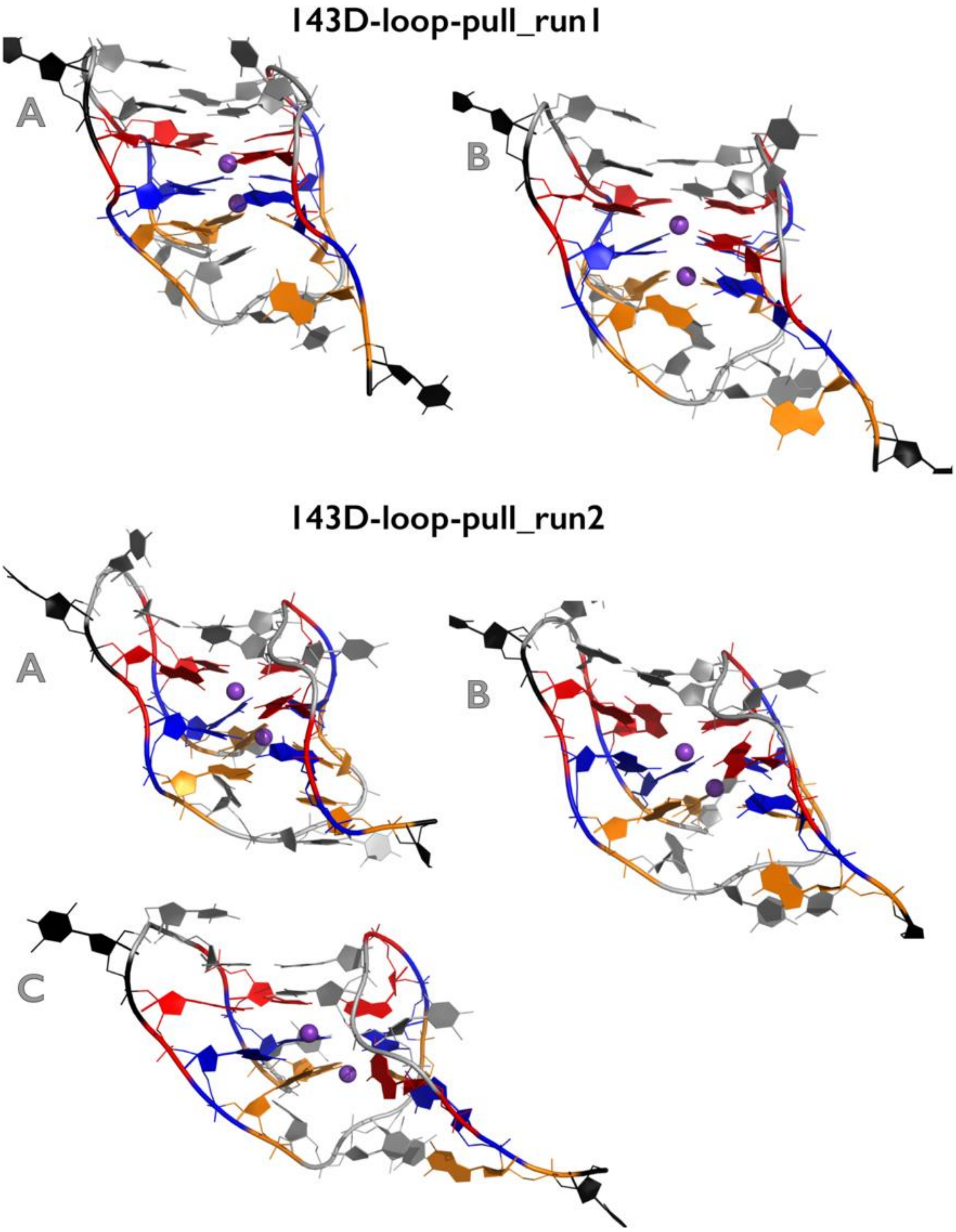

**Figure S7B:** Most important structural events during first and second independent *fast pulling* simulations of $143D_{\text{loop-pull}}$ GQ system. See legend of Figure S1B for more details.

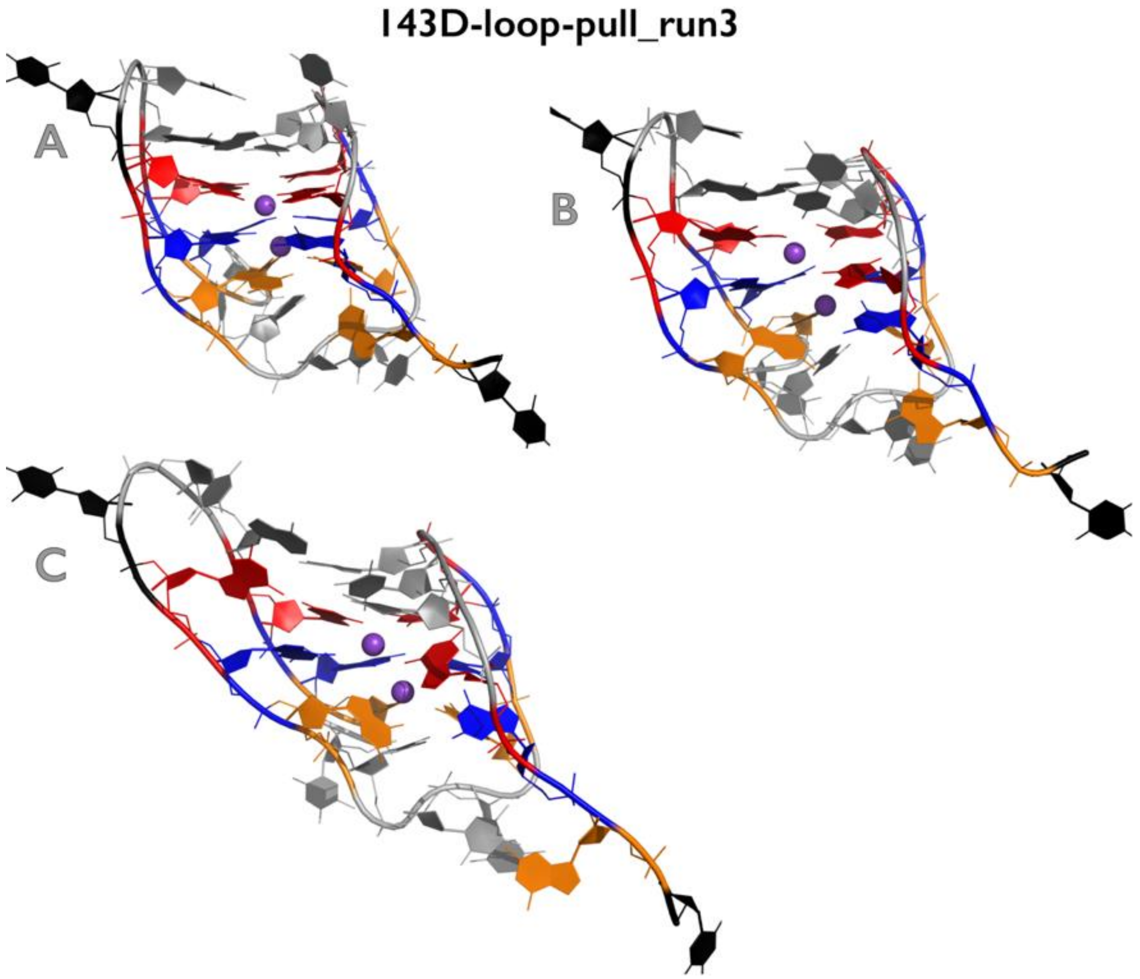

**Figure S7C:** Most important structural events during third independent *fast pulling* simulation of 143D$_{loop\text{-}pull}$ GQ system. See legend of Figure S1B for more details.

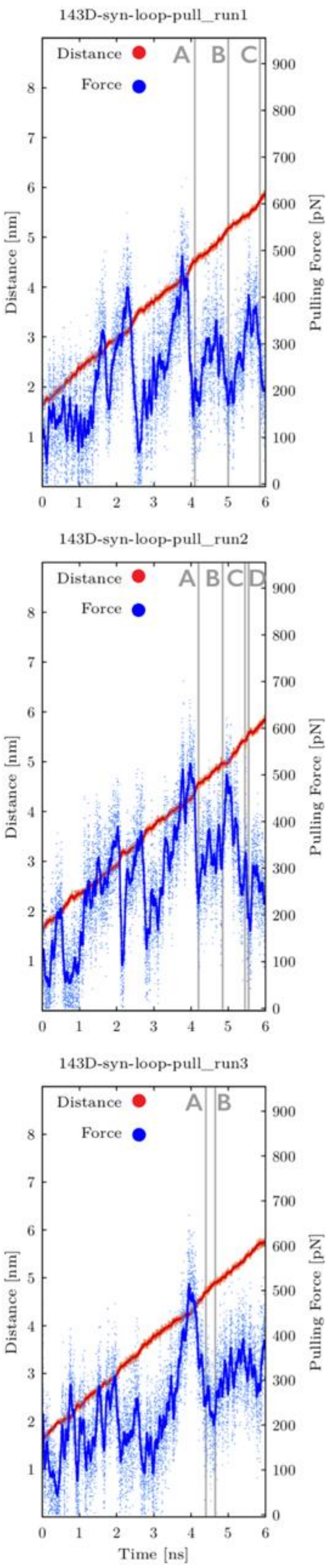

**Figure S8A:** Time evolution of distance between pulling centers and pulling force during three independent *fast pulling* simulations of $143D_{syn\_loop-pull}$ GQ system (see legend of Figure S1A for more details). See Figures S8B and S8C for inspection of structures corresponding to main structural events.

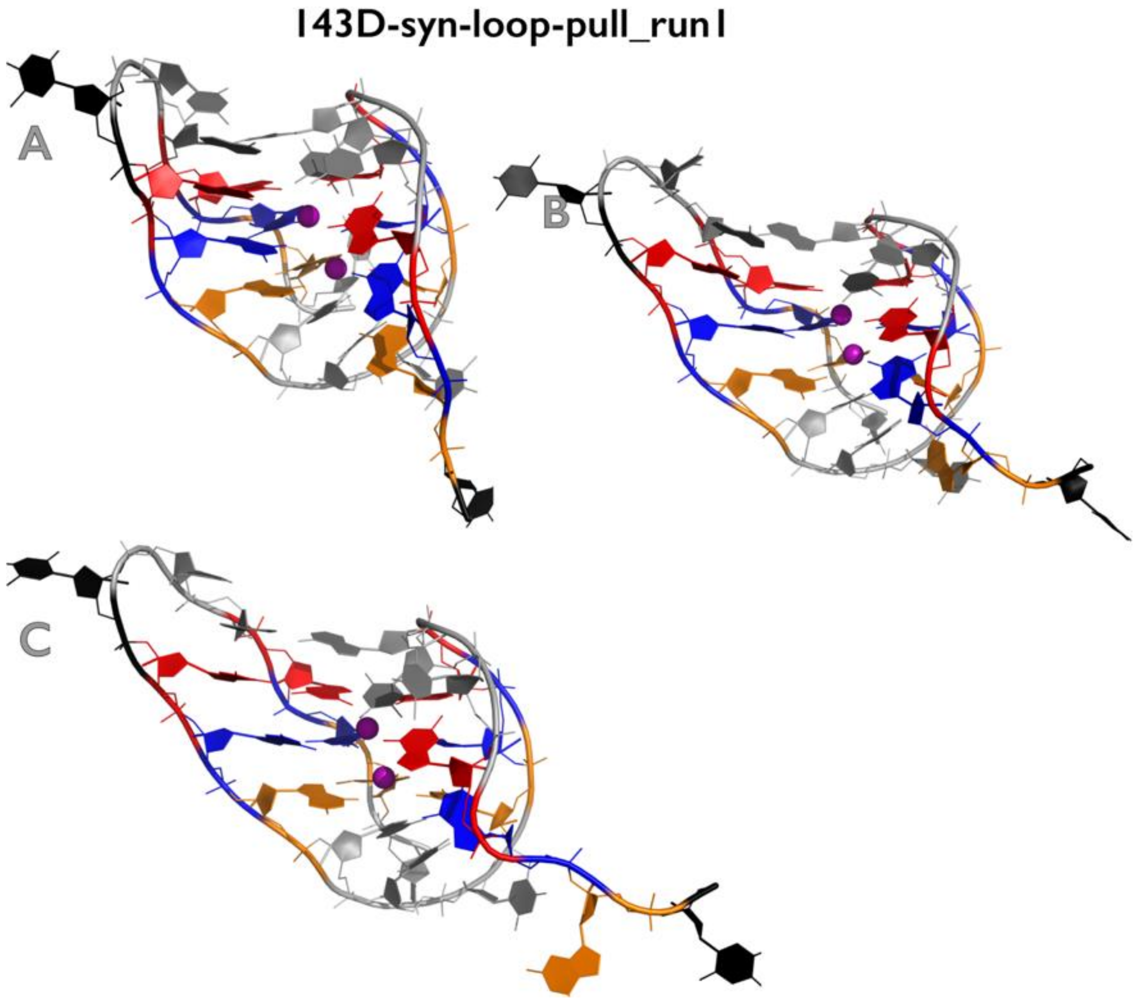

**Figure S8B:** Most important structural events during first independent *fast pulling* simulation of 143D$_{syn_loop\text{-}pull}$ GQ system. See legend of Figure S1B for more details.

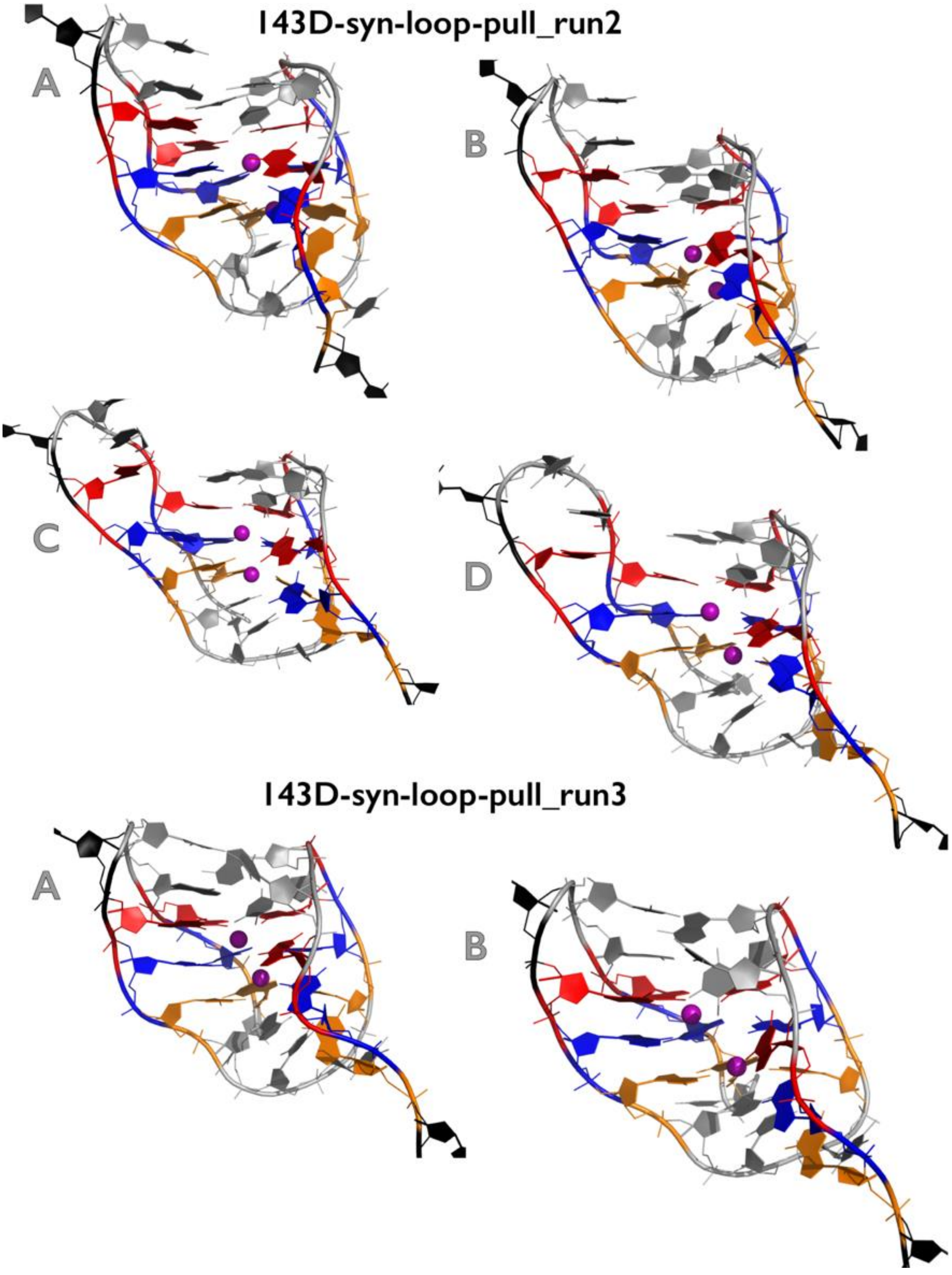

**Figure S8C:** Most important structural events during second and third independent *fast pulling* simulations of 143D$_{syn_loop-pull}$ GQ system. See legend of Figure S1B for more details.

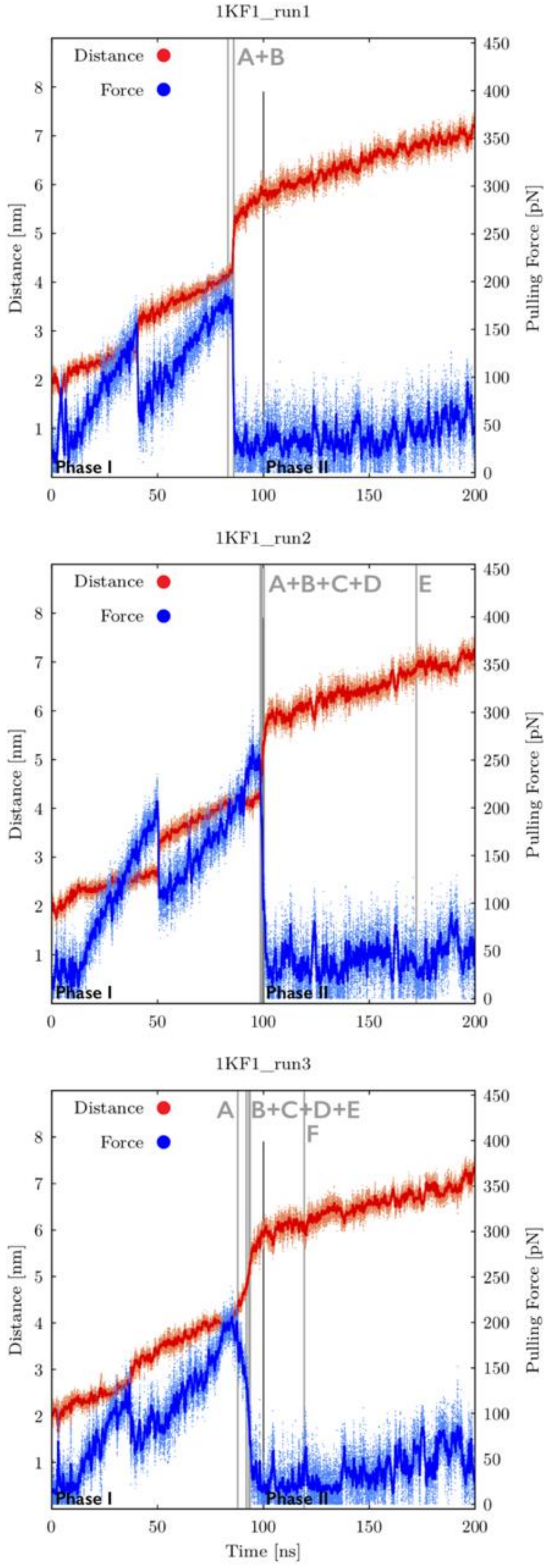

**Figure S9A:** Time evolution of distance between pulling centers and pulling force during three independent *slow zig-zag pulling* simulations of 1KF1 GQ system. Snapshots were saved every 5 ps and plots are showing both instantaneous values (orange and light-blue dots for distance and force, respectively) and smoothing, i.e., averaging over 100 consecutive snapshots (red and blue lines for distance and force, respectively). Pulling phases are marked (see Methods in the main text for details) and main structural events are highlighted as grey vertical lines with labels (capital letters). See Figures S9B-S9D for inspection of structures corresponding to main structural events. Note that first major drops of the pulling force before the GQ unfolding event “A” (and notable prolongation of end-to-end distances) are connected with repositioning of terminal T residues.

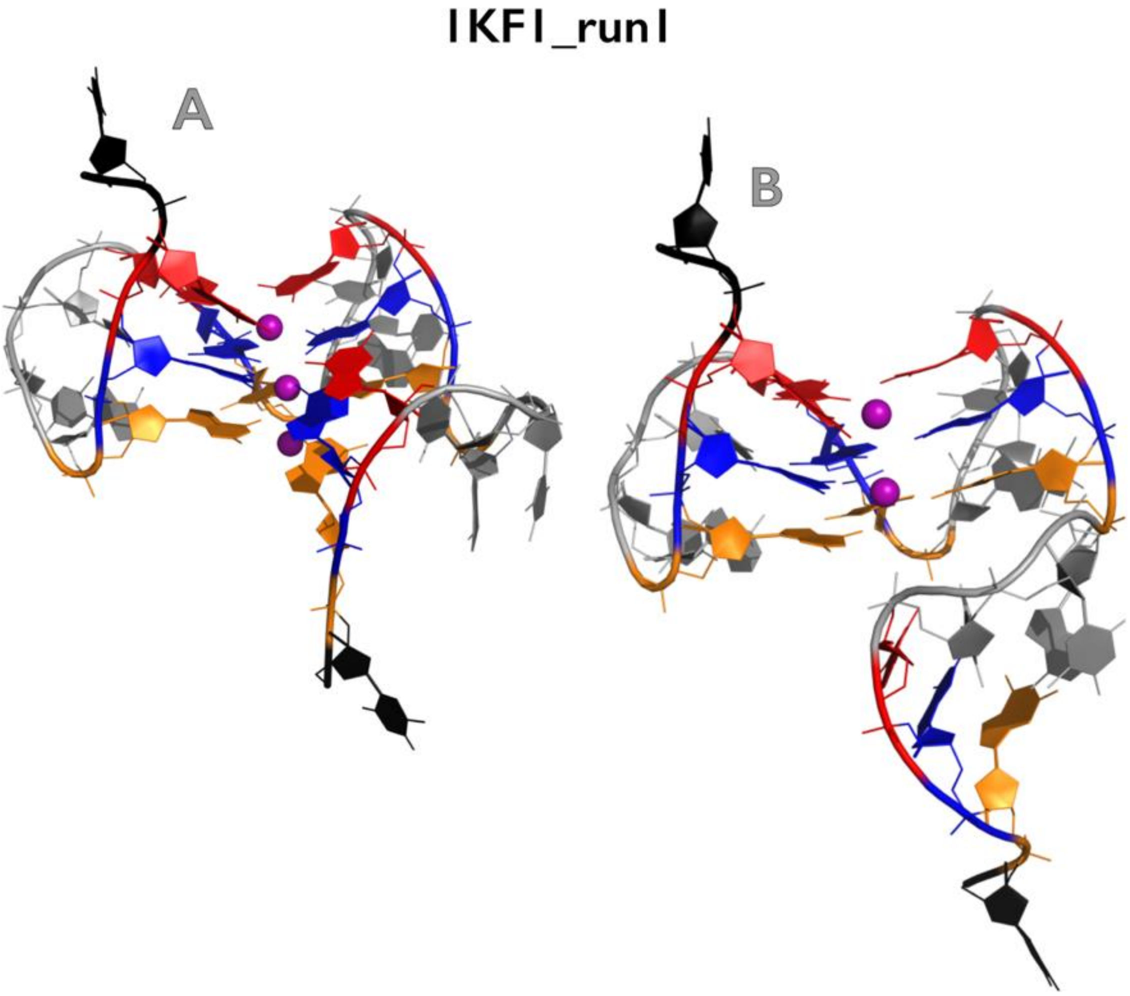

**Figure S9B:** Most important structural events during first independent *slow zig-zag pulling* simulation of 1KF1 GQ system. See legend of Figure S1B for more details.

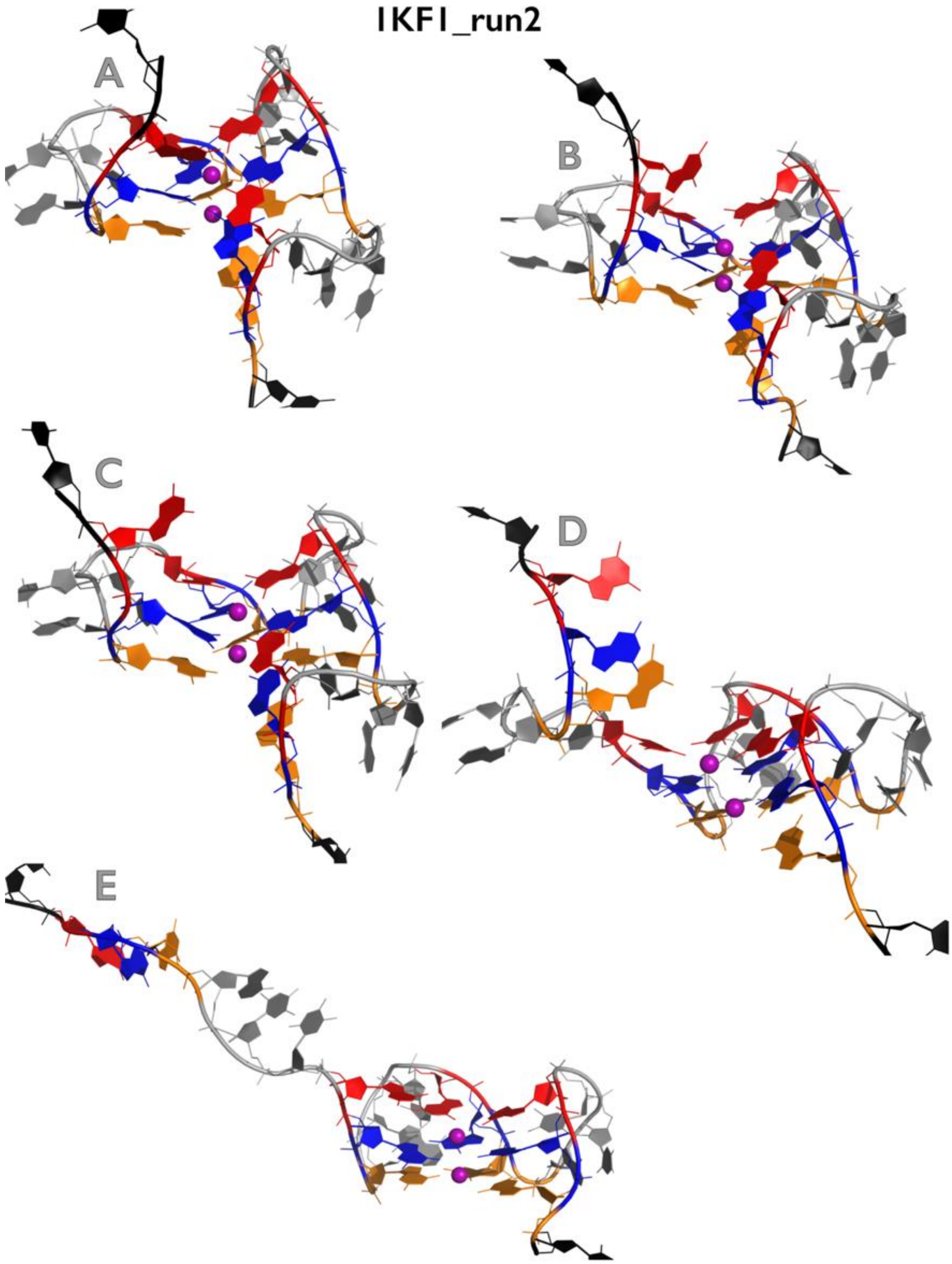

**Figure S9C:** Most important structural events during second independent *slow zig-zag pulling* simulation of 1KF1 GQ system. See legend of Figure S1B for more details.

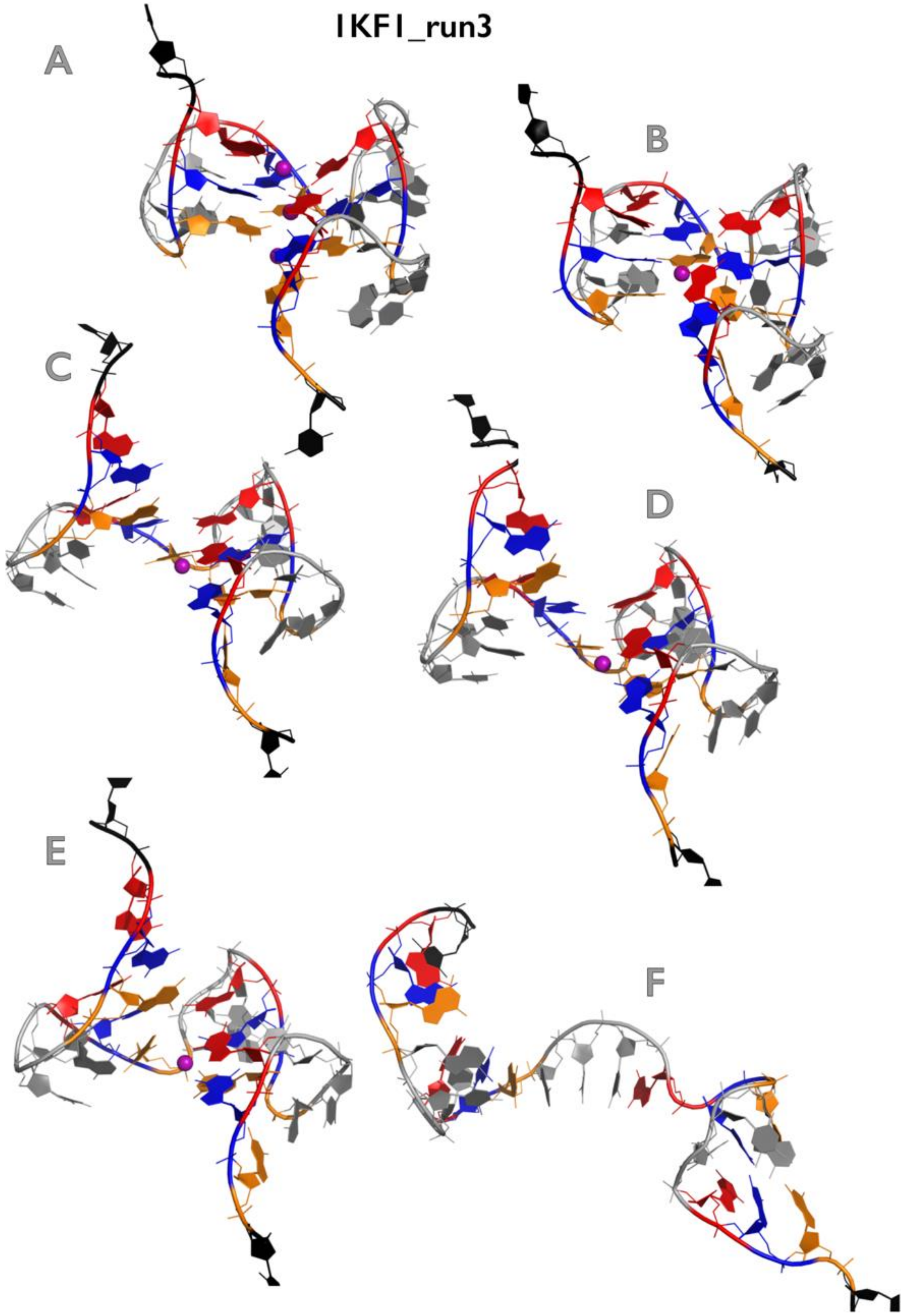

**Figure S9D:** Most important structural events during third independent *slow zig-zag pulling* simulation of 1KF1 GQ system. See legend of Figure S1B for more details.

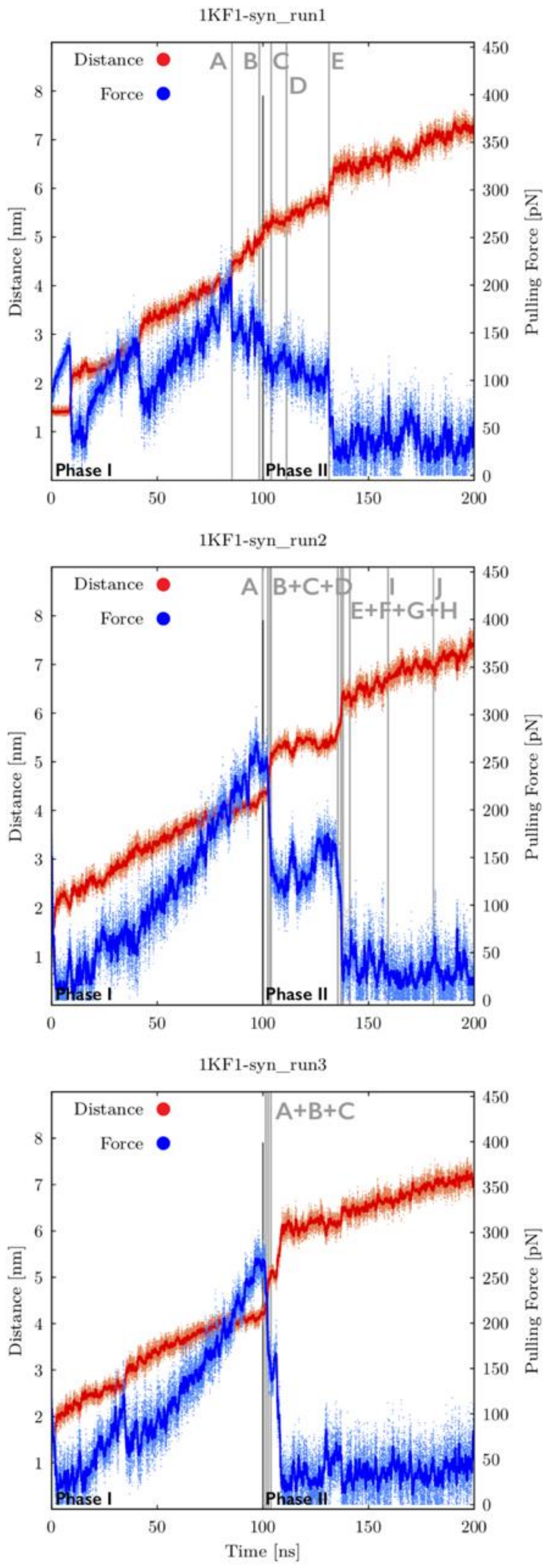

**Figure S10A:** Time evolution of distance between pulling centers and pulling force during three independent *slow zig-zag pulling* simulations of $1KF1_{syn}$ GQ system (see legend of Figure S9A for more details). See Figures S10B-S10D for inspection of structures corresponding to main structural events.

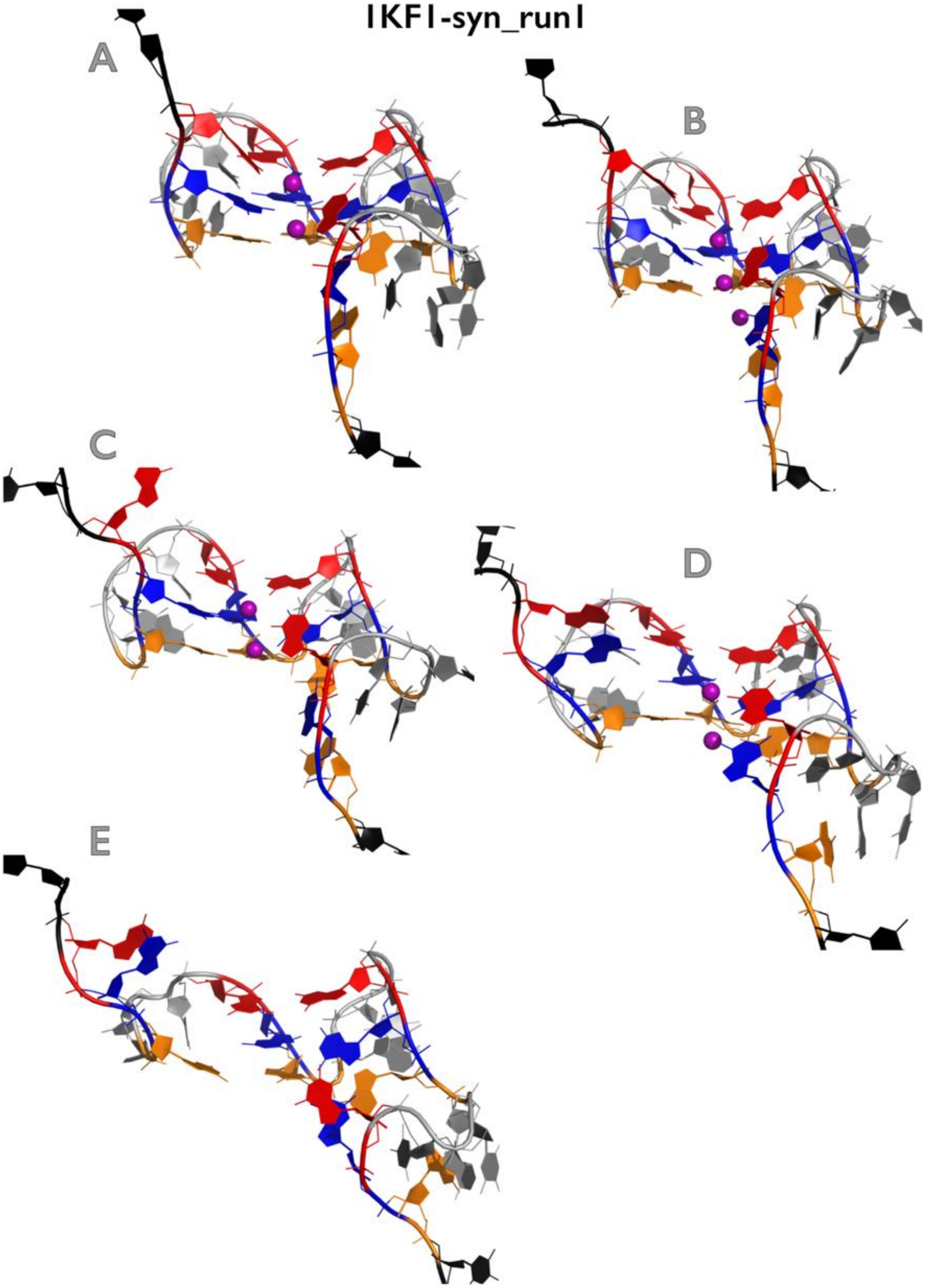

**Figure S10B:** Most important structural events during first independent *slow zig-zag pulling* simulation of $1KF1_{syn}$ GQ system. See legend of Figure S1B for more details.

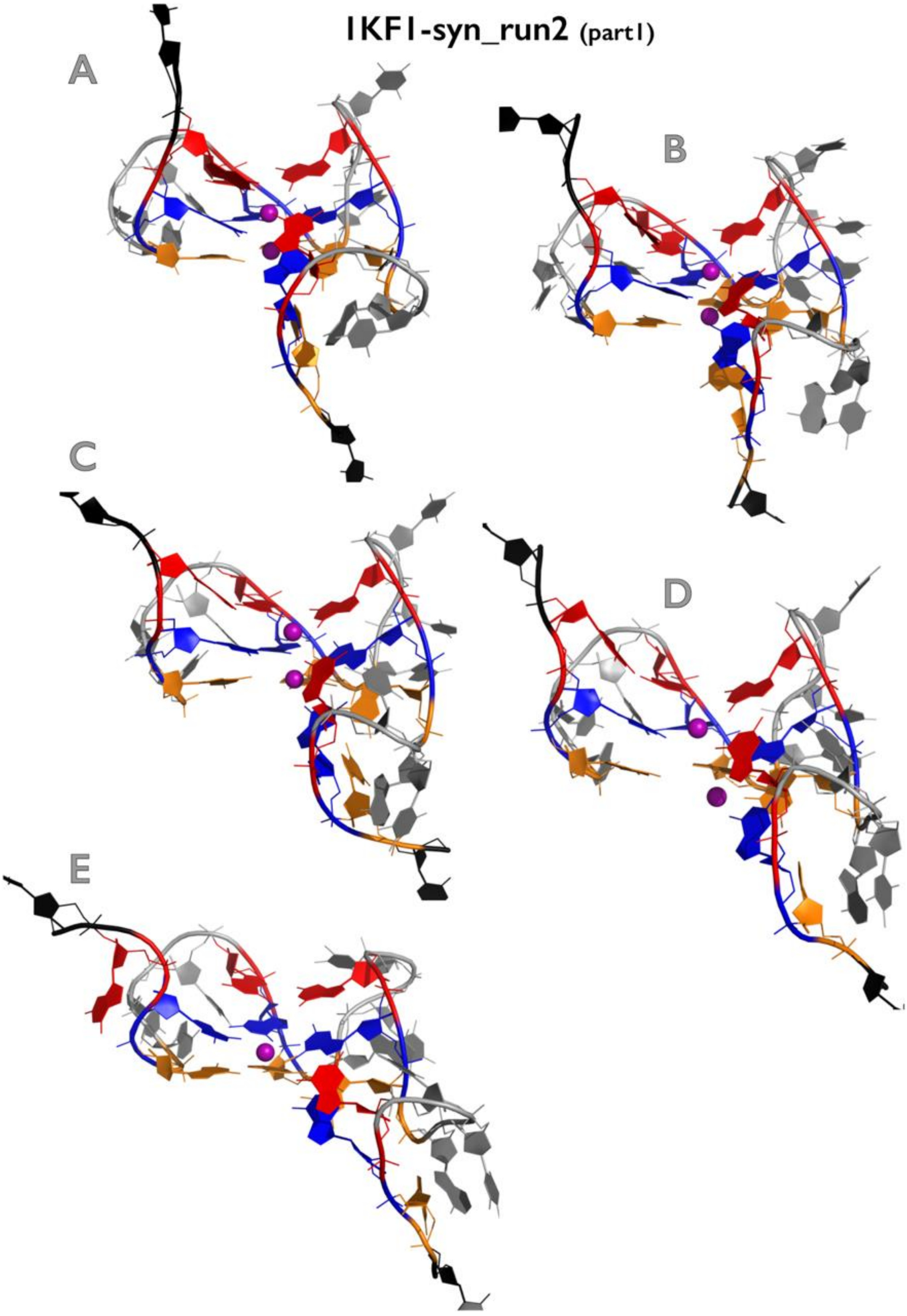

Figure continuing on the next page

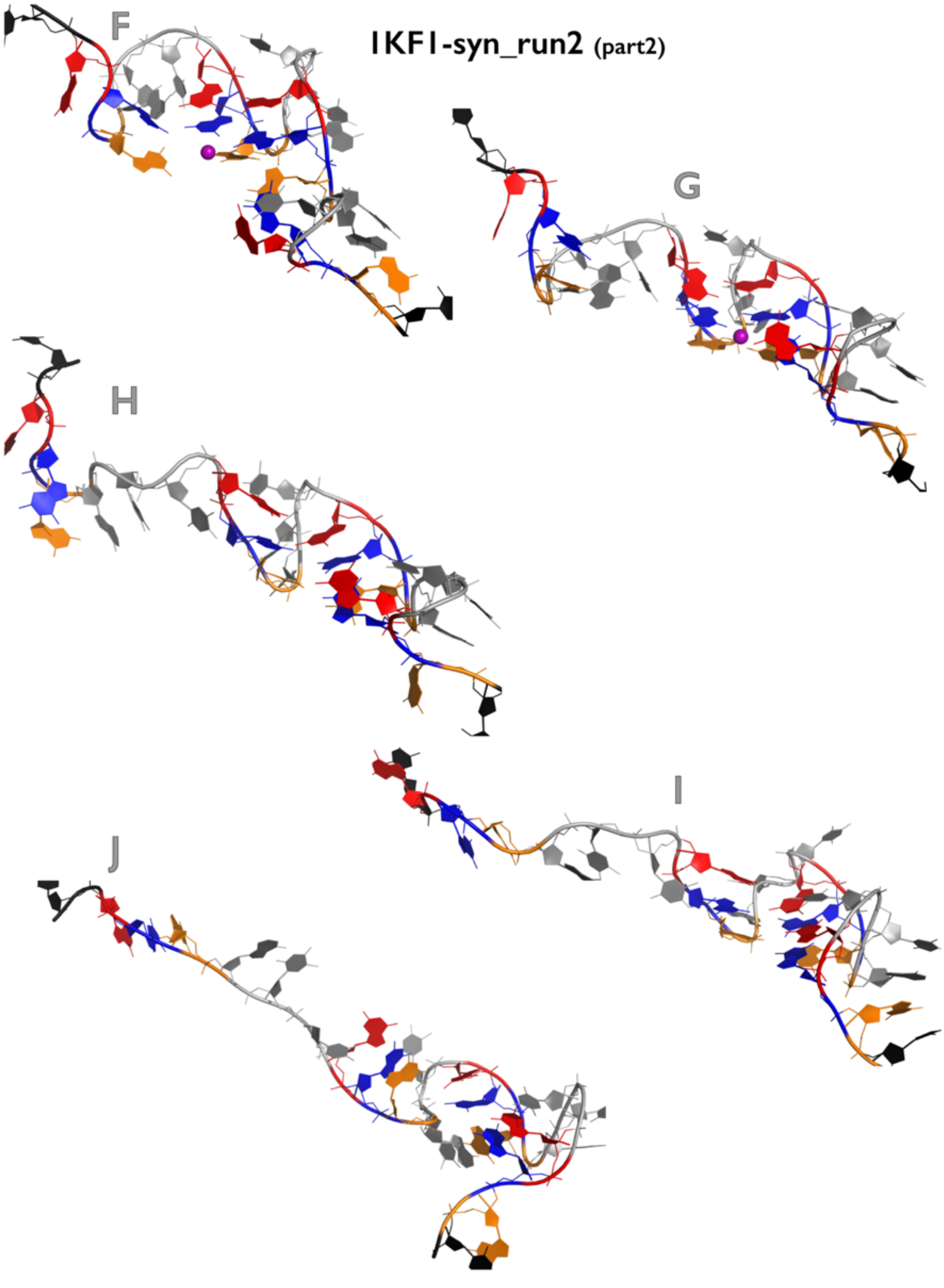

**Figure S10C:** Most important structural events during second independent *slow zig-zag pulling* simulation of 1KF1$_{syn}$ GQ system. See legend of Figure S1B for more details.

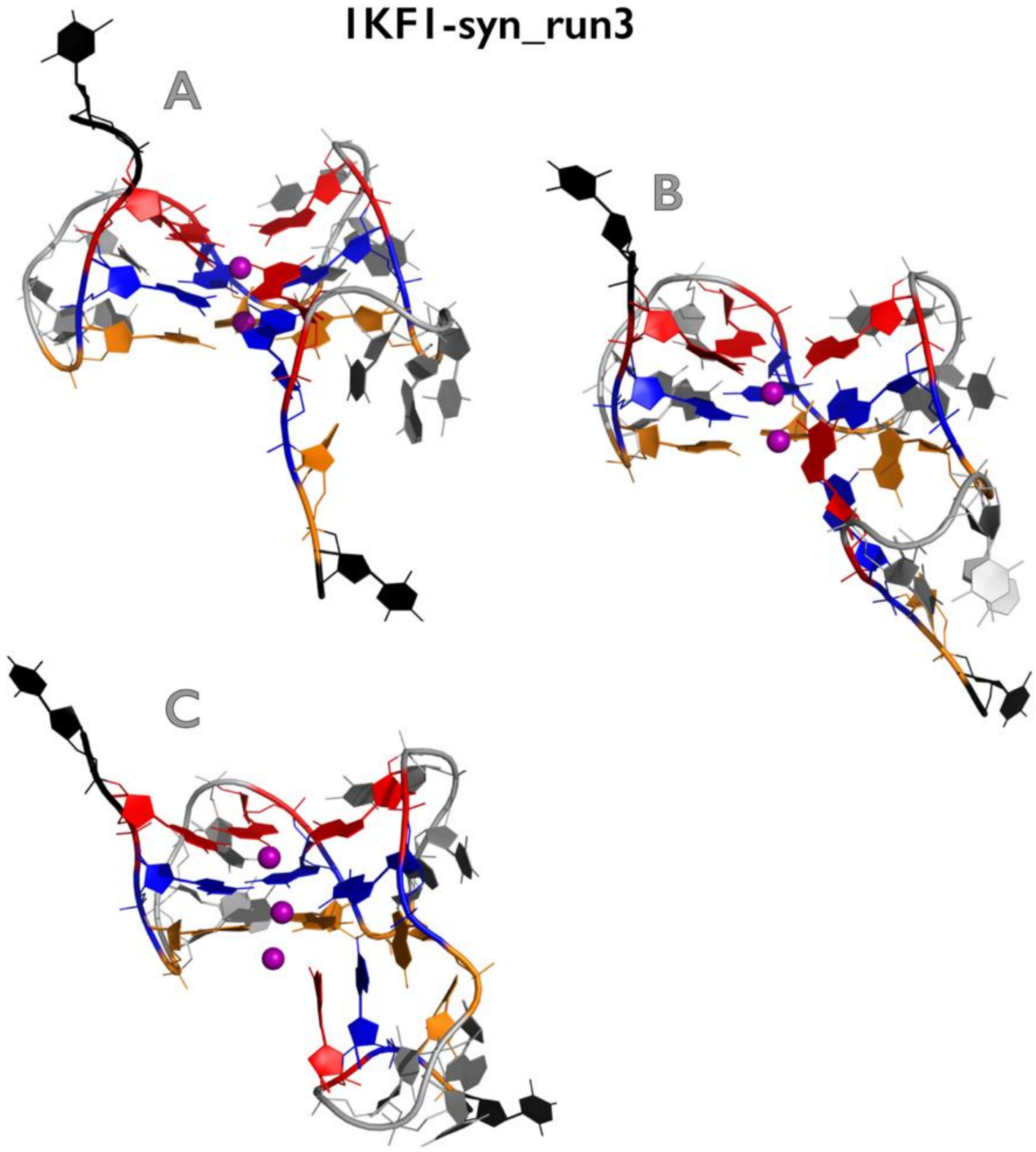

**Figure S10D:** Most important structural events during third independent *slow zig-zag pulling* simulation of $1KF1_{syn}$ GQ system. See legend of Figure S1B for more details.

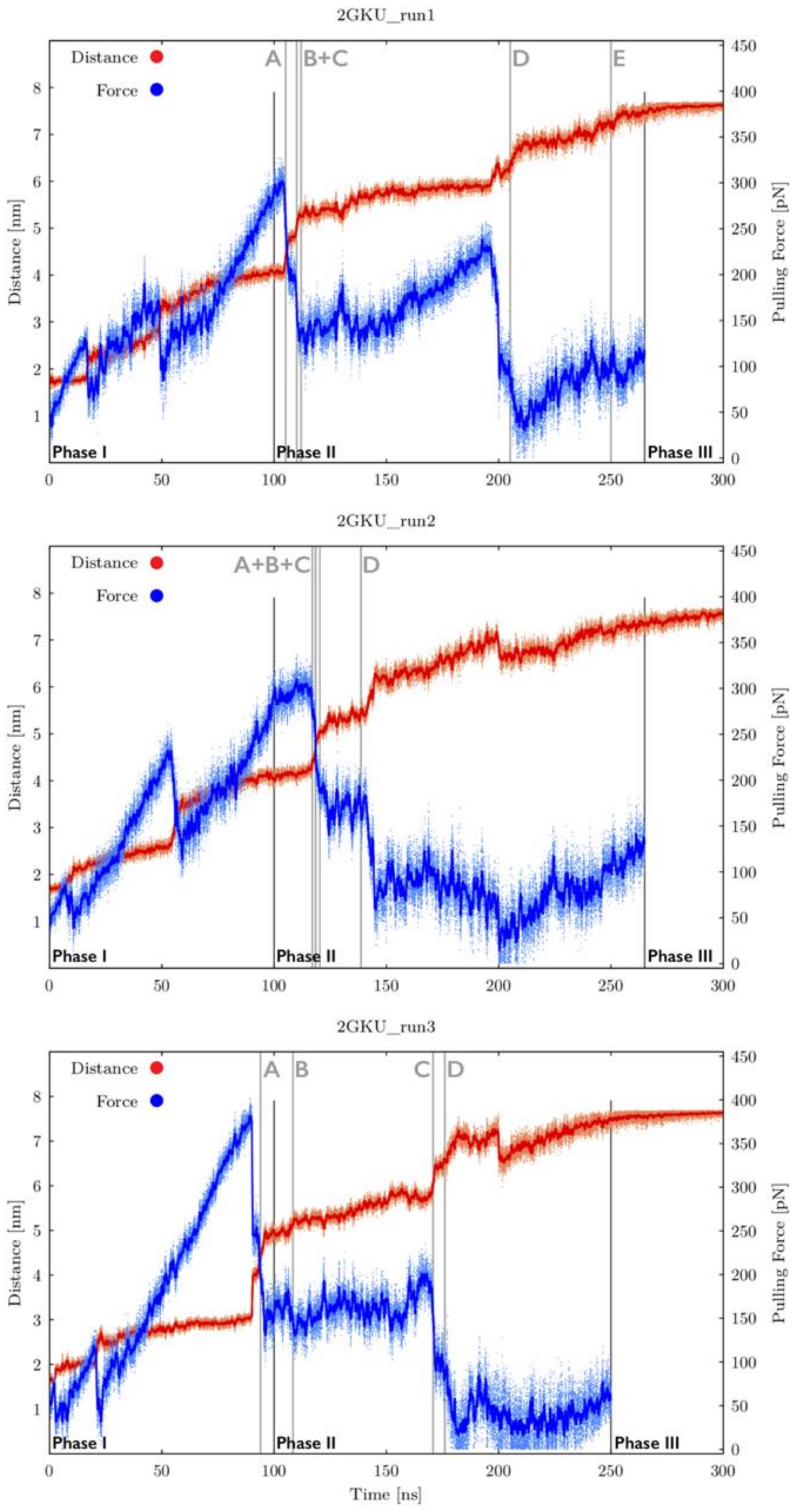

**Figure S11A:** Time evolution of distance between pulling centers and pulling force during three independent *slow zig-zag pulling* simulations of 2GKU GQ system (see legend of Figure S9A for more details). See Figures S11B-S11D for inspection of structures corresponding to main structural events.

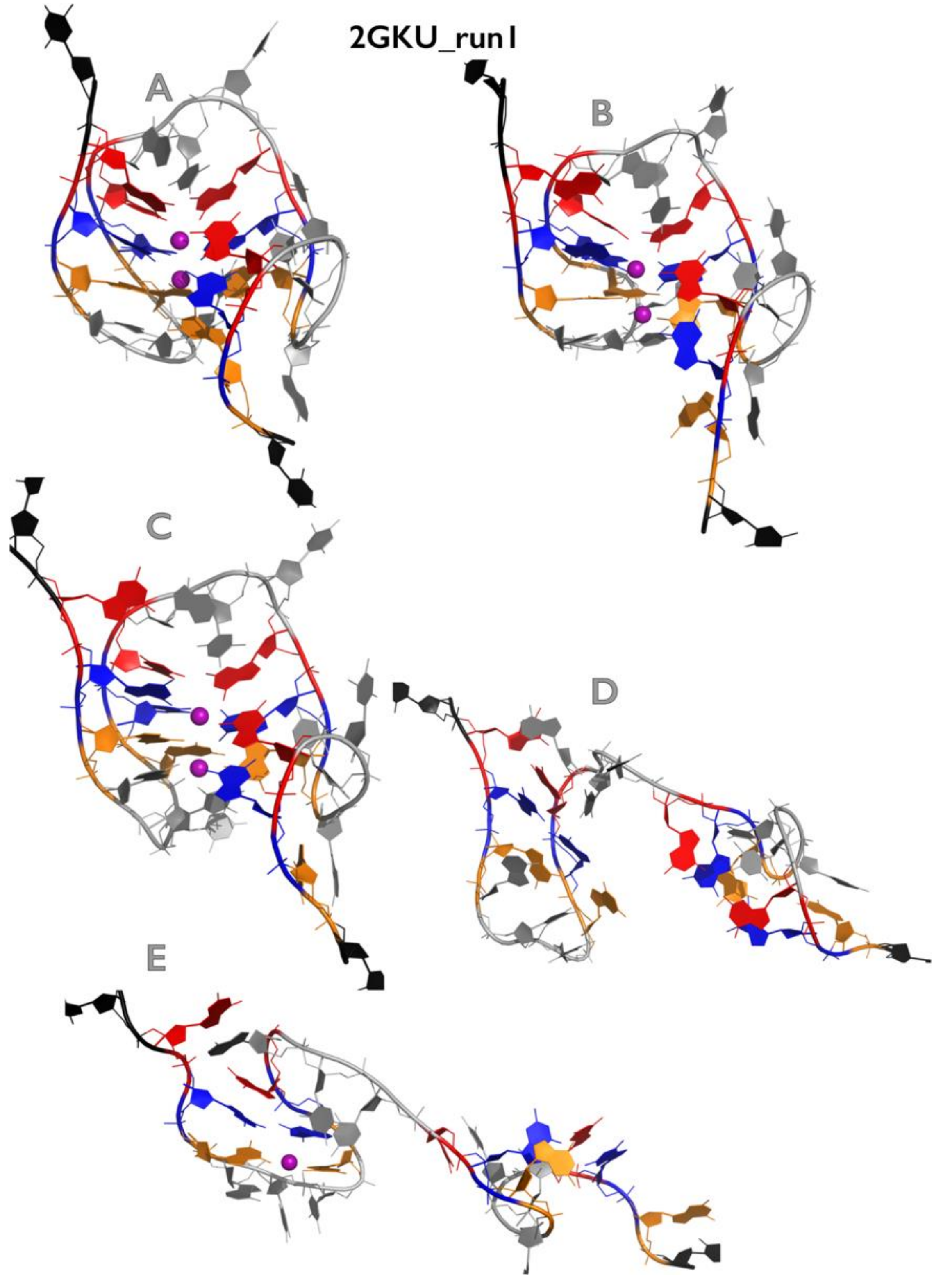

**Figure S11B:** Most important structural events during first independent *slow zig-zag pulling* simulation of 2GKU GQ system. See legend of Figure S1B for more details.

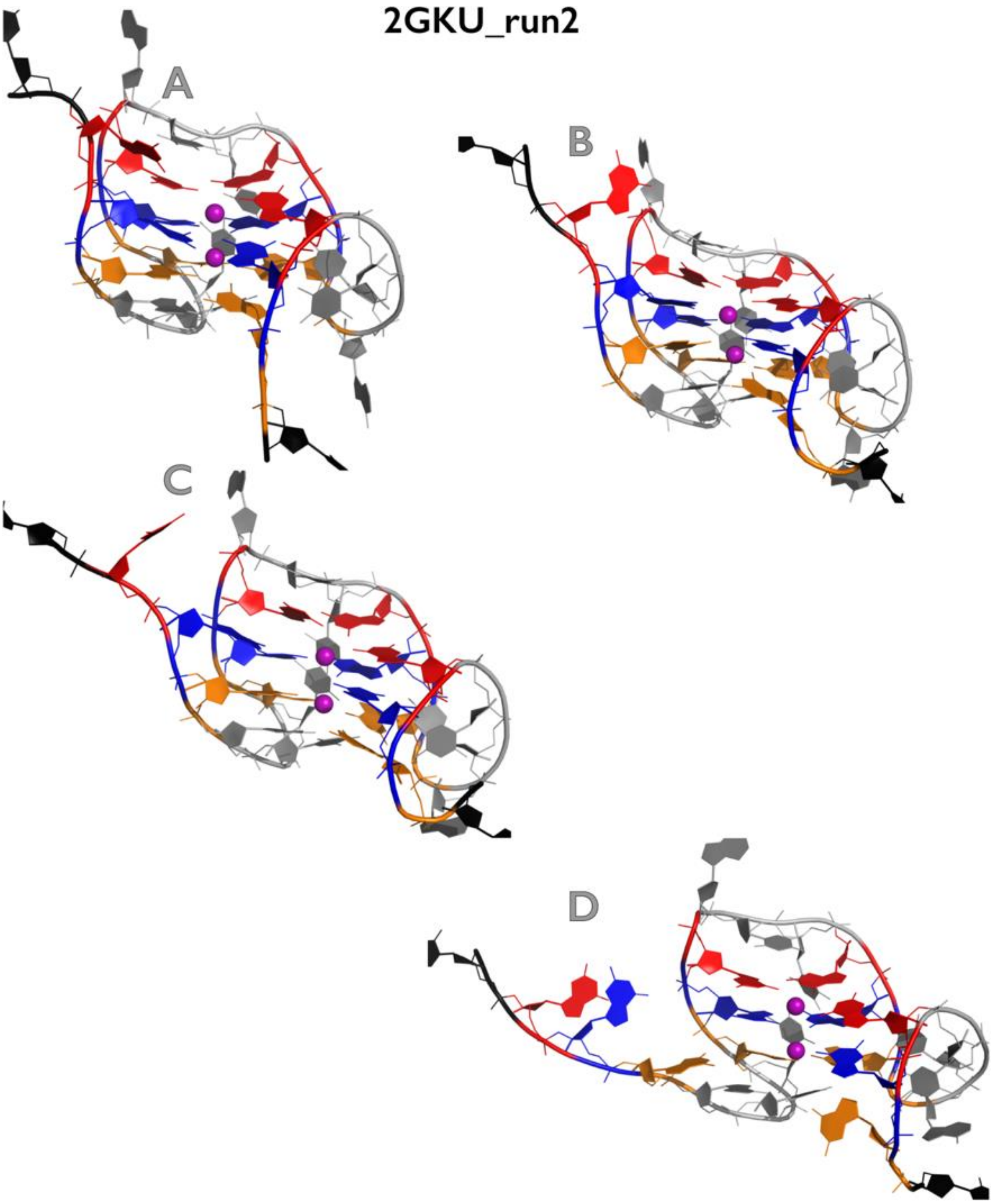

**Figure S11C:** Most important structural events during second independent *slow zig-zag pulling* simulation of 2GKU GQ system. See legend of Figure S1B for more details.

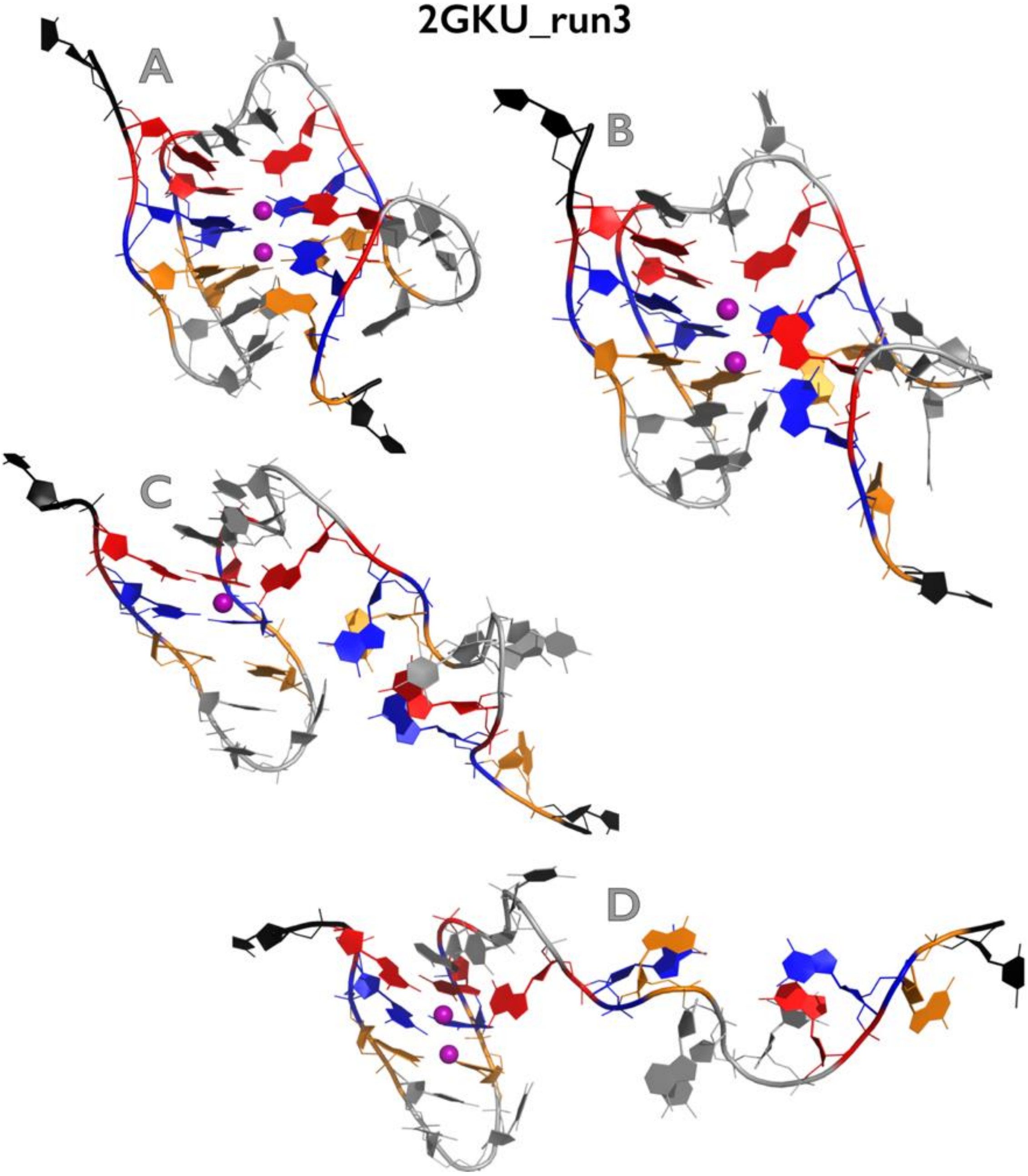

**Figure S11D:** Most important structural events during third independent *slow zig-zag pulling* simulation of 2GKU GQ system. See legend of Figure S1B for more details.

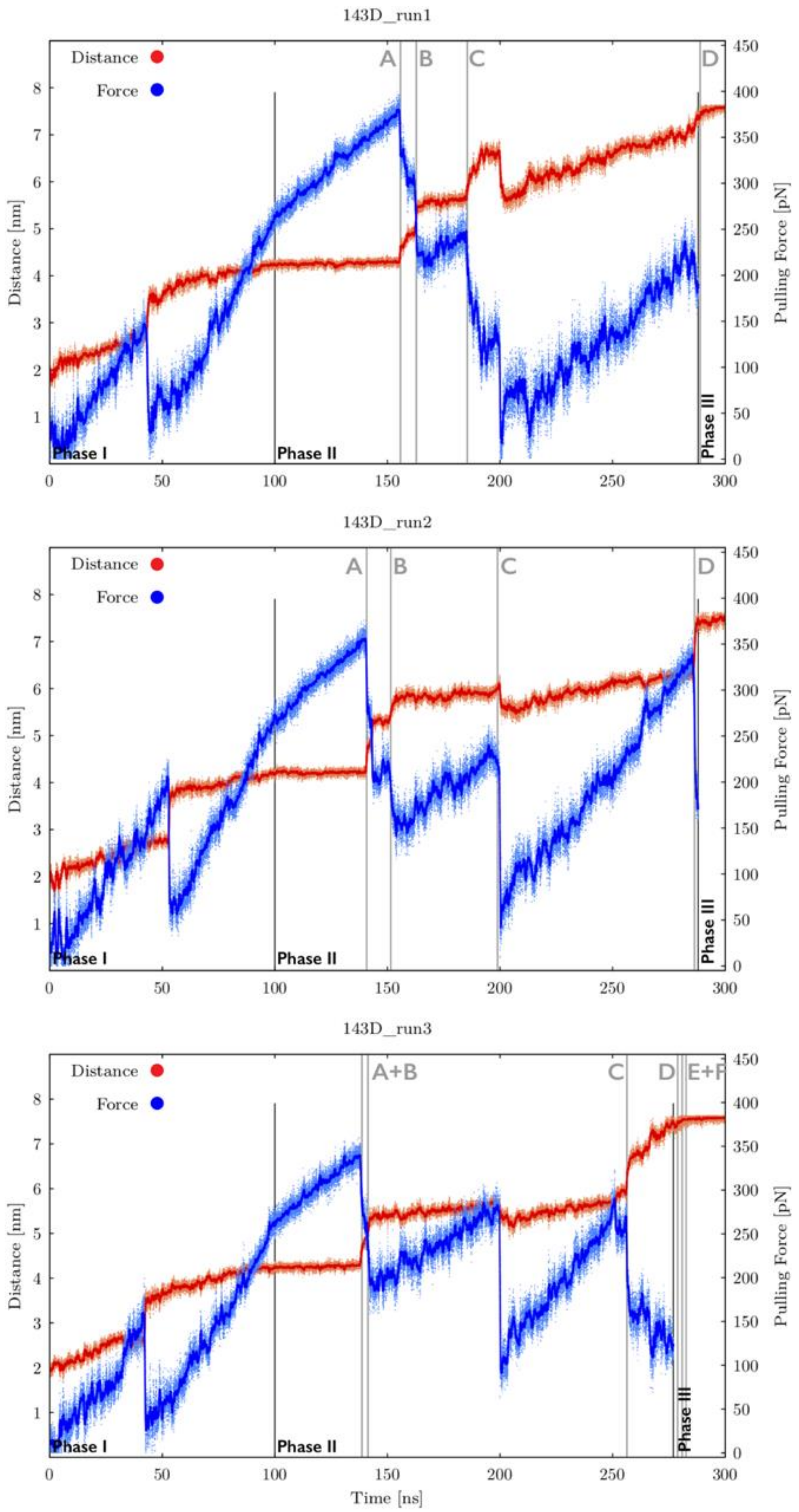

**Figure S12A:** Time evolution of distance between pulling centers and pulling force during three independent *slow zig-zag pulling* simulations of 143D GQ system (see legend of Figure S9A for more details). See Figures S12B-S12D for inspection of structures corresponding to main structural events.

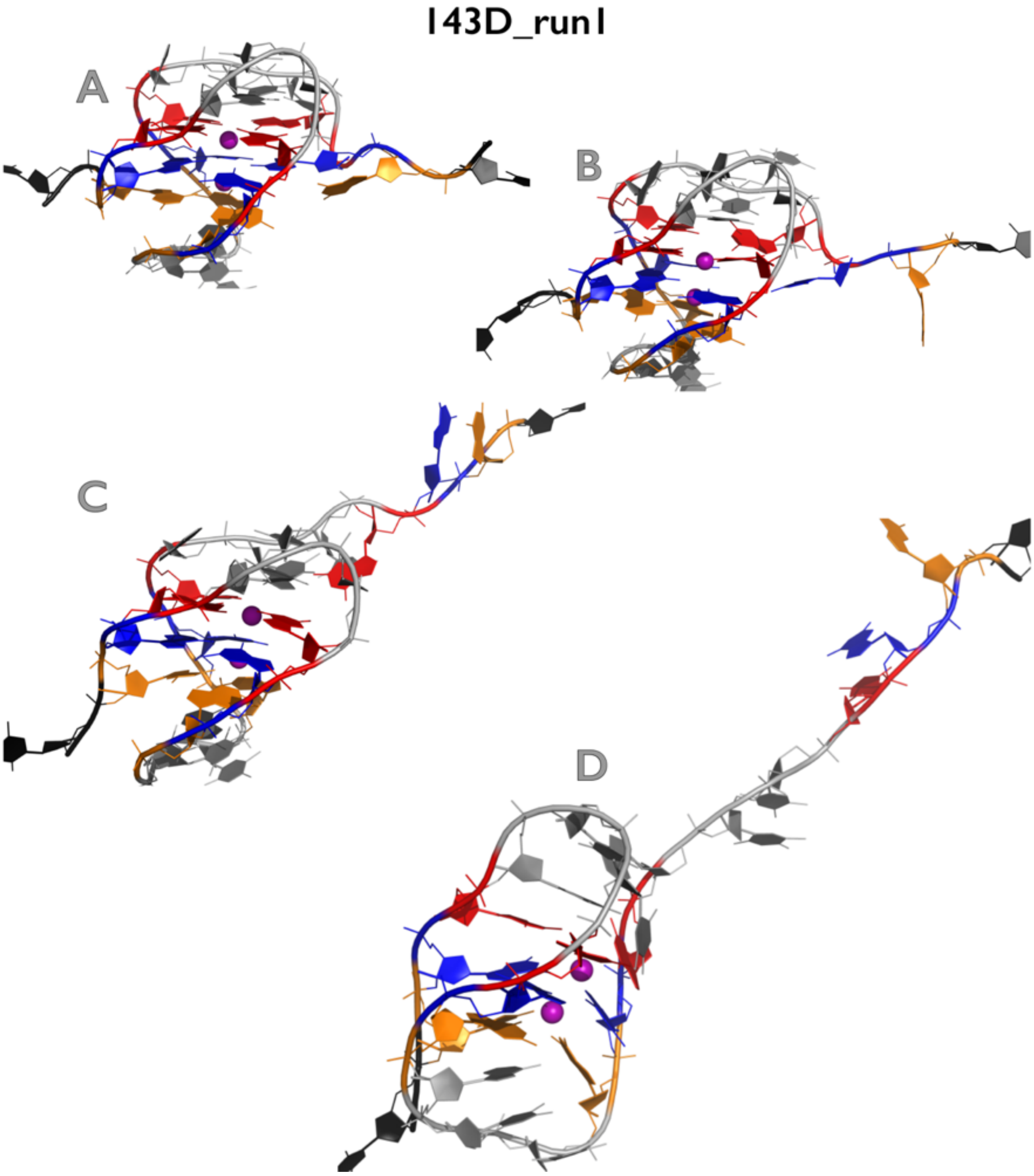

**Figure S12B:** Most important structural events during first independent *slow zig-zag pulling* simulation of 143D GQ system. See legend of Figure S1B for more details.

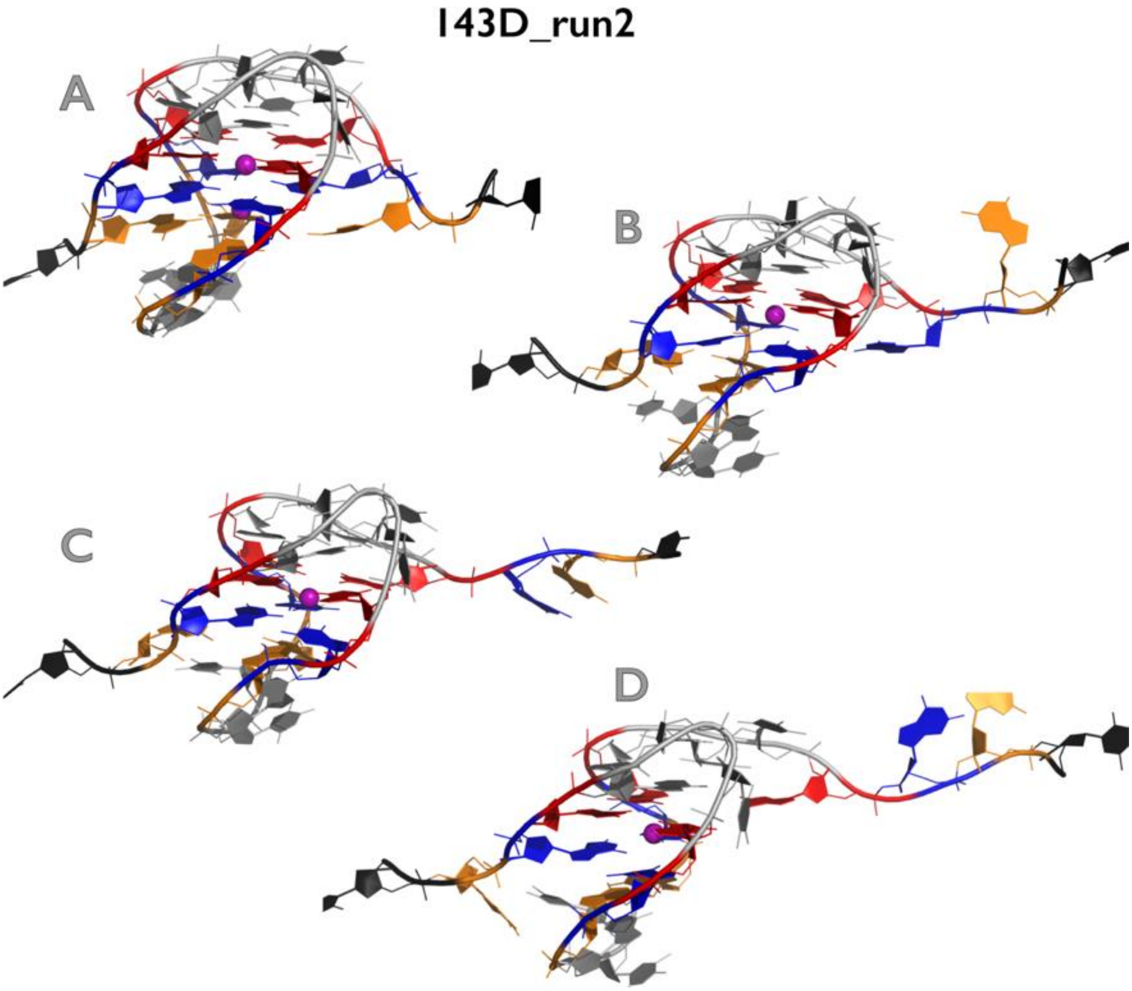

**Figure S12C:** Most important structural events during second independent *slow zig-zag pulling* simulation of 143D GQ system. See legend of Figure S1B for more details.

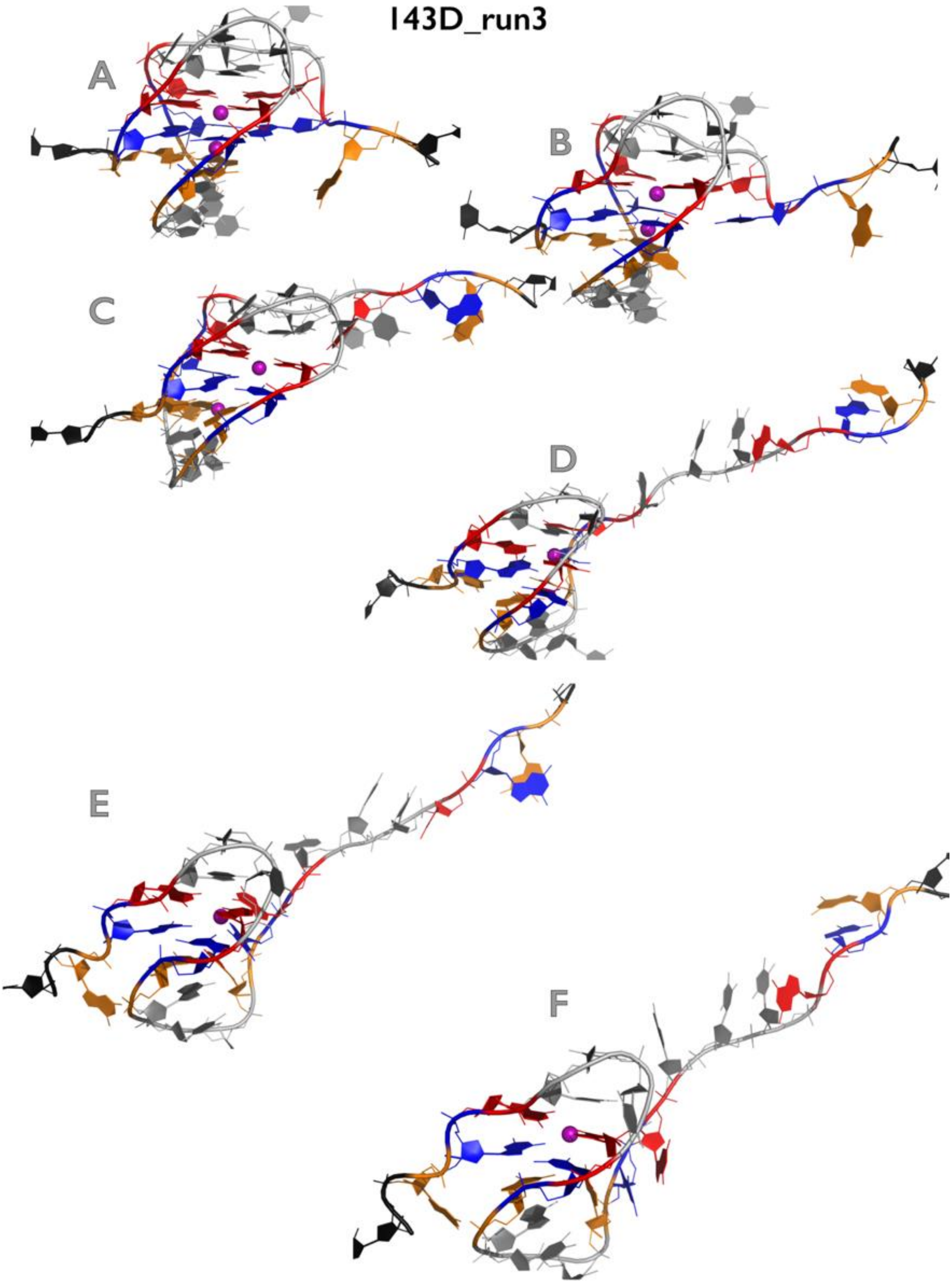

**Figure S12D:** Most important structural events during third independent *slow zig-zag pulling* simulation of 143D GQ system. See legend of Figure S1B for more details.

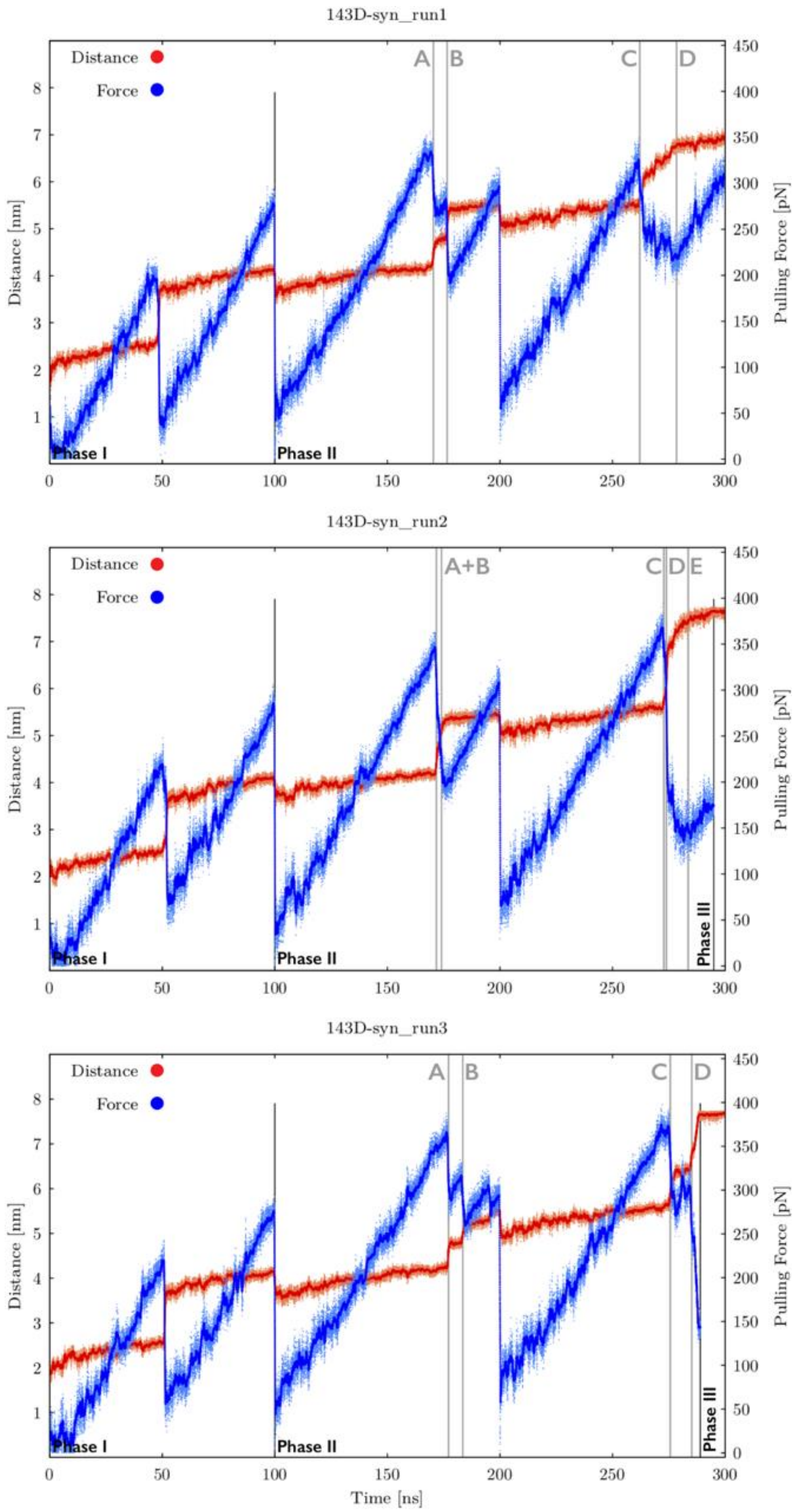

**Figure S13A:** Time evolution of distance between pulling centers and pulling force during three independent *slow zig-zag pulling* simulations of 143D$_{syn}$ GQ system (see legend of Figure S9A for more details). See Figures S13B-S13D for inspection of structures corresponding to main structural events.

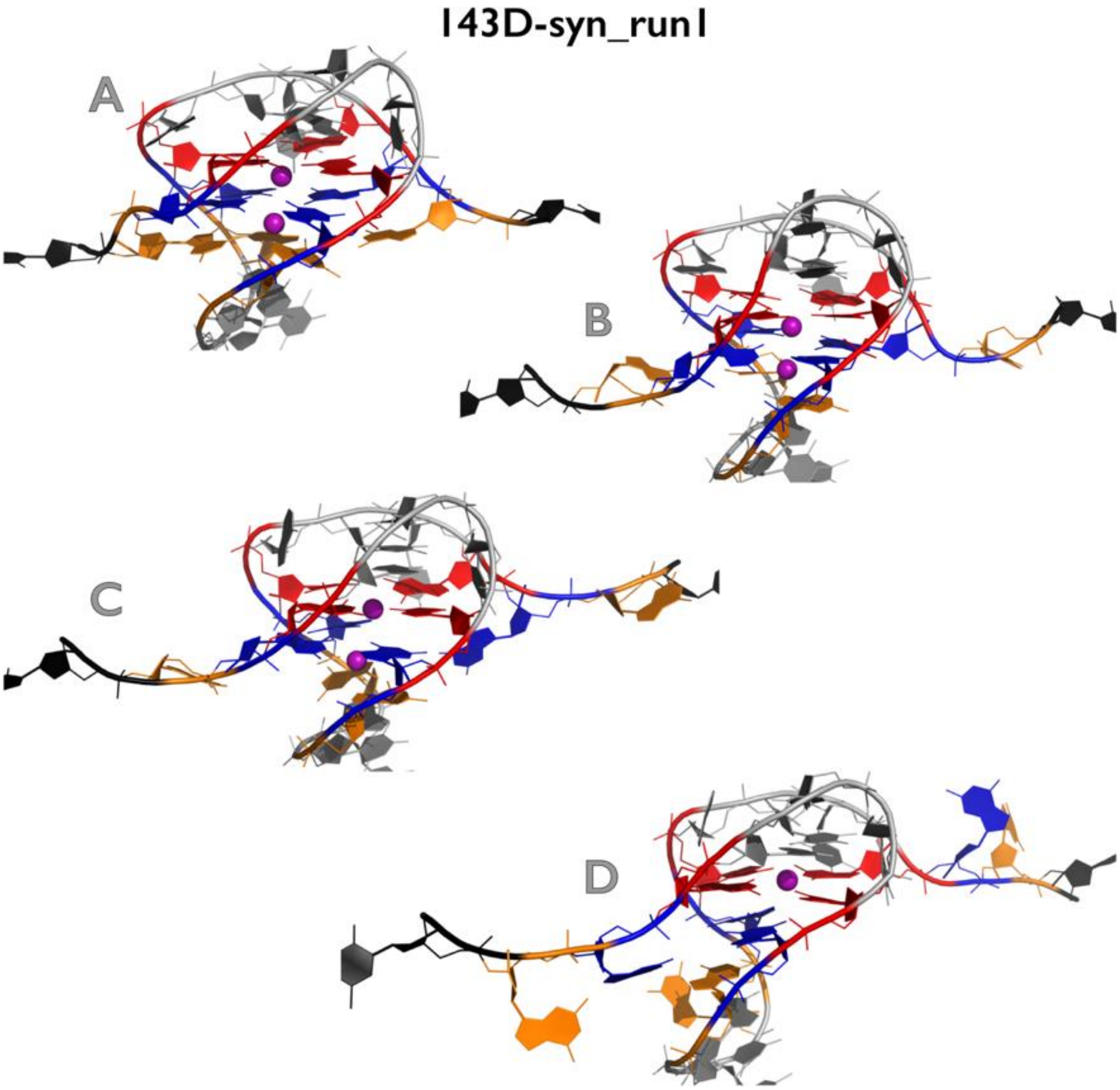

**Figure S13B:** Most important structural events during first independent *slow zig-zag pulling* simulation of $143D_{syn}$ GQ system. See legend of Figure S1B for more details.

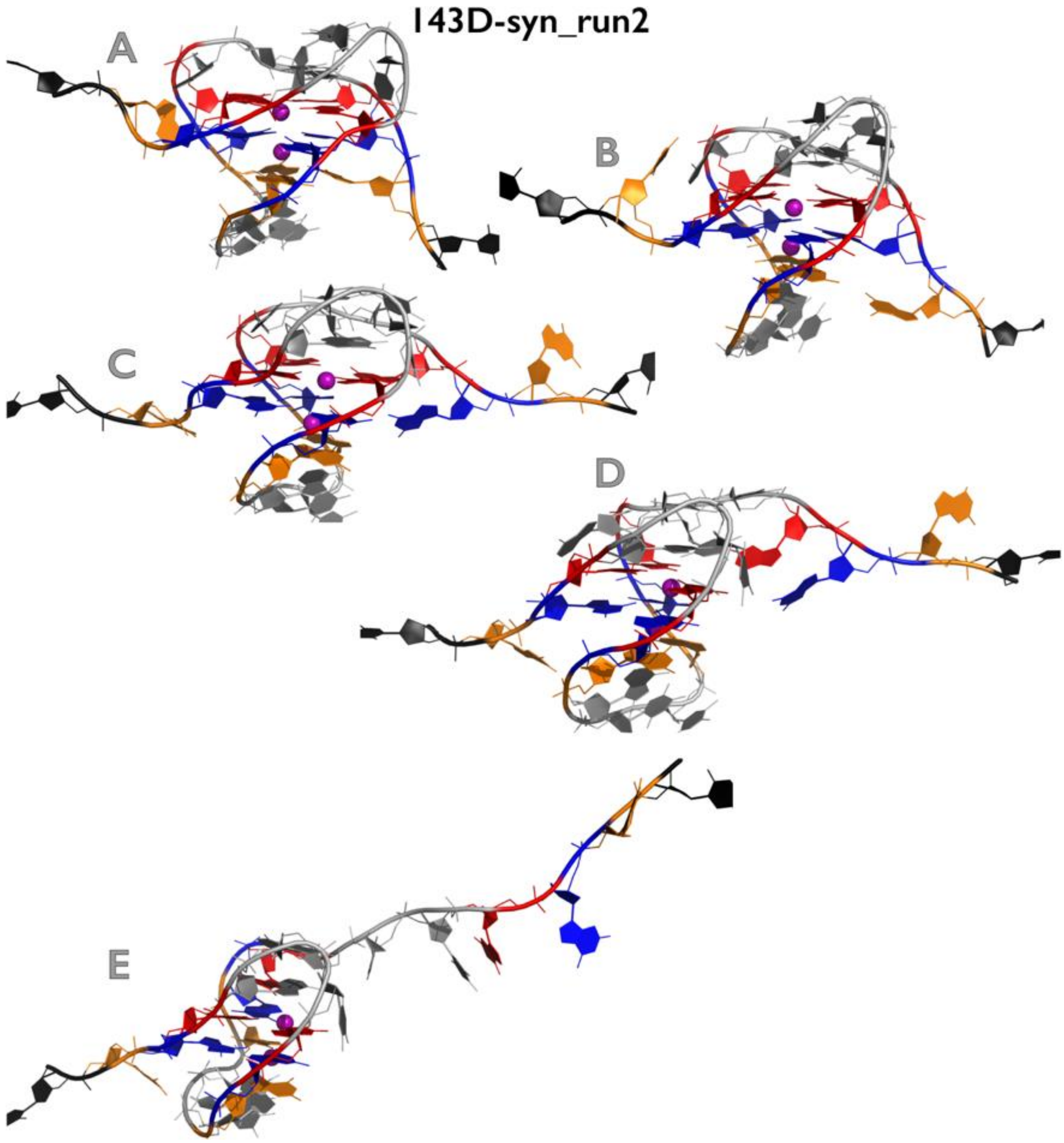

**Figure S13C:** Most important structural events during second independent *slow zig-zag pulling* simulation of $143D_{syn}$ GQ system. See legend of Figure S1B for more details.

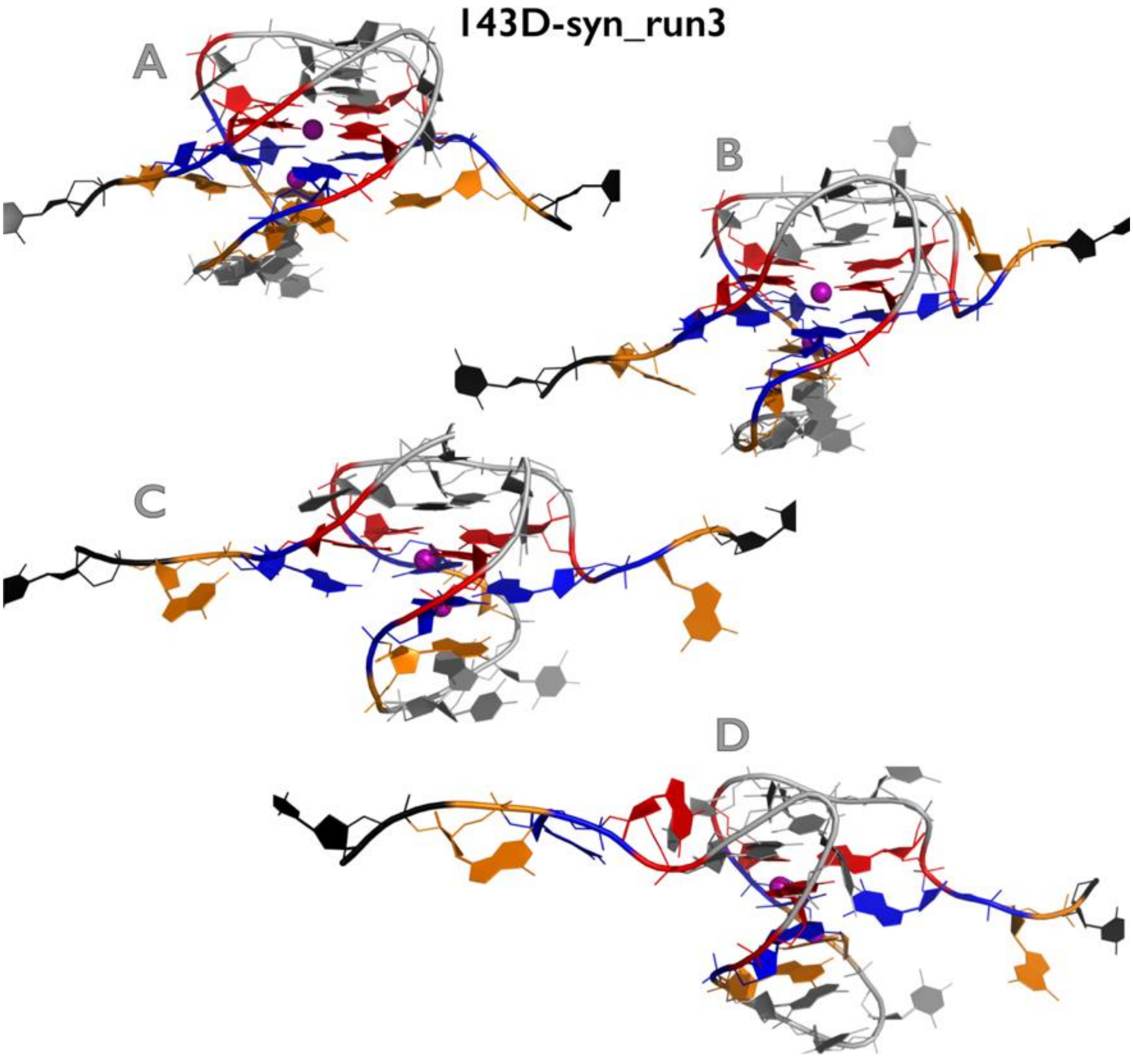

**Figure S13D:** Most important structural events during third independent *slow zig-zag pulling* simulation of 143D$_{syn}$ GQ system. See legend of Figure S1B for more details.

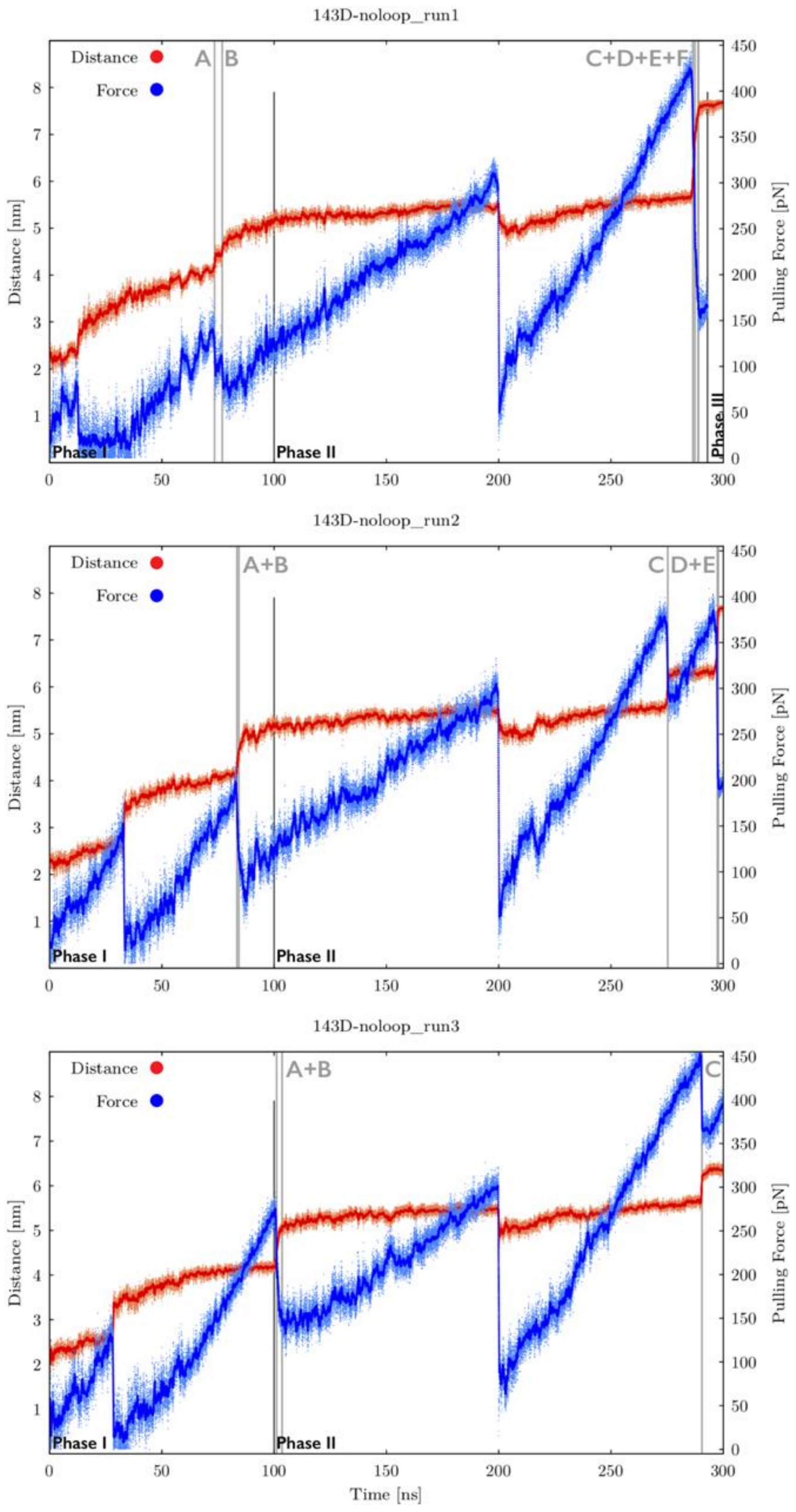

**Figure S14A:** Time evolution of distance between pulling centers and pulling force during three independent *slow zig-zag pulling* simulations of $143D_{noloop}$ GQ system (see legend of Figure S9A for more details). See Figures S14B-S14D for inspection of structures corresponding to main structural events.

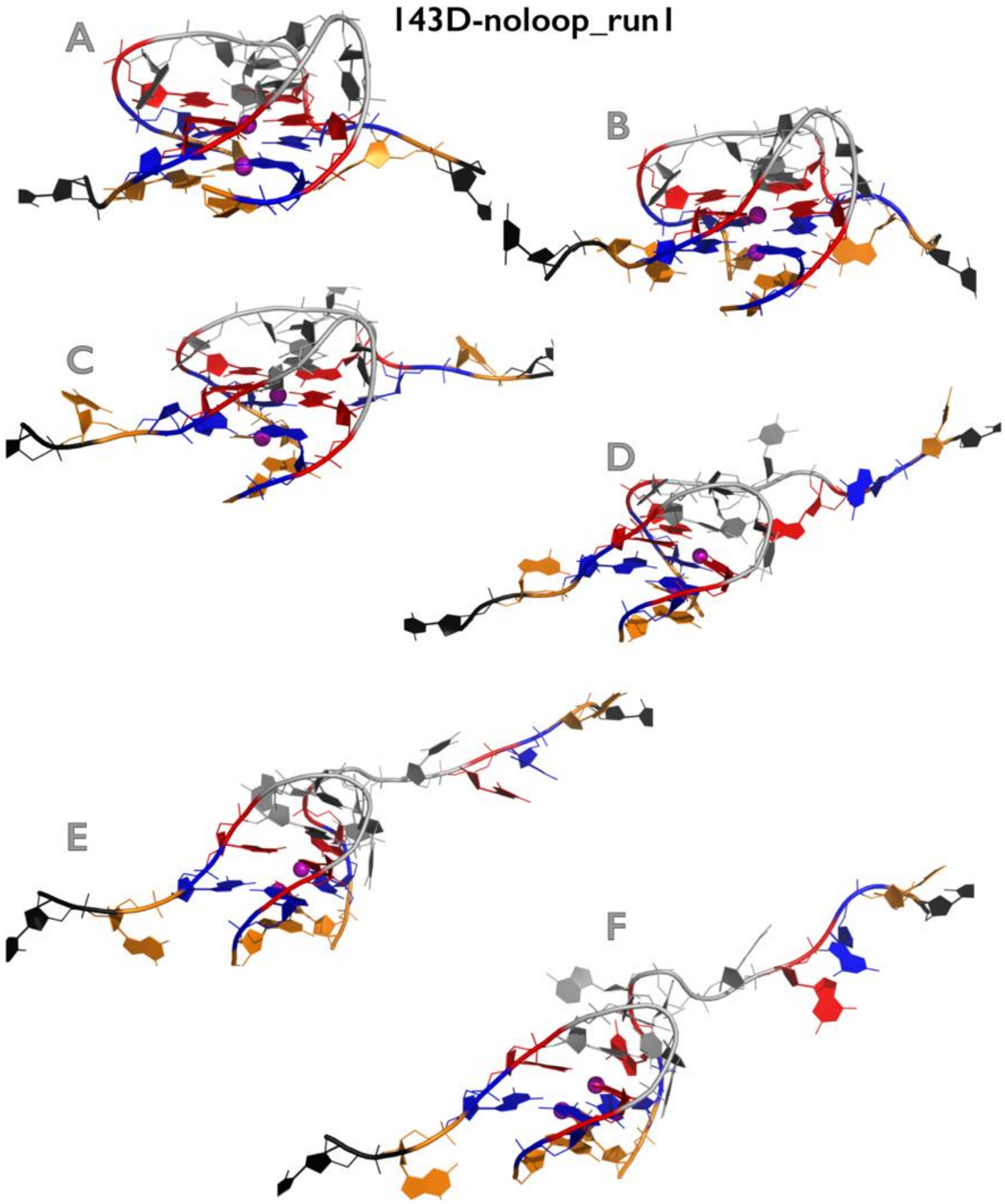

**Figure S14B:** Most important structural events during first independent *slow zig-zag pulling* simulation of $143D_{noloop}$ GQ system. See legend of Figure S1B for more details.

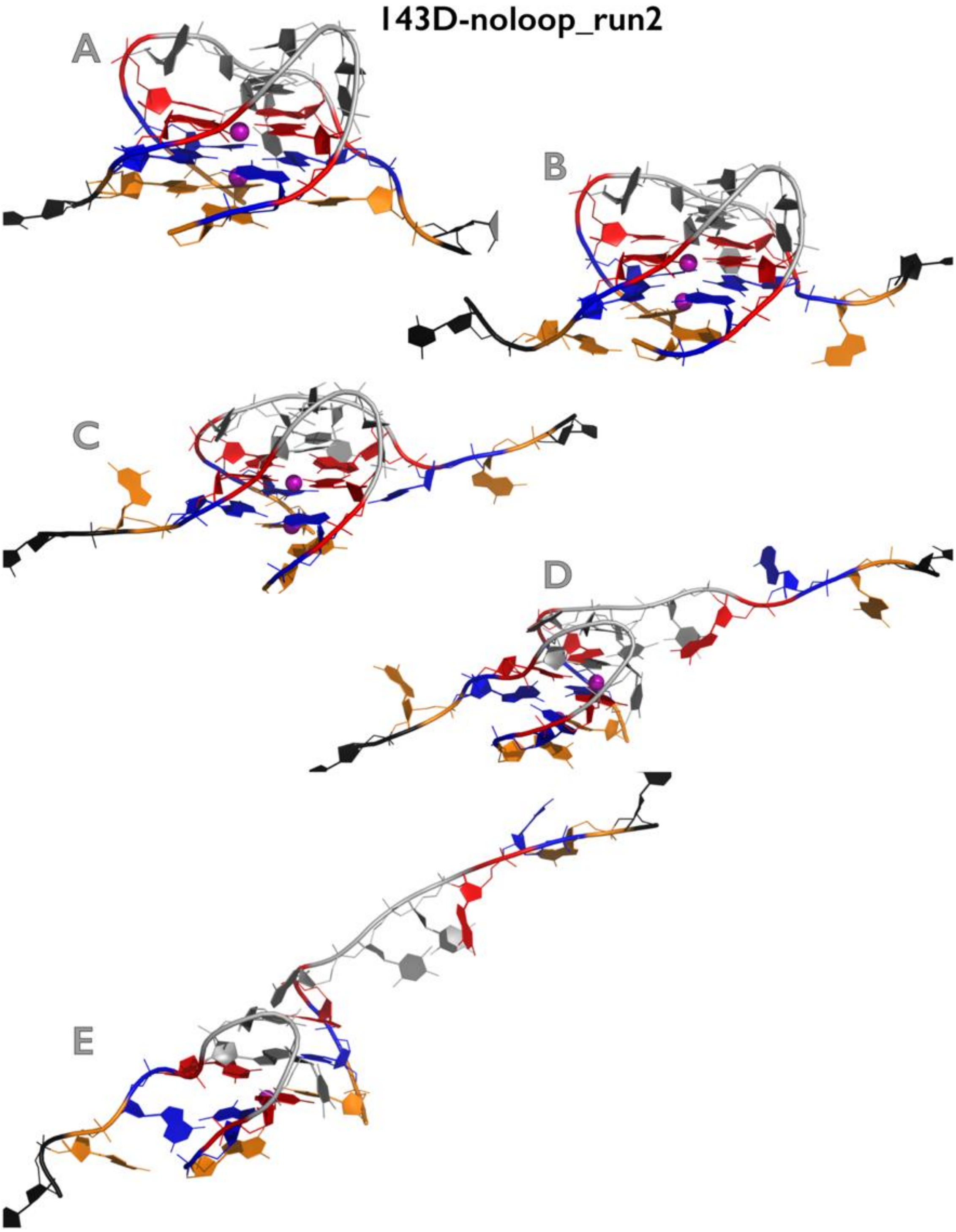

**Figure S14C:** Most important structural events during second independent *slow zig-zag pulling* simulation of 143D$_{noloop}$ GQ system. See legend of Figure S1B for more details.

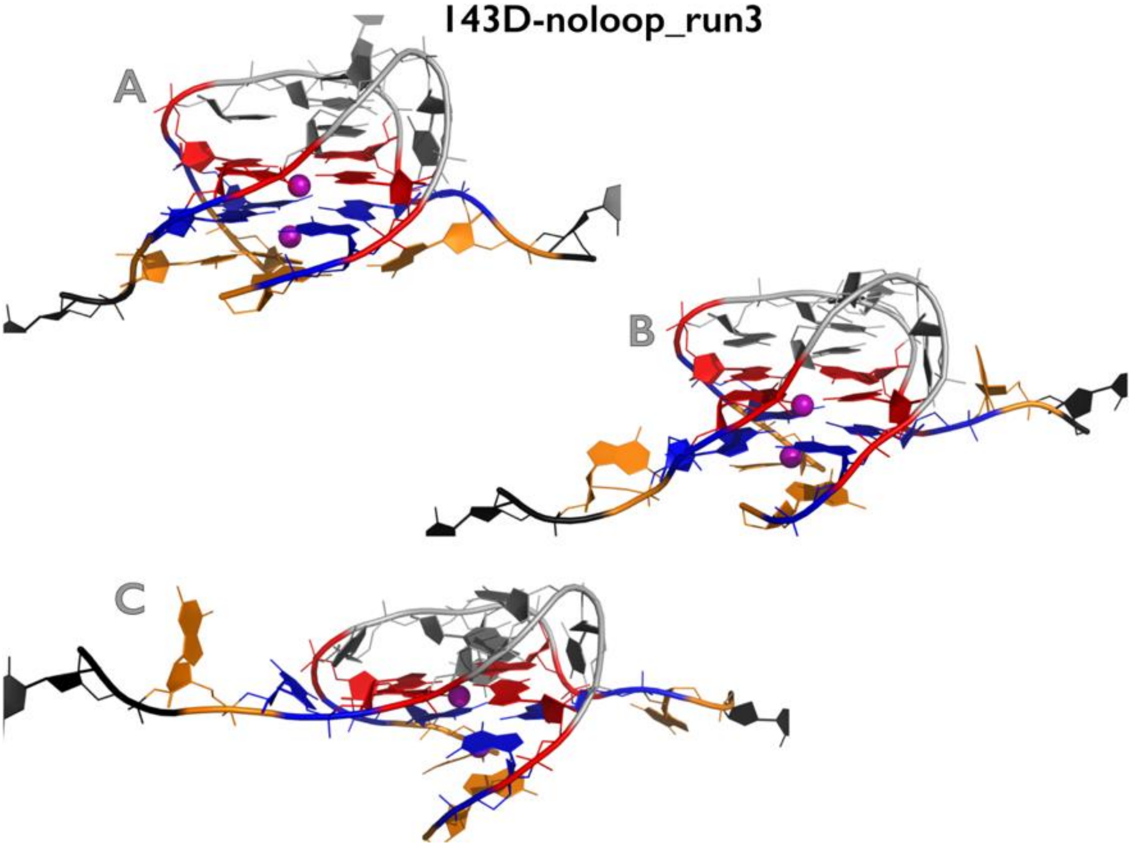

**Figure S14D:** Most important structural events during third independent *slow zig-zag pulling* simulation of $143D_{noloop}$ GQ system. See legend of Figure S1B for more details.

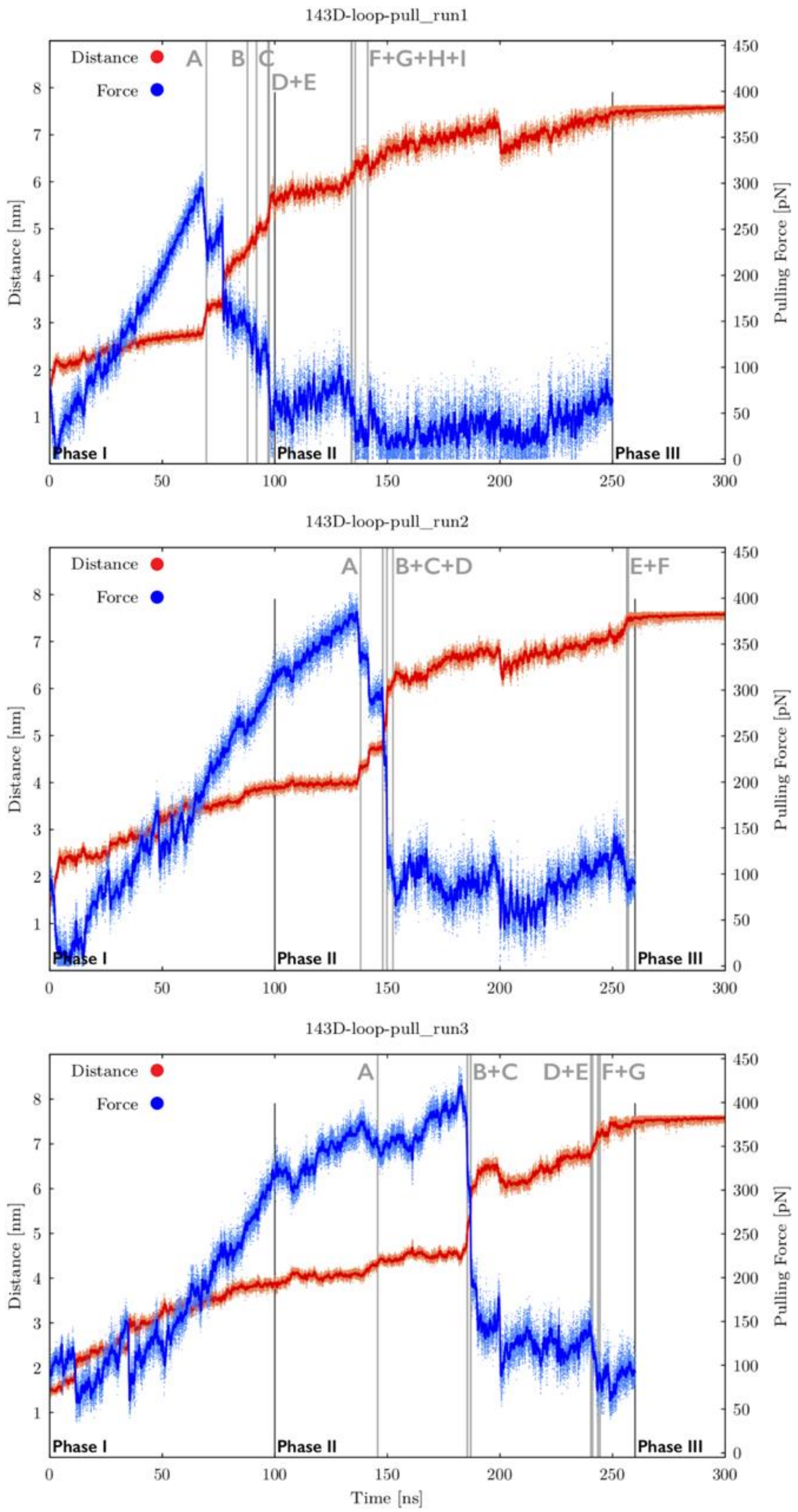

**Figure S15A:** Time evolution of distance between pulling centers and pulling force during three independent *slow zig-zag pulling* simulations of 143D$_{\text{loop-pull}}$ GQ system (see legend of Figure S9A for more details). See Figures S15B-S15D for inspection of structures corresponding to main structural events.

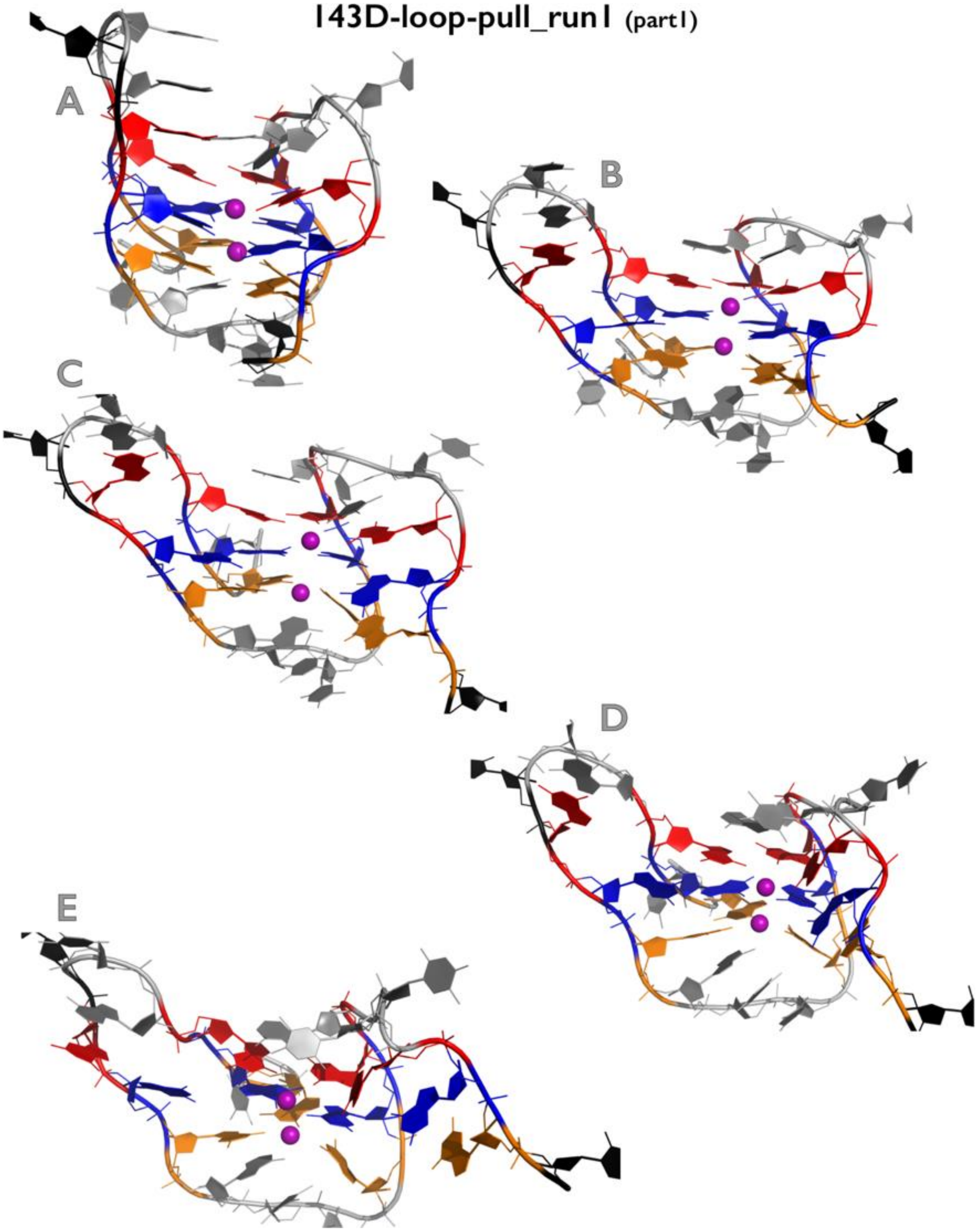

Figure continuing on the next page

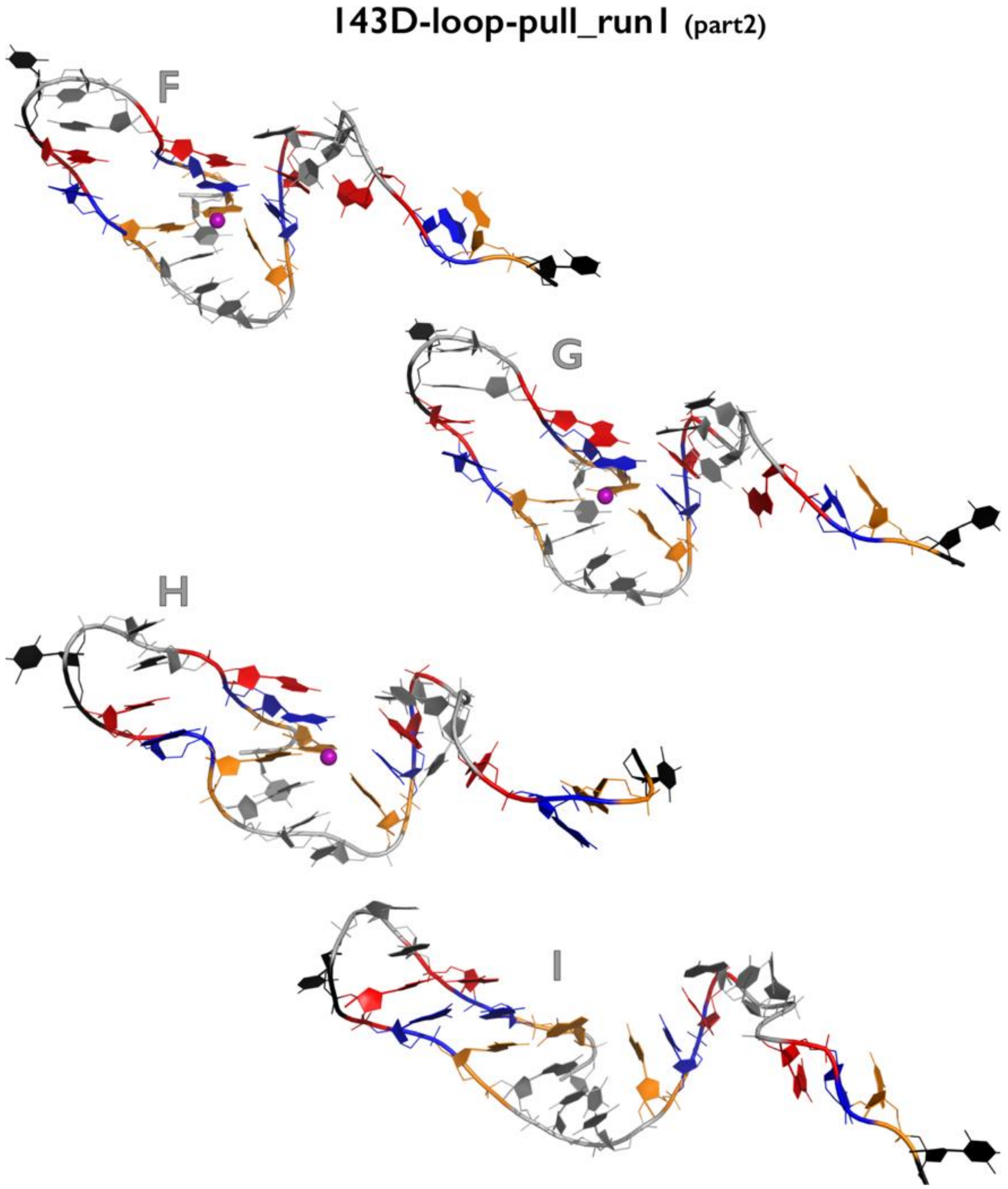

**Figure S15B:** Most important structural events during first independent *slow zig-zag pulling* simulation of $143D_{loop\text{-}pull}$ GQ system. See legend of Figure S1B for more details.

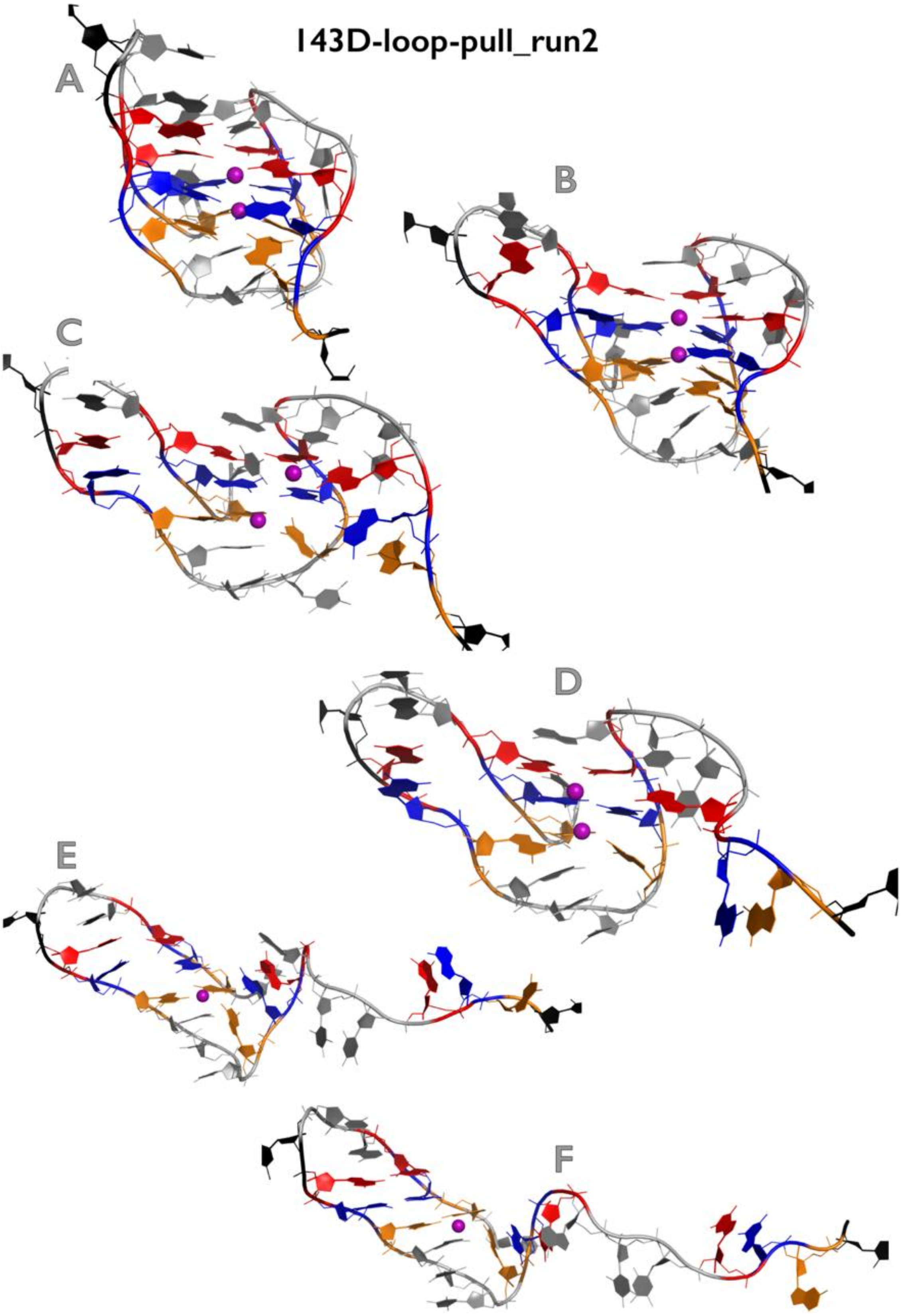

**Figure S15C:** Most important structural events during second independent *slow zig-zag pulling* simulation of 143D$_{loop\text{-}pull}$ GQ system. See legend of Figure S1B for more details.

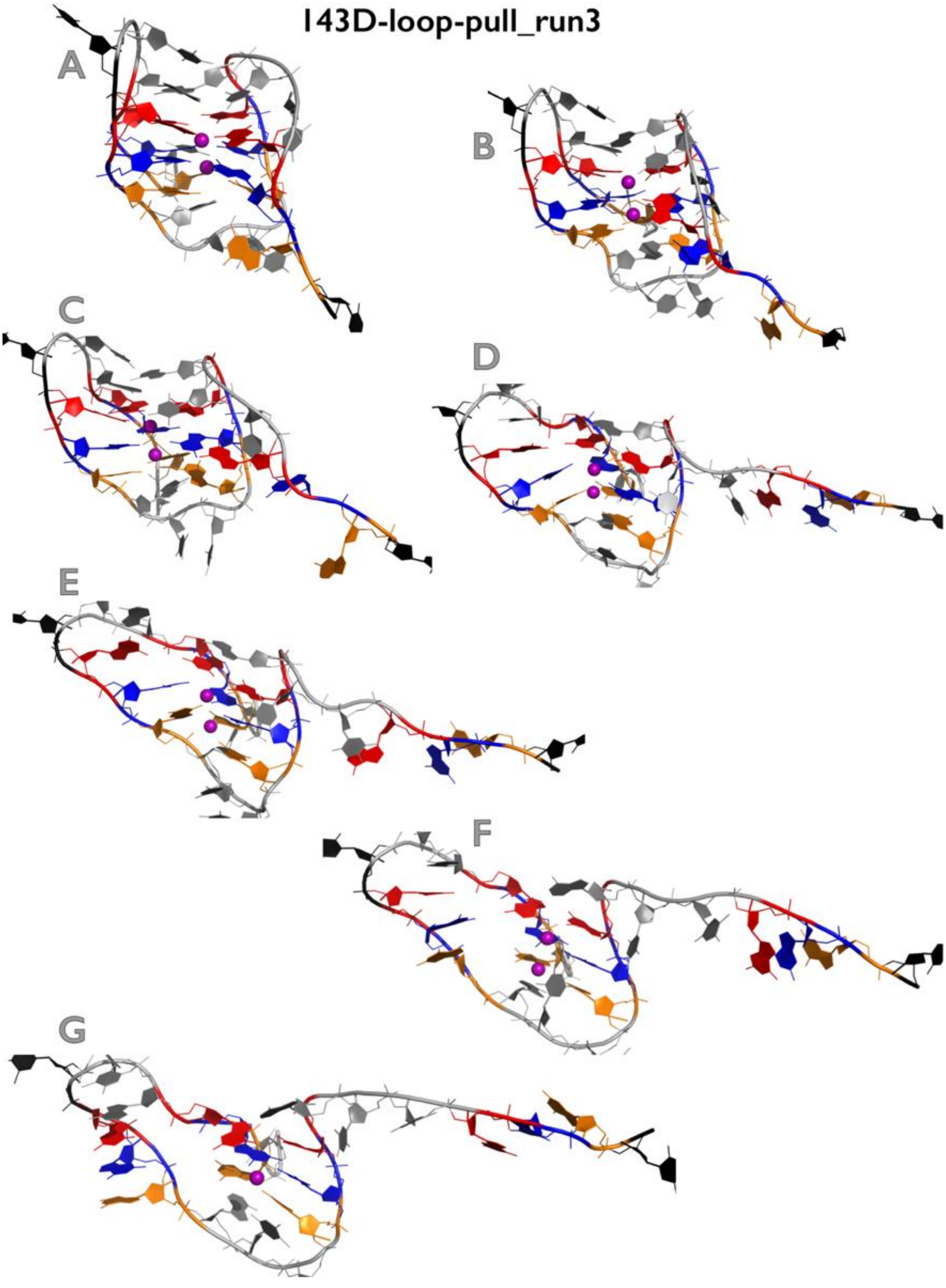

**Figure S15D:** Most important structural events during third independent *slow zig-zag pulling* simulation of $143D_{loop\text{-}pull}$ GQ system. See legend of Figure S1B for more details.

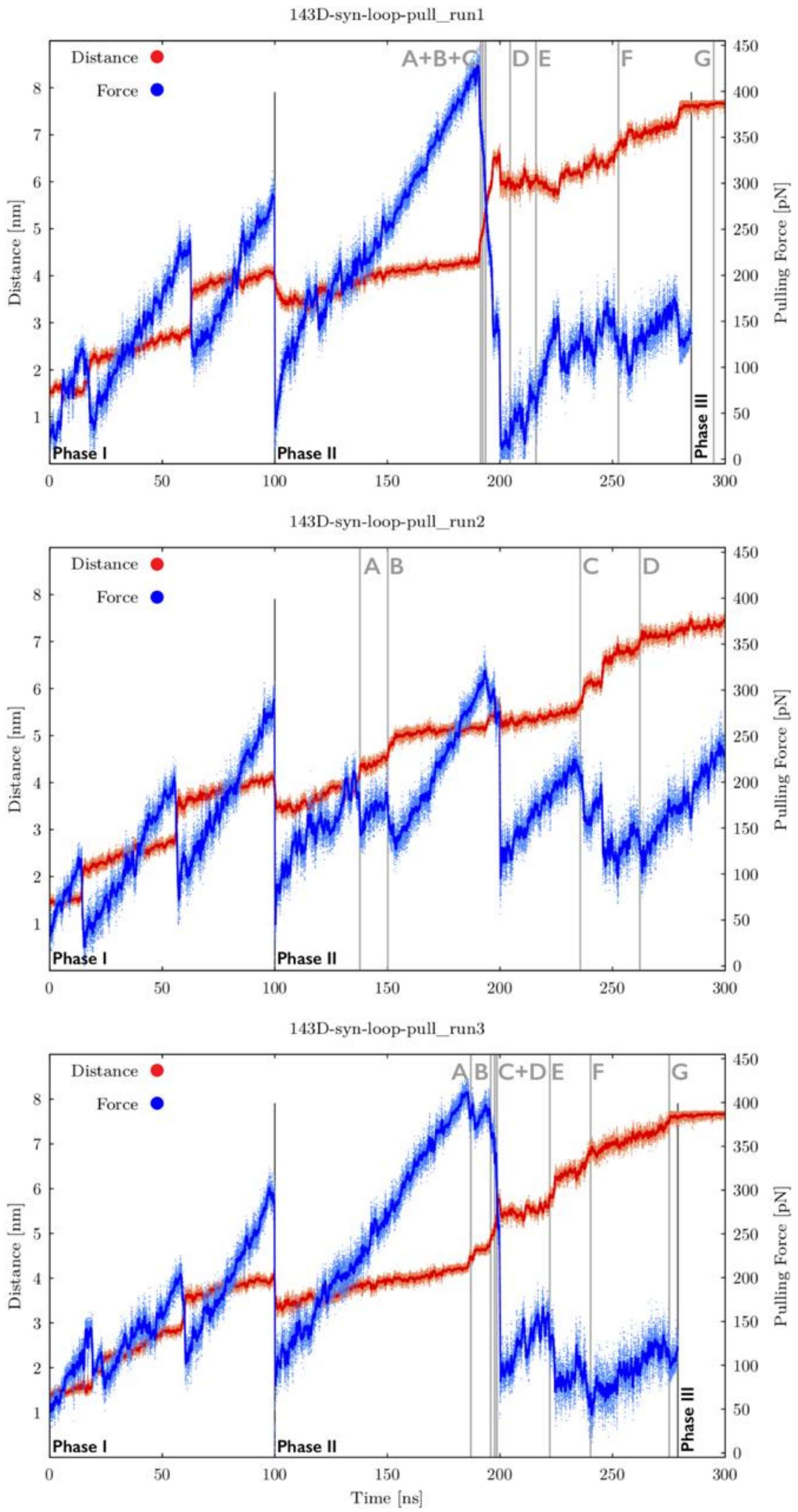

**Figure S16A:** Time evolution of distance between pulling centers and pulling force during three independent *slow zig-zag pulling* simulations of $143D_{syn\_loop-pull}$ GQ system (see legend of Figure S9A for more details). See Figures S16B-S16D for inspection of structures corresponding to main structural events.

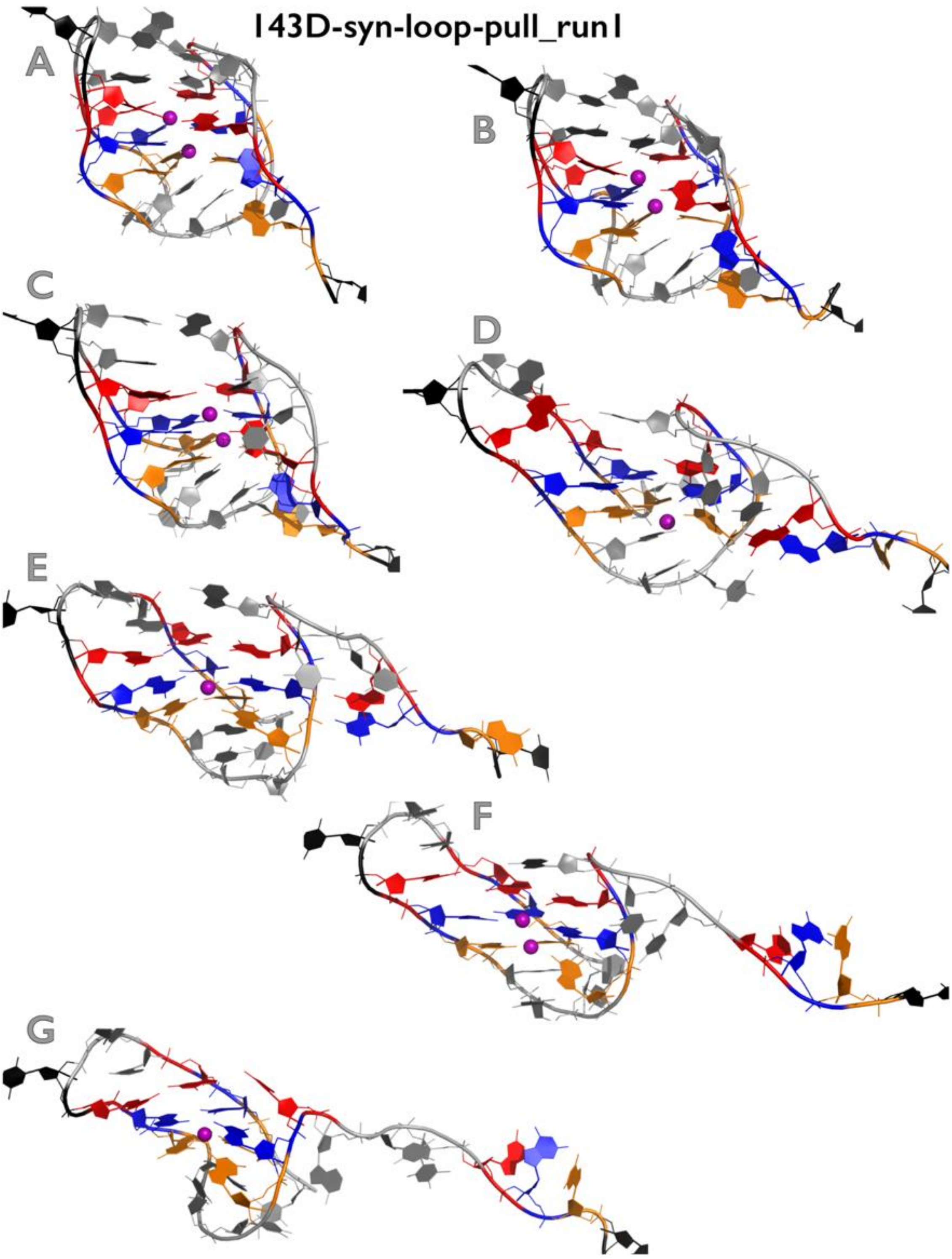

**Figure S16B:** Most important structural events during first independent *slow zig-zag pulling* simulation of 143D$_{syn_loop\text{-}pull}$ GQ system. See legend of Figure S1B for more details.

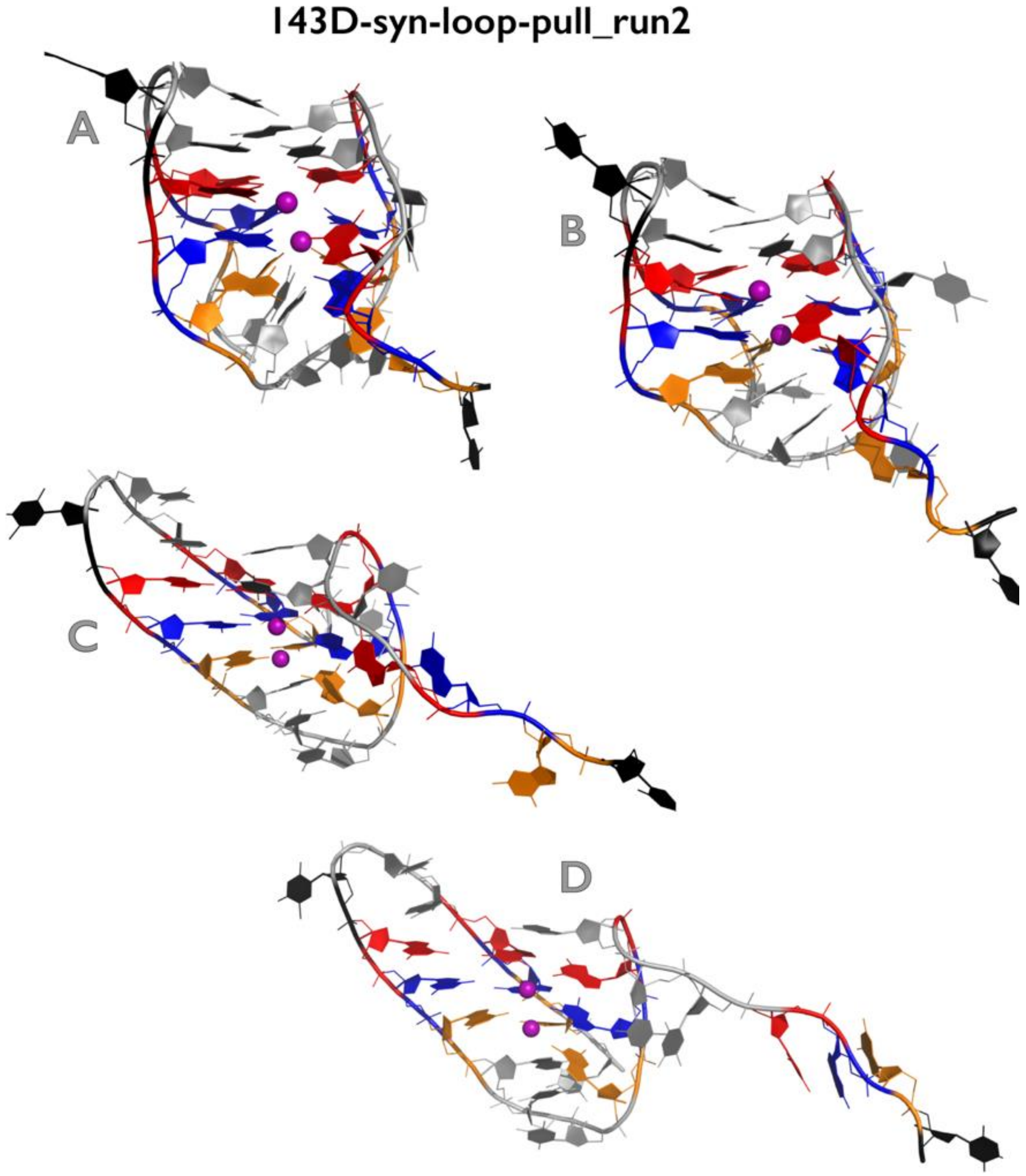

**Figure S16C:** Most important structural events during second independent *slow zig-zag pulling* simulation of $143D_{syn\_loop\text{-}pull}$ GQ system. See legend of Figure S1B for more details.

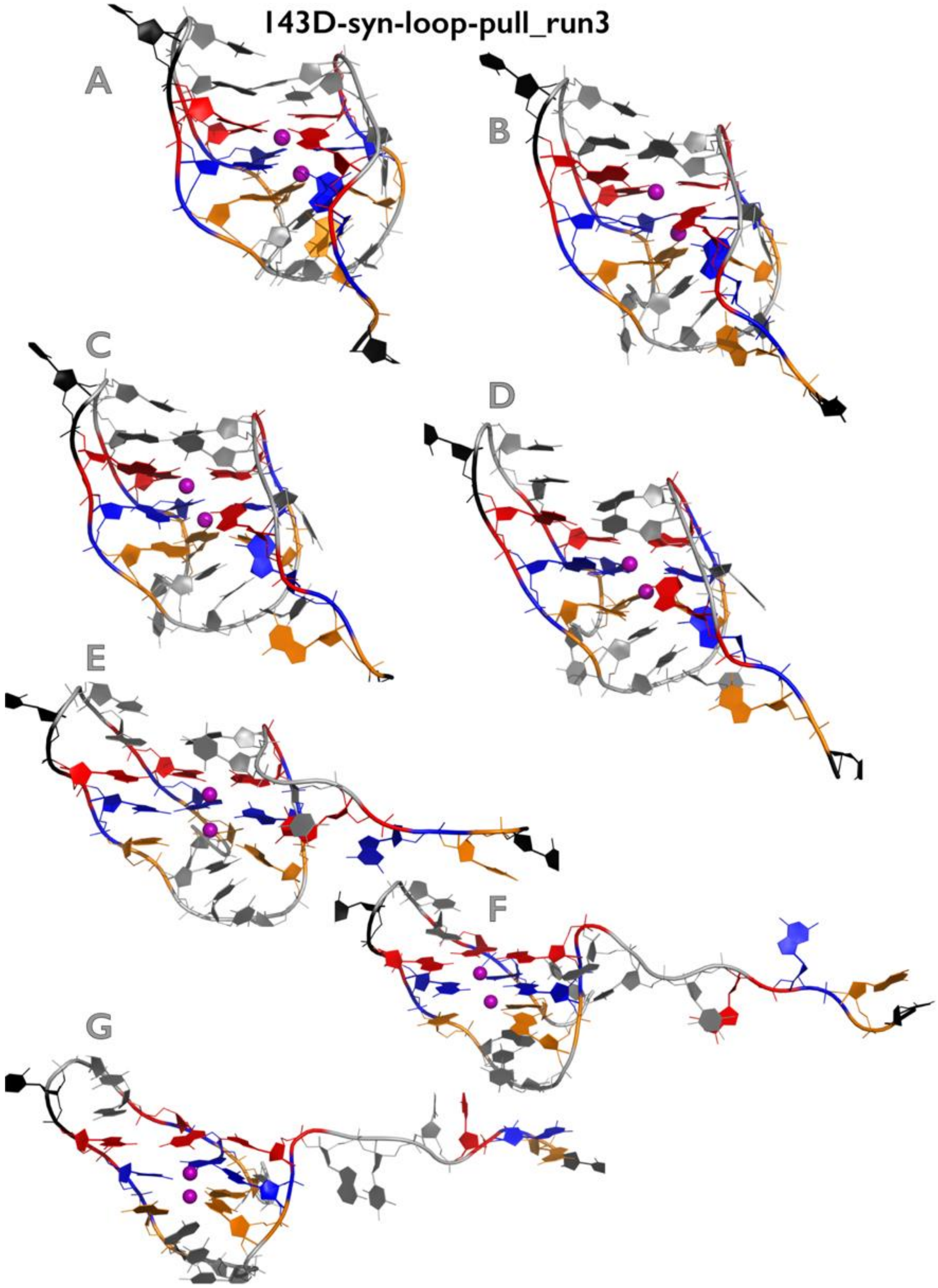

**Figure S16D:** Most important structural events during third independent *slow zig-zag pulling* simulation of 143D$_{syn_loop\text{-}pull}$ GQ system. See legend of Figure S1B for more details.

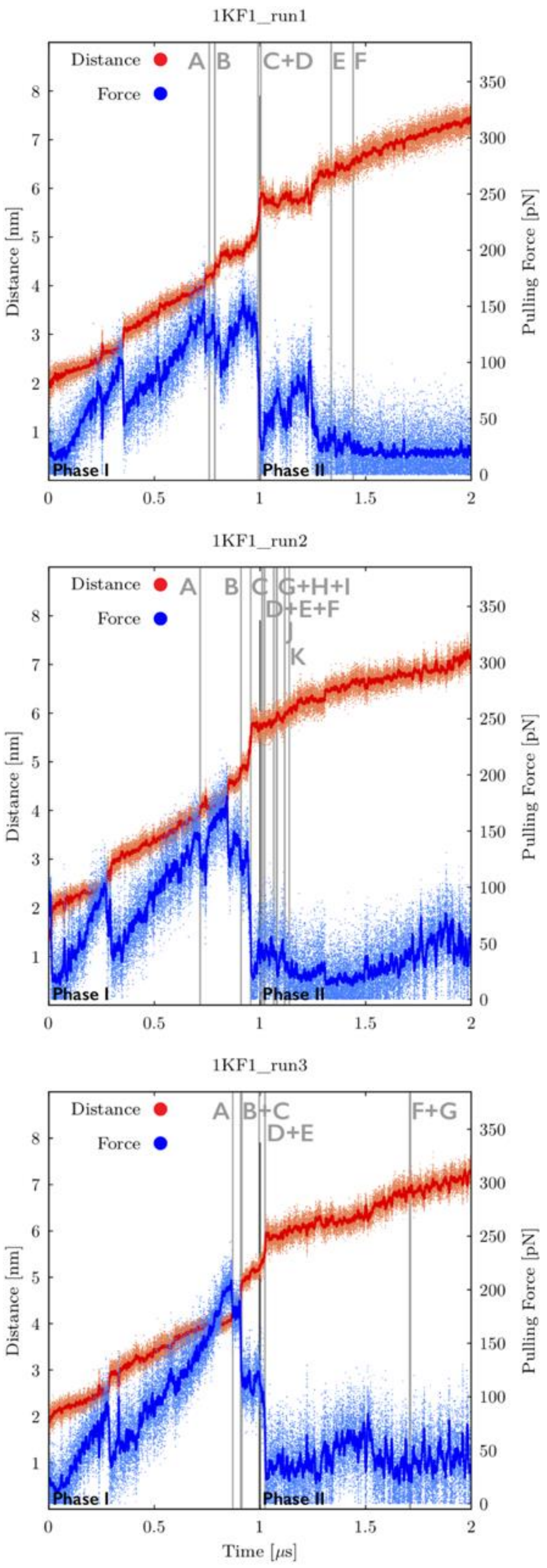

**Figure S17A:** Time evolution of distance between pulling centers and pulling force during three independent *very slow zig-zag pulling* simulations of 1KF1 GQ system. Snapshots were saved every 50 ps and plots are showing both instantaneous values (orange and light-blue dots for distance and force, respectively) and smoothing, i.e., averaging over 100 consecutive snapshots (red and blue lines for distance and force, respectively). Pulling phases are marked (see Methods in the main text for details) and main structural events are highlighted as grey vertical lines with labels (capital letters). See Figures S17B-S17D for inspection of structures corresponding to main structural events. Note that first major drops of the pulling force before the GQ unfolding event "A" (and notable prolongation of end-to-end distances) are connected with repositioning of terminal T residues.

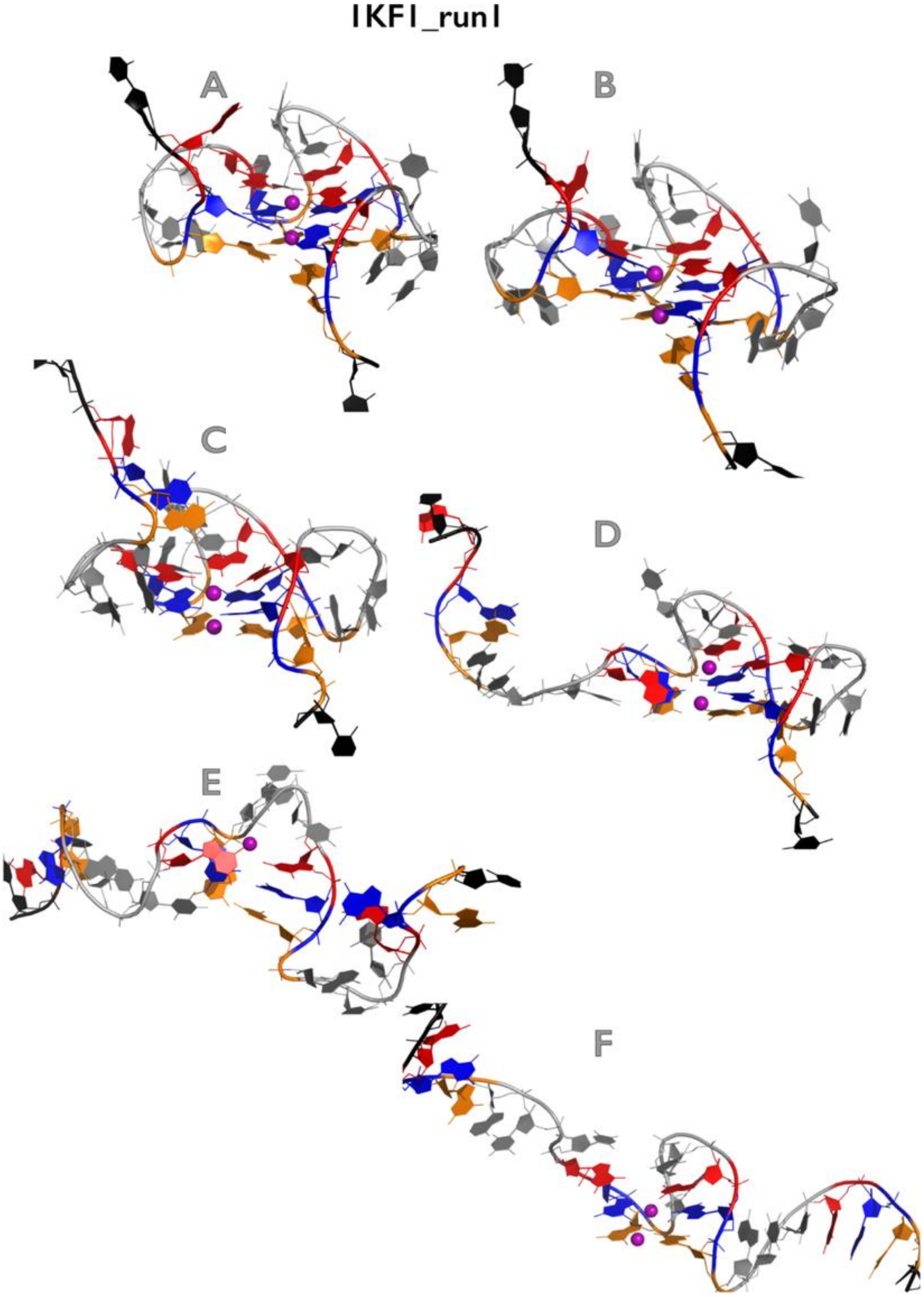

**Figure S17B:** Most important structural events during first independent *very slow zig-zag pulling* simulation of 1KF1 GQ system. See legend of Figure S1B for more details.

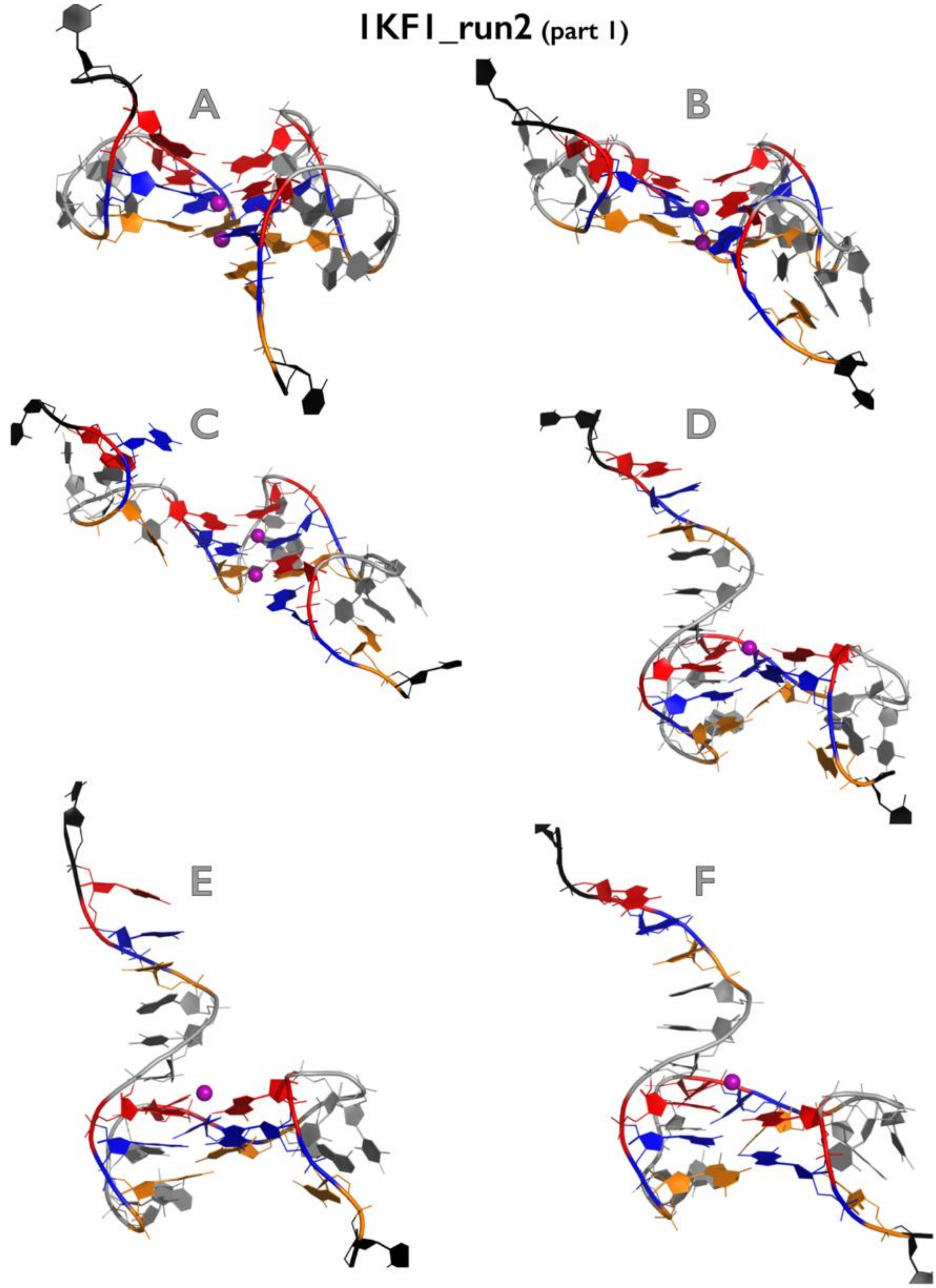

Figure continuing on the next page

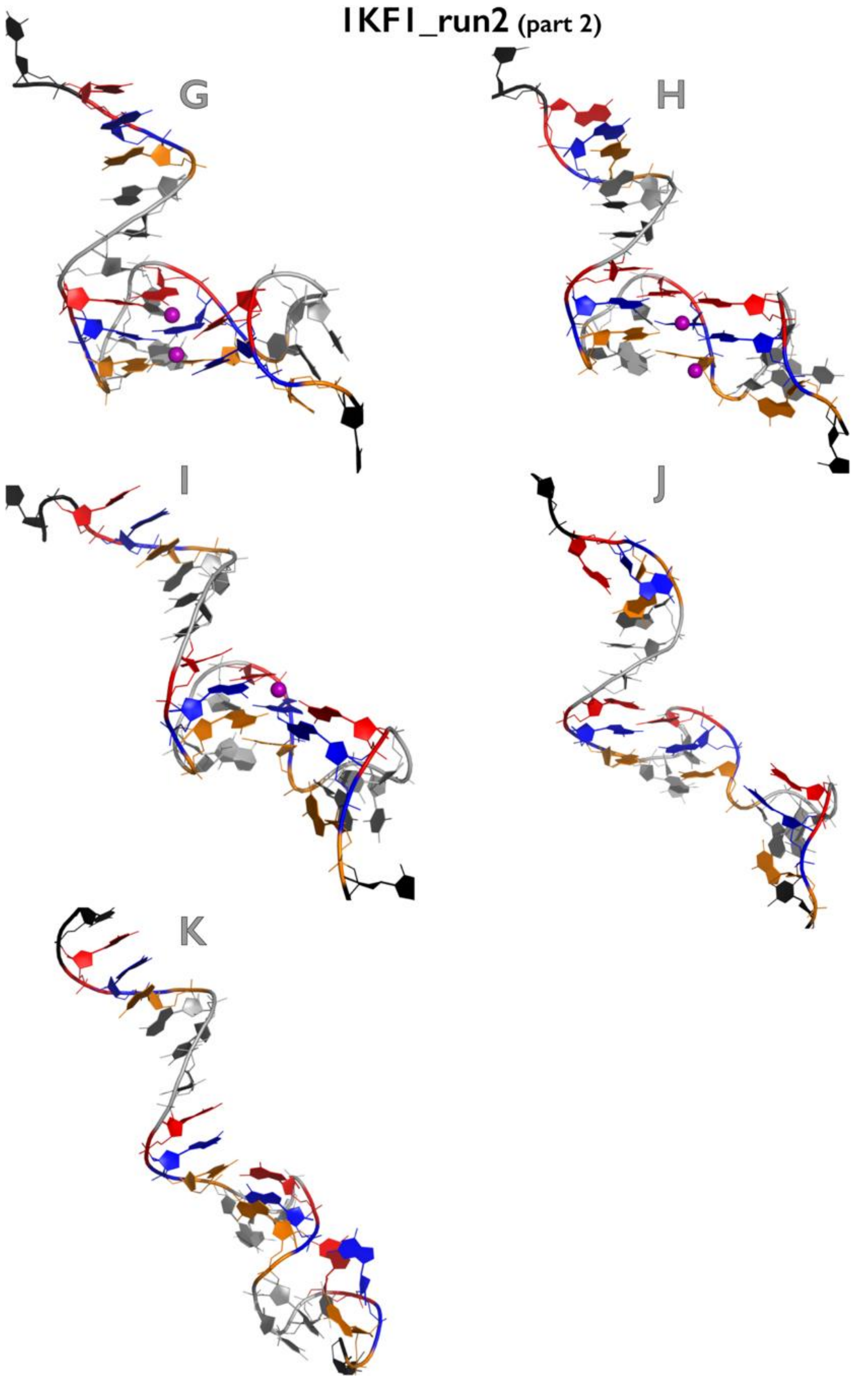

**Figure S17C:** Most important structural events during second independent *very slow zig-zag pulling* simulation of 1KF1 GQ system. See legend of Figure S1B for more details.

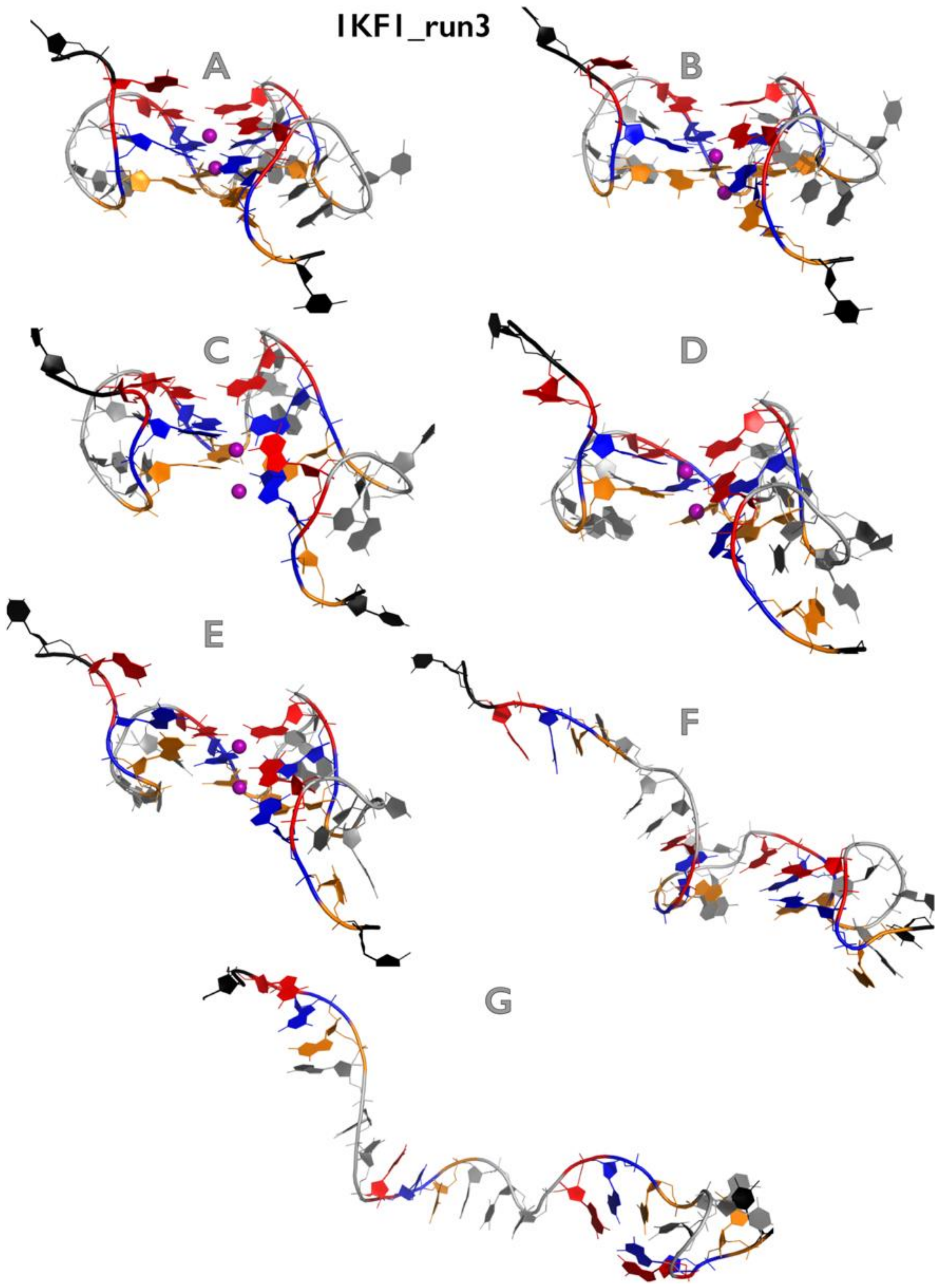

**Figure S17D:** Most important structural events during third independent *very slow zig-zag pulling* simulation of 1KF1 GQ system. See legend of Figure S1B for more details.

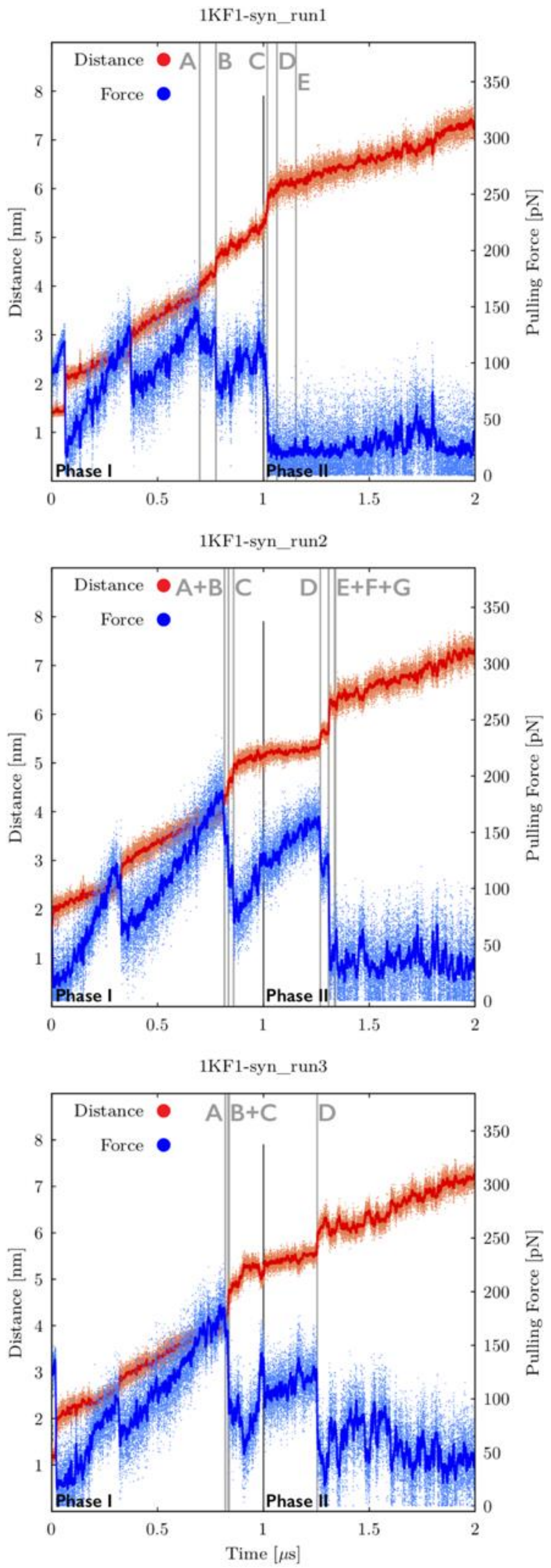

**Figure S18A:** Time evolution of distance between pulling centers and pulling force during three independent *very slow zig-zag pulling* simulations of 1KF1$_{syn}$ GQ system (see legend of Figure S17A for more details). See Figures S18B-S18D for inspection of structures corresponding to main structural events.

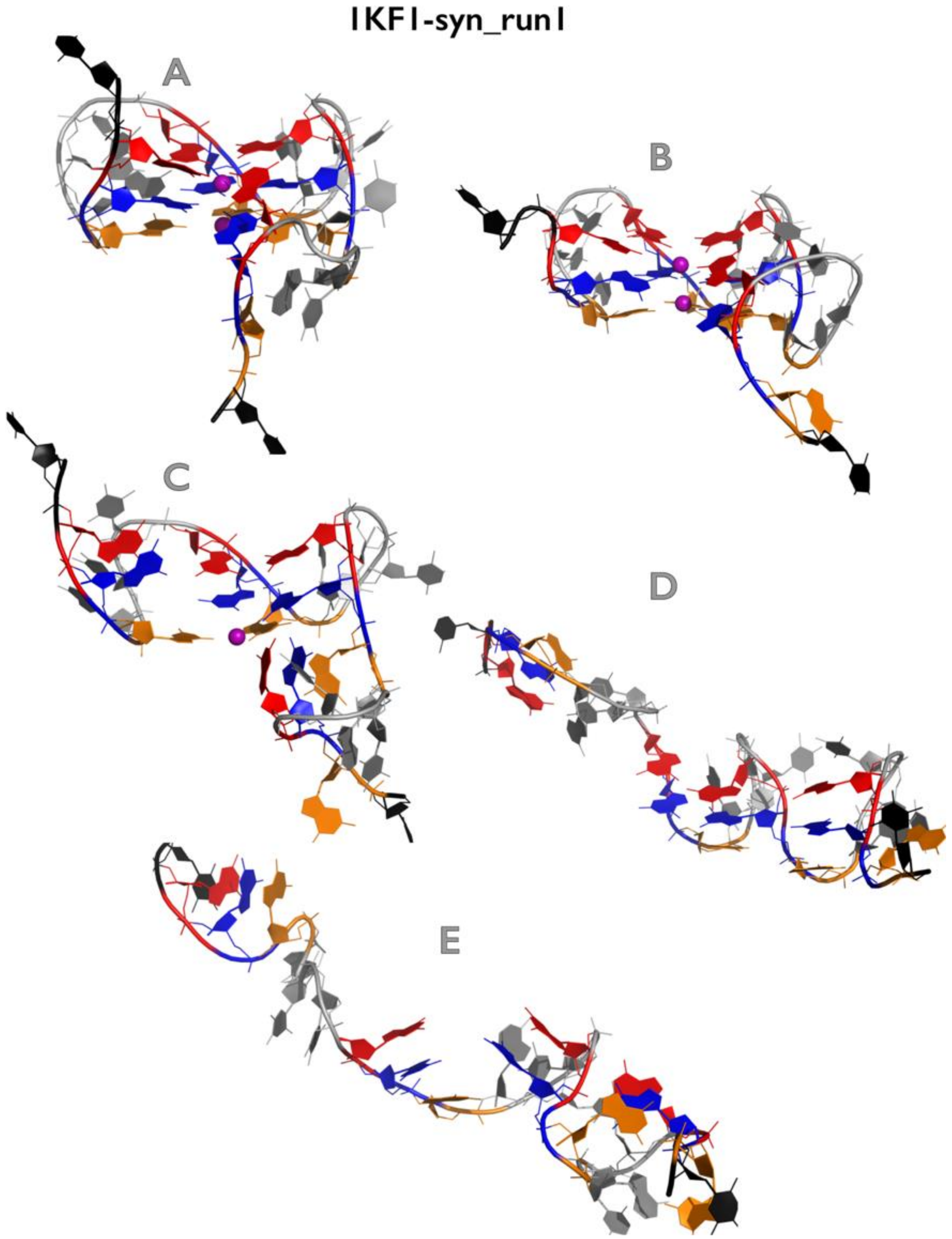

**Figure S18B:** Most important structural events during first independent *very slow zig-zag pulling* simulation of $1KF1_{syn}$ GQ system. See legend of Figure S1B for more details.

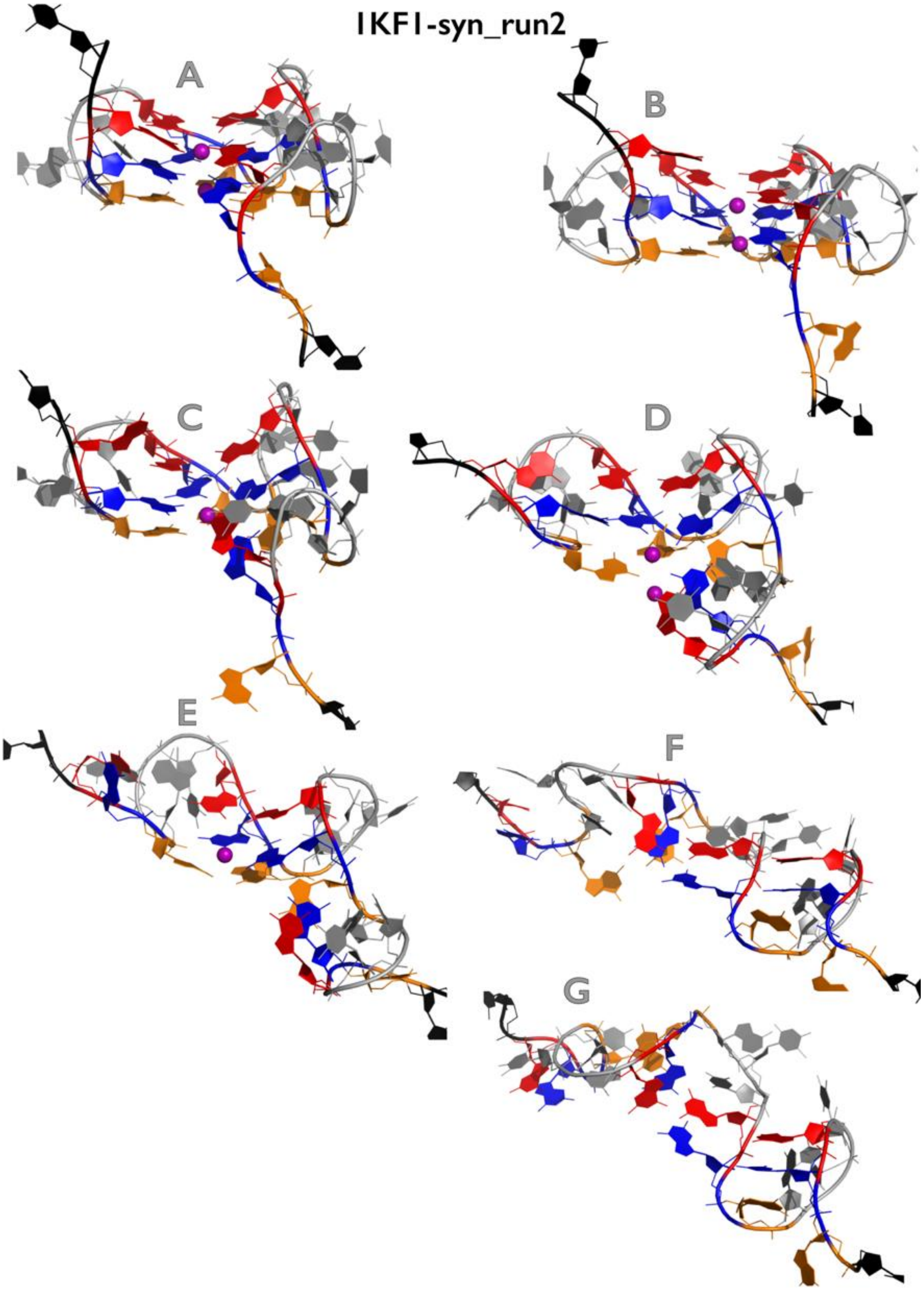

**Figure S18C:** Most important structural events during second independent *very slow zig-zag pulling* simulation of $1KF1_{syn}$ GQ system. See legend of Figure S1B for more details.

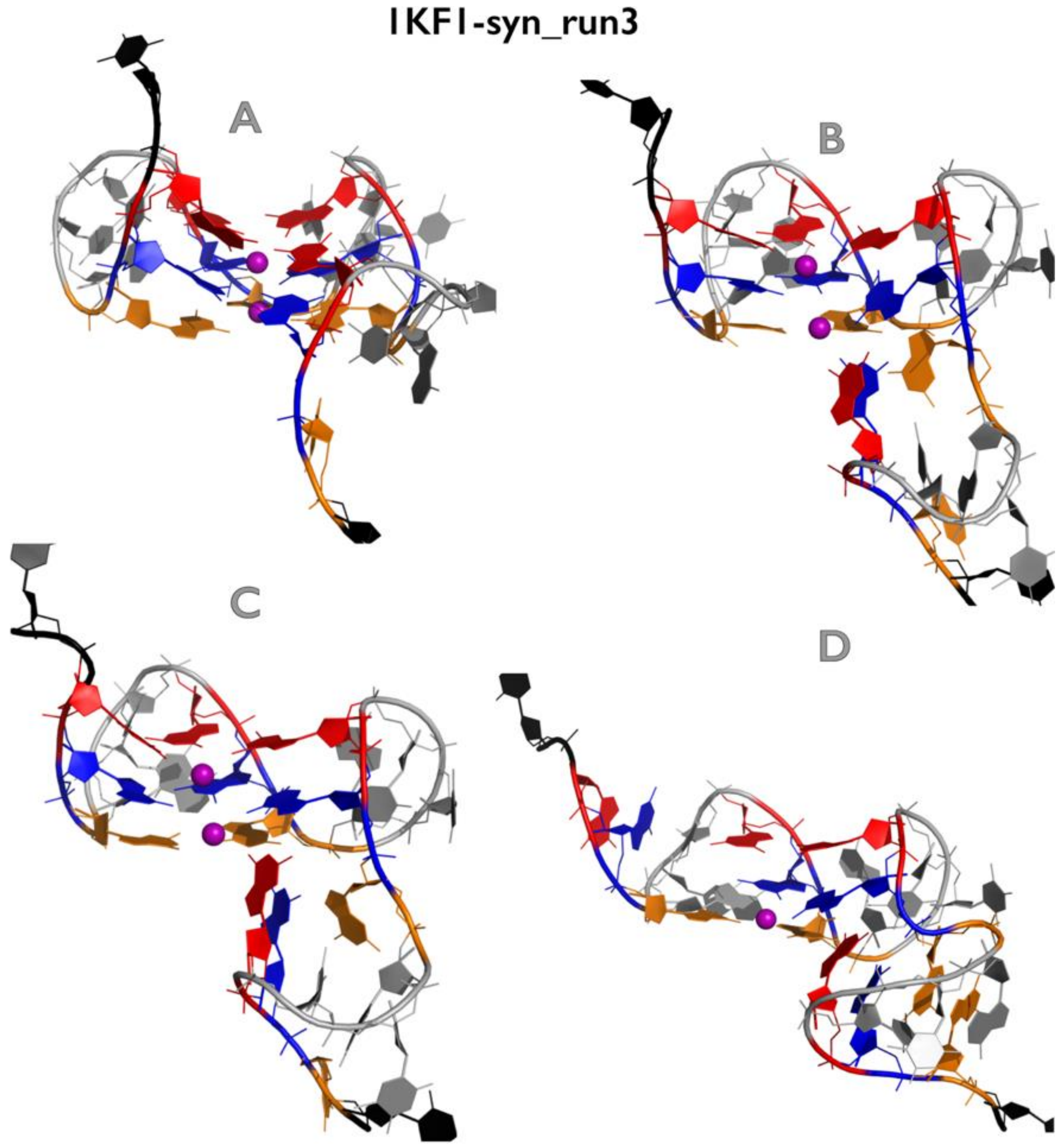

**Figure S18D:** Most important structural events during third independent *very slow zig-zag pulling* simulation of $1KF1_{syn}$ GQ system. See legend of Figure S1B for more details.

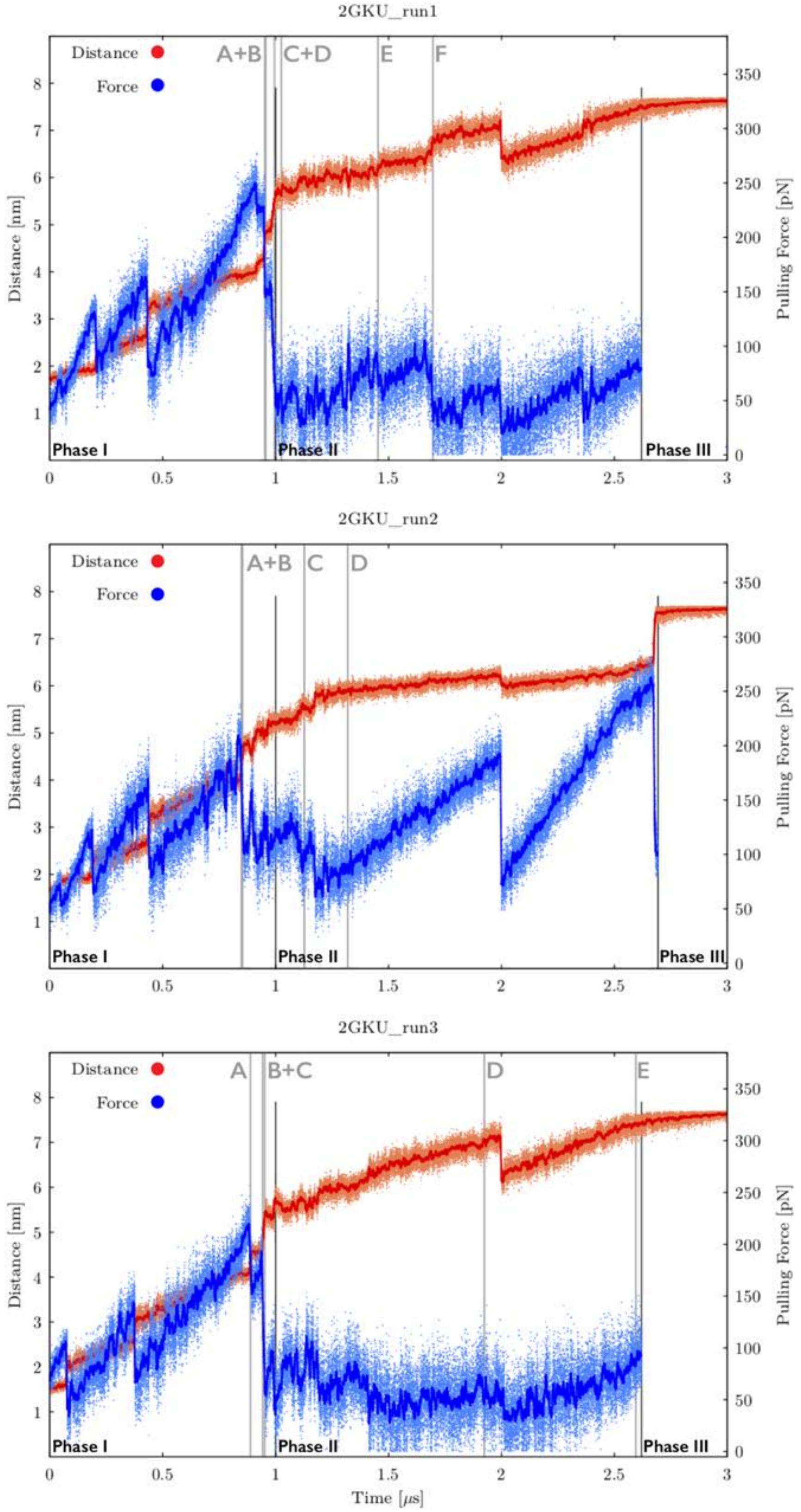

**Figure S19A:** Time evolution of distance between pulling centers and pulling force during three independent *very slow zig-zag pulling* simulations of 2GKU GQ system (see legend of Figure S17A for more details). See Figures S19B-S19D for inspection of structures corresponding to main structural events.

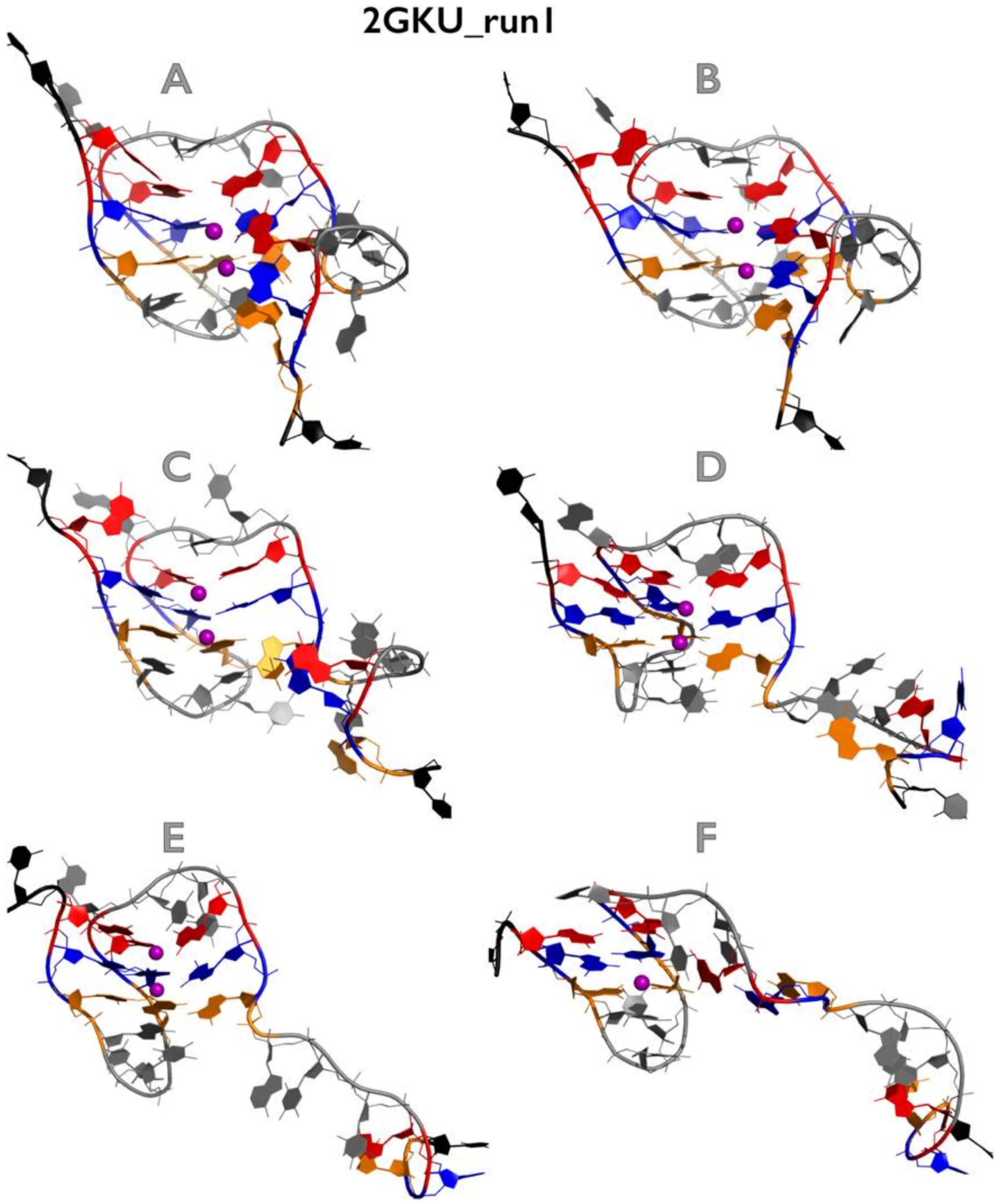

**Figure S19B:** Most important structural events during first independent *very slow zig-zag pulling* simulation of 2GKU GQ system. See legend of Figure S1B for more details.

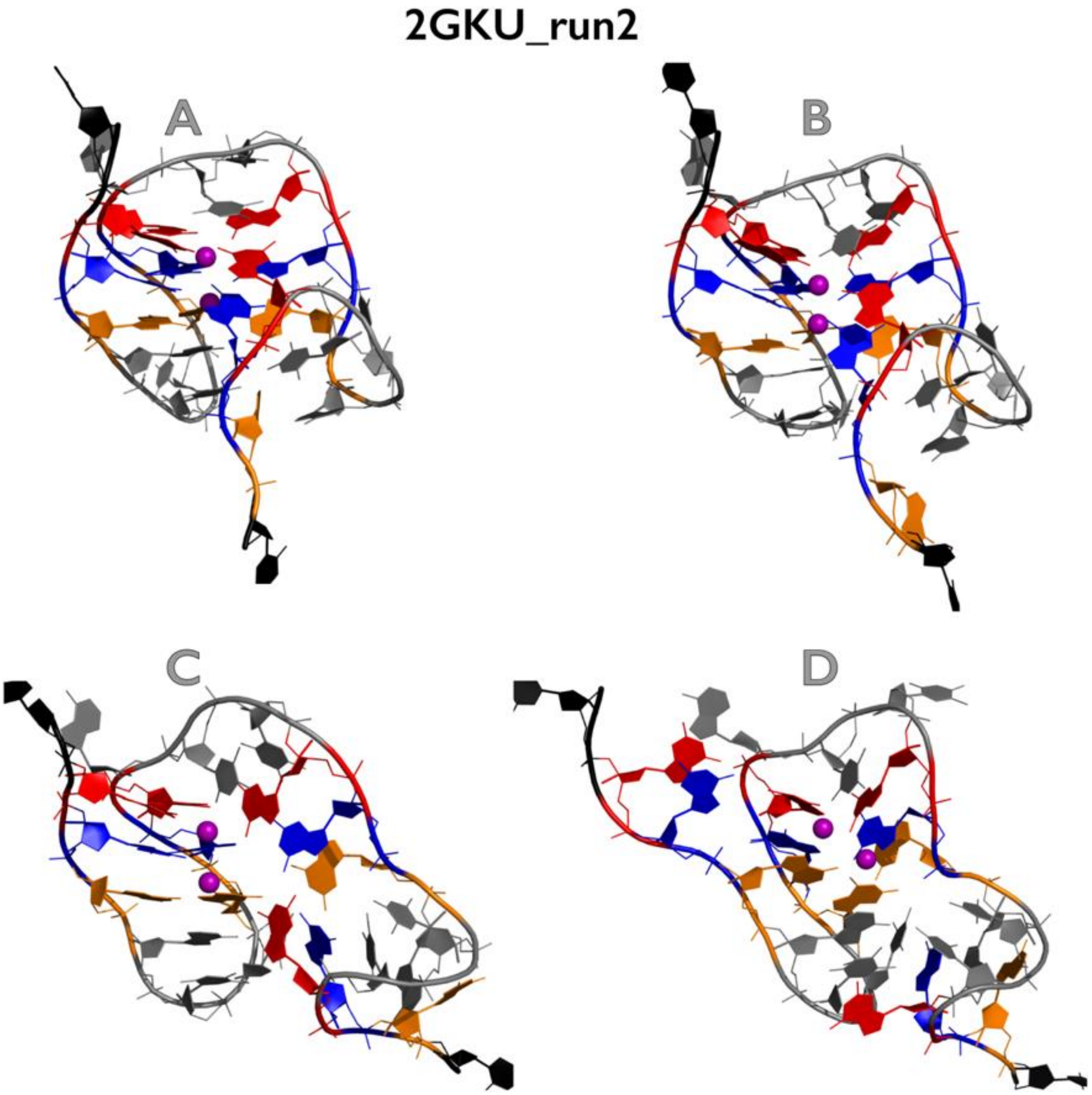

**Figure S19C:** Most important structural events during second independent *very slow zig-zag pulling* simulation of 2GKU GQ system. See legend of Figure S1B for more details.

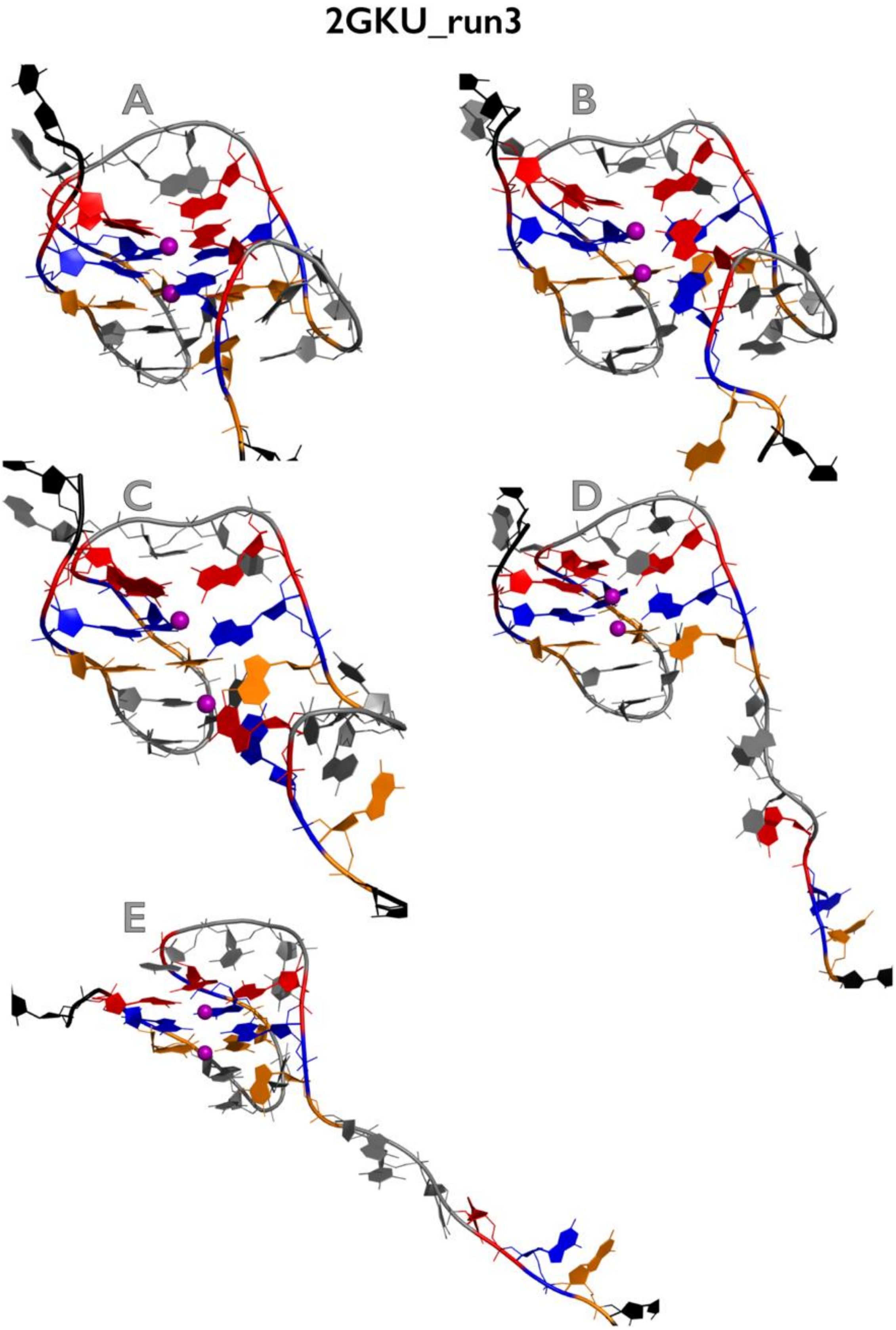

**Figure S19D:** Most important structural events during third independent *very slow zig-zag pulling* simulation of 2GKU GQ system. See legend of Figure S1B for more details.

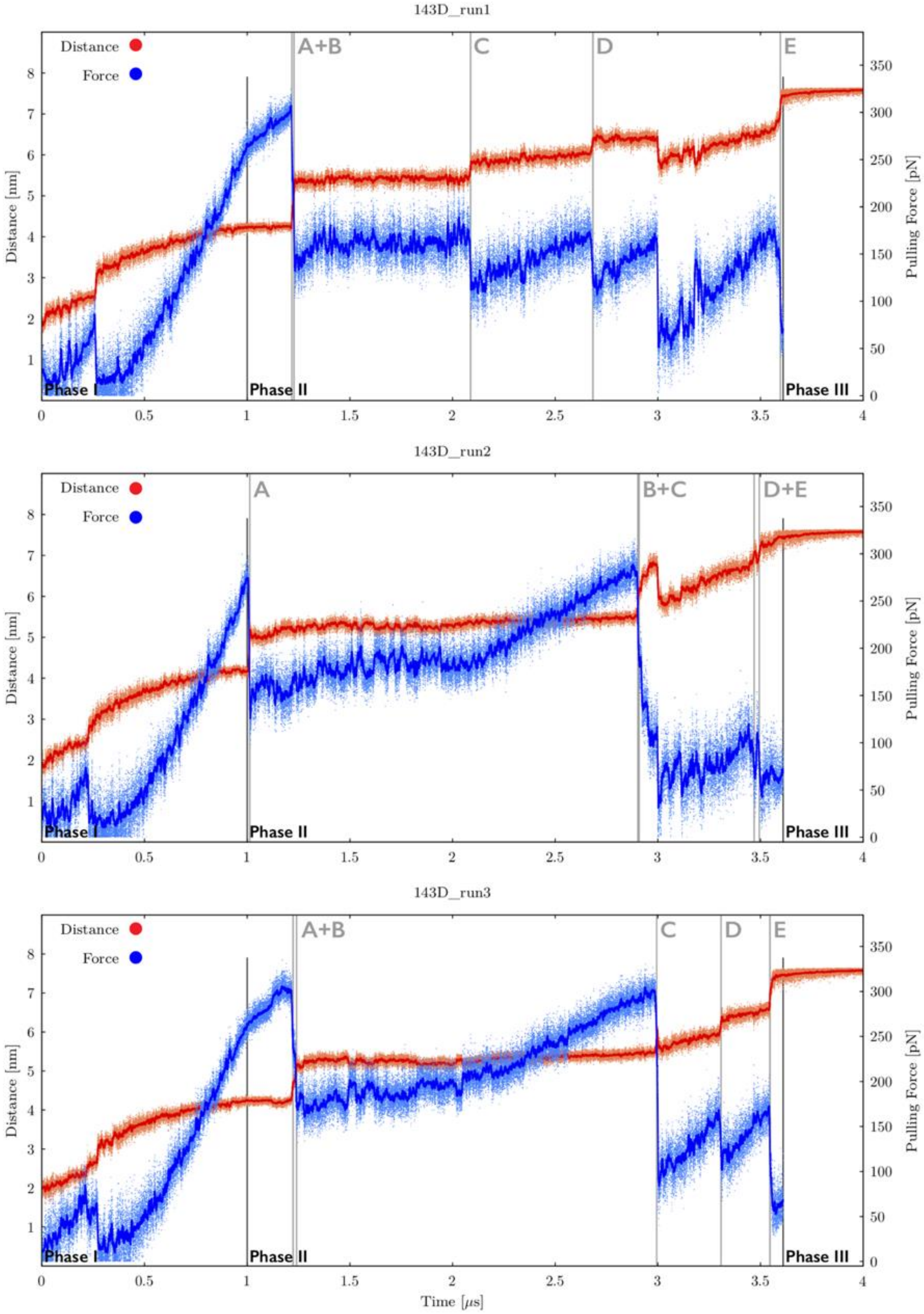

**Figure S20A:** Time evolution of distance between pulling centers and pulling force during three independent *very slow zig-zag pulling* simulations of 143D GQ system (see legend of Figure S17A for more details). See Figures S20B-S20D for inspection of structures corresponding to main structural events.

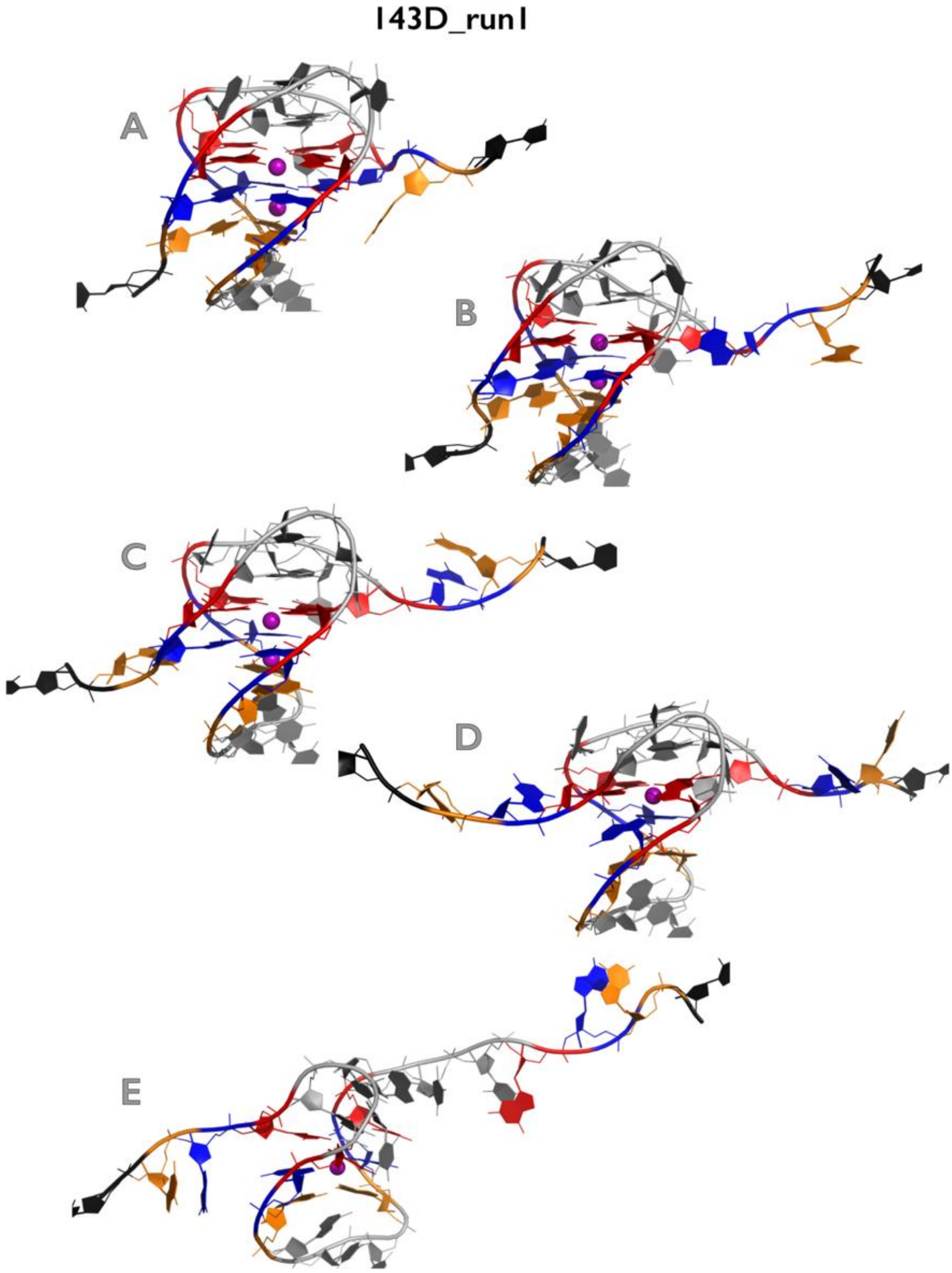

**Figure S20B:** Most important structural events during first independent *very slow zig-zag pulling* simulation of 143D GQ system. See legend of Figure S1B for more details.

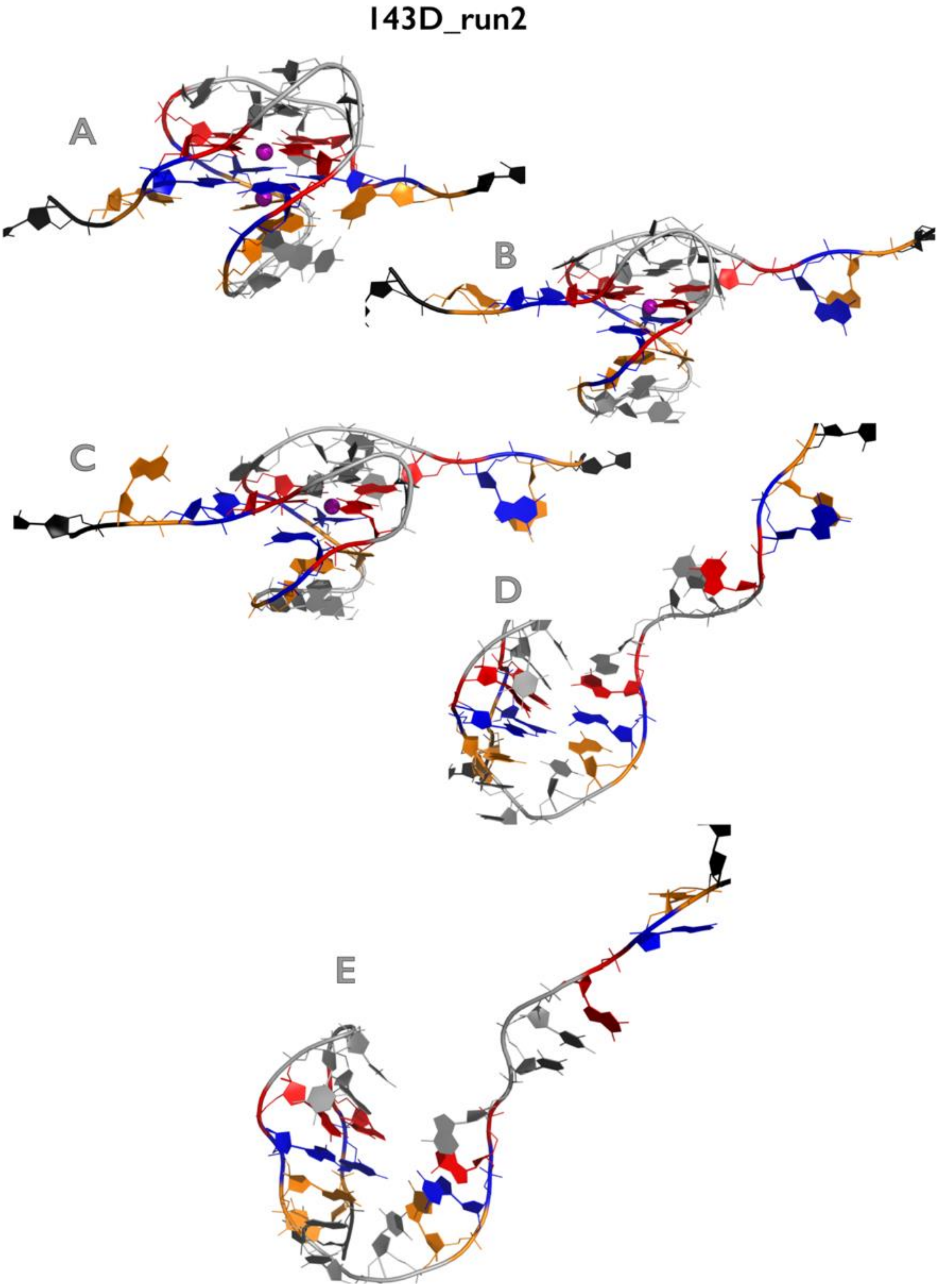

**Figure S20C:** Most important structural events during second independent *very slow zig-zag pulling* simulation of 143D GQ system. See legend of Figure S1B for more details.

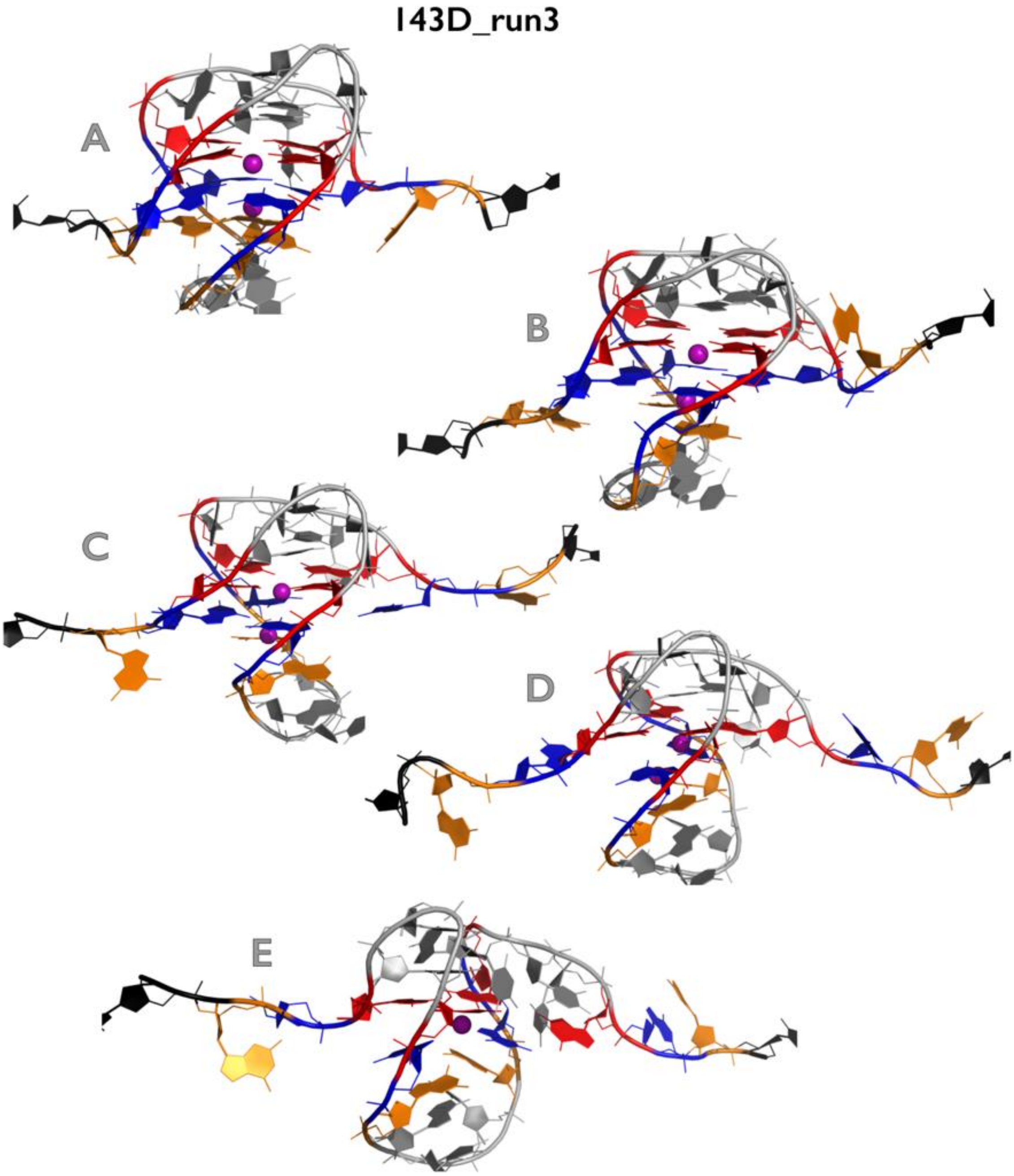

**Figure S20D:** Most important structural events during third independent *very slow zig-zag pulling* simulation of 143D GQ system. See legend of Figure S1B for more details.

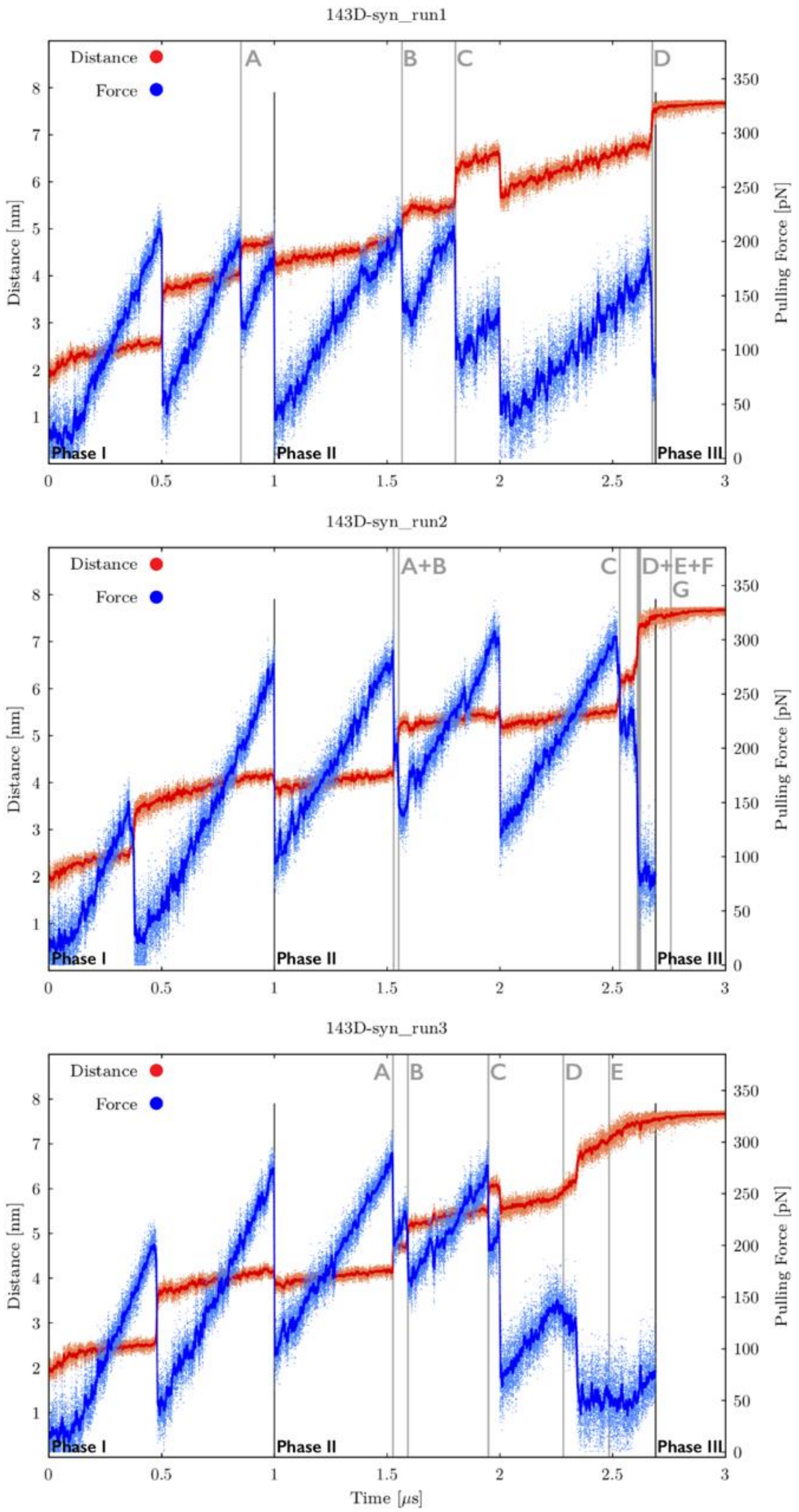

**Figure S21A:** Time evolution of distance between pulling centers and pulling force during three independent *very slow zig-zag pulling* simulations of $143D_{syn}$ GQ system (see legend of Figure S17A for more details). See Figures S21B-S21D for inspection of structures corresponding to main structural events.

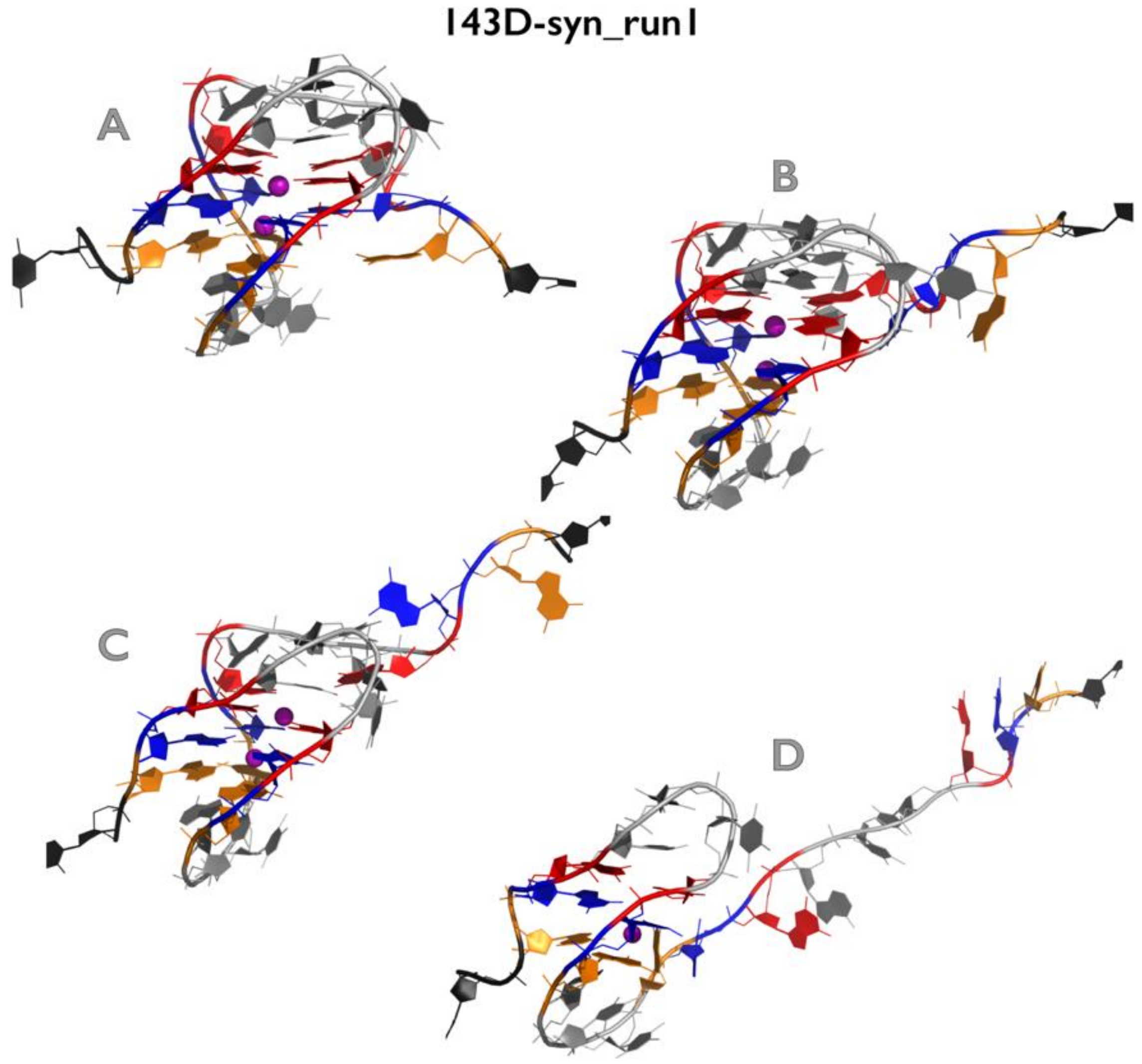

**Figure S21B:** Most important structural events during first independent *very slow zig-zag pulling* simulation of $143D_{syn}$ GQ system. See legend of Figure S1B for more details.

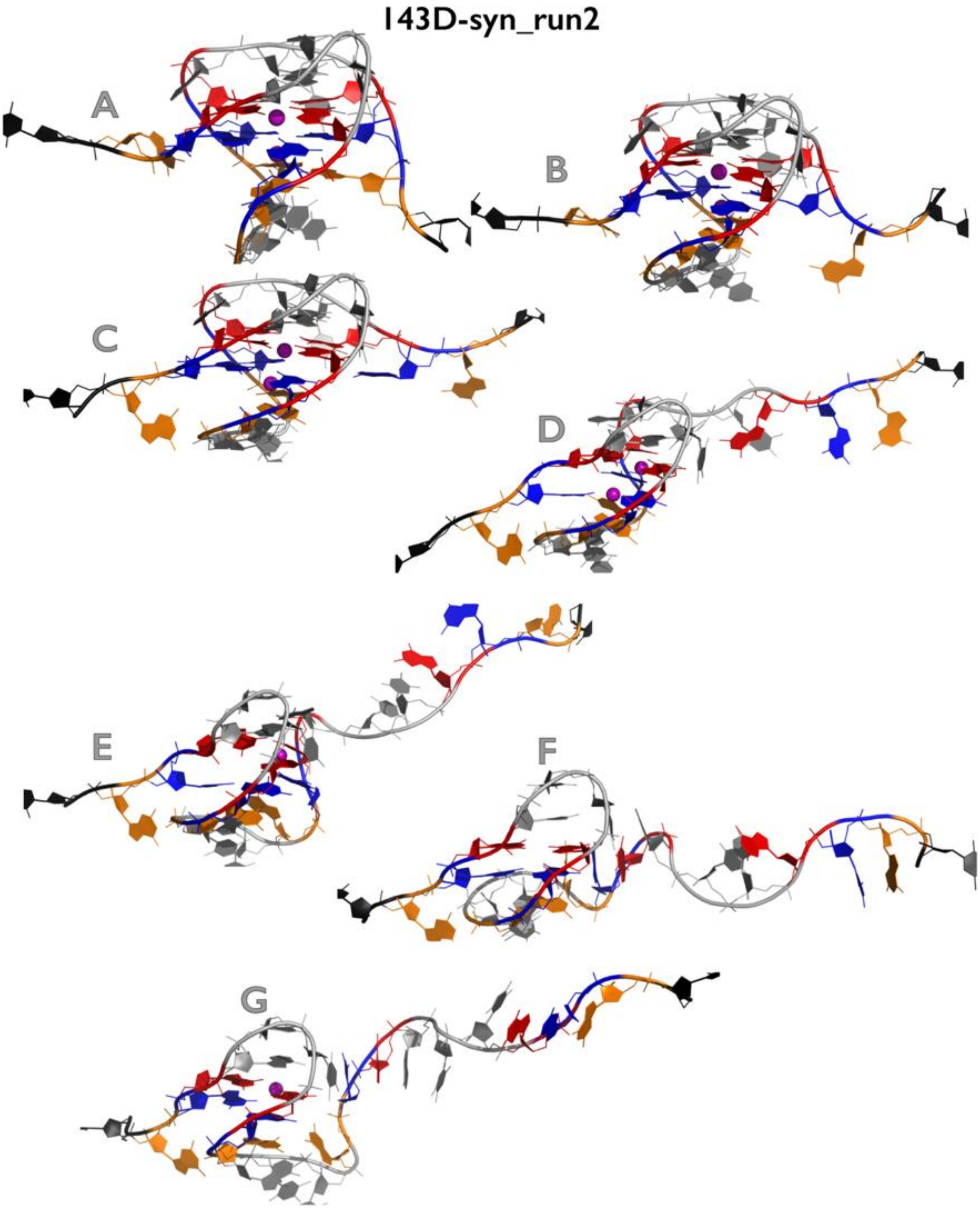

**Figure S21C:** Most important structural events during second independent *very slow zig-zag pulling* simulation of $143D_{syn}$ GQ system. See legend of Figure S1B for more details.

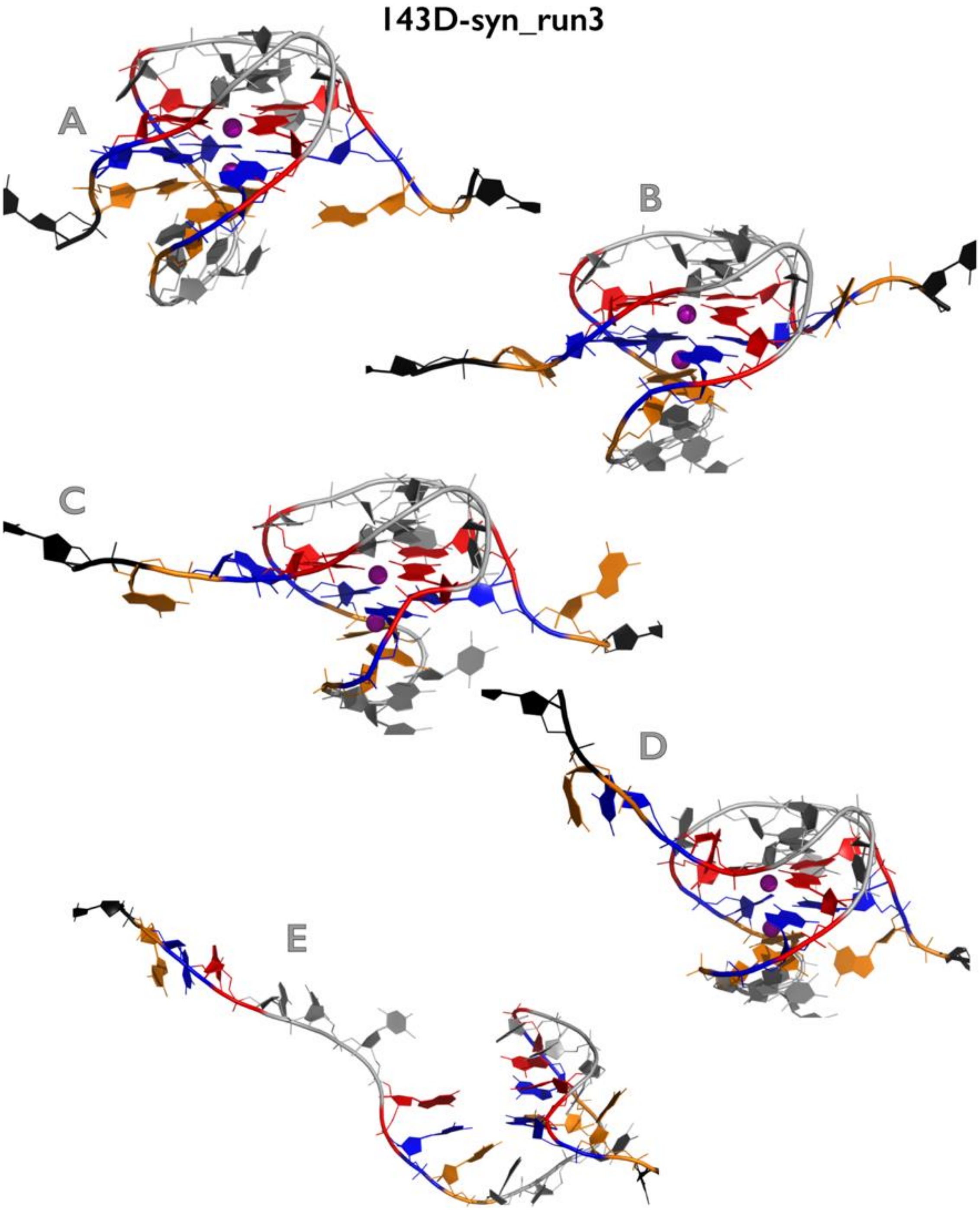

**Figure S21D:** Most important structural events during third independent *very slow zig-zag pulling* simulation of $143D_{syn}$ GQ system. See legend of Figure S1B for more details.

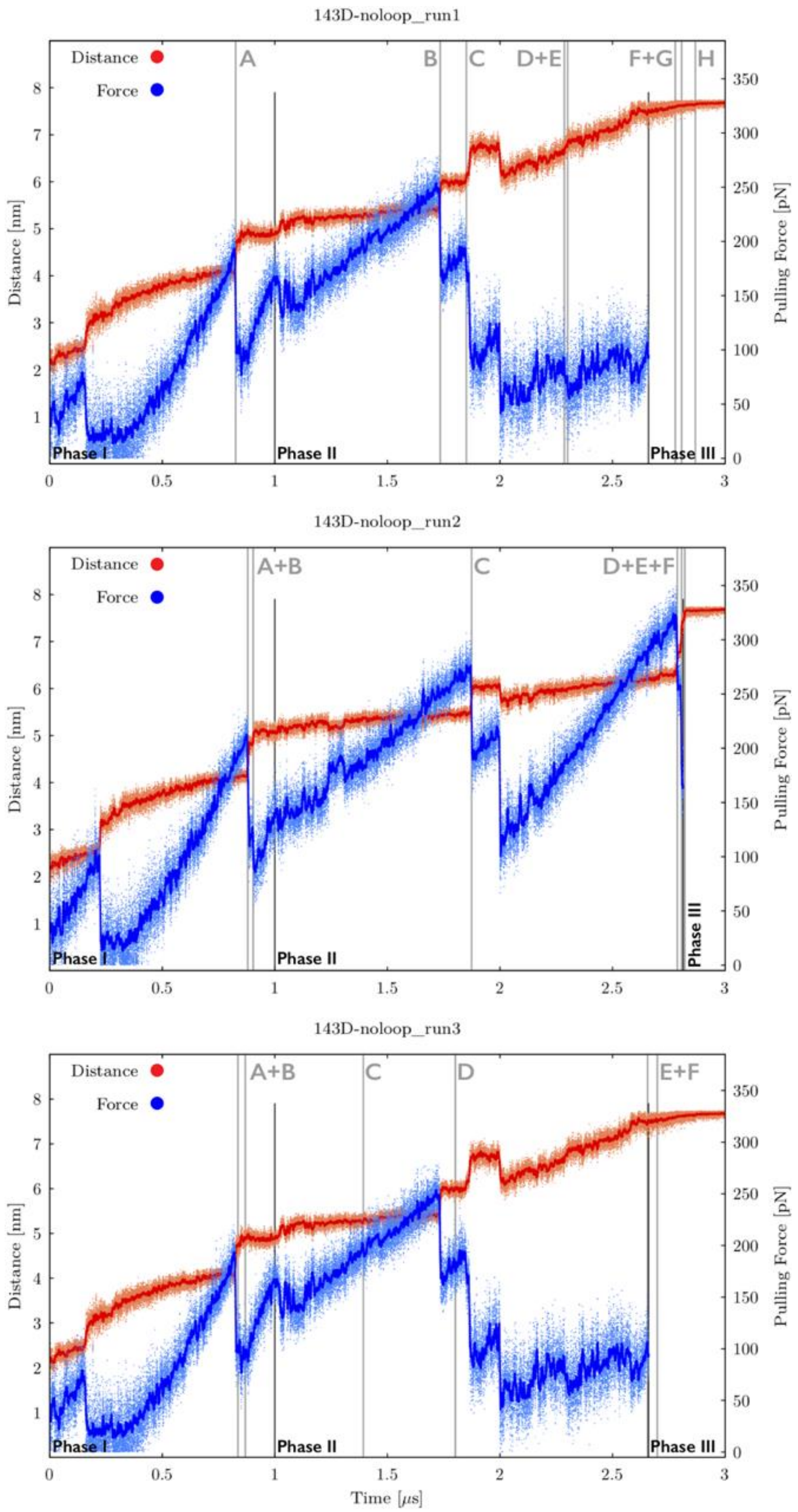

**Figure S22A:** Time evolution of distance between pulling centers and pulling force during three independent *very slow zig-zag pulling* simulations of 143D$_{noloop}$ GQ system (see legend of Figure S17A for more details). See Figures S22B-S22D for inspection of structures corresponding to main structural events.

**143D-noloop_run1 (part1)**

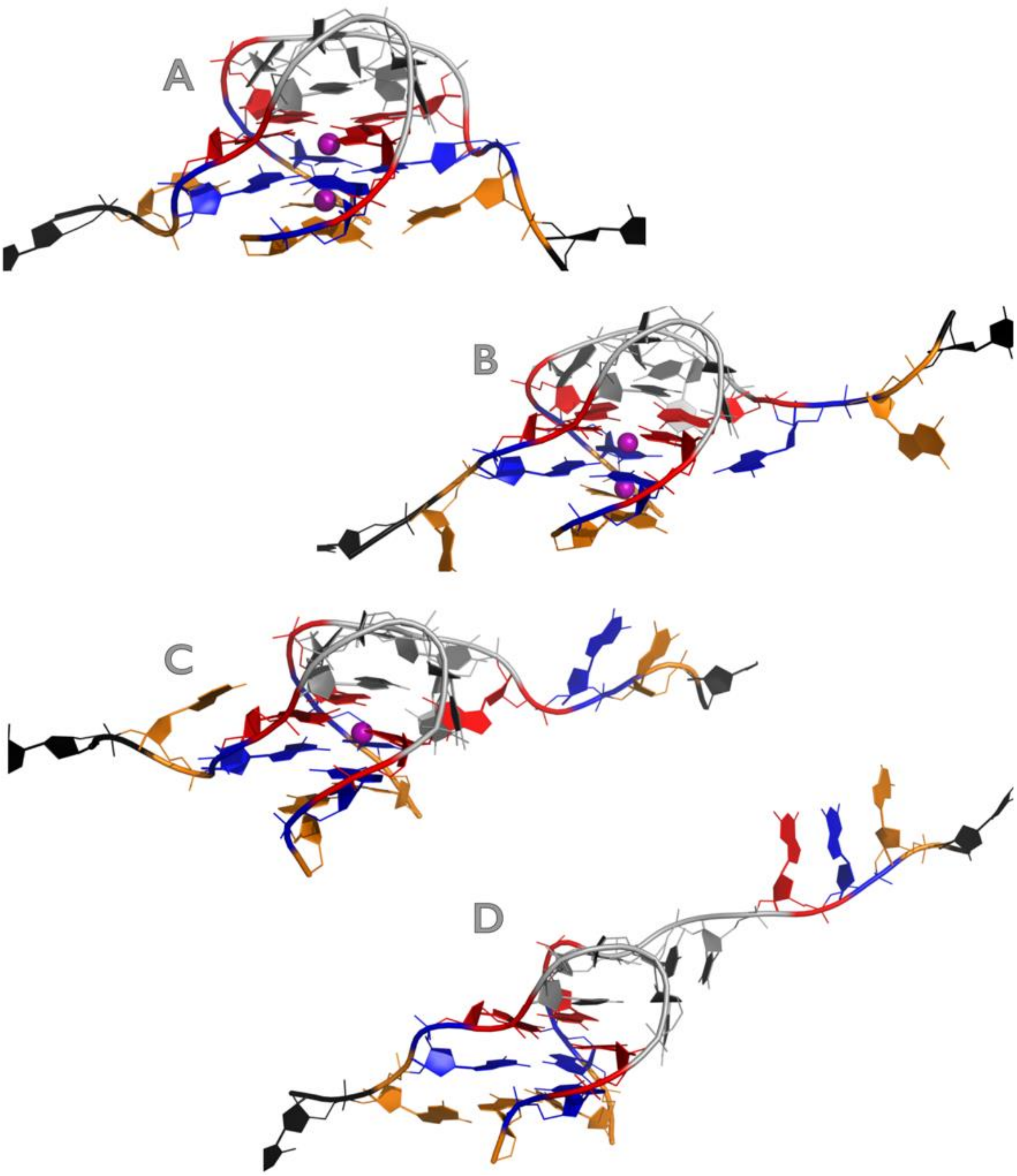

Figure continuing on the next page

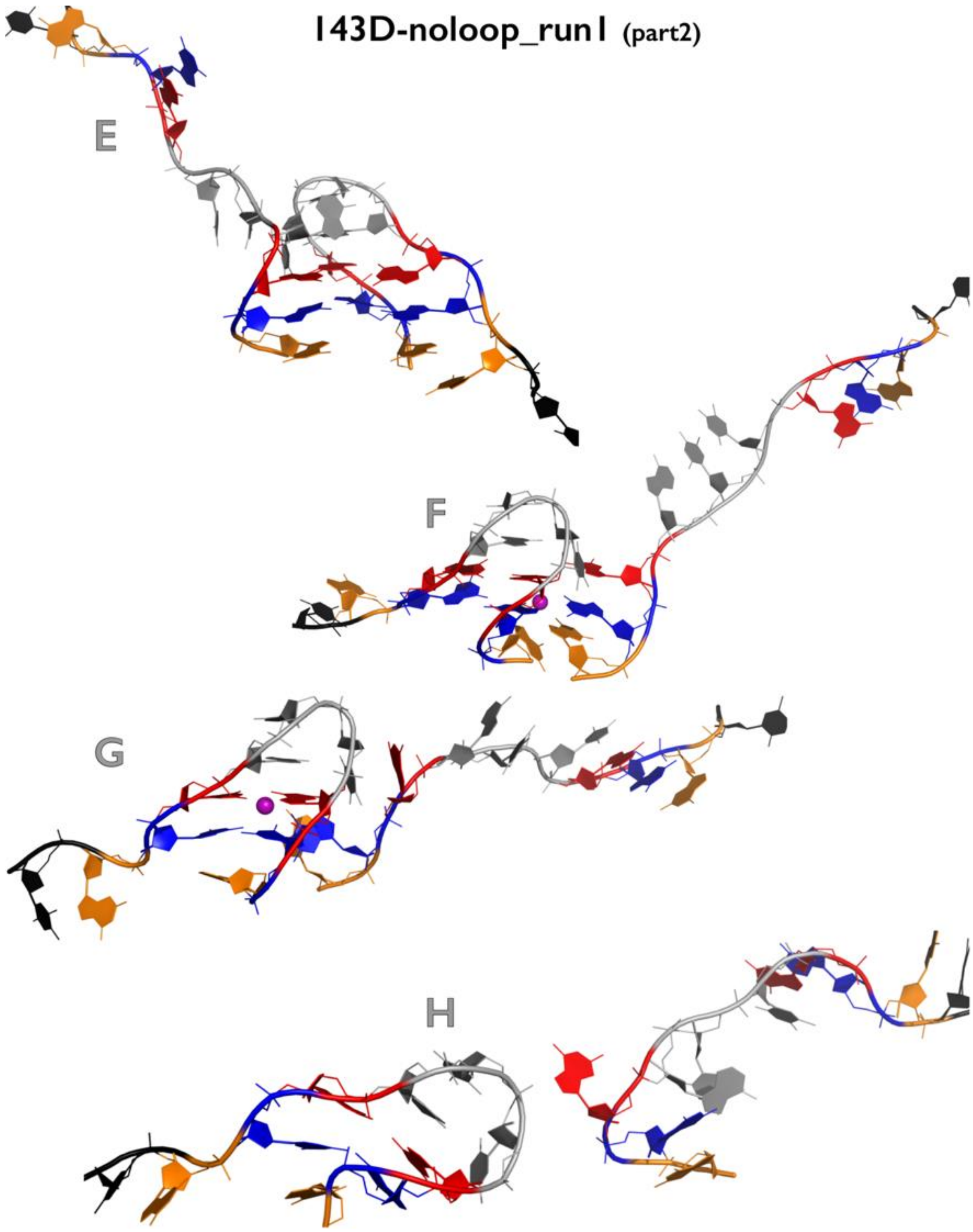

**Figure S22B:** Most important structural events during first independent *very slow zig-zag pulling* simulation of 143D$_{noloop}$ GQ system. See legend of Figure S1B for more details.

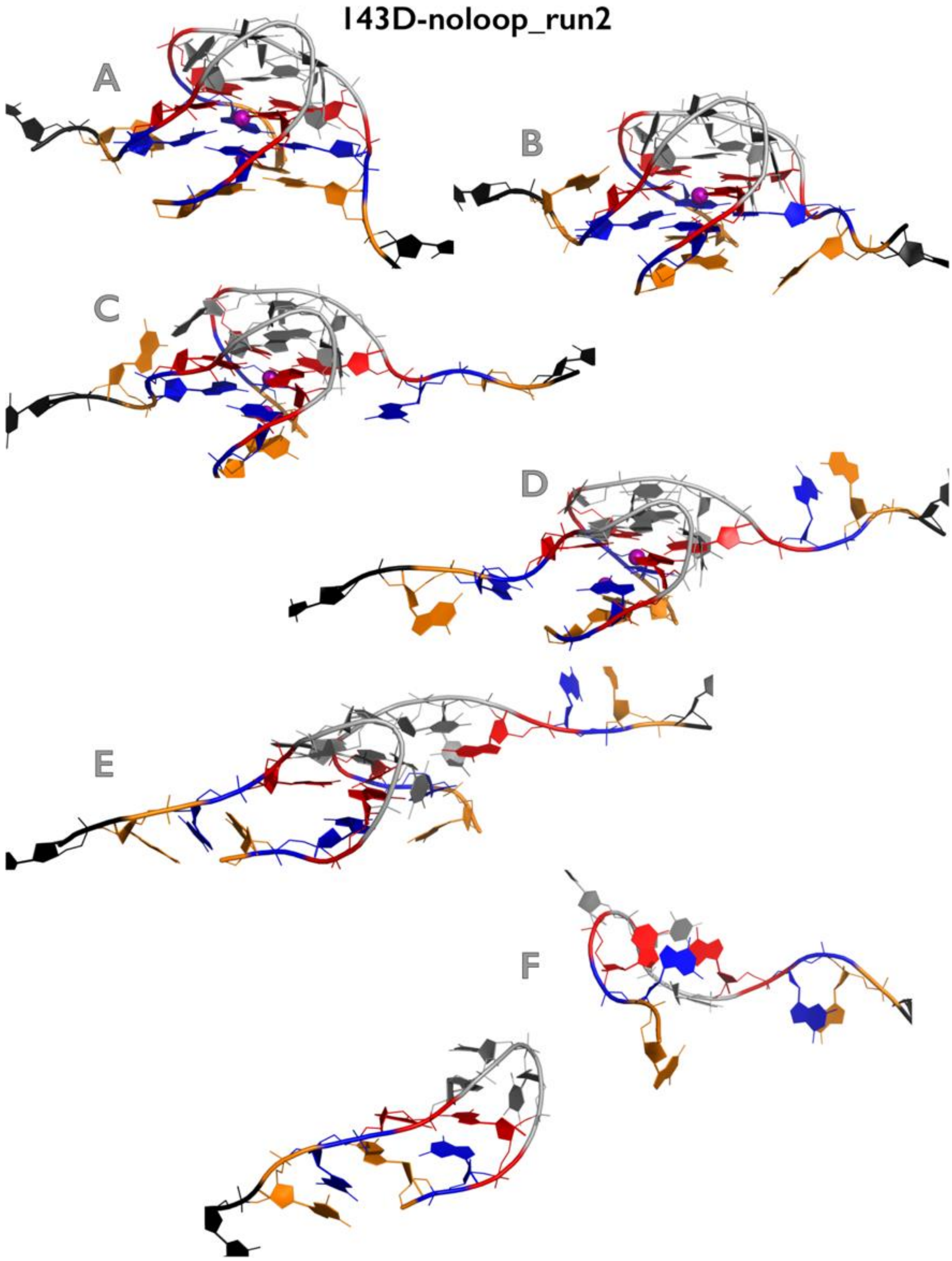

**Figure S22C:** Most important structural events during second independent *very slow zig-zag pulling* simulation of 143D$_{noloop}$ GQ system. See legend of Figure S1B for more details.

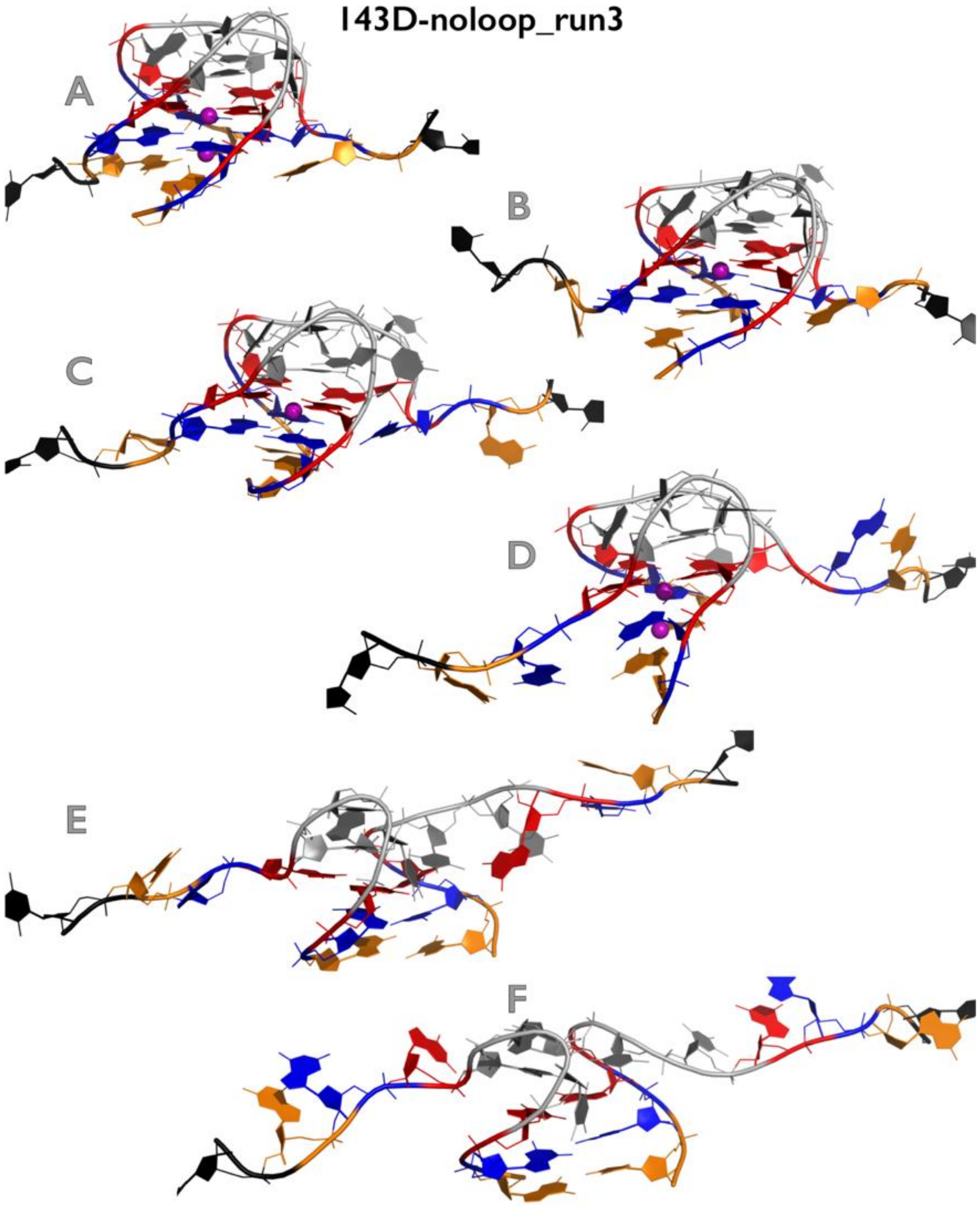

**Figure S22D:** Most important structural events during third independent *very slow zig-zag pulling* simulation of $143D_{noloop}$ GQ system. See legend of Figure S1B for more details.

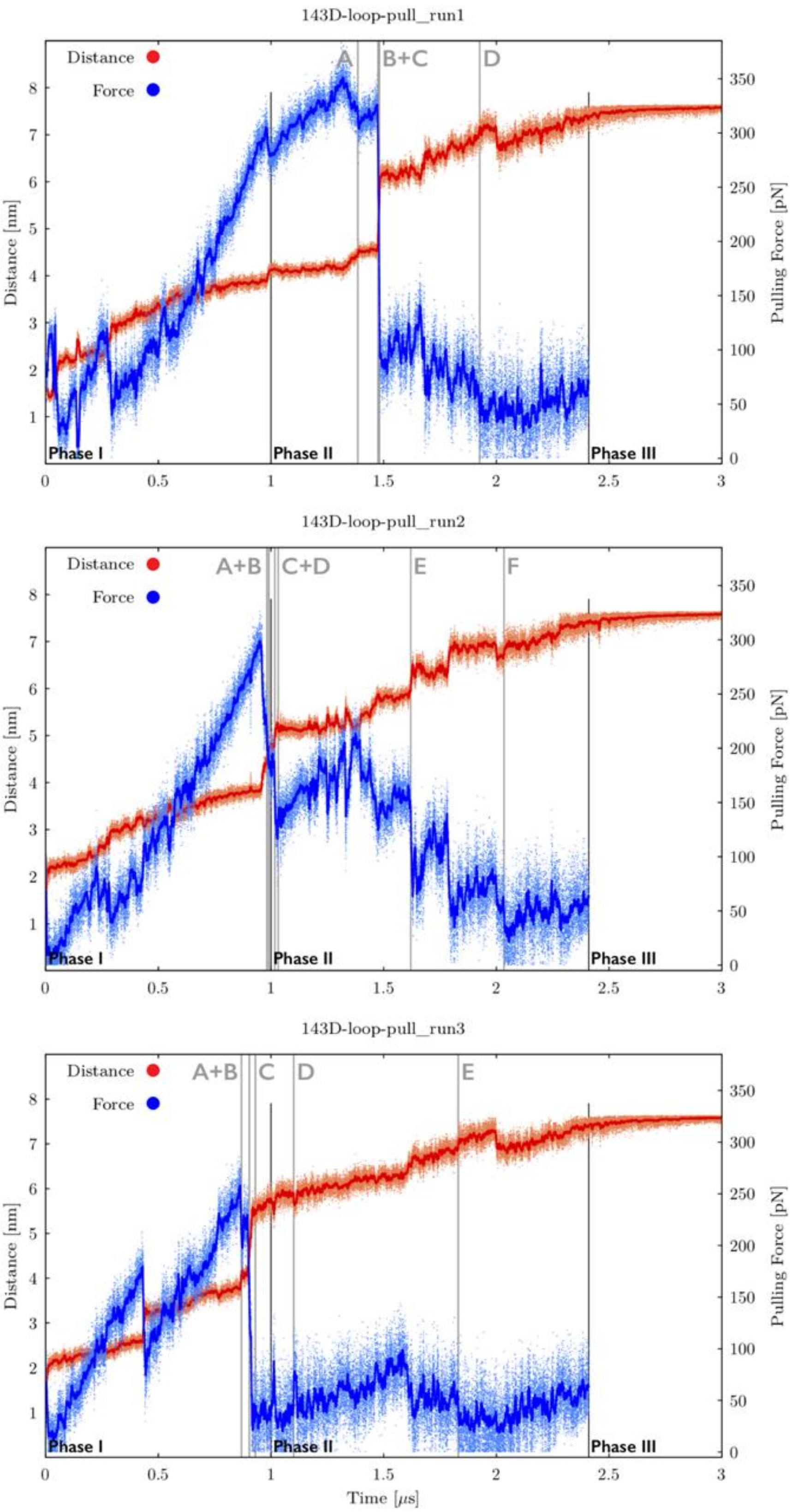

**Figure S23A:** Time evolution of distance between pulling centers and pulling force during three independent *very slow zig-zag pulling* simulations of $143D_{loop\text{-}pull}$ GQ system (see legend of Figure S17A for more details). See Figures S23B-S23D for inspection of structures corresponding to main structural events.

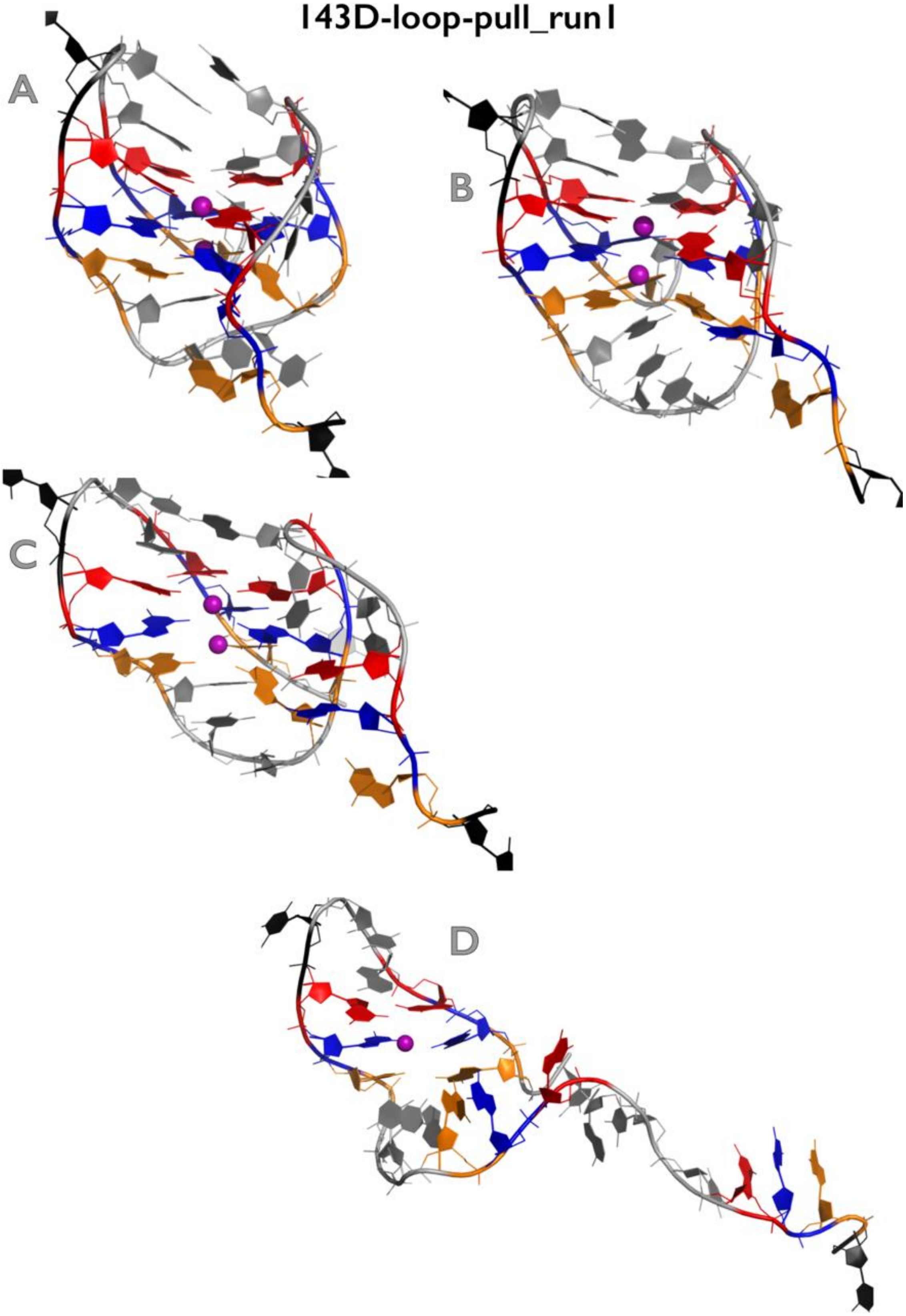

**Figure S23B:** Most important structural events during first independent *very slow zig-zag pulling* simulation of $143D_{loop\text{-}pull}$ GQ system. See legend of Figure S1B for more details.

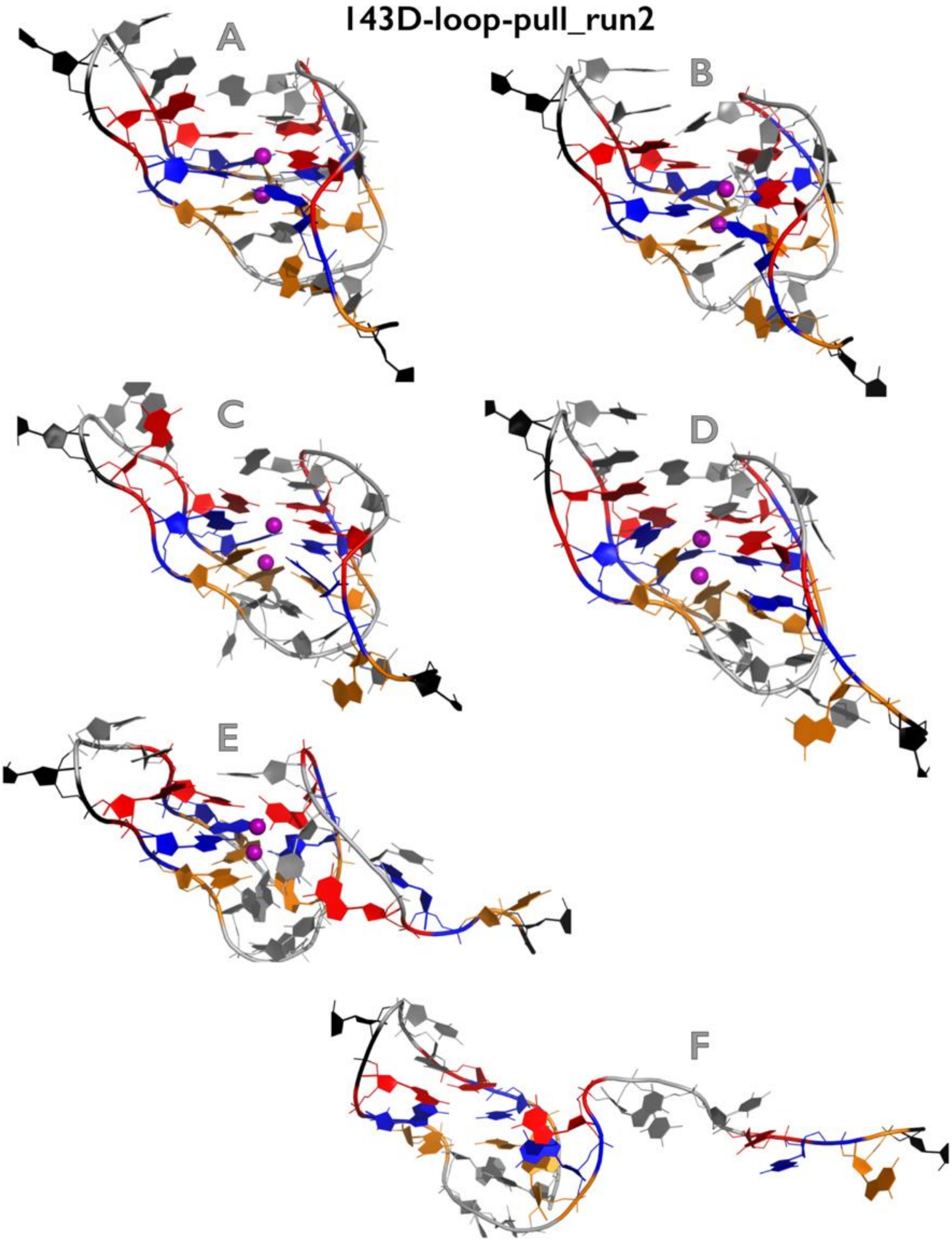

**Figure S23C:** Most important structural events during second independent *very slow zig-zag pulling* simulation of $143D_{loop-pull}$ GQ system. See legend of Figure S1B for more details.

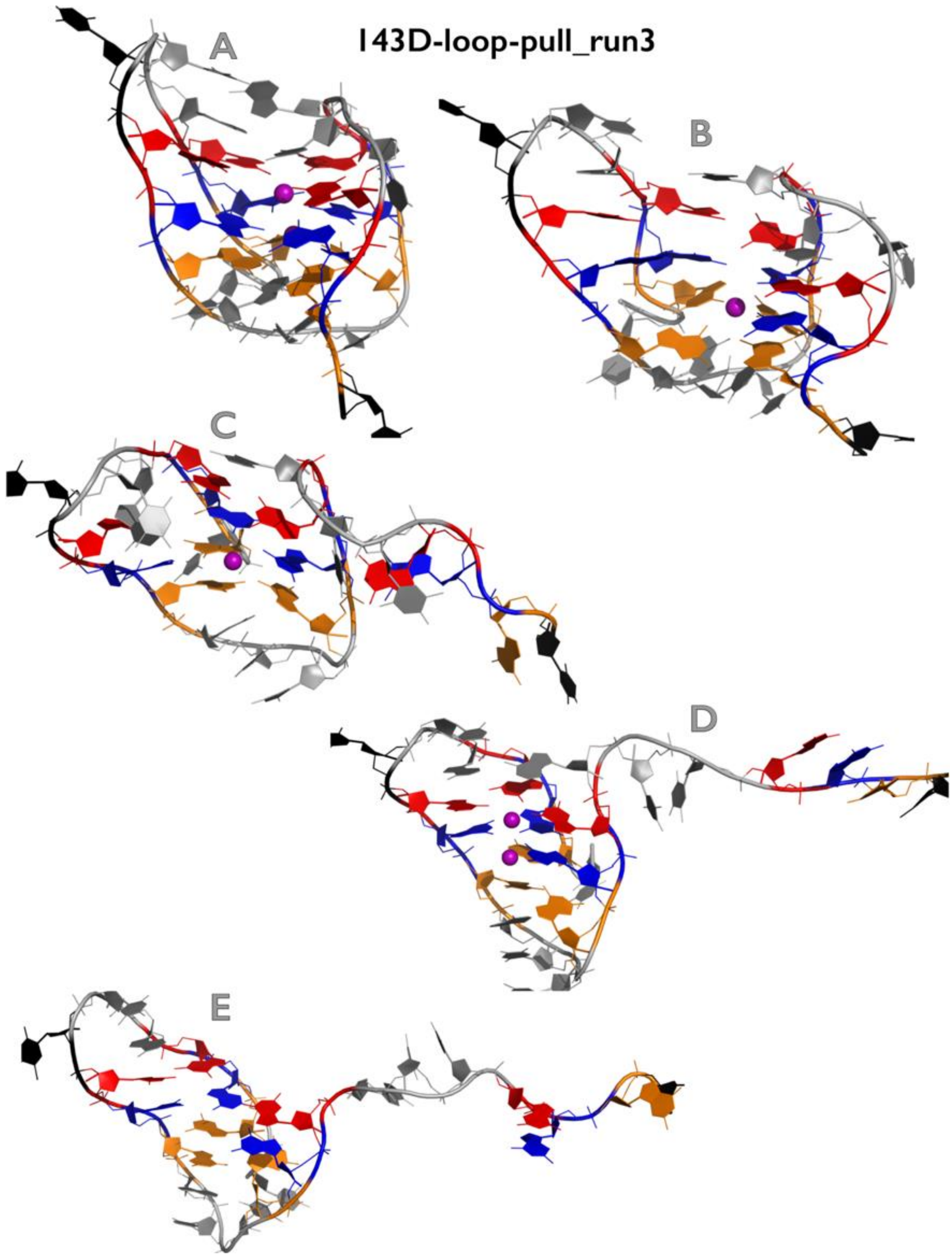

**Figure S23D:** Most important structural events during third independent *very slow zig-zag pulling* simulation of $143D_{loop\text{-}pull}$ GQ system. See legend of Figure S1B for more details.

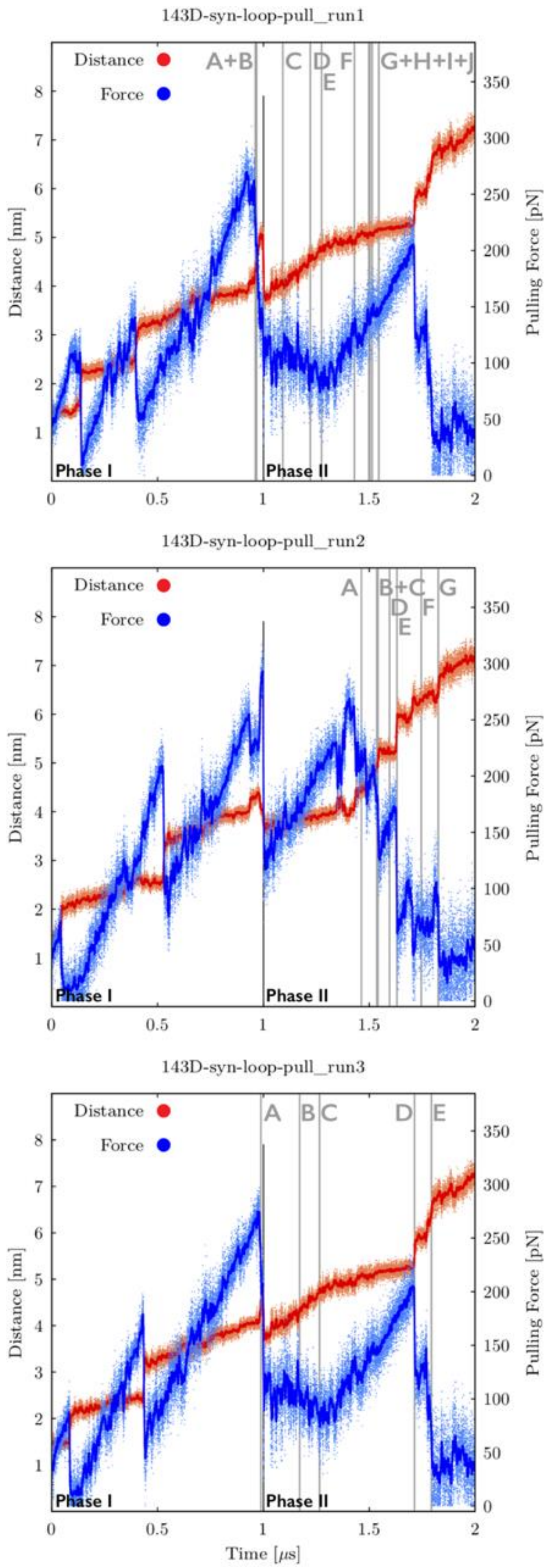

**Figure S24A:** Time evolution of distance between pulling centers and pulling force during three independent *very slow zig-zag pulling* simulations of 143D$_{syn_loop\text{-}pull}$ GQ system (see legend of Figure S17A for more details). See Figures S24B-S24D for inspection of structures corresponding to main structural events.

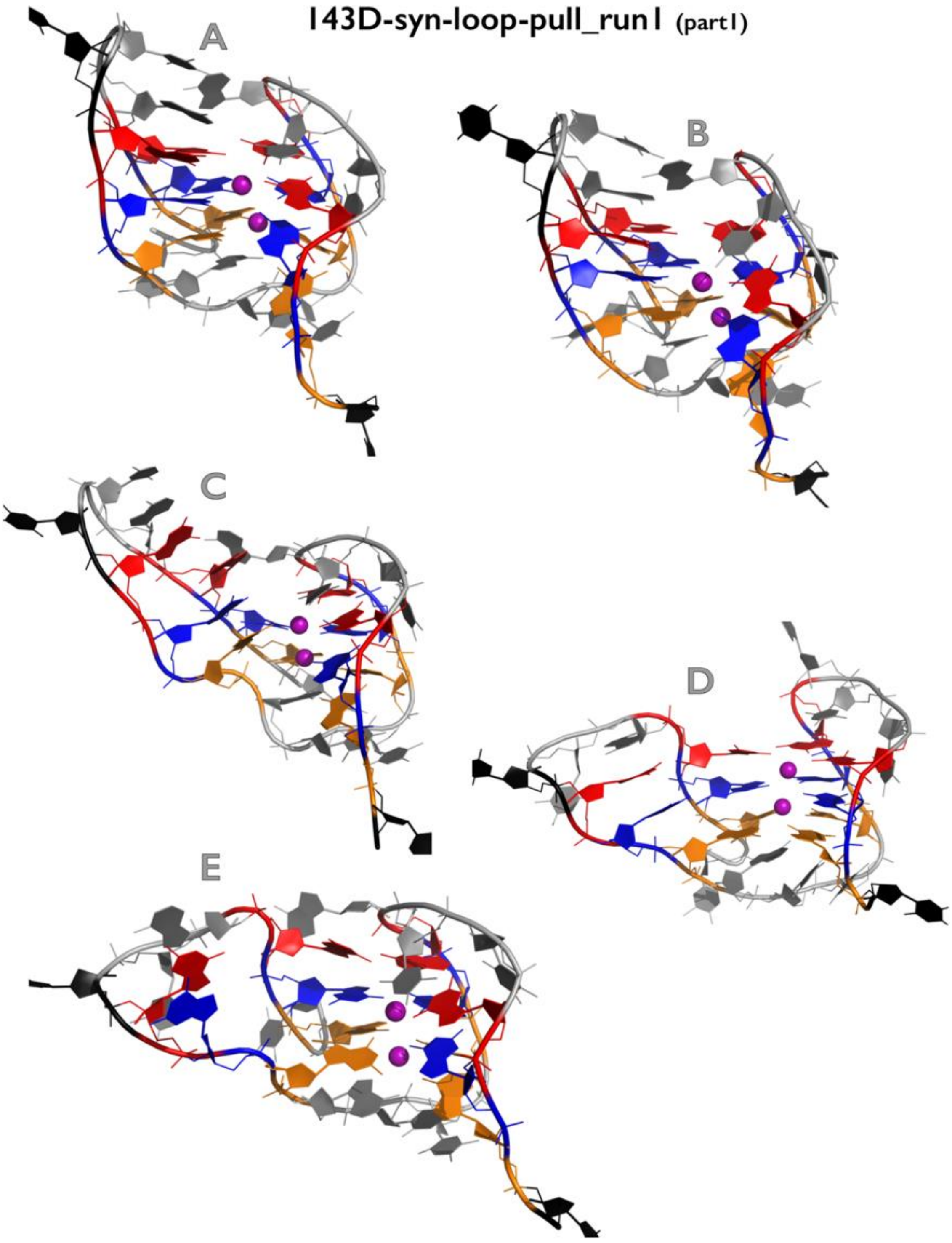

Figure continuing on the next page

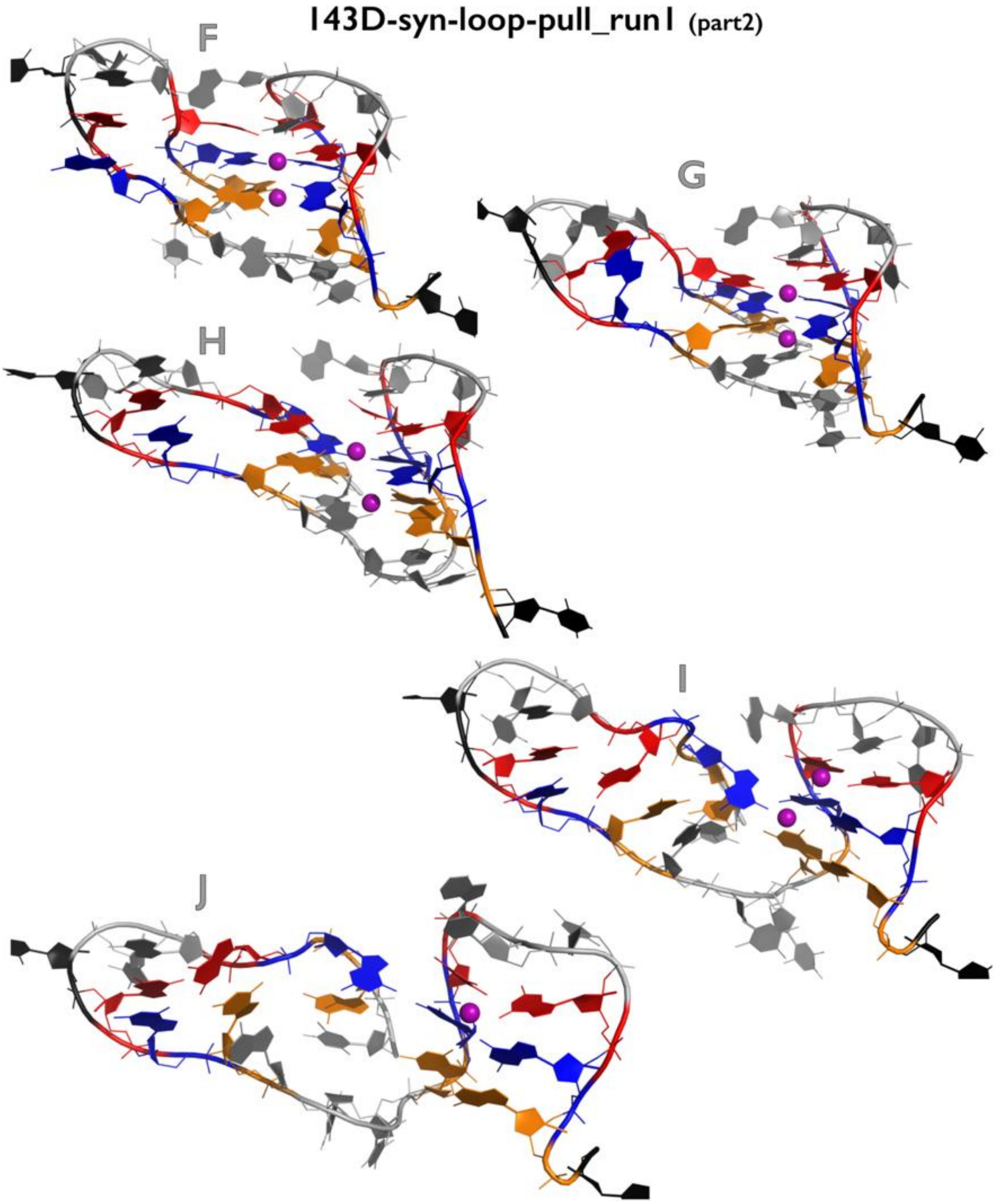

**Figure S24B:** Most important structural events during first independent *very slow zig-zag pulling* simulation of $143D_{syn\_loop\text{-}pull}$ GQ system. See legend of Figure S1B for more details.

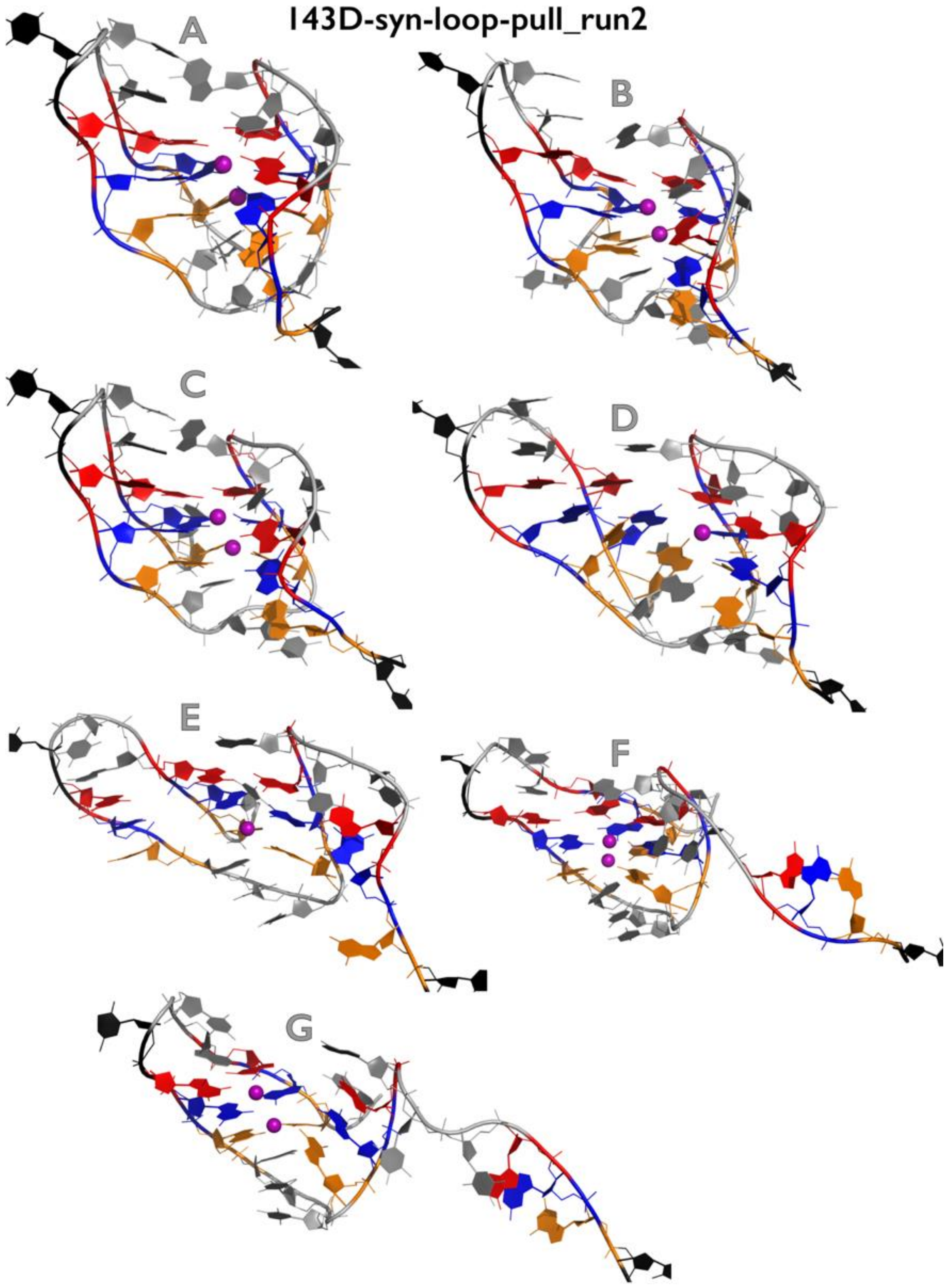

**Figure S24C:** Most important structural events during second independent *very slow zig-zag pulling* simulation of 143D$_{syn_loop\text{-}pull}$ GQ system. See legend of Figure S1B for more details.

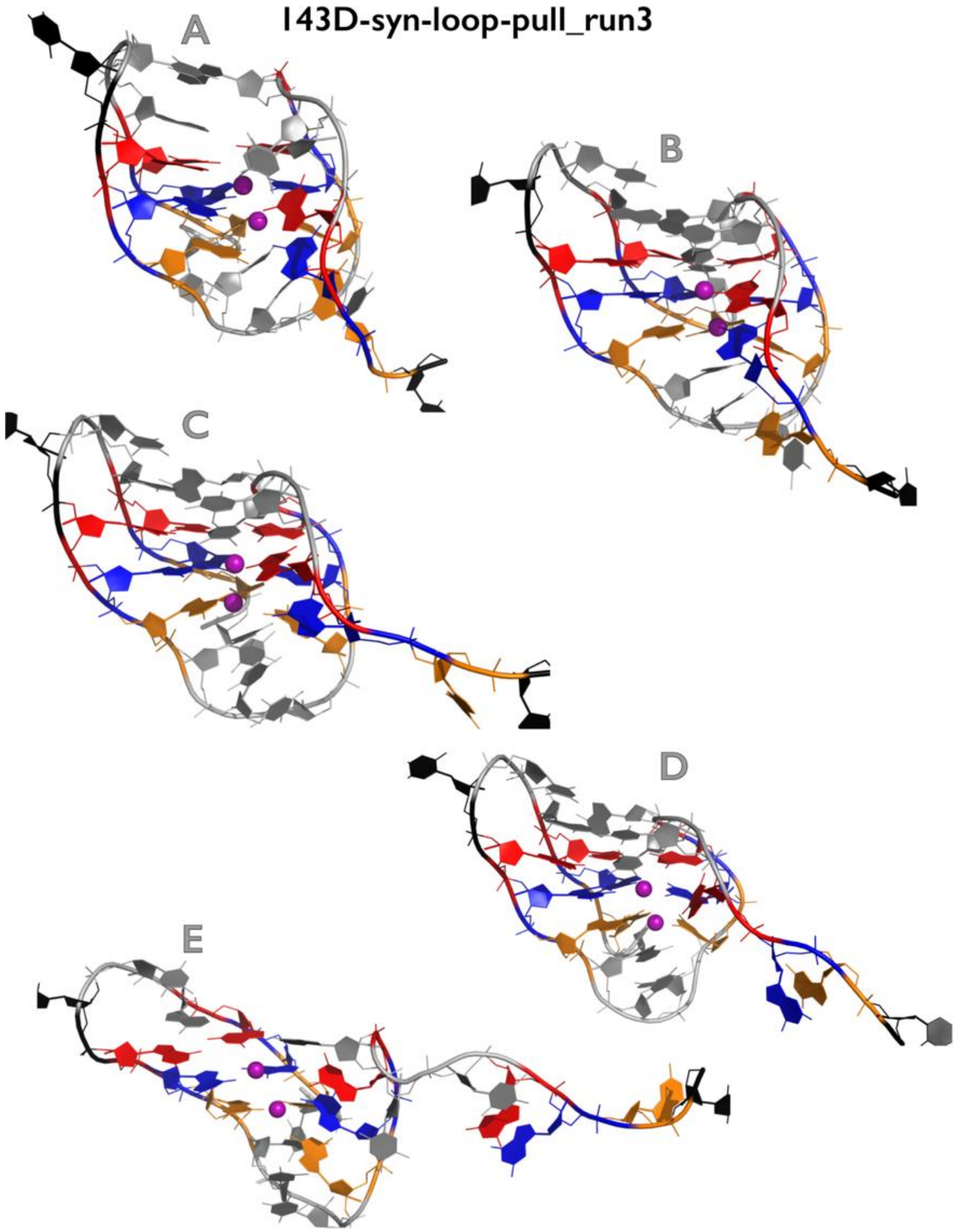

**Figure S24D:** Most important structural events during third independent *very slow zig-zag pulling* simulation of 143D$_{syn_loop\text{-}pull}$ GQ system. See legend of Figure S1B for more details.

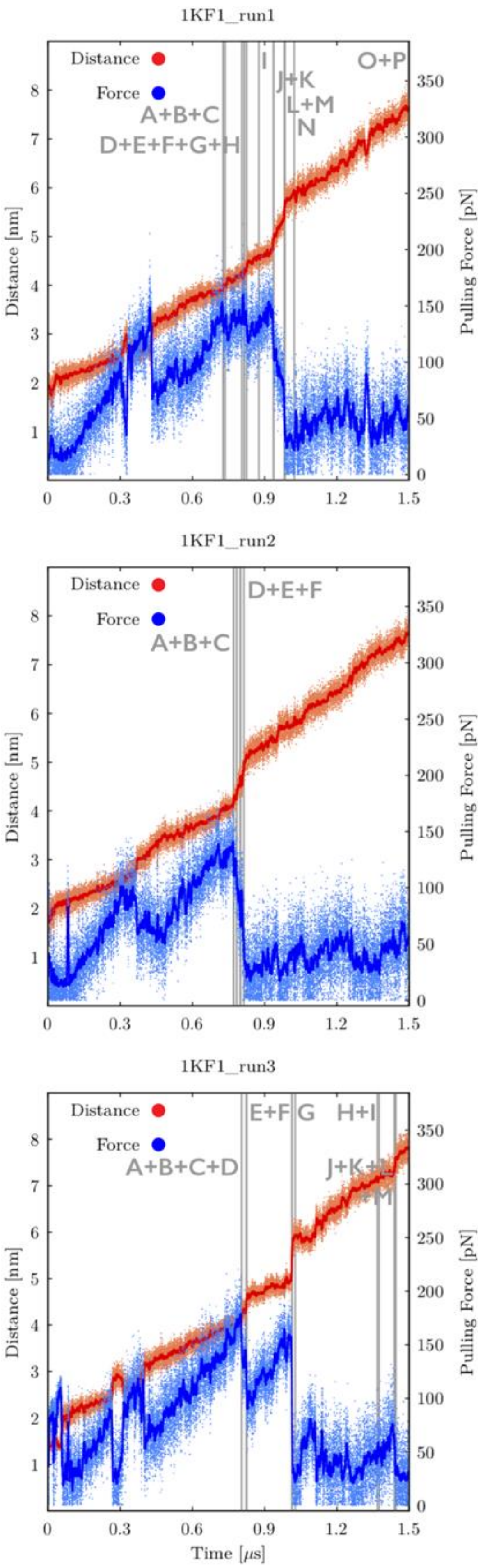

**Figure S25A:** Time evolution of distance between pulling centers and pulling force during three independent *very slow pulling* simulations of 1KF1 GQ system. Snapshots were saved every 50 ps and plots are showing both instantaneous values (orange and light-blue dots for distance and force, respectively) and smoothing, i.e., averaging over 100 consecutive snapshots (red and blue lines for distance and force, respectively). Main structural events are highlighted as grey vertical lines with labels (capital letters). See Figures S25B-S25D for inspection of structures corresponding to main structural events. Note that first major drops of the pulling force before the GQ unfolding event "A" (and notable prolongation of end-to-end distances) are connected with repositioning of terminal T residues.

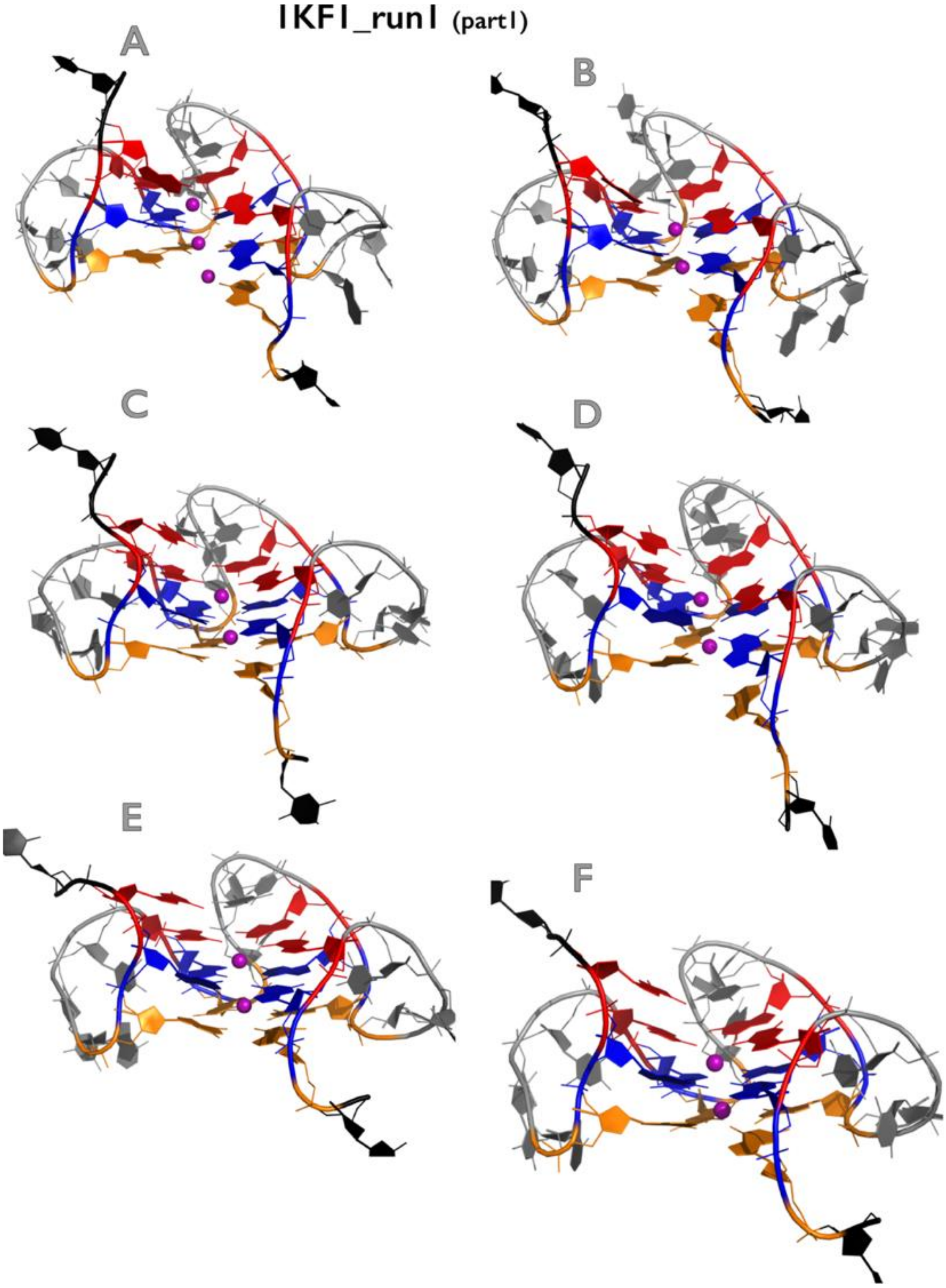

Figure continuing on the next page

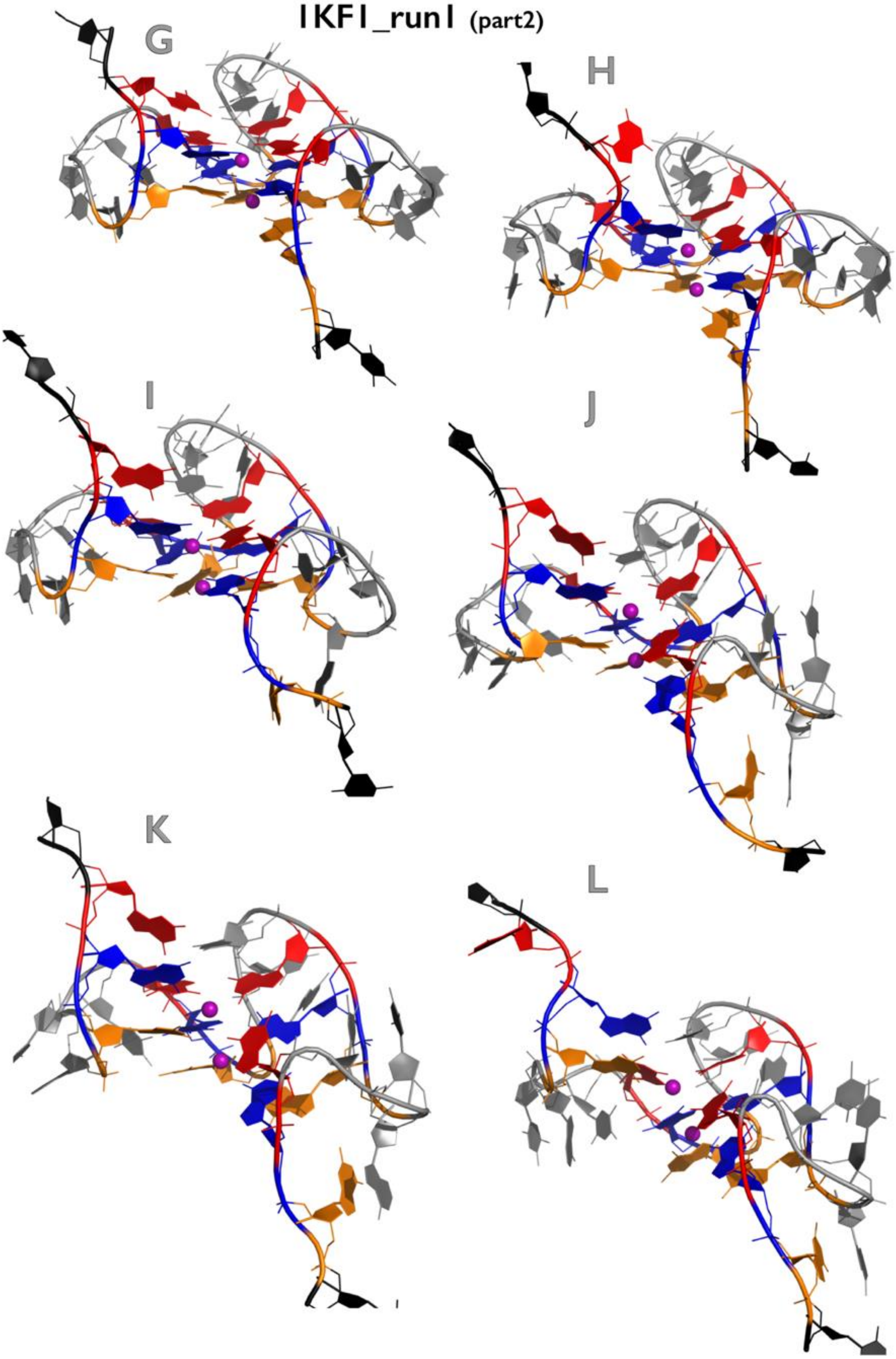

Figure continuing on the next page

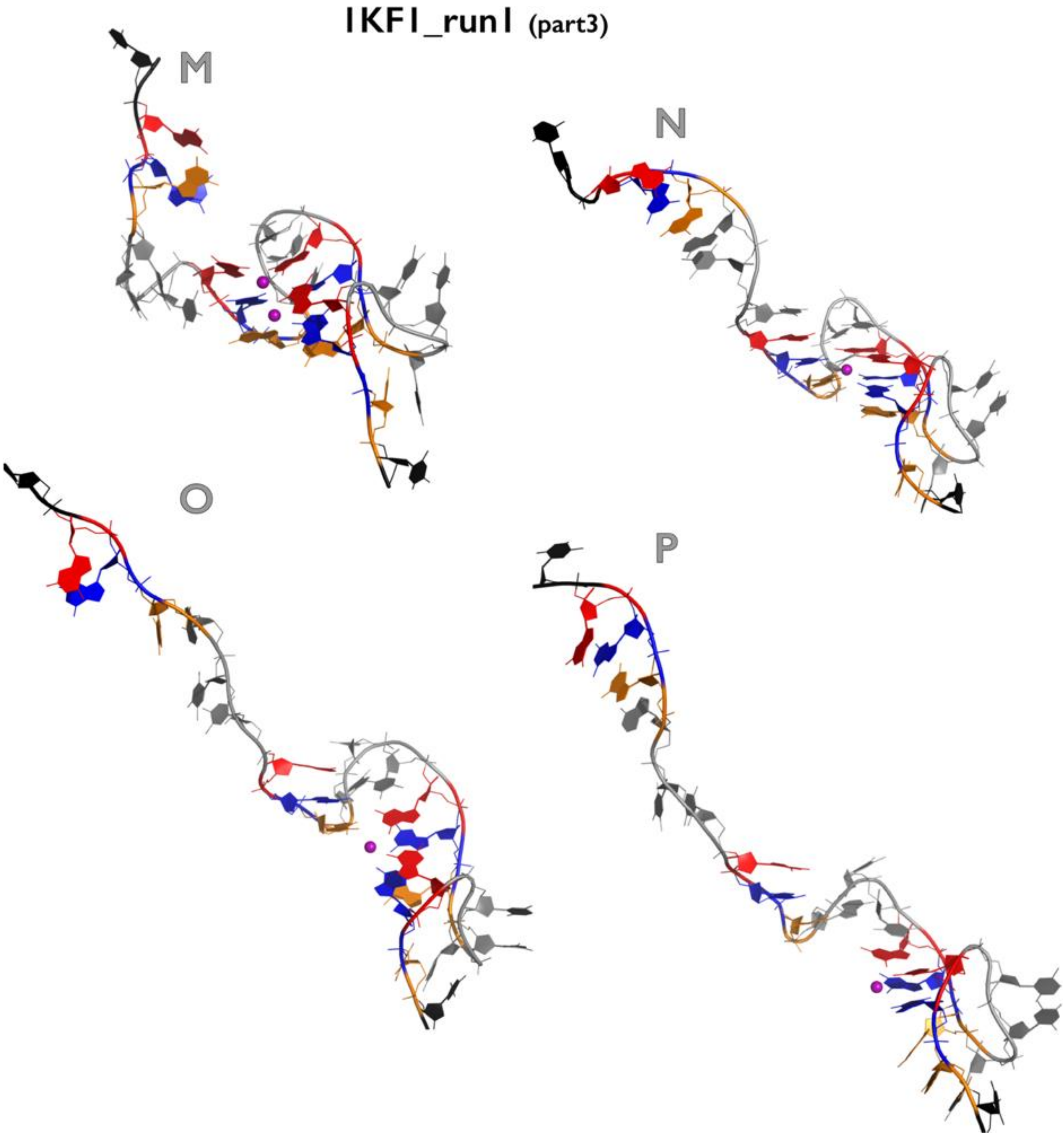

**Figure S25B:** Most important structural events during first independent *very slow pulling* simulation of 1KF1 GQ system. See legend of Figure S1B for more details.

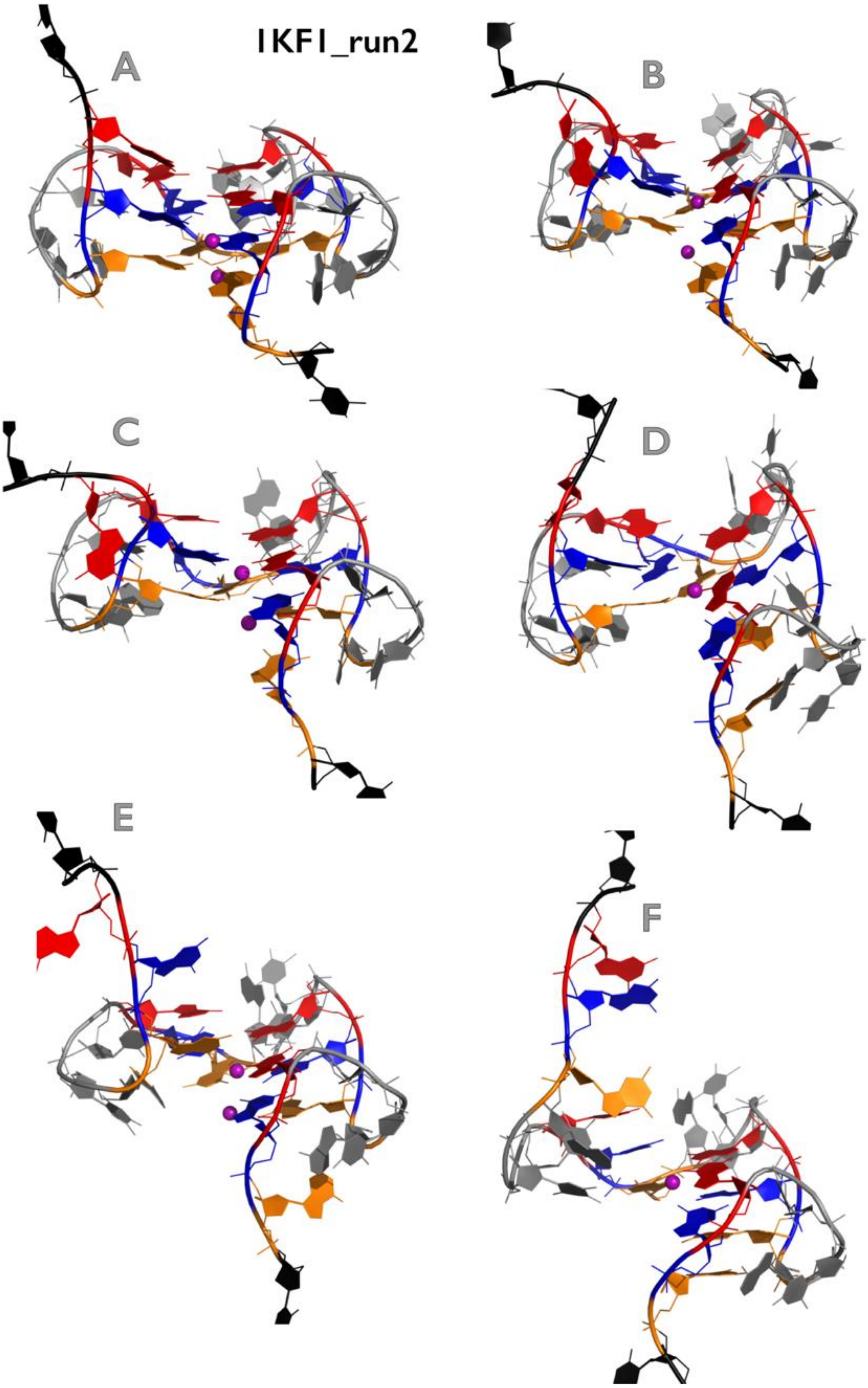

**Figure S25C:** Most important structural events during second independent *very slow pulling* simulation of 1KF1 GQ system. See legend of Figure S1B for more details.

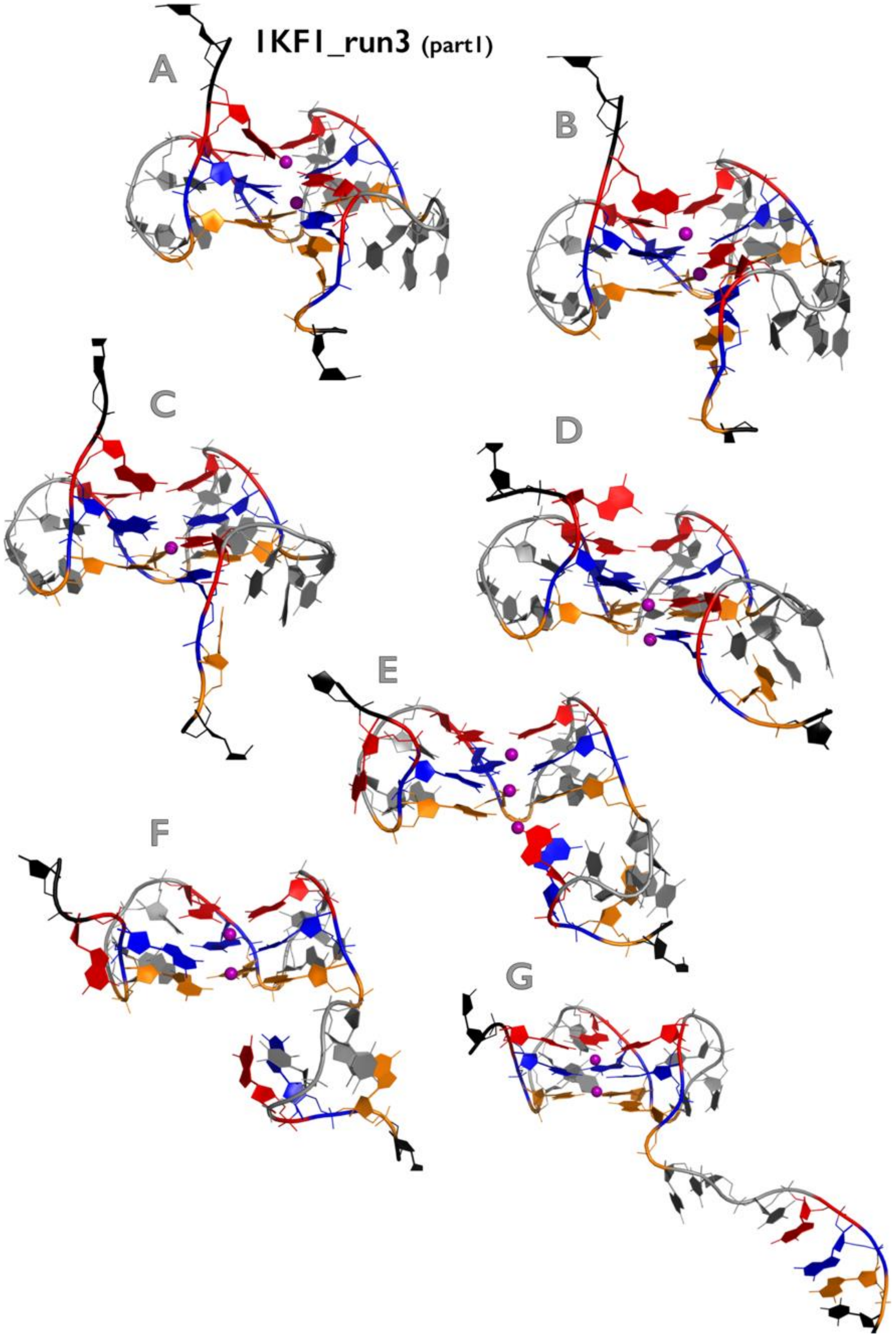

Figure continuing on the next page

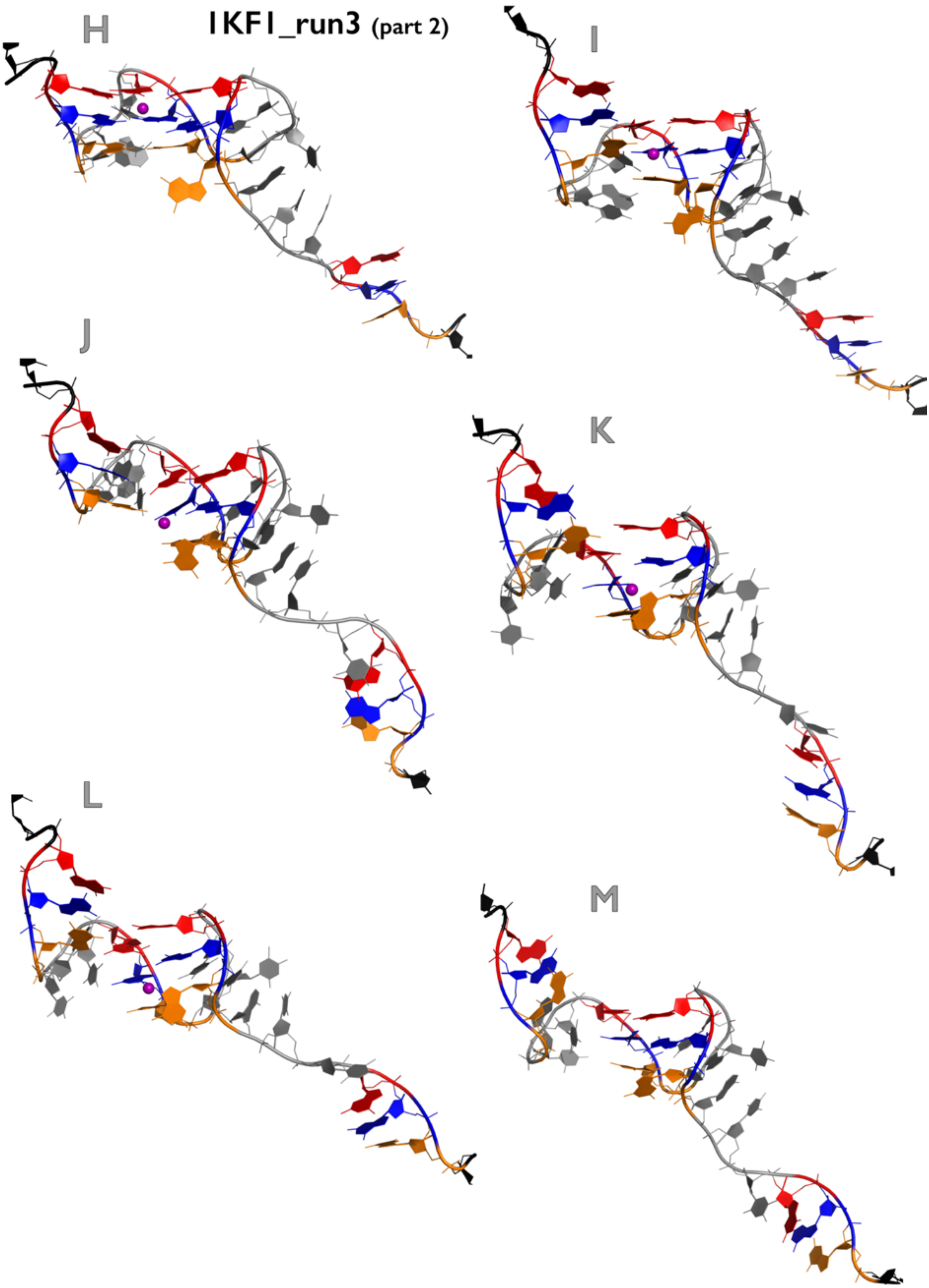

**Figure S25D:** Most important structural events during third independent *very slow pulling* simulation of 1KF1 GQ system. See legend of Figure S1B for more details.

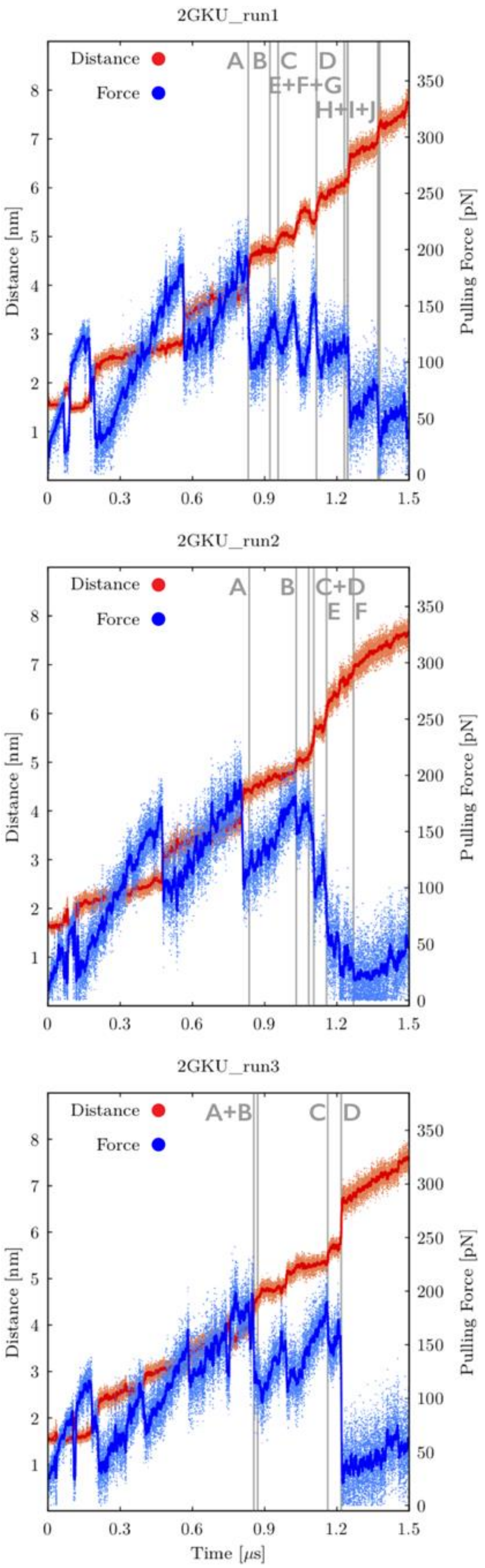

**Figure S26A:** Time evolution of distance between pulling centers and pulling force during three independent *very slow pulling* simulations of 2GKU GQ system (see legend of Figure

S25A for more details). See Figures S26B-26D for inspection of structures corresponding to main structural events.

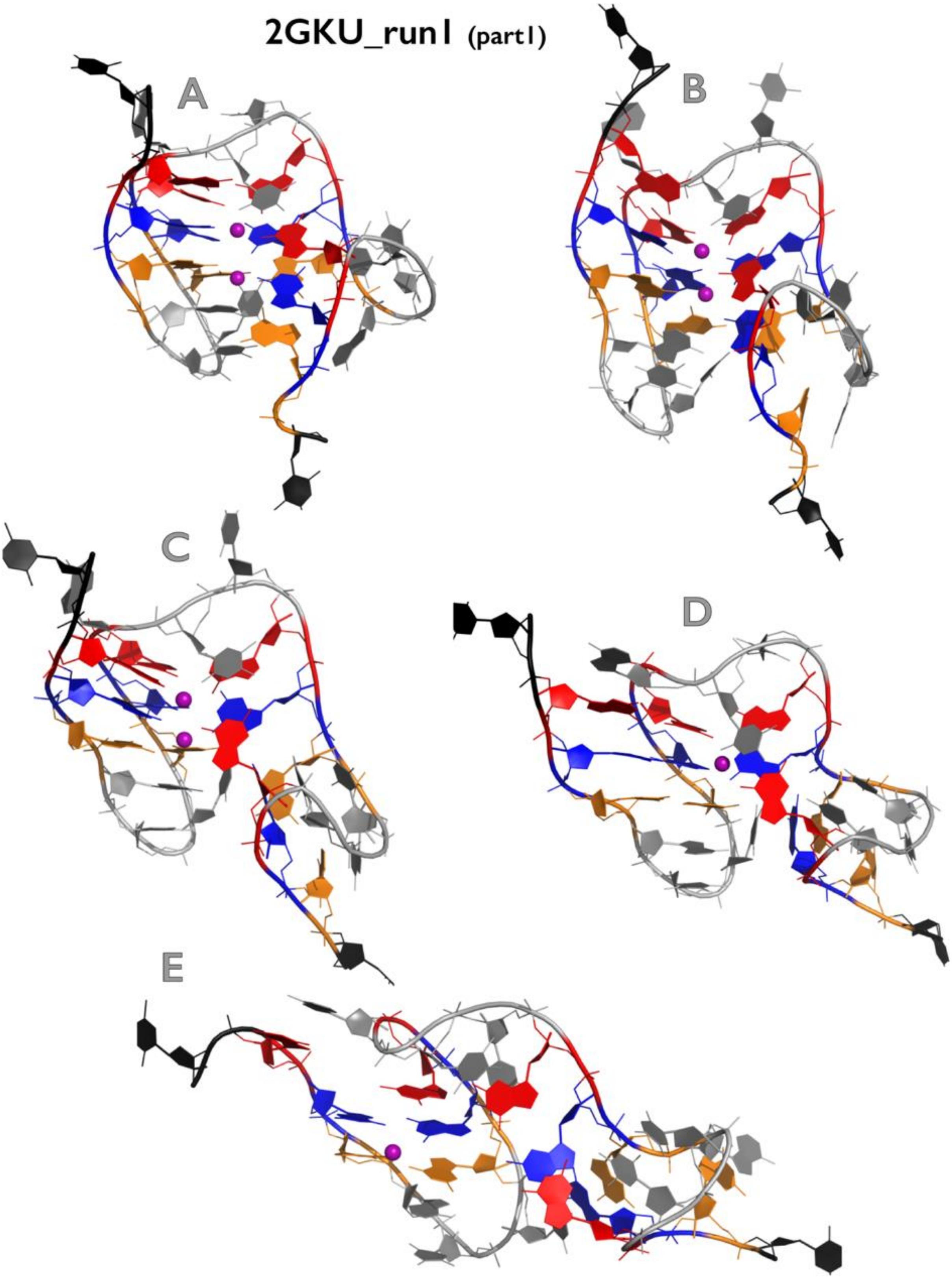

Figure continuing on the next page

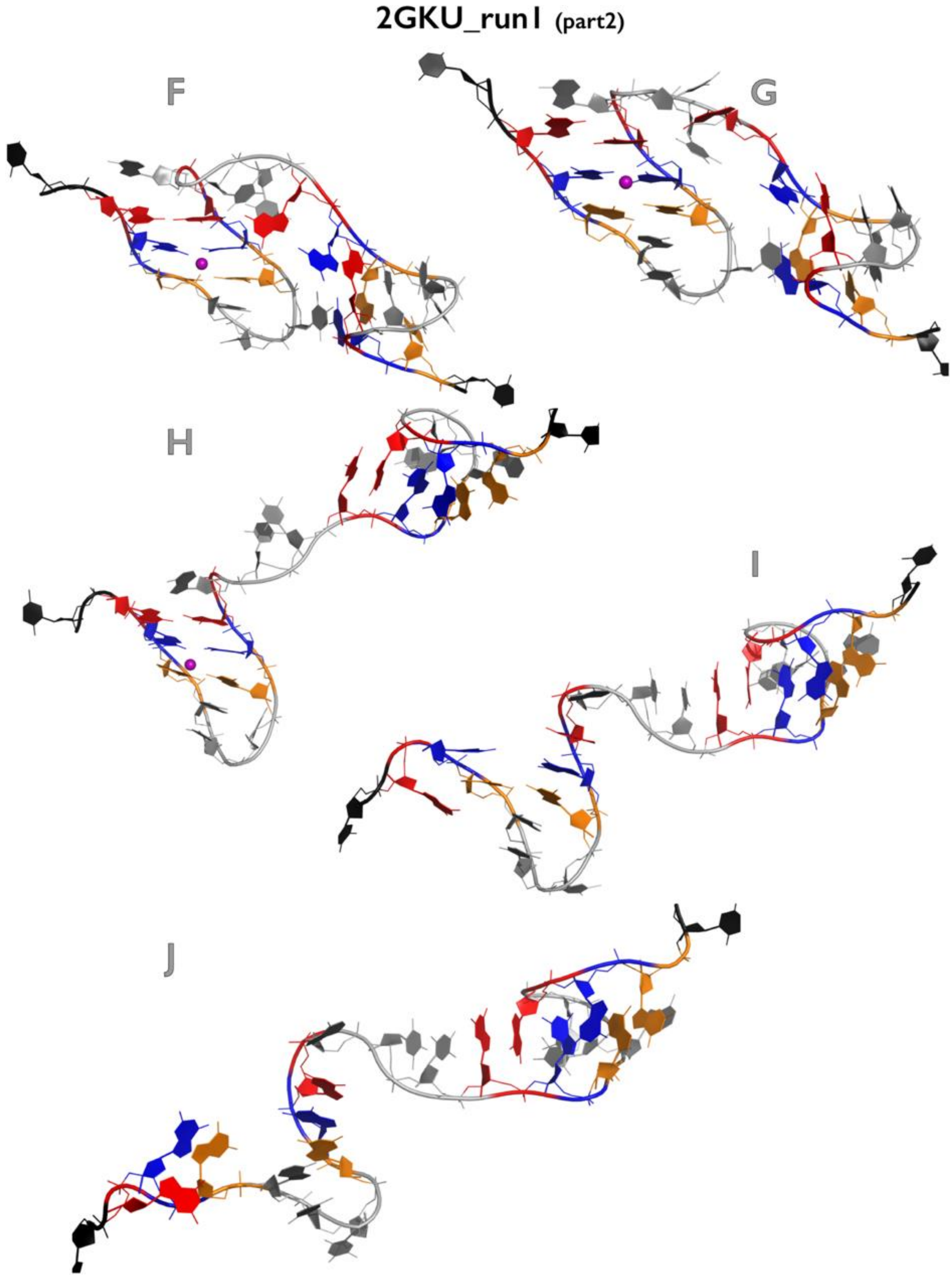

**Figure S26B:** Most important structural events during first independent *very slow pulling* simulation of 2GKU GQ system. See legend of Figure S1B for more details.

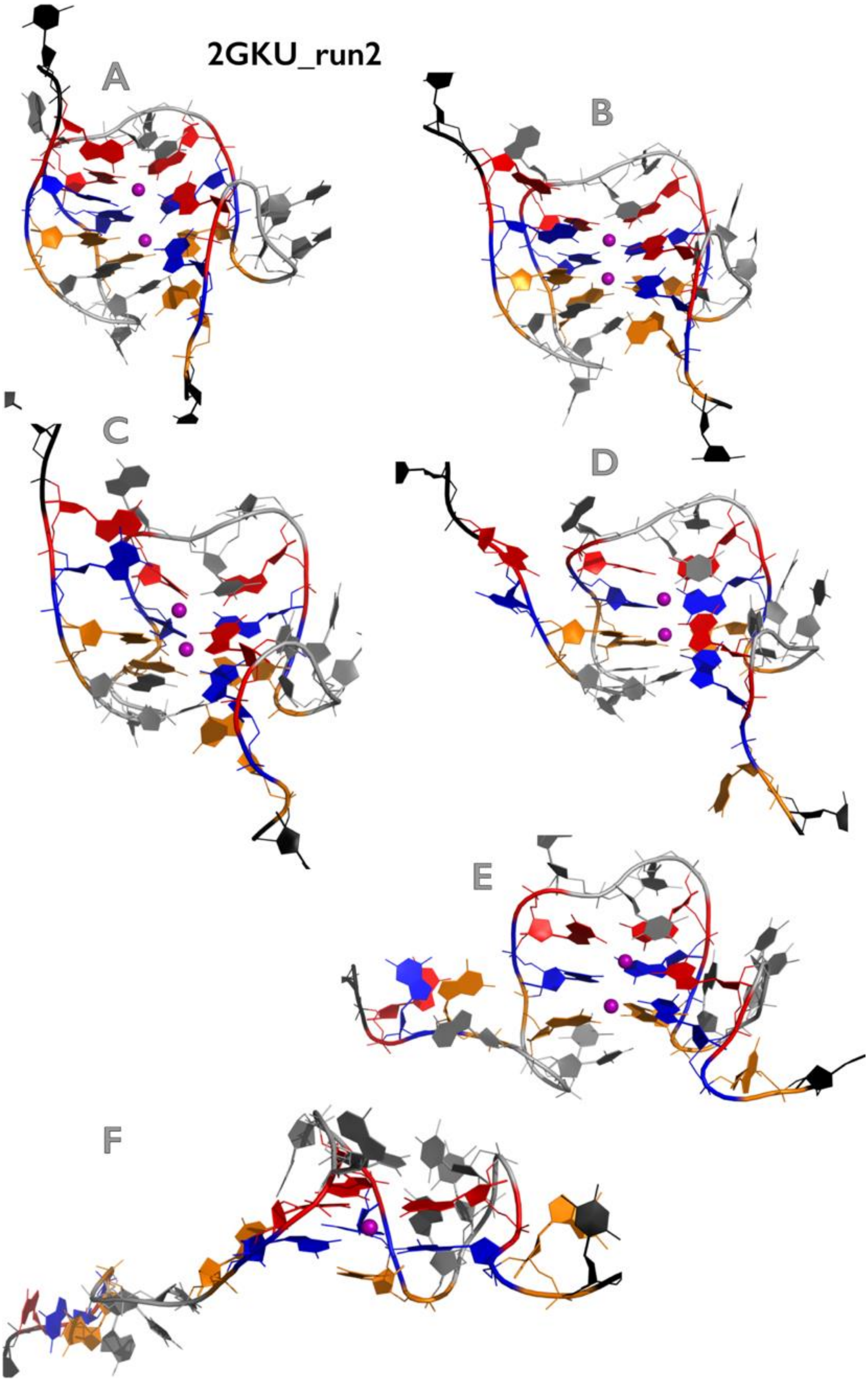

**Figure S26C:** Most important structural events during second independent *very slow pulling* simulation of 2GKU GQ system. See legend of Figure S1B for more details.

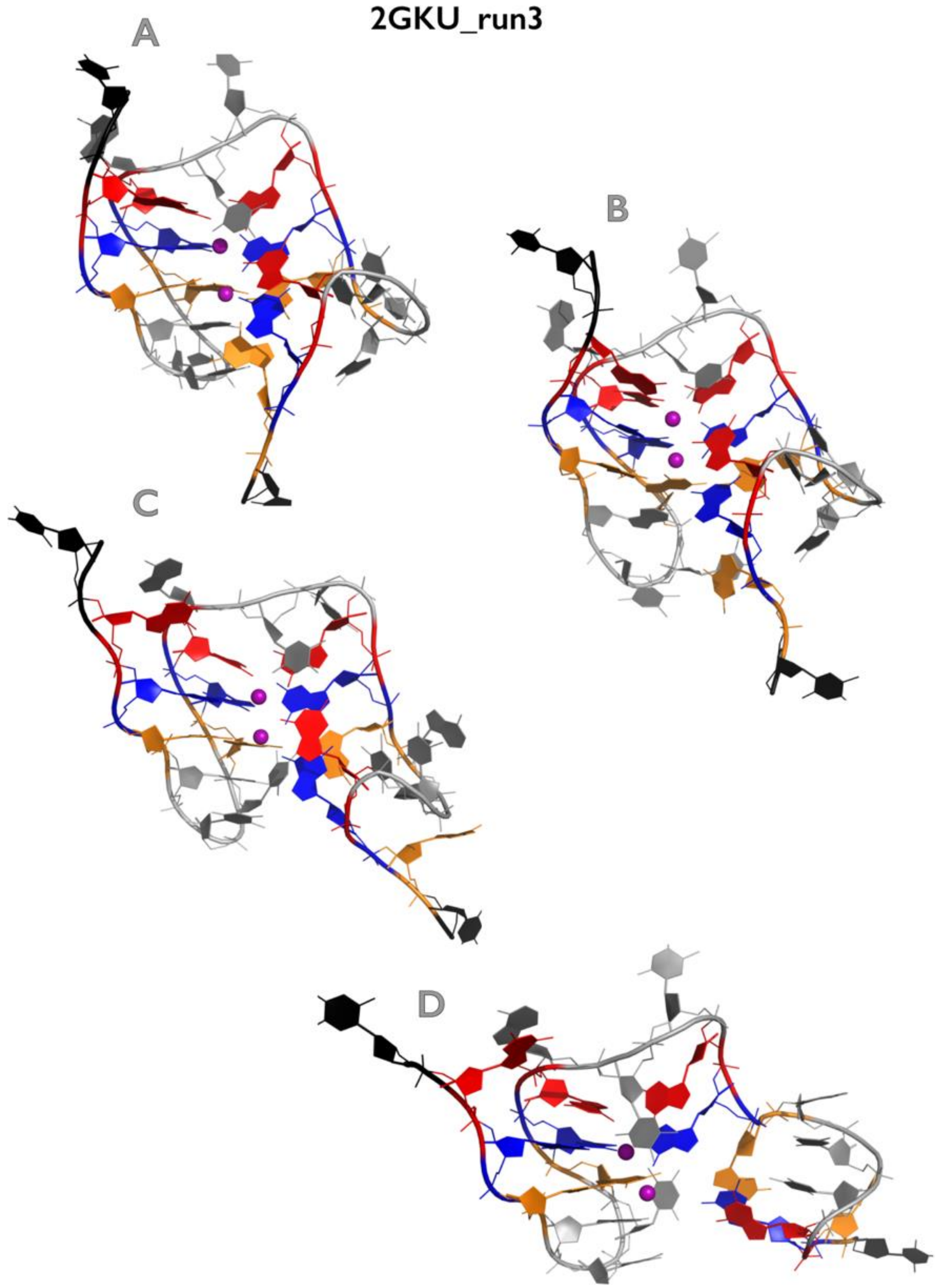

**Figure S26D:** Most important structural events during third independent *very slow pulling* simulation of 2GKU GQ system. See legend of Figure S1B for more details.

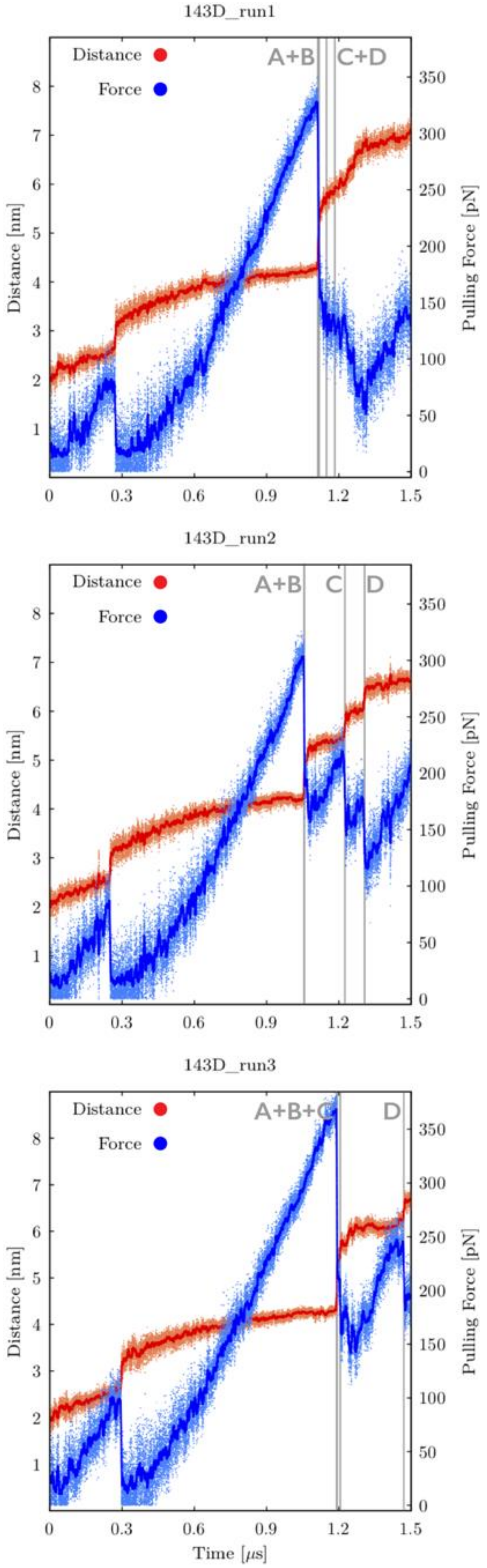

**Figure S27A:** Time evolution of distance between pulling centers and pulling force during three independent *very slow pulling* simulations of 143D GQ system (see legend of Figure

S25A for more details). See Figures S27B-S27D for inspection of structures corresponding to main structural events.

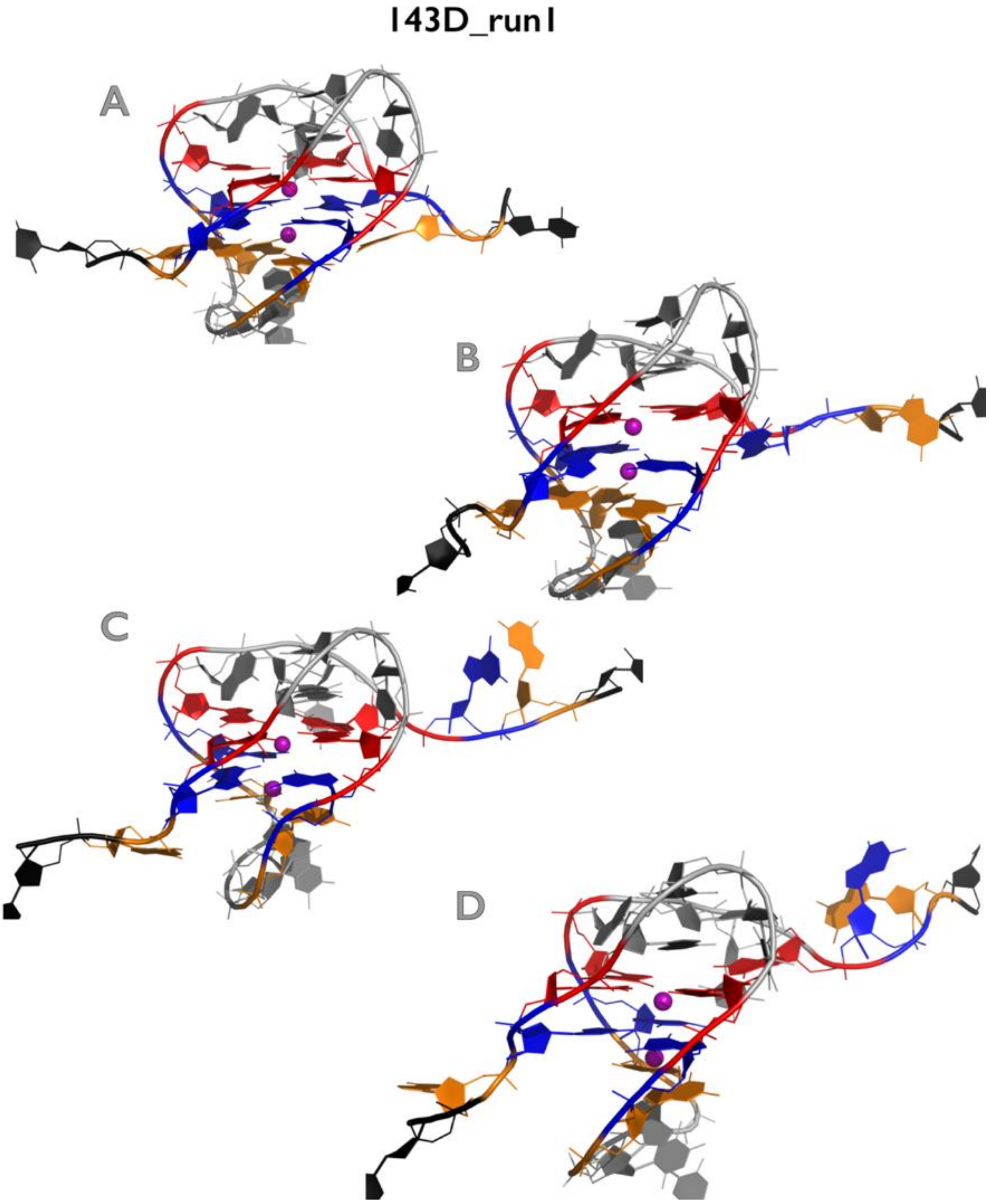

**Figure S27B:** Most important structural events during first independent *very slow pulling* simulation of 143D GQ system. See legend of Figure S1B for more details.

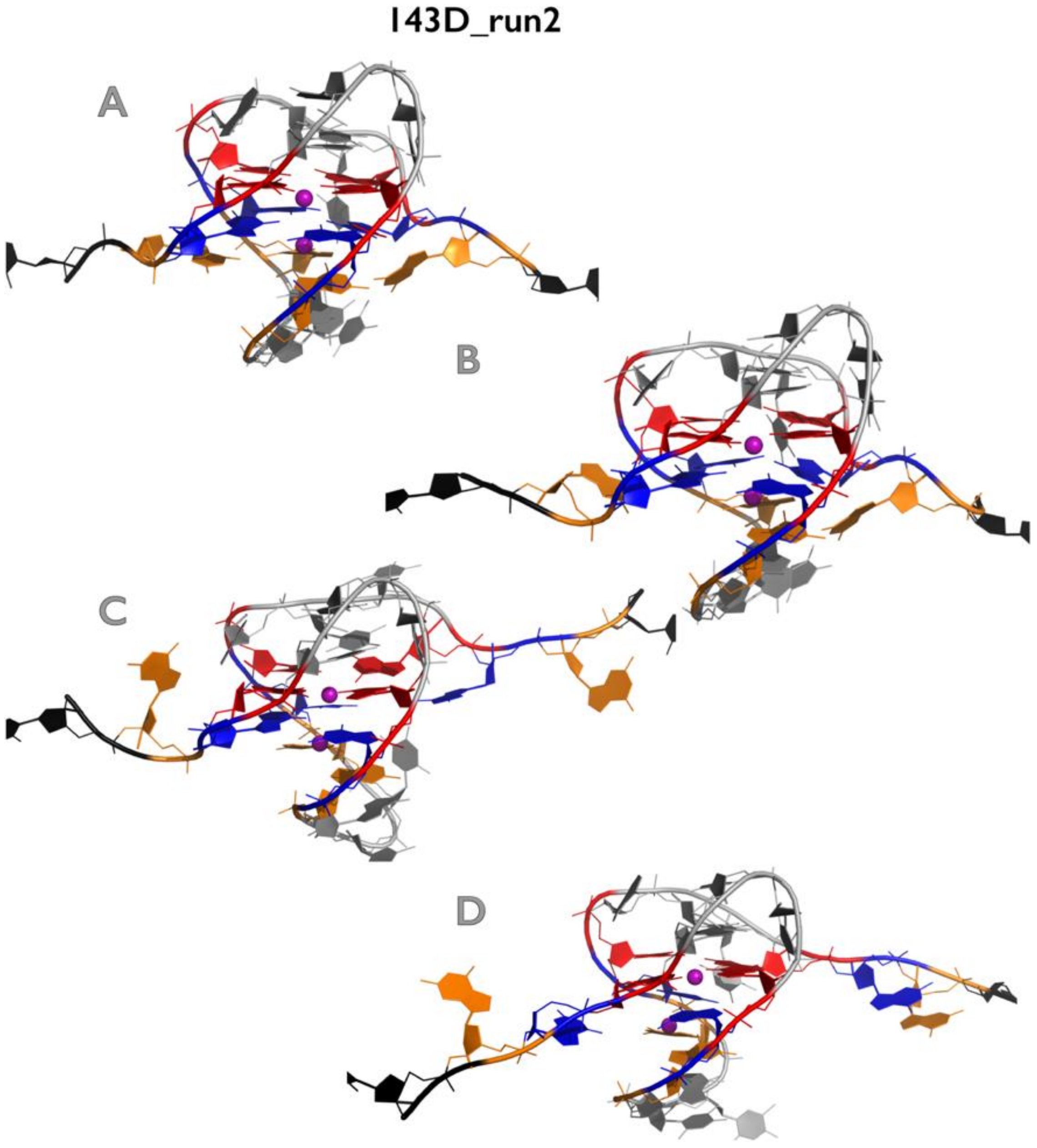

**Figure S27C:** Most important structural events during second independent *very slow pulling* simulation of 143D GQ system. See legend of Figure S1B for more details.

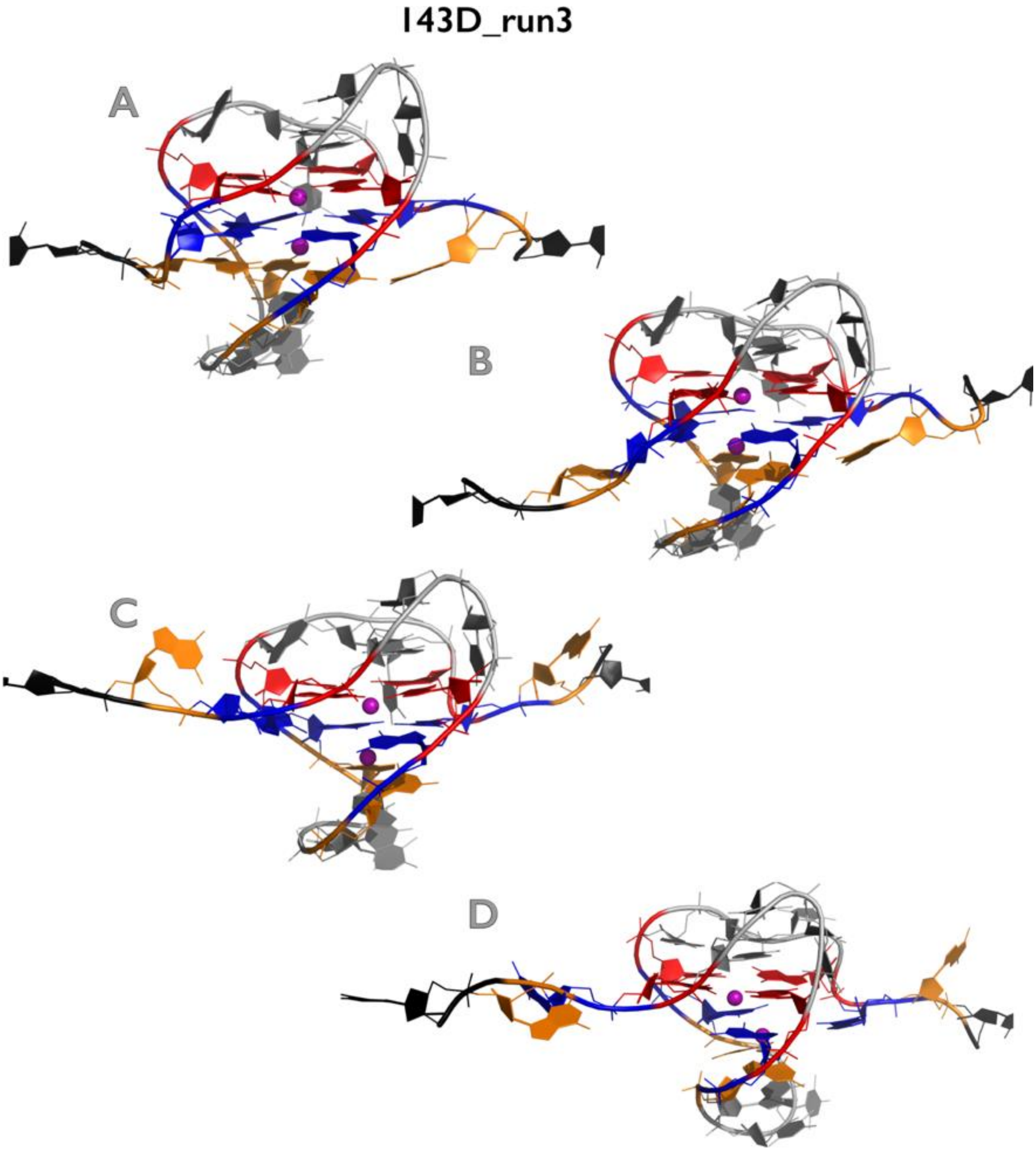

**Figure S27D:** Most important structural events during third independent *very slow pulling* simulation of 143D GQ system. See legend of Figure S1B for more details.

### REFERENCES

1. Cang, X.H., Sponer, J. and Cheatham, T.E. (2011) Explaining the Varied Glycosidic Conformational, G-Tract Length and Sequence Preferences for Anti-Parallel G-Quadruplexes. *Nucleic Acids Research*, **39**, 4499-4512.
2. Sponer, J., Mladek, A., Spackova, N., Cang, X.H., Cheatham, T.E. and Grimme, S. (2013) Relative Stability of Different DNA Guanine Quadruplex Stem Topologies Derived Using Large-Scale Quantum-Chemical Computations. *Journal of the American Chemical Society*, **135**, 9785-9796.
